# RNA-seq of Human T-Cells After Hematopoietic Stem Cell Transplantation Identifies Linc00402 as a Novel Regulator of T-Cell Alloimmunity

**DOI:** 10.1101/2020.04.16.045567

**Authors:** Daniel Peltier, Molly Radosevich, Guoqing Hou, Cynthia Zajac, Katherine Oravecz-Wilson, David Sokol, Israel Henig, Julia Wu, Stephanie Kim, Austin Taylor, Hideaki Fujiwara, Yaping Sun, Pavan Reddy

## Abstract

Mechanisms governing allogeneic T-cell responses after allogeneic hematopoietic stem cell (HSC) and solid organ transplantation are incompletely understood. Long non-coding RNAs (lncRNA) do not code for, but control gene expression with tissue specificity. However, their role in T-cell alloimmunity is unknown. We performed RNA-seq on donor T-cells from HSCT patients and found that increasing strength of allogeneic stimulation caused greater differential expression of lncRNAs. The differential expression was validated in an independent patient cohort, and also following ex vivo allogeneic stimulation of healthy human T-cells. Linc00402, a novel, conserved lncRNA, was identified as the most differentially expressed and was enriched 88 fold in human T-cells. Mechanistically, it was mainly located in the cytoplasm, and its expression was rapidly reduced following T-cell activation. Consistent with this, tacrolimus preserved the expression of Linc00402 following T-cell activation, and lower levels of Linc00402 were found in patients who subsequently went on to develop acute graft versus host disease (GVHD). The dysregulated expression of Linc00402 was also validated in murine T-cells, both in vitro and in vivo. Functional studies using multiple modalities to deplete Linc00402 in both mouse and human T-cells, demonstrated a critical role for Linc00402 in the T-cell proliferative response to an allogeneic stimulus but not a non-specific anti-CD3/CD28 stimulus. Thus, our studies identified Linc00402 as a novel, conserved regulator of allogeneic T-cell function. Because of its T-cell specific expression and its impact on allogeneic T-cell responses, targeting Linc00402 may improve outcomes after allogeneic HSC and solid organ transplantation.

**One sentence summary:** LncRNAs are differentially expressed by allogeneic antigen-stimulated T-cells, and the novel lncRNA, Linc00402, is a specific regulator of mouse and human allogeneic T-cells.

## INTRODUCTION

Allogeneic solid organ and hematopoietic stem cell transplantation are life-saving treatments for organ failure and high-risk malignancies, non-malignant hematologic disorders, and inherited metabolic disorders, respectively. Their efficacy is limited by T-cell mediated alloimmune reactions resulting in GVHD and allograft organ rejection, which are mitigated by non-specific immunosuppression that have significant side-effects. Better understanding of T-cell alloimmunity could lead to more specific and better tolerated therapies for mitigating the undesirable effects caused by allogeneic T-cells.

Non-coding RNAs (ncRNA) fine-tune gene expression and control cellular processes. Long non-coding RNAs (lncRNA) are a subset of ncRNAs that are 200 nucleotides or longer and are devoid of functional open reading frames (ORF) (*1, 2*). LncRNAs regulate gene expression in a variety of ways including regulating transcription (in cis or trans), translation, as competing endogenous RNAs, or by regulating the stability of specific mRNAs (*3, 4*). Importantly, in contrast to other ncRNA subsets such as micro-RNAs, lncRNAs possess tissue and context-specific expression (*5, 6*), which could make them attractive targets for anti-sense oligonucleotide based medicines.

The expression of lncRNAs in T-cells is both activation and subset-specific (*5–7*), yet their expression and role in T-cells following allogeneic HSCT remains unknown. In this report, we used unbiased RNA sequencing of donor T-cells from healthy controls and well-controlled patient samples following HSCT to investigate the role of lncRNAs in human allogeneic T-cells. We found that lncRNAs are differentially expressed by allogeneic T-cells and identified a novel lncRNA, Linc00402, as a specific regulator of both mouse and human allogeneic T-cells.

## RESULTS

### Increasing allo-stimulation causes differential expression of lncRNAs in human T-cells

To determine whether lncRNAs are differentially expressed by human alloimmune T-cells in a clinically relevant in vivo context, we performed RNA-seq of donor T-cells status-post clinical HSCT that received increasing degrees of allogeneic stimulation in stringently controlled hosts. Specifically, T-cells were isolated from 4 groups of well-controlled hematopoietic stem cell transplantation (HSCT) recipients, namely: 1) healthy controls; 2) autologous (Auto) recipients, where T-cells received proliferation signals due to lymphopenia; 3) matched unrelated donor recipients (MUD), where T-cells received proliferation signals due to minor histocompatibility driven allo-stimulation in the context of lymphopenia and immune suppression; and 4) mis-matched unrelated donor (MMUD) recipients, where T-cells received proliferation signals driven by both minor and major histocompatibility allo-stimulation in the context of lymphopenia and immune suppression (**Figure 1A**). All patients were controlled for absence of clinically significant acute GVHD, absence of infections, absence of corticosteroid therapy, type of immunosuppression (T-cell suppressive regimen with tacrolimus), similar myeloablative conditioning, stem cell source (PBMCs) without in vivo or ex vivo T-cell depletion, and time post HSCT (approximately 30 days post-HSCT) (**Table 1**). The transplants occurred from August 2006-October 2012. There were no major differences in age distribution or gender among the groups. The most common primary diagnosis in the Auto group was multiple myeloma, whereas for the MUD and MMUD groups the indication for transplant skewed more towards myeloid malignancies and myelodysplasia. Fludarabine and busulphan (Flu Bu4) were used as the preparative regimen for the MUD and MMUD patients. Tacrolimus with low dose methotrexate was the predominant GVHD prophylaxis used in both the MUD and MMUD groups. Percent CD3 donor chimerism on day +30 was not significantly different between the allogeneic groups (MUD mean 90%, MMUD mean 77%, *p* = 0.17). The MMUD patients were mismatched at HLA-DQ and HLA-A.

**Figure 1:**
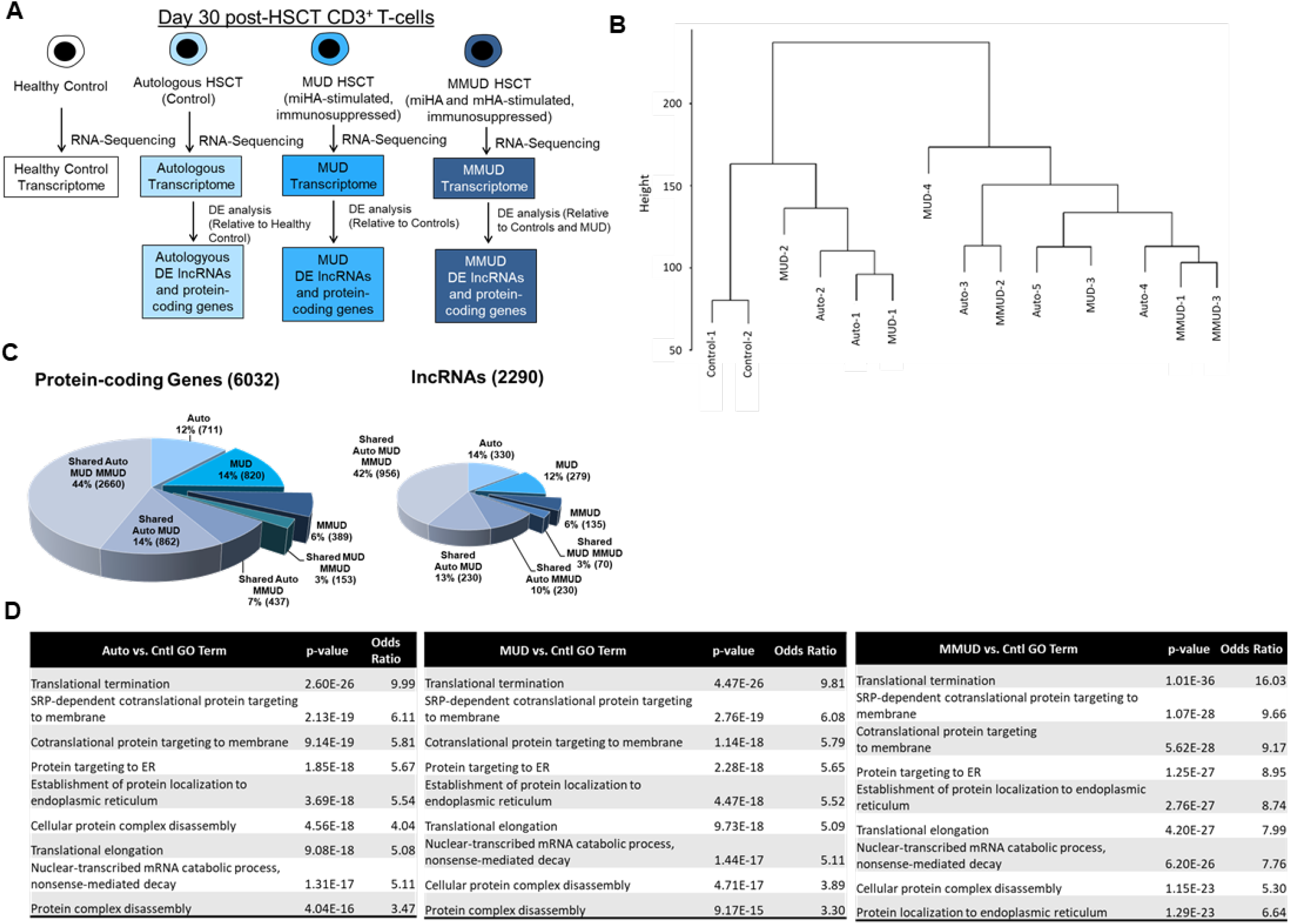
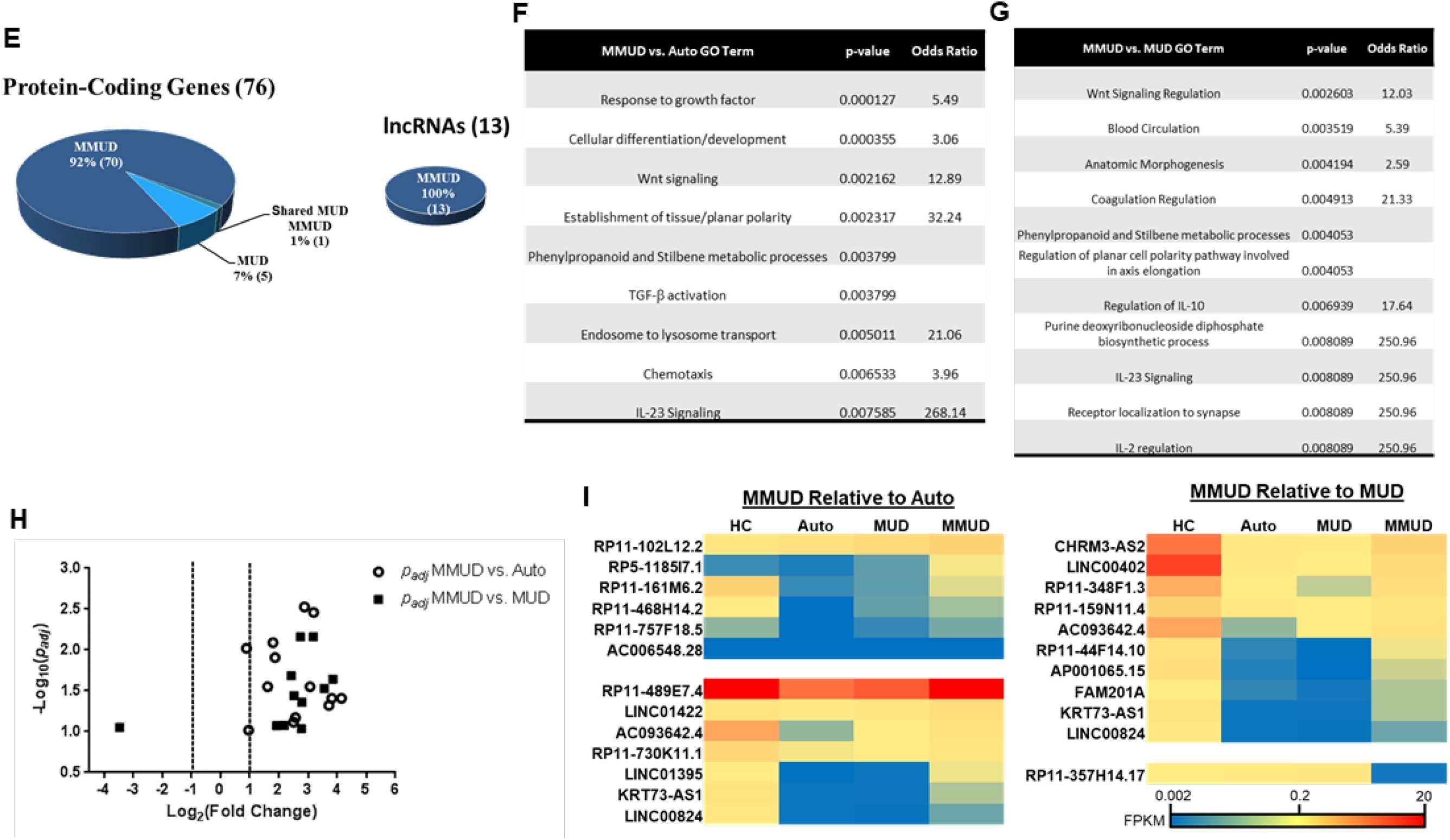
RNA-seq analysis post HSCT identifies differentially expressed lncRNAs in human allogeneic T cells. Total RNA sequencing was performed on healthy control CD3^+^ cells or CD3^+^ cells 30 days after either autologous, MUD, or MMUD HSCT. The experimental and analysis schema are shown in **A**. Hierarchical clustering analysis is shown in **B**. Differentially expressed (*p_adj_* < 0.1, *p* < 0.05, and fold change > 2) protein-coding (left pie charts) and lncRNA (right pie charts) genes (number of genes in parentheses) relative to healthy controls (**C**) or autologous HSCT controls (**E**) are shown. The top 10 enriched gene ontology terms for the differentially expressed protein-coding genes in the Auto, MUD, and MMUD groups relative to healthy controls is shown in **D**. A full list of enriched GO terms is show in **Supplemental Tables 1-3**. Summaries of the enriched gene ontology terms for the differentially expressed protein-coding genes in the MUD and MMUD groups relative to the Auto controls is shown in **F and G,** respectively. A full list of enriched GO terms is show in **Supplemental Tables 4-5**. A volcano plot of differentially expressed (*p* < 0.05, *p_adj_ <* 0.1) lncRNAs in the MMUD group relative to either autologous or MUD controls is shown in **H**. Heat maps showing differentially expressed lncRNAs in the MMUD group relative to autologous (left) or MUD (right) BMT patients (*p*<0.05, *p_adj_<*0.1) are shown in **I**. lncRNAs are grouped by expression pattern. Those underlined were differentially expressed relative to both Auto and MUD groups.

**Table 1.** RNA-seq patient characteristics.

|  | Transplant Group | Age | Sex | Primary Diagnosis | Conditioning | GVHD Prophylaxis | Stem cell Source | BMT Date | Day Post BMT | Donor CD3% Day 30 | HLA Mismatch |
| --- | --- | --- | --- | --- | --- | --- | --- | --- | --- | --- | --- |
| HC 1 | Healthy Control | 28 | Male | None |  |  |  |  |  |  |  |
| HC 2 | Healthy Control | 32 | Female | None |  |  |  |  |  |  |  |
| Auto 1 | Autologous | 52 | Female | Multiple Myeloma | Melphalan | None | PBSC | 2007 | 30 |  |  |
| Auto 2 | Autologous | 54 | Female | Multiple Myeloma | Melphalan | None | PBSC | 2006 | 30 |  |  |
| Auto 3 | Autologous | 68 | Female | Plasma Cell Leukemia | Melphalan | None | PBSC | 2006 | 32 |  |  |
| Auto 4 | Autologous | 47 | Male | Diffuse Large B-Cell Lymphoma | R-BEAM | None | PBSC | 2007 | 35 |  |  |
| Auto 5 | Autologous | 53 | Male | Mantle Cell Lymphoma | R-CVP | None | PBSC | 2007 | 26 |  |  |
| MUD 1 | Matched Unrelated | 56 | Male | Multiple Myeloma | Flu Bu4 | Tacro/MTX | PBSC | 2012 | 29 | 93 | None (10/10) |
| MUD 2 | Matched Unrelated | 55 | Male | CLL | R-Flu Bu4 | Tacro/MTX | PBSC | 2012 | 38 | 95 | None (8/8) |
| MUD 3 | Matched Unrelated | 67 | Female | MDS | Flu Bu4 | Tacro/MMF/Enbrel/ECP | PBSC | 2012 | 30 | 95 | None (10/10) |
| MUD4 | Matched Unrelated | 42 | Male | AML | Flu Bu4 | Tacro/MTX | PBSC | 2012 | 31 | 77 | None (8/8) |
| MMUD 1 | Mis-Matched Unrelated | 29 | Male | MDS | Flu Bu4 | Tacro/MTX/Enbrel | PBSC | 2009 | 30 | 91 | A (9/10) |
| MMUD 2 | Mis-Matched Unrelated | 62 | Female | Myelofibrosis | Flu Bu4 | Tacro/MTX | PBSC | 2012 | 34 | 65 | DQ (9/10) |
| MMUD 3 | Mis-Matched Unrelated | 43 | Male | AML | Flu Bu4 | Tacro/MTX | PBSC | 2012 | 26 | 75 | DQ (9/10) |

AML: acute myeloid leukemia. BEAM: Rituxan, carmustine, etoposide, cytarabine, melphalan. CLL: chronic lymphocytic leukemia. ECP: extracorporeal photopheresis. Flu Bu 4: fludarabine, busulphan. MDS: myelodysplastic syndrome. MMF: mycophenolic acid. MTX: methotrexate. PBSC: peripheral blood stem cells. R-R-CVP: Rituxan, cyclophosphamide, vincristine, prednisone. Tacro: tacrolimus.

To assess for global differences in differential protein-coding and lncRNA gene expression among the groups, we performed RNA-seq on purified CD3^+^ T-cells followed by unsupervised hierarchical clustering. The unsupervised clustering showed distinct grouping of the healthy controls, a relative skewing of the MMUD samples to that opposite of the healthy controls, and an intermingling of the autologous and MUD samples (**Figure 1B**). These data indicated that the transcriptomic profile of T-cells dramatically changes after any HSCT; however, among transplanted T-cells, those that received the greatest amount of allogeneic stimulation, the MMUD samples, differed the most. The most enriched gene ontology (GO) terms for differentially expressed protein-coding genes (*p_adj_* < 0.05) in the Auto, MUD, and MMUD samples relative to the healthy controls overlapped significantly and were largely related to transcriptional and translational regulation (**Figure 1D** and **Supplemental Tables 1-3**). Further, multiple terms pertaining to T-cell functions were enriched in all groups relative to healthy controls (**Supplemental Tables 1-3**), indicating proliferative and activated T-cells following any HSCT. Overall, these enriched GO terms were similar to those observed in a rhesus macaque GVHD model that used microarrays to profile the transcriptome of CD3^+^ T-cells post-HSCT (*8*).

Next, we assessed the number of differentially expressed protein-coding genes in the MUD and MMUD groups relative to the autologous (Auto) controls. Strikingly, 92% (70/76) of the differentially expressed protein coding genes were exclusively differentially expressed in the MMUD group relative to the Auto controls (**Figure 1E**). These data suggested that the protein-coding Auto and MUD transcriptomes were more similar and that increasing allogeneic stimulation, as in the MMUD group, enhanced differential expression of protein-coding transcripts.

To characterize the biologic properties associated with the differentially expressed protein-coding genes in the MMUD group, we performed GO analysis on differentially expressed protein-coding genes (*p_adj_* < 0.05) in the MMUD group relative to the MUD group or the Auto group. Because expression increased in the MMUD group for nearly all of the protein-coding genes (**Supplemental Figure 1**), we included any differentially expressed protein coding gene, regardless of its direction, in the GO analysis (**Figure 1F and G**, and **Supplemental Tables 4** and **5**). Several terms consistent with T-cell activation were enriched including IL-2 regulation, IL-23 signaling, and chemotaxis. Terms associated with immunologic regulation, such as IL-10 and TGF-β signaling, were also enriched. Extending previous data from macaque CD3^+^ T-cells post-allogeneic HSCT, we additionally observed the enrichment of GO terms for Wnt signaling, phenylpropanoid and stilbene metabolic processes in human T-cells (*8*).

Because lncRNA differential expression has never been assessed in allogeneic T-cells, we next examined the expression patterns of the differentially expressed lncRNAs among transplanted T-cells. We assessed the number of differentially expressed lncRNAs in the MUD and MMUD groups relative to the Auto controls. Similar to the protein coding genes, their expression generally increased with increasing allo-antigen stimulation (**Figures 1H and I**). Strikingly, 100% (13/13) of the differentially expressed lncRNAs were exclusively differentially expressed in the MMUD group relative to the Auto controls (**Figure 1E**). By contrast, just one lncRNA was decreased when we compared the MMUD and MUD groups (**Figure 1H**). However, no lncRNAs were differentially expressed in the Auto group compared to the MUD group. These data suggested that increasing allogeneic stimulation, as in the MMUD group (MHC disparate), influenced differential expression of lncRNAs when compared with the non-allogeneic-stimulated autologous or minor allogeneic antigen-stimulated (MUD) groups. Notably, fewer differentially expressed lncRNAs were identified than protein coding genes (**Figure 1H and I** vs **Supplemental Figure 1**) reflecting the lower amount of annotated lncRNAs (15,767) than protein coding genes (19,950) in Genecode 25, utilized as our reference annotation.

### Validation of lncRNA differential expression in an independent patient cohort

Next, to confirm the sequencing data, we performed qRT-PCR from the RNA that was used for sequencing. Eight of 11 (73%) tested lncRNAs with RNA-seq *p_adj_* < 0.1 were validated using qRT-PCR with ANOVA *p* < 0.1 (**Figure 2A**, **Supplemental Table 6**). Similar results were obtained for protein coding genes (**Supplemental Figure 2A-O, Supplemental Table 7**). Overall, these data further illustrate that increasing allo-antigen stimulation of human T-cells following HSCT predominantly increases both protein coding and long non-coding RNA expression.

**Figure 2:**
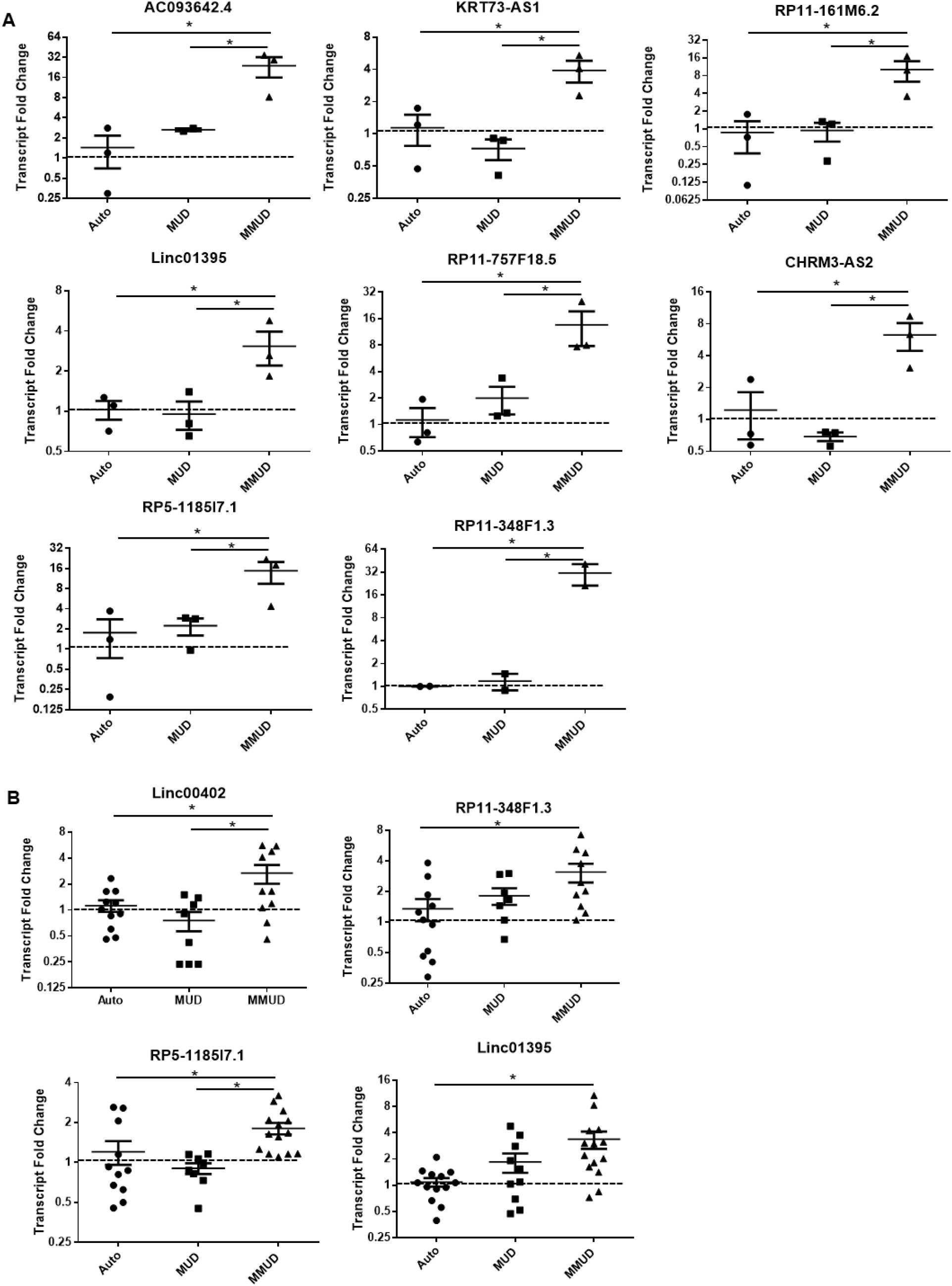
Confirmatory expression of lncRNAs in the RNA-seq patient cohort and their validation in an independent patient cohort. **A.** qRT-PCR was performed on total RNA derived from CD3^+^ peripheral blood T cells from patients detailed in Table 1 and the relative expression of the indicated transcripts was measured relative to β-actin. **B.** qRT-PCR was performed on cryogenically preserved CD3^+^ peripheral blood T cells from patients detailed in Table 2. The relative expression of the indicated transcripts is shown relative to β-actin. Error bars represent the mean +/− the SEM. \**p* < 0.05 by multiple comparison using original FDR method of Benjamini and Hochberg following a one-way ANOVA in which *p* <0.1 (See supplemental tables 6 and 8 for further details).

**Table 2.** Validation cohort characteristics.

|  | Transplant Group | Age | Sex | Primary Diagnosis | Conditioning | GVHD Prophylaxis | Stem cell Source | BMT Date | Day Post BMT | Donor CD3% Day 30 | HLA Mismatch |
| --- | --- | --- | --- | --- | --- | --- | --- | --- | --- | --- | --- |
| C-Auto 1 | Autologous | 51 | Female | MM | Mel |  | PBSC | 5/4/2006 | 36 |  |  |
| C-Auto 2 | Autologous | 63 | Male | MM | Mel |  | PBSC | 6/12/2008 | 35 |  |  |
| C-Auto 3 | Autologous | 50 | Female | MM | Mel |  | PBSC | 8/17/2006 | 34 |  |  |
| C-Auto 4 | Autologous | 52 | Male | MM | Mel |  | PBSC | 5/23/2008 | 28 |  |  |
| CAuto 5 | Autologous | 44 | Female | MM | Mel |  | PBSC | 2/14/2008 | 28 |  |  |
| C-Auto 6 | Autologous | 68 | Male | MM | Mel |  | PBSC | 1/5/2007 | 31 |  |  |
| C-Auto 7 | Autologous | 48 | Male | MM | Mel |  | PBSC | 5/4/2006 | 32 |  |  |
| C-Auto 8 | Autologous | 54 | Male | MM | Mel |  | PBSC | 2/29/2008 | 28 |  |  |
| C-Auto 9 | Autologous | 61 | Female | MM | Mel |  | PBSC | 2/28/2008 | 32 |  |  |
| C-Auto 10 | Autologous | 72 | Male | MM | Mel |  | PBSC | 1/13/2006 | 25 |  |  |
| C-Auto 11 | Autologous | 61 | Male | MM | Mel |  | PBSC | 11/9/2006 | 34 |  |  |
| C-Auto 12 | Autologous | 62 | Male | MM | Mel |  | PBSC | 3/20/2008 | 33 |  |  |
| C-Auto 12 | Autologous | 62 | Male | MM | Mel |  | PBSC | 3/20/2008 | 33 |  |  |
| C-Auto 13 | Autologous | 68 | Male | MM | Mel |  | PBSC | 2/15/2008 | 26 |  |  |
| C-Auto 14 | Autologous | 61 | Male | MM | Mel |  | PBSC | 4/18/2008 | 31 |  |  |
| C-MUD 1 | Matched Unrelated | 30 | Male | MDS-T | Flu Bu4 | Tacro/MTX | PBSC | 7/24/2012 | 35 | 76% | None (10/10) |
| C-MUD 2 | Matched Unrelated | 50 | Female | TR B-ALL | Flu Mel | Tacro/MMF | PBSC | 9/6/2012 | 34 | 95% | None (8/8) |
| C-MUD 3 | Matched Unrelated | 66 | Male | MDS | Flu Bu4 | Tacro/MTX | PBSC | 8/8/2012 | 36 | 90% | None (8/8) |
| C-MUD 4 | Matched Unrelated | 55 | Male | MF | Flu Bu4 | Tacor/MTX | PBSC | 10/18/2012 | 32 | 57% | None (10/10) |
| C-MUD 5 | Matched Unrelated | 53 | Male | MF | Flu Bu4 | Tacro/MTX | PBSC | 10/24/2012 | 33 | 30% | None (10/10) |
| C-MUD 6 | Matched Unrelated | 45 | Male | FL | R-Flu Bu4 | Tacro/MTX | BM | 10/11/2012 | 33 | 90% | None (10/10) |
| C-MUD 7 | Matched Unrelated | 29 | Female | T-ALL/HLH | Cy/TBI | Tacro/MTX | PBSC | 6/28/2012 | 31 | 84% | None (10/10) |
| C-MUD 8 | Matched Unrelated | 51 | Male | AML | Flu Bu4 | Tacro/MTX | PBSC | 4/3/2012 | 30 | 95% | None (10/10) |
| C-MUD 9 | Matched Unrelated | 58 | Male | AML | Flu Bu4 | Tacro/MTX | PBSC | 9/27/2012 | 39 | 64% | None (8/8) |
| C-MUD 10 | Matched Unrelated | 48 | Male | DLBCL | R-Flu Bu4 | Tacro/MTX | PBSC | 12/13/2011 | 27/30 | 95% | None (10/10) |
| C-MUD 11 | Matched Unrelated | 651 | Female | AML | Flu Bu4 | Tacro/MTX | PBSC | 8/21/2012 | 27 | 95% | None (10/10) |
| C-MMUD 1 | Mis-matched Unrelated Donor | 64 | Male | AML | Flu Bu4 | Tacro/MTX | PBSC | 11/24/2010 | 28 | 92% | A (7/8) |
| C-MMUD 2 | Mis-matched Unrelated Donor | 69 | Male | ALL | Flu Bu4 | Tacro/MTX | PBSC | 7/2/2013 | 34 | 81% | B (7/8) |
| C-MMUD 3 | Mis-matched Unrelated Donor | 52 | Male | ALL | Cy/TBI | Tacro/MTX | PBSC | 2/10/2012 | 32 | 91% | DQ (9/10) |
| C-MMUD 4 | Mis-matched Unrelated Donor | 63 | Female | ALL | Flu Bu4 | Tacro/MTX | PBSC | 1/22/2013 | 35 | 93% | A (9/10) |
| C-MMUD 5 | Mis-matched Unrelated Donor | 59 | Male | CTCL | FluMel | Tacro/MTX | PBSC | 3/6/2013 | 21 | 100% | C (9/10) |
| C-MMUD 6 | Mis-matched Unrelated Donor | 63 | Male | MDS | Flu Bu4 | Tacro/MTX/Vorinostat | PBSC | 11/18/2013 | 23 | 74% | DQ (9/10) |
| C-MMUD 7 | Mis-matched Unrelated Donor | 61 | Male | DLBCL | R-Flu Bu2 TBI | Tacro/MMF/Enbrel | PBSC | 9/16/2009 | 30 | 96% | C (9/10) |
| C-MMUD 8 | Mis-matched Unrelated Donor | 55 | Female | MCL | Flu Bu2 TBI | Tacro/MTX/Vorinostat | PBSC | 9/15/2009 | 38 | 76% | DRB1 (9/10) |
| C-MMUD 9 | Mis-matched Unrelated Donor | 52 | Female | AML | Flu Bu4 | Tacro/MTX | PBSC | 12/17/2013 | 30 | 51% | C (9/10) |
| C-MMUD 10 | Mis-matched Unrelated Donor | 54 | Male | AML | Flu Bu4 | Tacro/MTX | PBSC | 9/21/2010 | 35 | 95% | A (9/10) |
| C-MMUD 11 | Mis-matched Unrelated Donor | 56 | Female | AML | Flu Bu4 | Tacro/MMF/Enbrel/ECP | PBSC | 7/18/2012 | 29 | 95% | A (9/10) |
| C-MMUD 12 | Mis-matched Unrelated Donor | 10 | Female | HgSS | Campath/Flu/Mel | Tacro/MMF/Methylprednisone | BM | 2/1/2012 | 23/32 | 95% | DQ (9/10) |
| C-MMUD 13 | Mis-matched Unrelated Donor | 23 | Male | SAA | Thymo/Flu/Cy/TBI | Cyclosporin/MTX | BM | 8/23/2012 | 31/36 | 95% | C (7/8) |
| C-MMUD 14 | Mis-matched Unrelated Donor | 66 | Male | MDS | Flu Bu4 | Tacro/MMF/Enbrel/ECP | PBSC | 8/26/2010 | 25 | 72% | C (7/8) |
| C-MMUD 15 | Mis-matched Unrelated Donor | 61 | Male | DLBCL | R-Flu Bu4 TBI | Tacro/MMF | PBSC | 5/27/2010 | 32/35 | 94% | A (7/8) |
| C-MMUD 16 | Mis-matched Unrelated Donor | 61 | Male | AML | Flu Bu4 | Tacro/MTX | PBSC | 8/28/2012 | 29 | 67% | DRB1 (9/10) |
| C-MMUD 17 | Mis-matched Unrelated Donor | 66 | Female | AML | Flu Bu2 TBI | Tacro/MMF/Enbrel/ECP | PBSC | 10/30/2009 | 33 | 95% | A (9/10) |

AML: acute myeloid leukemia. BM: Bone marrow. CTCL: Cutaneous T-cell lymphoma. Cy: Cyclophosphamide. DLBCL: Diffuse large B-cell lymphoma. ECP: extracorporeal photopheresis. Flu Bu 4: fludarabine, busulphan. HgSS: Sickle cell disease. MCL: Mantle cell lymphoma. MDS: myelodysplastic syndrome. MDS-T: Therapy related myelodysplastic syndrome. Mel: Melphalan. MF: Myelofibrosis. MM: multiple myeloma. MMF: mycophenolic acid. MTX: methotrexate. PBSC: peripheral blood stem cells. R: Rituxan. SAA: Severe aplastic anemia. Tacro: tacrolimus. TBI: Total body irradiation. T-ALL/HLH: Treatment related T-cell acute lymphoblastic leukemia/hemophagocytic lymphohistiocytosis. TR B-ALL: Treatment related B-cell acute lymphoblastic leukemia. Thymo: Thymoglobulin.

To further validate and enhance the generalizability of lncRNA differential expression from the RNA-seq data, we performed qRT-PCR on T-cells isolated from an independent cohort of HSCT patients that was similar to the sequencing cohort patient characteristics (**Table 2**). To also appropriately and stringently control for sample quality, we also monitored the CD3^+^ percentage of live lymphocytes and the RNA integrity of each sample and observed no significant difference among the groups for either (**Supplemental Figures 3A** and **B**). We then validated the differential expression of 4 most differentially regulated lncRNAs from the RNA-seq data, such as Linc00402, RP11-348F1.3, RP5-1185I7.1, and Linc01395 (**Figures 2B**, and **Supplemental Table 8**). We additionally validated our RNA-seq test cohort for the differential expression of 7 protein-coding genes in the independent cohort (**Supplemental Figures 4A-G**, and **Supplemental Table 9**).

### Characterization and expression of the novel lncRNA, Linc00402, in human T-cells

To characterize further the tissue-specific expression of our identified lncRNAs, we queried the FANTOM-CAT database (*9*). Of the 4 lncRNAs that were differentially expressed and validated in the independent cohort, Linc00402 and RP11-348F1.3 were enriched in primary human T-cells 88.3 and 17.8 fold, respectively (**Figure 3A**). By contrast, RP5-1185I7.1 was not enriched in T-cells, and Linc01395 was not annotated in the FANATOM-CAT database (**Figure 3A**). Additionally, the FANTOM-CAT database, using 4 different published algorithms, predicted that Linc00402 and RP11-348F1.3 did not possess protein coding potential (**Figure 3B**). Given these results, we next characterized and analyzed the function of these novel T-cell specific lncRNAs that are differentially expressed in allogeneic human T-cells.

**Figure 3:**
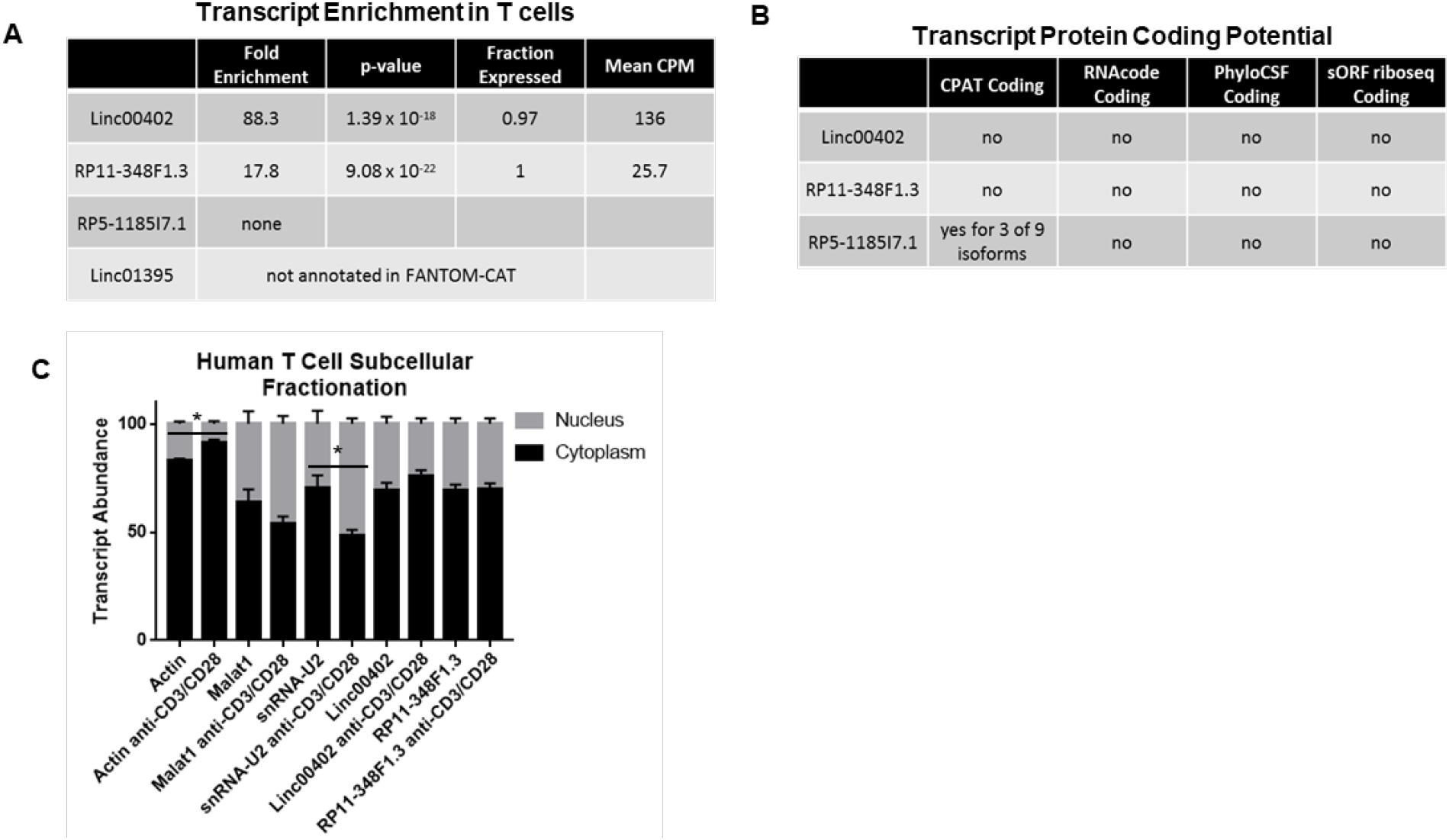
Linc00402 expression is enriched in human T-cells and is mainly localized to the cytoplasm. **A.** LncRNA fold enrichment in human T-cells was queried using the publically available FANTOM-CAT database. **B.** LncRNA coding potential was queried using multiple *in silico* algorithms using the publically available database FANTOM-CAT. **C.** qRT-PCR for the indicated transcripts was performed on naïve or activated human T-cell cytosolic and nuclear fractions. β-actin and snRNA U2 were included as positive cytosolic and nuclear controls, respectively (n=2-4 per group, \**p* < 0.05 using a two-tailed t-test).

Because subcellular localization of lncRNAs often correlates with their molecular function, we next determined the subcellular localization of the two T-cell specific lncRNAs, Linc00402 and RP11-348F1.3 (*10*). We fractionated unstimulated and anti-CD3/CD28 stimulated healthy human T-cells and extracted RNA for qRT-PCR. Actin and snRNA U2 were included as cytoplasmic and nuclear controls, respectively, both of which shifted more towards their expected subcellular compartments following anti-CD3/CD28 stimulation (**Figure 3C**). By contrast, the majority of Linc00402 and RP11-348F1.3 transcripts localized to the cytoplasm and this was unaffected by anti-CD3/CD28 stimulation (**Figure 3C**). These results are consistent with another T-cell lncRNA, Malat-1 (*6, 11*), and also in agreement with two independent in silico lnRNA subcellular localization prediction algorithms (*10, 12*).

### In vitro allogeneic stimulation differentially regulates Linc00402 expression

Because differential expression of lncRNAs correlated with the increasing strength of allogeneic stimulation in vivo, we next tested the hypothesis that allogeneic stimulation regulated Linc00402 expression in vitro using mixed lymphocyte reactions (MLR). Negatively selected T-cells from healthy donors were mixed with either positively selected, lethally-irradiated (30 Gy) autologous monocytes or a pool of 2-3 unrelated (i.e. allogeneic) donor peripheral blood monocytes. In addition, T-cells were also incubated with anti-CD3/CD28-coated beads as a positive control for T-cell activation. Following incubation, the T-cells were separated from the remaining irradiated monocytes via negative selection and analyzed for differential expression of lncRNAs via qRT-PCR. As controls for both up-regulated and down-regulated genes, we included the confirmed protein coding gene SV2A and the well-characterized lncRNA MIAT, respectively. As expected, SV2A’s expression increased following allo-stimulation or T-cell activation and MIAT’s expression decreased (**Figure 4A**). We then confirmed the differential expression of Linc00402, RP11-348F1.3, RP5-1185I7.1, and Linc01395 whose expression increased following in vivo allogeneic stimulation from both our sequencing and confirmatory patient cohorts. Surprisingly, and in contrast to in vivo allogeneic stimulation following clinical HSCT, in vitro allogeneic stimulation resulted in decreased expression of all 4 of these lncRNAs. The in vitro down regulation of Linc00402 and RP11-348F1.3 also occurred rapidly following anti-CD3/CD28 stimulation (**Figures 4B**), suggesting that their expression was directly regulated by T-cell activation. Thus, T-cell differential expression of Linc00402 and RP11-348F1.3 was opposite in vivo versus in vitro. We, therefore, next explored the putative reason for this difference.

**Figure 4:**
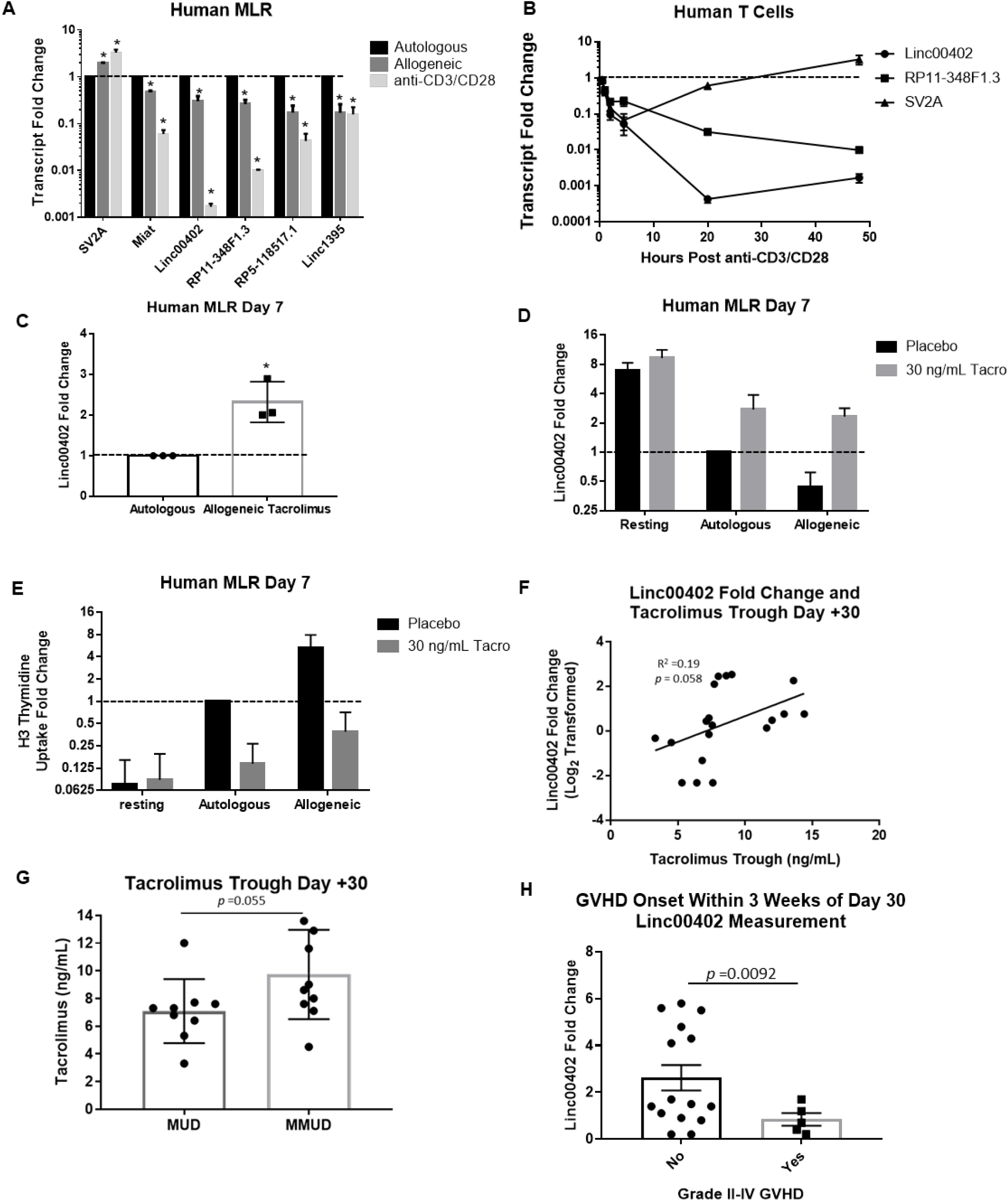
Linc00402 is differentially expressed in ex vivo allo-stimulated human T-cells and its expression is effected by tacrolimus-mediated inhibition of T-cell activation. Human T-cells were incubated with irradiated (30 Gy) autologous or allogeneic monocytes for 6 days or anti-CD3/CD28 Dynabeads for 2 days. The T-cells were then re-isolated via negative magnetic selection and subject to qRT-PCR (**A**) (\**p* < 0.05, one-way ANOVA, original FDR method of Benjamini and Hochberg, n=3-6 per group). **B.** Human T-cells were activated with anti-CD3/CD28 Dynabeads for the indicated times and then the expression of the indicated transcripts was assessed via qRT-PCR (n=2-4). Error bars represent the mean +/− the SEM. Human T-cells were pre-incubated with tacrolimus (30 ng/mL) or placebo for 1 hour prior to mixing them with irradiated (30 Gy) autologous or allogeneic monocytes or placebo. Seven days later, T-cells were negatively magnetically isolated and subjected to qRT-PCR analysis of Linc00402 with fold changes set relative to autologous placebo-treated controls (**C-D**) (n = 2-3 per condition, \**p*<0.05). **E.** Proliferation was assessed by uptake of tritiated thymidine relative to autologous placebo control uptake (n = 3 per condition, \**p*<0.05). **F.** The correlation between Linc00402 expression (Log_2_ fold change from autologous controls) in MUD and MMUD patient T-cells described in figure 2B and their day plus 30 tacrolimus trough level. **G.** Average day plus 30 tacrolimus trough levels from MUD and MMUD patients described in figure 2B (error bars represent the mean +/− SEM). **H.** Average day plus 30 Linc00402 expression (fold change from autologous controls) in T-cells described in Figure 2B for those patients who did and did not develop grade II-IV GVHD within 3 weeks of their day plus 30 Linc00402 expression measurement (Welch’s *t*-test was used to calculate the *p*-value).

### Inhibition of T-cell activation by tacrolimus alters Linc00402 expression

A key difference between the in vivo allogeneic T-cell patient samples (MMUD and MUD samples) and the in vitro allogeneic T-cells is that the patient samples were exposed to tacrolimus for GVHD prophylaxis. Tacrolimus is a calcineurin inhibitor that blocks T-cell activation by inhibiting the nuclear translocation of nuclear factor of activated T-cells (NFAT) (*13*). Therefore, we hypothesized that its presence may affect the expression of Linc00402 in T-cells. To test this, we performed human in vitro MLRs in the absence and presence of tacrolimus. Consistent with this hypothesis, tacrolimus-treated allo-stimulated T-cells expressed more Linc00402 than placebo-treated autologous-stimulated T-cells (**Figure 4C**), analogous to the in vivo results between the MMUD and autologous patient groups (**Figure 2B**). In fact, resting, placebo-treated T-cells expressed the greatest amount of Linc00402 whose expression then declined with increasing amounts of stimulation provided by irradiated autologous, or allogeneic monocytes (**Figures 4D**). In agreement with tacrolimus influencing the expression of Linc00402 by modulating T-cell activation, T-cell proliferation inversely correlated with Linc00402 expression in MLRs (**Figure 4E**). As for Linc00402, tacrolimus also preserved the expression of RP11-348F1.3 in allogeneic T-cells relative to placebo-treated autologous-stimulated human T-cells (**Supplemental Figures 5A and B**). Consistent with tacrolimus limiting T-cell activation and thereby preserving Linc00402 expression, there was a trend towards a positive correlation between day +30 tacrolimus troughs and Linc00402 levels in MUD and MMUD patients (**Figure 4F**).

These data suggested that inhibition of T-cell activation by tacrolimus resulted in higher levels of Linc00402 expression in allogeneic antigen-stimulated T-cells both in vivo (MMUD patients) and in vitro (MLR). However, this still did not account for the lack of increased Linc00402 expression in the MUD patient samples, which were also exposed to tacrolimus and received an allogeneic stimulation, albeit a milder allogeneic stimulus than the MMUD samples. To address this conundrum, we reasoned that physicians kept tacrolimus levels higher in MMUD patients due to their increased risk of GVHD relative to MUD patients (*14*). To test this, we analyzed whether higher tacrolimus levels in the MMUD patients would limit T-cell activation and may account for the higher Linc00402 expression in MMUD patient T-cells relative to MUD patient T-cells. Consistent with this notion, we observed a trend towards higher day +30 tacrolimus troughs in the MMUD patients relative to the MUD patients (**Figure 4G**).

These data collectively suggested that Linc00402 expression was inversely proportional to the activation status of T-cells. We further correlated the clinical consequence of this by determining whether Linc00402 expression inversely correlated with the development of clinically significant acute GVHD (grade II-IV) within 3 weeks of the day +30 Linc00402 expression value measured in MUD and MMUD patients. We restricted the analysis to acute GVHD onset within the subsequent 3 weeks after the day +30 Linc00402 expression measurement because low calcineurin inhibitor troughs were previously shown to increase the risk of developing acute GVHD in the following week (*15*). Consistent with this, we observed a significantly higher Linc00402 expression in those who did not go on to develop GVHD in the subsequent 3 weeks (**Figure 4H**). Collectively, these resulted showed that tacrolimus impacts Linc00402 expression. Nonetheless, given the complexity of the in vivo clinical context and limitations imposed by multiple variables and patient numbers, we next sought to determine whether Linc00402 expression was truly biologically relevant by analyzing its role in murine T-cells.

### Linc00402 is conserved in mouse T-cells and is regulated by T-cell activation both in vitro and in vivo

Sequence conservation is generally considered a hallmark of functionally important genes; however, only about 40-70% of lncRNAs are conserved between humans and mice (*9, 16, 17*). To assess for murine orthologues, we queried the FANTOM-CAT database, which indicated Linc00402’s transcription initiation region and exons were conserved (**Figure 5A**), and an orthologous region on mouse chromosome 14 was identified using TransMap Ensembl Mappings Version 4. In contrast, RP11-348F1.3 was not conserved. This orthologous region maintained synteny with the transcription factor Kruppel like factor 12 (KLF12) (**Figure 5B**); maintenance of synteny and sequence conservation are characteristics associated with functional lncRNAs (*18*). Therefore, we next assessed for the expression of these transcripts in mouse T-cells. Primers were designed for highly conserved regions of each (marked by arrows in Figure 5B) and used to conduct qRT-PCR on mouse T-cells. A significant signal above a non-template control was obtained for Linc00402, but not for RP11-348F1.3 (**Figure 5C**). Linc00402 expression in murine T-cells was further confirmed via northern blotting using an RNA probe (**Figure 5D**). A 2.5 kb band, which approximates the 3 kb human transcript, was present in murine resting T-cells but not in negative-control human HeLa cells. Furthermore, this 2.5 kb band was down-regulated following anti-CD3/CD28 Dynabead stimulation (**Figure 5D**). These data demonstrated for the first time that Linc00402, a novel lncRNA, is present and conserved in T-cells from both humans and mice.

**Figure 5:**
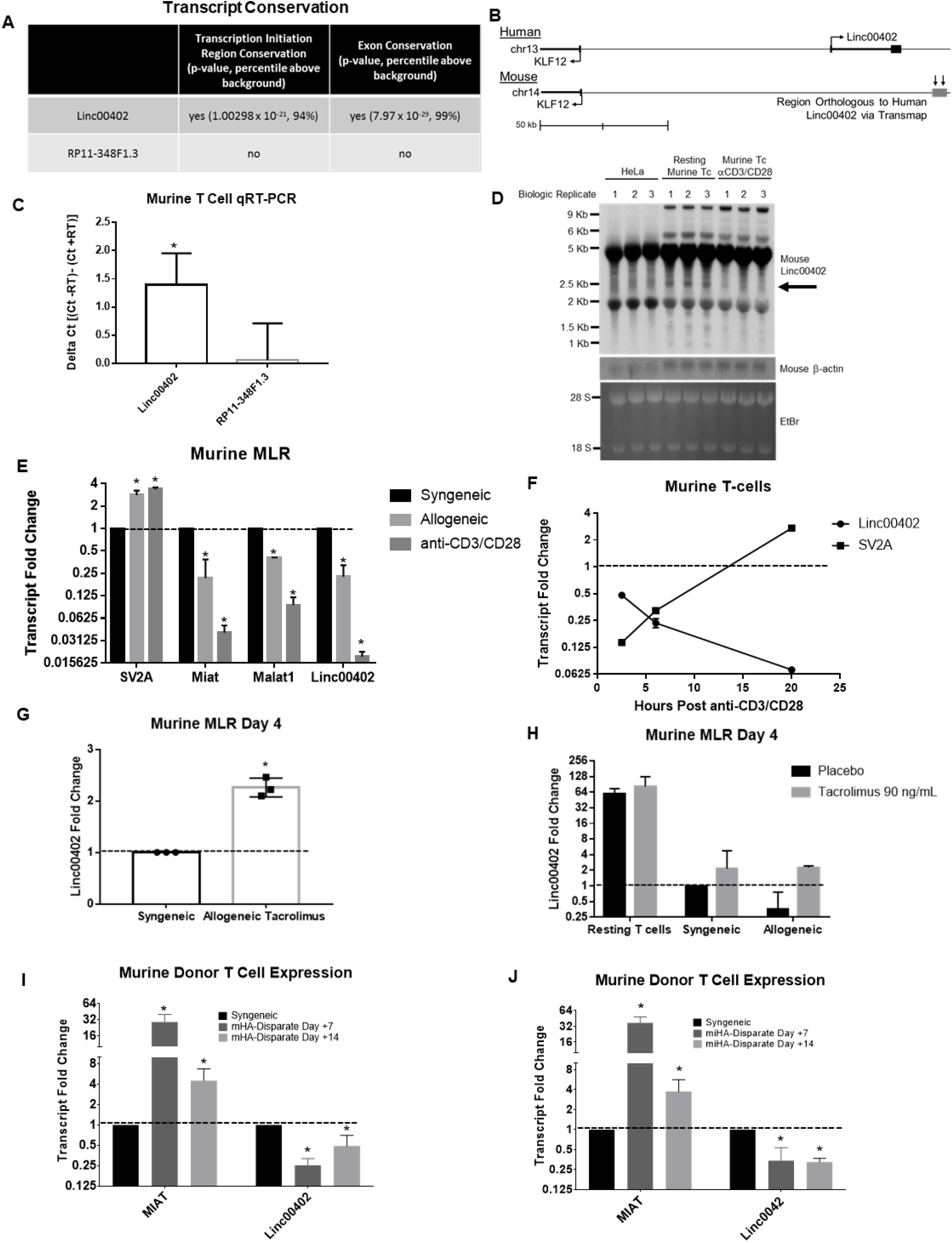
Linc00402 is differentially expressed in murine T-cells. LncRNA conservation was queried via the FANTCOM-CAT database (**A**). **B.** Depiction of the human and mouse Linc00402 locus on chromosome 13 and 14, respectively. **C.** qRT-PCR was performed on murine T-cells using primers for regions predicted by the Ensembl genome browser to be orthologous to the indicated human lncRNA exonic region. Data depict the subtraction of qRT-PCR threshold cycles from threshold cycles obtained from negative reverse transcriptase controls (\**p* < 0.05 using a two-tailed t-test). **D.** Ten μg of total RNA from resting mouse T-cells or mouse T-cells stimulated for 48 hours with anti-CD3/CD28 Dyna beads was analyzed via northern blot using probes specific for murine Linc00402 and β-actin. Total RNA from HeLa cells was run as a negative control. Triplicate biologic replicates are shown. The arrow points to a 2.5 kb band that is similar in size to human Linc00402, absent in HeLa cells, and decreases in mouse T-cells following anti-CD3/CD28 stimulation. **E.** C57Bl/6 splenic T-cells were incubated with irradiated (30 Gy) syngeneic or allogeneic (BALB/c) splenocytes or anti-CD3/CD28 Dynabeads. T-cells were then re-isolated and subjected to qRT-PCR (\**p* < 0.05, one-way ANOVA, original FDR method of Benjamini and Hochberg, n=2-7). **F.** Mouse T-cells were activated with anti-CD3/CD28 Dynabeads for the indicated times and then the expression of the indicated transcripts was assessed via qRT-PCR (n=2-4). Error bars represent the mean +/− the SEM. **G-H.** Mouse T-cells were pre-incubated with tacrolimus (90 ng/mL) or placebo for 1 hour prior to mixing them with irradiated (30 Gy) syngeneic or allogeneic splenic monocytes or placebo. Seven days later, T-cells were negatively magnetically isolated and subjected to qRT-PCR analysis of Linc00402 with fold changes set relative to syngeneic placebo-treated controls (n = 3 per condition, \**p*<0.05). **I-J.** The differential expression of selected transcripts from allogeneic splenic donor T-cells was assessed via qRT-PCR on days +7 and +14 from mHA-disparate (**I**) and multiple miHA-disparate (**J**) recipients (\**p* < 0.05, one-way ANOVA, original FDR method of Benjamini and Hochberg, n=3-4 per group). Error bars represent the mean +/− the SEM.

We next assessed whether Linc00402 in murine T-cells was also regulated by allo-stimulation as in human T-cells. We first tested this in vitro with murine MLRs similar to that described for the human MLRs except that T-cells and irradiated monocytes were derived from mouse spleens. As in the human MLRs, the control protein coding transcript, SV2A, showed increased expression upon allo-stimulation or non-specific T-cell activation and the two well-characterized lncRNAs Miat and Malat1 were down-regulated in response to an allogeneic or anti-CD3/CD28 stimulus (**Figure 5E**). Similar to that observed in the human MLR, murine Linc00402 expression decreased with allo-stimulation and T-cell activation (**Figure 5E**). Furthermore, just as with human Linc00402, murine Linc00402 expression in T-cells rapidly declined following stimulation with anti-CD3/CD28 suggesting that its expression is regulated by T-cell activation (**Figure 5F**).

We further explored whether incubation of murine allogeneic T-cells with tacrolimus also affected Linc00402 expression as in human allogeneic T-cell. Consistently, tacrolimus inhibited murine T-cell activation thereby causing the expression of murine Linc00402 to be higher in mouse T-cells treated with tacrolimus and stimulated with allogeneic monocytes relative to the expression observed in placebo-treated mouse T-cells stimulated with syngeneic monocytes (**Figures 5G-H**). These data demonstrated that the in vitro regulation of Linc00402 expression in allogeneic T-cells was conserved in humans and mice.

Next, we hypothesized that Linc00402 would be similarly regulated in allogeneic T-cells following experimental murine in vivo allogenic HSCT. To determine this, we measured the expression of Linc00402 in donor T-cells at 7 and 14 days post HSCT in two distinct murine models of GVHD (B6γBalb/c and B6γC3H.SW). Specifically, the expression of murine Linc00402 decreased 2.5 to 13 fold (*p* < 0.01) depending on the time point and model (**Figure 5I and J**). The decreased expression of murine Linc00402 in allo-antigen stimulated T-cells was consistent with the lack of tacrolimus in these models and demonstrated that Linc00402 expression is down-regulated upon murine T-cell activation both in vitro and in vivo, similar to human T-cells.

### Anti-sense-mediated knock-down of Linc00402 impairs human and murine allogeneic T-cell proliferation

Having established the differential expression of Linc00402 in human and mouse allogeneic T-cells both in vitro and in vivo, we next determined the functional relevance of Linc00402 in T-cells. To determine the function of Linc00402, we knocked it down utilizing multiple approaches. First, we utilized GapmeR anti-sense oligonucleotides because previous work had shown their efficiency in knocking down transcripts in primary T-cells both in vitro and in vivo (*19, 20*). To assess the efficiency of GapmeR uptake in resting T-cells, varying concentrations of a non-targeting fluorescently-labeled GapmeR were incubated with human or mouse T-cells for 72 or 48 hours, respectively (**Figure 6A**). Cryogenically preserved peripheral blood human T-cells and freshly isolated mouse splenic T-cells efficiently took up the GapmeRs within 72 and 48 hours, respectively (**Figure 6A**). Next, we designed GapmeRs targeting human and mouse Linc00402 and, also for human RP11-348F1.3 as a control. The GapmeRs did not cause any significant cytotoxicity (**Supplemental Figures 6A and B**). We then confirmed the ability of these GapmeRs to reduce the expression of their target transcripts in human and mouse T-cells relative to a non-targeting control (**Figures 6B and D**) and then tested their impact on T-cell proliferation in MLRs. The knock-down of Linc00402 in both human and murine allogeneic T-cell inhibited their proliferation in response to an allogeneic stimulus, but had no effect on proliferation in response to an anti-CD3/CD28 stimulus (**Figures 6C and E**). Knock-down of the previously characterized, ubiquitously expressed, pro-proliferative, lncRNA, murine Malat-1, inhibited T-cell proliferation following both allogeneic and anti-CD3/CD28 stimulations (**Figure 6E**) (*21*). Similar to Linc00402, knock-down of RP11-348F1.3, inhibited human allogeneic T-cell proliferation but had no effect on T-cell proliferation following anti-CD3/CD28 stimulation (**Figure 6C**). Because there is no mouse orthologue for RP11-348F1.3, we could not test its influence on murine allogeneic T-cell proliferation. Together, these data demonstrated the conserved pro-proliferative function of the novel lncRNA, Linc00402, following an allogeneic stimulus in both human and murine T-cells.

**Figure 6:**
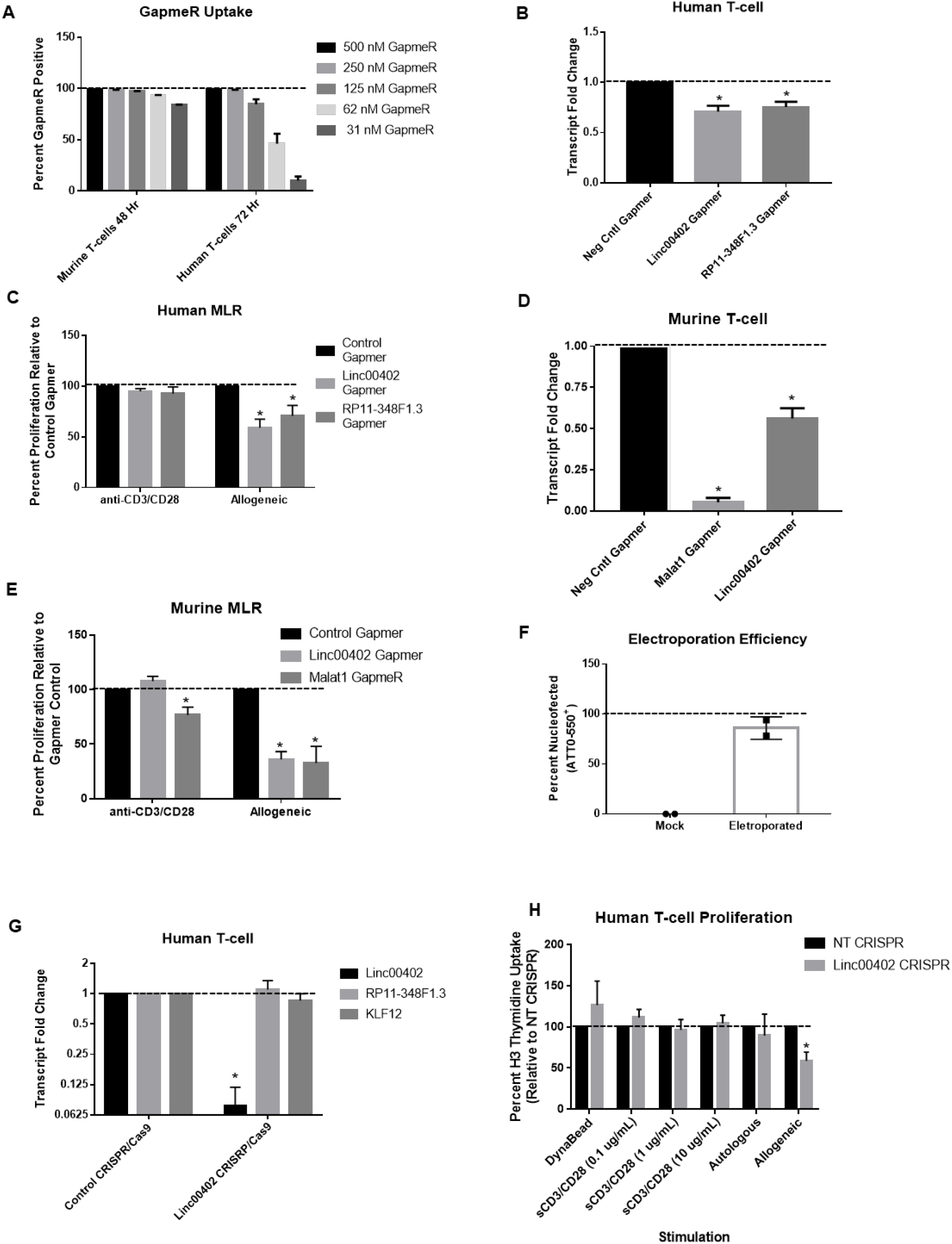
Depletion of Linc00402 and its predicted murine orthologue in T-cells inhibits allogeneic T-cell proliferation. Human and mouse T-cells were incubated with varying concentrations of a 5’ FAM-labeled non-targeting GapmeR for 72 or 48 hours, respectively. The percentage of FAM^+^ T-cells was assessed via flow cytometry (**A**) (n=2 per group). **B** and **D** Human and mouse T-cells were incubated with 500 or 250 nM of the indicated GapmeRs for either 72 or 48 hours, respectively. qRT-PCR was then performed for the indicated transcripts (\**p* < 0.05, one-way ANOVA, original FDR method of Benjamini and Hochberg, n= 4-7 per group). Human (**C**) or mouse (**E**) T-cells were incubated with the indicated GapmeRs as in B and D and then MLRs were performed as described in figures 3 and 4, respectively. T-cell proliferation was then assessed by 3H thymidine incorporation (\**p* < 0.05, one-way ANOVA, original FDR method of Benjamini and Hochberg, n=3-7 per group). Error bars represent the mean +/− the SEM. **F.** Human T-cells were electroporated with fluorescently-labeled CRISPR-Cas9 RNPs. Electroporation efficiency was assessed 24 hours later via flow cytometry relative to mock electroporated controls. **G.** The expression of Linc00402 and KLF12 were assessed via qRT-PCR in human T-cells 72 hours after electroporation of Linc00402-targeting or non-targeting control CRISPR-Cas9 RNPs (n=3-4 per group, \**p* < 0.05). **H.** Human T-cells were electroporated with non-targeting control or Linc00402-targeting CRISPR-Cas9 RNPs and then stimulated with Dynabeads or varying concentrations of soluble anti-CD3/CD28 antibodies for 48 hours. Alternatively, electroporated cells were stimulated with irradiated autologous or allogeneic monocytes and incubated for 14 days. Proliferation was measured via tritiated thymidine incorporation (\**p* < 0.05, one-way ANOVA, original FDR method of Benjamini and Hochberg, n=2-3 per group).

### CRISPR-Cas9-mediated genomic deletion of Linc00402 impairs allogeneic T-cell proliferation

Next, we confirmed our GapmeR knock-down results by deleting Linc00402 in human T-cells via nucleoporation of CRISPR-Cas9 ribonucleoprotein complex (RNP) pools (*22*). Using this approach, we regularly achieved high nucleoporation efficiencies (**Figure 6F**) and robust depletion of Linc00402 without effecting the expression of its neighboring gene KLF12 (**Figure 6G**). CRISPR-Cas9-mediated depletion of Linc00402 inhibited the ability of T-cells to proliferate in response to an allogeneic stimulus, but had no effect on their ability to proliferate following an anti-CD3/CD28 Dynabead stimulation or various concentrations of soluble anti-CD3/CD28 antibodies (**Figure 6H**). These data demonstrated that lncRNAs, specifically the conserved lncRNA, Linc00402, regulated only allogeneic T-cell proliferation in mice and humans.

## DISCUSSION

The mechanisms governing human allogeneic T-cell-mediated toxicities remain incompletely understood, which limits the wider application of allogeneic HSC and solid organ transplantation. Here we identified a novel lncRNA, Linc00402, that regulates allogeneic T-cells. Specifically, we found that transplanted T-cells altered the expression of thousands of lncRNAs relative to healthy control T-cells. Among transplanted T-cells, however, only those that received the greatest allogeneic stimulation demonstrated the largest amount of differentially expressed lncRNAs compared to T-cells proliferating in lymphopenic hosts or allogeneic T-cells responding to minor histocompatibility antigens. These results were confirmed in an independent patient cohort. Several of these lncRNAs were also dysregulated in both human and murine MLRs as well as in multiple murine models of HSCT. Because of its specific expression in T-cells, its sequence conservation, and its maintained synteny, we focused on Linc00402, and validated its functional relevance in both human and murine T-cells.

LncRNAs possess remarkable tissue, species and context-specific expression relative to mRNAs (*5, 16, 17, 23-25*). Consistently, we observed mainly increased Linc00402 expression with increasing allo-stimulation of T-cells from patients 30 days after HSCT. The dysregulation of Linc00402 in response to an allogeneic stimulus remained consistent in ex vivo human and murine T-cells as well as in vivo murine T-cells following multiple models of experimental HSCT. We made the interesting observation that the in vivo differential expression of Linc00402 in allo-stimulated T-cells exposed to immunosuppression had the opposite effect on its expression to that observed in the absence of immunosuppression. We demonstrated that these differences were due to tacrolimus exposure in the allogeneic patient T-cells, which inhibited T-cell activation leading to greater expression of Linc00402. We further demonstrated that the expression level of Linc00402 was directly proportional to the trough concentration of tacrolimus in patients. This was consistent with higher Linc00402 expression in MMUD patient T-cells where tacrolimus levels were kept higher than in MUD patients. The correlation between calcineurin inhibitor concentration and Linc00402 expression was likely related to suppression of T-cell activation, which will need to be examined further in future studies.

T-cells are required for the development of GVHD, and the agents that form the backbone of GVHD prophylaxis inhibit T-cell activation (*26*). Previous studies demonstrated that sub-optimal inhibition of T-cell activation increased the risk of subsequent GVHD (*15, 27, 28*). Here we showed that T-cell activation down-regulated Linc00402 expression; therefore, we wondered if low Linc00402 expression was associated with the subsequent development of acute GVHD. While acknowledging our small sample size and the retrospective nature of this analysis, we observed significantly lower Linc00402 expression in those patients who then went on to develop grade II-IV acute GVHD in the next 3 weeks. These studies will need to be repeated in a prospective manner with larger sample sizes, but nonetheless, they highlight the potential of lncRNAs as biomarkers for important outcomes post-HSCT. However, given its conservation and expression in murine and human T-cells, its differential expression in response to allogeneic stimulation both in vitro and in vivo, and its influence on T-cell proliferation following allogeneic antigen stimulation, our data demonstrate that Linc00402 is a novel regulator of T-cell alloimmunity. Furthermore, solid organ allogeneic transplantation is often performed across MHC mismatches (*29*), targeting Linc00402 may have significance for solid organ allogeneic transplantation.

The molecular mechanisms mediating lncRNA regulation of gene expression vary (*4*). In general, lncRNAs that primarily localize to the nucleus regulate transcription, whereas those that primarily localize to the cytoplasm regulate gene expression via a variety of post-transcriptional mechanisms (*4*). Here we found that Linc00402 localized primarily to the cytoplasm in human T-cells. Future work will seek to characterize their molecular interactions in order to understand their molecular mechanisms of action.

In addition to providing evidence that lncRNAs regulate allogeneic T-cells, our RNA-seq data also established the first unbiased human T-cell transcriptome post-HSCT, which will likely prove to be a valuable hypothesis-generating resource. For instance, in addition to differentially expressed lncRNAs, we also found many interesting and novel differentially expressed protein-coding genes whose further study might provide useful insights into allogeneic T-cell biology.

To our knowledge, the role lncRNAs play on allogeneic T-cell responses following HSCT was previously uncharacterized. However, several observations supported the hypothesis that lncRNAs influenced allogeneic T-cell function, including that lncRNAs are transcribed by T-cell subtype-specific transcription factors (*5*), they regulate T-cell functions critical for alloimmunity including cytokine expression and migration (*5-7, 30, 31*), they are associated with solid organ allograft rejection (*32–34*), they regulate Th1 effector responses in the setting of cardiac allograft rejection (*33*), and they favor murine tolerogenic dendritic cells following allogeneic cardiac transplantation (*11*). Here we showed that lncRNAs, such as Linc00402, are differentially expressed by allogeneic T-cells and regulate their function. Importantly, the tissue and context specific expression of lncRNAs, such as Linc00402, may make them potentially attractive therapeutic targets for improving outcomes after allogeneic HSC and solid organ transplantation.

## MATERIALS AND METHODS

### Human T-cell isolation

Human peripheral blood was obtained via routine venipuncture according to IRB protocol UMCC 2001.0234 (HUM00043287), Long-Term Evaluation of the Biology and Outcomes of Hematopoietic Stem Cell Transplantation, which collects clinical data and blood samples serially throughout the transplant course as well as at the time of significant clinical complications, such as GVHD, at our center. Alternatively, heparinized whole blood was purchased from Innovative Research. Peripheral blood mononuclear cells (PBMC) were purified from heparinized whole blood via density centrifugation with Ficoll-Paque Premium (GE Healthcare) per manufacturer’s recommendations. PBMCs were then treated with red blood cell (RBC) lysis buffer (Sigma) per manufacturer’s instructions, and cryopreserved in heat-shocked fetal calf serum (Gibco) containing 10% DMSO (Sigma). PBMCs were thawed in a 37°C water bath and immediately mixed with complete cell media (RPMI 1640, 10% HS-FBS, penicillin, streptomycin, L-glutamine) supplemented with 50 U/mL of Benzonase (EMDMillipore). Cells were rinsed once with media and then CD3^+^ T-cells were isolated via positive magnetic selection or pan T-cells were isolated via negative selection according to the manufacturer’s protocol (Miltenyi Biotec).

### RNA-seq

RNA was isolated from CD3^+^ T-cells using the Qiagen RNeasy Plus Mini Kit or EN AllPrep Mini Kit with on column DNaseI (Qiagen) digestion. We required samples to have an RNA integrity number (RIN) greater than 7 as determined by Bioanalyzer analysis (Agilient). Illumina sequencing libraries were constructed using the Kapa Stranded RNA-seq Kit with RiboErase (Kapa Biosystems) and 100 ng of total RNA. The samples were sequenced on an Illumina HiSeq 2500, and we obtained approximately 65 million reads per sample. The reads were mapped and assembled into transcripts with STAR software (*35*) using the reference human genome 38 for alignment and Genecode 25 for transcript annotation. The transcripts were quantified using featureCounts (*36*), and differential expression analysis was performed for both lncRNAs and protein-coding genes using DESeq2 (*37*). GOStats was utilized for gene ontology analysis (*38*). All BAM sequencing files were deposited in the sequence read archive (SRA) at the NCBI (accession number PRJNA550422).

### qRT-PCR

RNA was isolated using the Qiagen RNeasy Plus Mini Kit. cDNA was generated using the SuperScript III cDNA synthesis kit (Invitrogen) with random hexamer primers according to the manufacturer’s protocol. qPCR was performed using PowerUp SYBR Green Master Mix (ThermoFisher), and 200 nM forward and reverse primers (**Supplemental Table 10**) (Integrated DNA Technologies) on a Mastercycler realplex^2^ thermocycler (Eppendorf). Primers were designed using the qPCR setting on Primer3Plus (*39*). All primers were checked for a single melting curve peak, the correct size band via agarose gel electrophoresis, and exponential amplification across a 1-10^5^ fold dilution series of cDNA. Fold changes were calculated according to the 2^(-ΔΔCt)^ method.

### Mice

Eight to 12 week-old female C57BL/6 (B6, H-2b, CD45.2) and BALB/c (H-2d, CD45.2) recipient mice were purchased from Charles River Laboratories. Eight to twelve week-old female C3H.SW (CD45.2) recipient and B6.SJL-*Ptprc^a^ Pepc^b^*/BoyJ (B6, CD45.1) donor mice were purchased from Jackson Laboratories. Animals were house under specific pathogen-free conditions. All animal studies were approved by the University Committee on Use and Care of Animals of the University of Michigan, based on University Laboratory Animal Medicine guidelines.

### Murine Allogeneic Bone Marrow Transplant

Murine bone marrow transplantation was performed as previous on 8-12 week-old animals (*20*). Briefly, splenic T-cells from donor mice were enriched using the Pan T-cell Isolation Kit II (Miltenyi Biotec), while bone marrow (BM) cells were depleted of T cells by using CD90.2 magnetic beads (Miltenyi Biotec). C57BL/6 (syngeneic), BALB/c (allogeneic) and C3H.SW (allogeneic) recipient mice were irradiated from a 137Cs source with 9.5, 8 (dose split equally 3-4 hours apart), or 10.5 Gy, respectively, on day - 1. On day 0, recipient C57BL/6, BALB/c, and C3H.SW mice received via tail vein injection 5 x 10^6^ T cell depleted (TCD) BM as well as 2.5, 1.0 or 3.0 x 10^6^ splenic T-cells from B6.SJL-*Ptprc^a^ Pepc^b^*/BoyJ mice, respectively. All recipients received pH 3.0 water starting day -1.

### Mixed Lymphocyte Reactions (MLR)

Cryogenically preserved healthy human peripheral blood T-cells or murine splenic T-cells were magnetically isolated by negative selection using Pan T-cell isolation kits from Miltenyi Biotec. Murine splenocytes underwent RBC lysis (Sigma) prior to negative T-cell enrichment. Human T-cells were then co-cultured with an equal amount of lethally irradiated (30 Gy) autologous or allogeneic monocytes (for allogeneic, pool of 2-3 unrelated donors) derived from the cell fraction bound to the Pan T cell negative selection column. Murine T-cells were co-cultured at a ratio of 2:1 (T-cells to monocytes) with lethally irradiated (30 Gy) C57Bl/6 (syngeneic) or BALB/c (allogeneic) splenic monocytes derived from the cell fraction bound to the Pan T cell negative selection column. 2 x 10^5^ T-cells were cultured per well on flat-bottom 96-well plates, which was scaled by surface area accordingly for larger well platforms as needed. Human MLRs were incubated for 6-7 days and murine MLRs were incubated for 4 days in complete cell media (RPMI 1640, 10% heat-shocked FBS, penicillin, streptomycin, L-glutamine, 2-mercaptoethanol). Where necessary, T-cells were re-isolated following completion of the MLR via negative magnetically-assisted enrichment. Where indicated, instead of incubating with irradiated monocytes, T-cells were incubated with murine or human anti-CD3/CD28 Dyna Beads (ThermoFisher) for 48 hours unless noted otherwise. Where indicated, mouse or human T-cells were cultured with mouse or human IL-7 (10 ng/mL) (R&D Systems) or tacrolimus (Sigma-Aldrich). IL-7 and/or tacrolimus-containing media was refreshed every 2-3 days. To assess T-cell proliferation, incorporation of 3H-thymidine (1 μCi/96 well) during the final 6 hours (murine MLR) or 15 hours (human MLR) of culture was measured by a TopCount (PerkinElmer).

### Flow Cytometry

Flow cytometry analysis was performed with fluorescently-labeled monoclonal antibodies to mouse CD4-APC (GK1.5, BioLegend) or CD8a-PE (53-6.7, BioLegend) or human CD3-PE (UCHT1, BioLegend), CD4-APC (SK3, BioLegend), and CD8a-PE (HIT8a, BiioLegend), as described previously (*40*). Dead cells were stained with Live/Dead Fixable Near-IR (ThermoFisher). Briefly, cells were washed in FACS staining buffer, stained with surface/dead cell markers, washed again and either immediately analyzed using an AttuneNxT flow cytometer or preserved for later analysis using Fixation/Permeabilization and Permeabilization buffers from eBioscience per manufacturer’s instruction.

### Subcellular fractionation

The Cytoplasmic and Nuclear RNA Purification Kit (Norgen Biotek) with on column DNaseI (Norgen Biotek) digestion was used to extract cytoplasmic and nuclear RNA.

### Northern blotting

Ten micrograms of total RNA (RIN > 8 for all samples) was mixed with Gly Loading Dye (Ambion) and heated to 48°C for 1 hour. RNA samples and the Century/Millenium RNA Marker (Ambion) were then loaded onto 1% agarose NorthernMax-Gly gels (Ambion) and electrophoresed at 135V for 2 hours. RNA integrity and loading was assessed via UV illumination. The gel was transferred to a Nytran Supercharge membrane (Whatman) per manufacturer protocol using a TurboBlotter and NorthernMax Transfer Buffer (Ambion) for 2 hours. Transfer efficiency was confirmed via UV visualization of the gel. The membrane was then UV linked and air dried. The membranes were rinsed 3 times with RNase-free water and prehybridized with a solution containing 6X SSC, 5X Denhardt’s solution, 0.5% SDS and 10 μg/ml sheared herring sperm DNA at 68°C for 48 hours. A ^32^P-labeled 368 nt transcript was generated from EcoRI digested probe temple (described below) using T7 RNA polymerase (New England Biolabs). The probe was diluted in the hybridization buffer described above, and hybridization was carried out at 68°C for 24 hours. The membranes were washed twice in a solution containing 2X SSC and 0.1% SDS at 68°C for 60 minutes followed by two rinses with 0.1X SSC and 0.1% SDS at 68°C for 30 minutes. The membranes were auto-radiographed at −80°C for 24h hours and developed. Next, the membranes were stripped (0.01X SSC and 0.1% SDS) at 68°C for 24 hours and re-probed with β-actin random-primed PCR probe generated from a 591 nt PCR template in hybridization solution at 42°C for 18 hours. The membranes were then rinsed twice (2X SSC and 0.1% SDS) at 42°C for 60 minutes, auto radiographed at −80°C for 72 hours, and developed for 3 minutes.

The murine Linc00402 probe template was generated by PCR (GoTaq Green 2X Master Mix, Promega) amplifying a 368 nt region from C57BL/6 T cell-derived DNA using the following forward and reverse primers containing EcoRI and HindIII sites, respectively (Forward: taagca<u>gaattc</u>TGTTAACCCGTGGCATCTTC, Reverse: taagca<u>aagctt</u>TTGTCCCACAGGAGAACTCAAG). The PCR fragment was cleaned up using the Wizard SV Gel and PCR Clean-up System (Promega). The fragment was digested with EcoRI-HF and HindIII-HF (New England Biolabs) per manufacturer instructions and gel purified. The vector pGEM-4Z (Promega) was then digested with EcoRI-HF and HindIII-HF, treated with calf intestinal phosphatase (New England Biolabs), and gel-purified. The digested PCR fragment and pGEM-4Z vector were then ligated using T4 DNA ligase (New England Biolabs) per manufacturer instruction. The probe-containing vector was then transformed into Stellar Competent Cells (Takara) via manufacturer protocol. Minipreps from individual colonies were generated using the Qiaprep Spin Mini Kit (Qiagen). The probe sequence was confirmed via Sanger Sequencing. The murine β-actin PCR probe template was amplified with GoTaq Green 2X Master Mix from C57BL/6 T-cell cDNA using the following primers (AGATGACCCAGATCATGTTTGAG, TCTGCATCCTGTCAGCAATG). The PCR probe template was gel purified prior to generating a ^32^P-labeled, random-primed PCR probe.

### GapmeR knockdown

HPLC purified and salt-exchanged GapmeR (Exiqon/Qiagen) antisense oligonucleotides were incubated with T-cells at the indicated concentration for the indicated time. Their sequences are: negative control-AACACGTCTATACGC, murine Gm34199-ACAGTATGATTAAGAT, human Linc00402-CATGACATAACAGAAA, and human RP11-348F1.3-CACTAATGAGATGAAC. The murine Malat-1 positive control GapmeR was purchased from Exiqon/Qiagen. To assess viability, the exclusion of 7-AAD and Annexin V (BioLegend) was assessed at various time points via flow cytometry per manufacturer recommendations.

### CRISPR-Cas9 ribonucleoprotein (RNP) nucleoporation of T-cells

CRISPR-Cas9 RNP nucleoporation was carried out essentially as previously described (*22*). Briefly, negative magnetically selected mouse or human T-cells, were incubated for 2-24 hours in the presence of IL-7 (10 ng/mL). Prior to nucleofection, cells were spun at 100 x g at room temperature for 10 minutes, rinsed with 37°C PBS (Gibco), and pelleted again. Rinsed cells were resuspended in buffer P4 (mouse) or P2 (human) at 1-10 x 10^6^ cells/20 μL (Lonza). Non-targeting Alt-R crRNA #1, Alt-R tracrRNA, and Alt-R ATTO-550-labeled tracrRNA were purchased from IDT. Custom designed human Linc00402-targeting Alt-R crRNAs were designed using IDT’s online design tool (Linc00402-1: ATCTCATTGACACTGCAAAC, Linc00402-2: GCTCTTAGGAAGATCCAAGC, and Linc00402-3: CAATGTGCTGGAGCATCCAC). crRNAs and tracrRNA were dissolved in Nuclease-Free Duplex Buffer (IDT) at 100 μM. Individual crRNAs and tracrRNA were mixed at equimolar concentrations in sterile PCR tubes and annealed by heating to 95°C in a thermocycler for 5 minutes followed by cooling for at least 10 minutes at room temperature. In another sterile PCR tube, 3 annealed duplexes (targeting human Linc00402) were mixed by adding 3 μL of each (150 pmol) and 6 μL (180 pmol) of TruCut Cas9 Protein v2 (Thermo Fisher Scientific) and incubated for 10 minutes at room temperature. Nine μL (450 pmol total) of the non-targeting control duplex was mixed with Cas9. Twenty μL of resuspended T-cells were then gently mixed with the RNP complexes and incubated for 2 minutes at room temperature. The entire volume of each T-cell/RNP mix was then transferred to a Lonza 4D Nucleofector 16-well cuvette. Mouse T-cells received pulse DS137 and human T-cells received pulse EH100. Post-nucleofection, pre-warmed IL-7-containing media (80-165 μL) was added directly to the cuvettes and incubated for 10 min at room temperature. Cells were gently resuspended and aliquotted into 96-wells containing pre-warmed IL-7-containing media at a density of 1-2 million cells/well in a final volume of 200 μL/well. Cells were then incubated in humidified 5% CO_2_ at 37°C for 6-15 hours before rinsing away the RNP complexes and re-culturing the cells in IL-7-containing media for a minimum of 72 total hours post-nucleofection prior to further experimentation.

### Statistics

GraphPad Prism7 was used to calculate all statistics. *P*-values were calculated using an ANOVA or unpaired, two-tailed t-test where appropriate. *P*-values less than 0.05 were considered significant unless noted elsewhere. To mitigate the effects of small sample size and heterogeneity in our confirmation cohort, samples with fold changes greater than +/−1.3 standard deviations from the mean (i.e. the approximate top and bottom 10% of the normal distribution) were eliminated from the analysis. At least three independent replicates of experiments were performed unless noted otherwise.

## Acknowledgements

DP was supported by the NIH T32 Training Grant in Molecular and Translational Hematology (4T32HL007622-30), NIH K12 Children’s Health Research Center Development Award (5K12HD028820-27), and the Hope From Harper St. Baldrick’s Foundation Fellowship. All other authors were supported by National Institutes of Health RO1 grants HL128046, CA203542, and CA217156 of which PR is the PI.

We’d like to thank the NY Genome Institute for their assistance with the human T-cell RNA-sequencing and analysis. We’d like to thank Lofstrand Labs for their assistance with northern blot analysis.

## Author Contribution

DP and PR conceived of the project.

DP performed, analyzed, and interpreted the experiments.

NR provided bioinformatics support.

GH, CJ, MR, KO, DS, IH, JW, SK, YS, and HF performed experiments.

DP and PR wrote and edited the manuscript.

## SUPPLEMENTARY MATERIALS

**Supplemental Figure 1:**
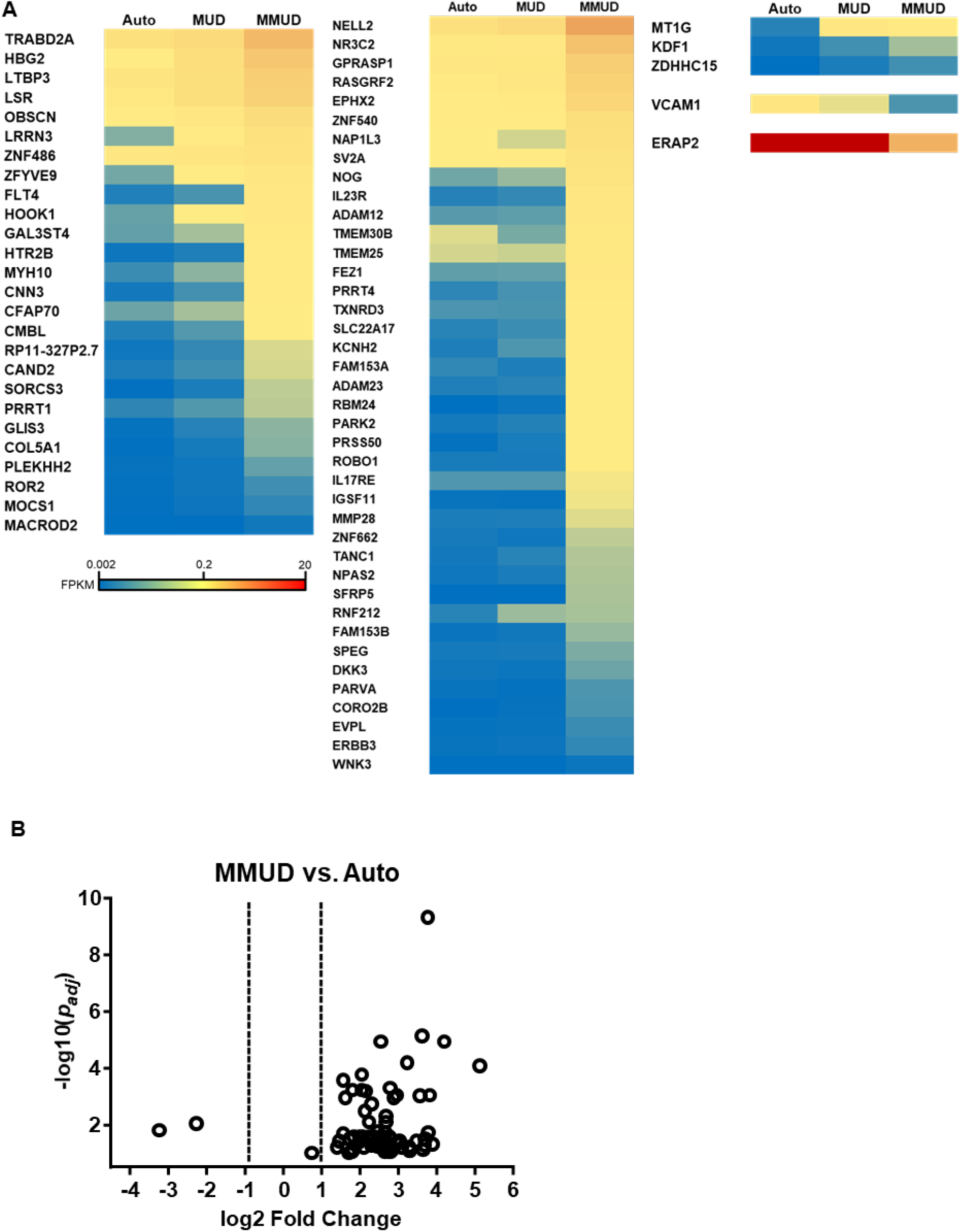
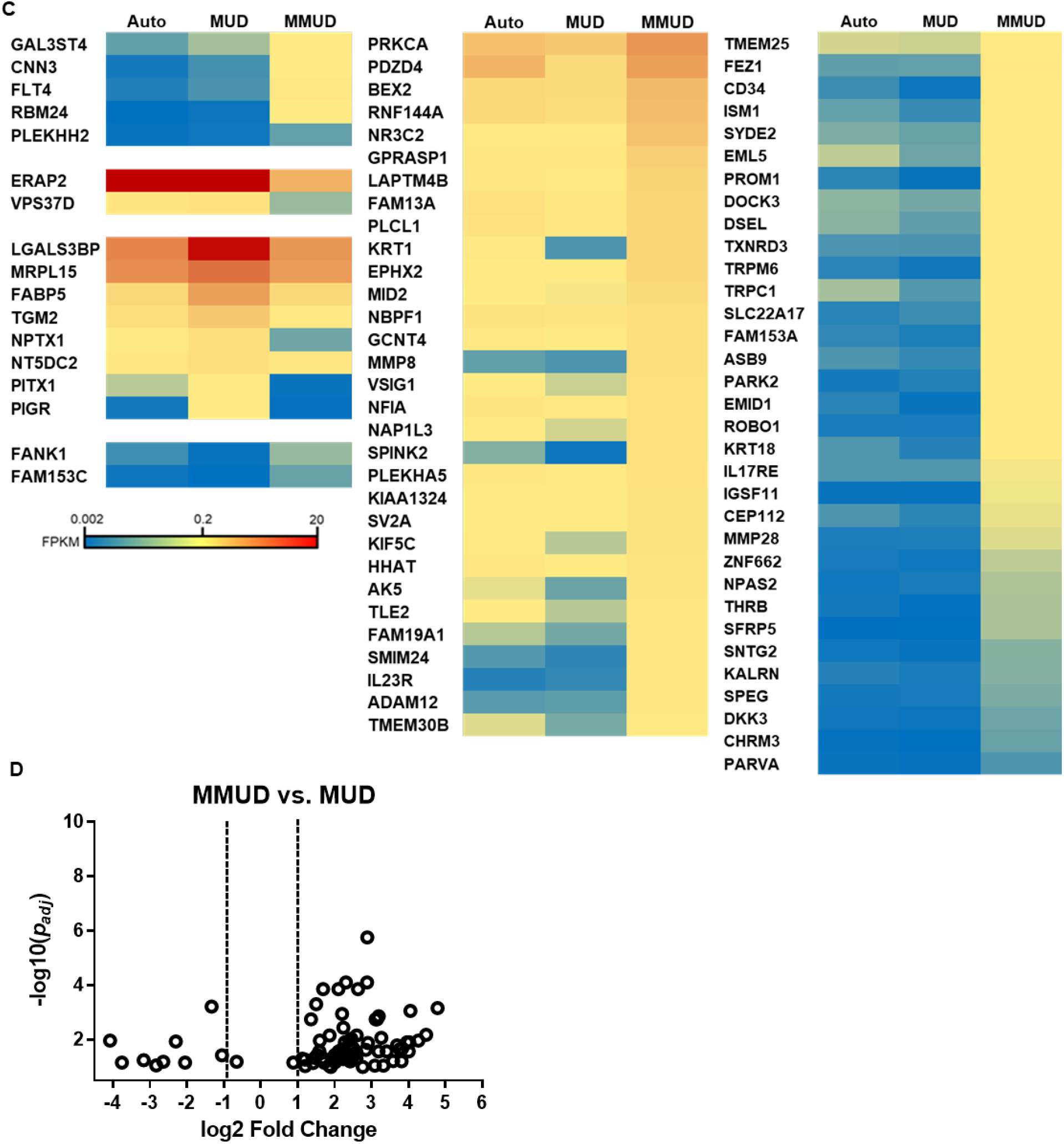
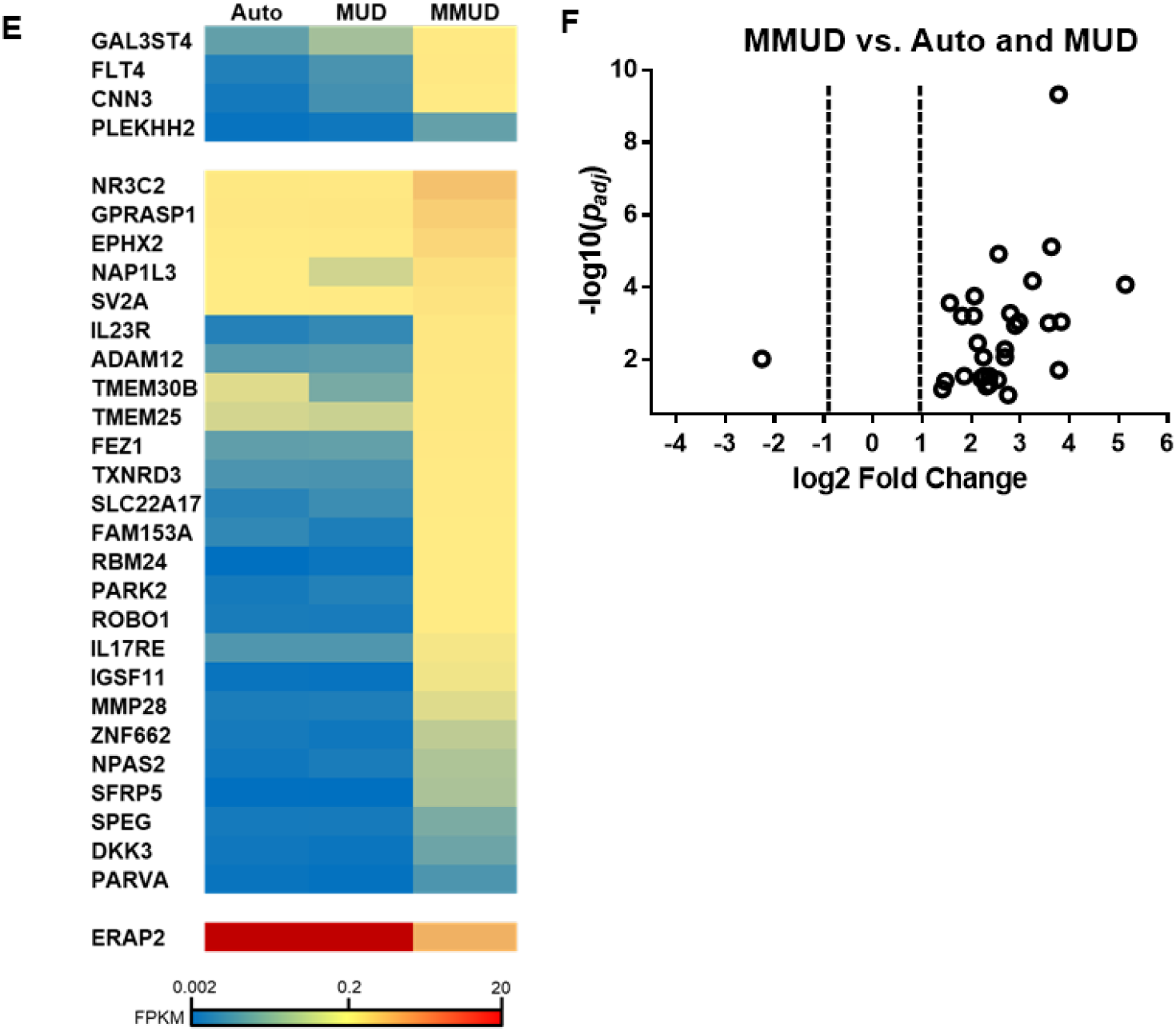
Differentially expressed protein-coding genes in human T-cells identified by RNA-seq. Heat maps showing differentially expressed MMUD protein-coding genes relative to autologous HSCT patients (*p*<0.05, *p_adj_<*0.1) (**A**), MUD HSCT patients (**C**), and those shared relative to both autologous and MUD patients (**E**) are shown. Protein-coding genes are grouped by expression pattern. Volcano plots of differentially expressed (*p* < 0.05, *p_adj_ <* 0.1) lncRNAs in the MMUD group relative to autologous controls (**B**), MUD controls (**D**), and those shared relative to both autologous and MUD patients (**F**) are shown.

**Supplemental Figure 2:**
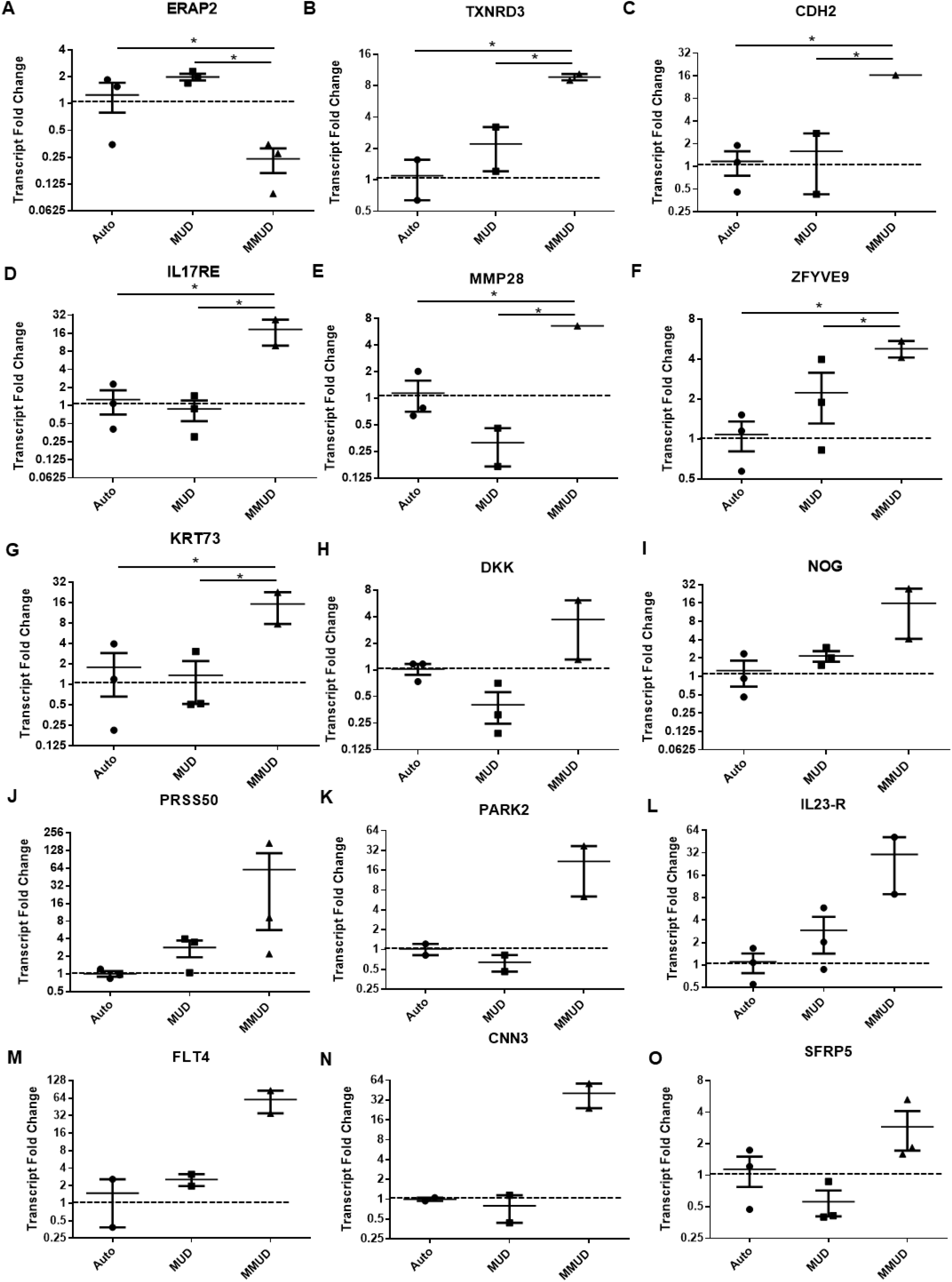
Confirmatory expression of protein-coding genes in RNA-seq patient cohort. qRT-PCR was performed on total RNA derived from CD3^+^ peripheral blood T-cells from patients detailed in Table 1. The relative expression of ERAP2 (**A**), TXNRD3 (**B**), CDH2 (**C**), IL17RE (**D**), MMP28 (**E**), ZFYVE9 (**F**), KRT73 (**G**), DKK (**I**), NOG (**J**), PRSS50 (**K**), PARK2 (**L**), IL23-R (**M**), FLT4 (**N**), CNN3 (**O**), and SFRP5 (**P**) are shown relative to β-actin. Error bars represent the mean +/− the SEM. \**p* < 0.05 by multiple comparison using original FDR method of Benjamini and Hochberg following a one-way ANOVA in which *p* <0.1 (See supplemental table for further details).

**Supplemental Figure 3:**
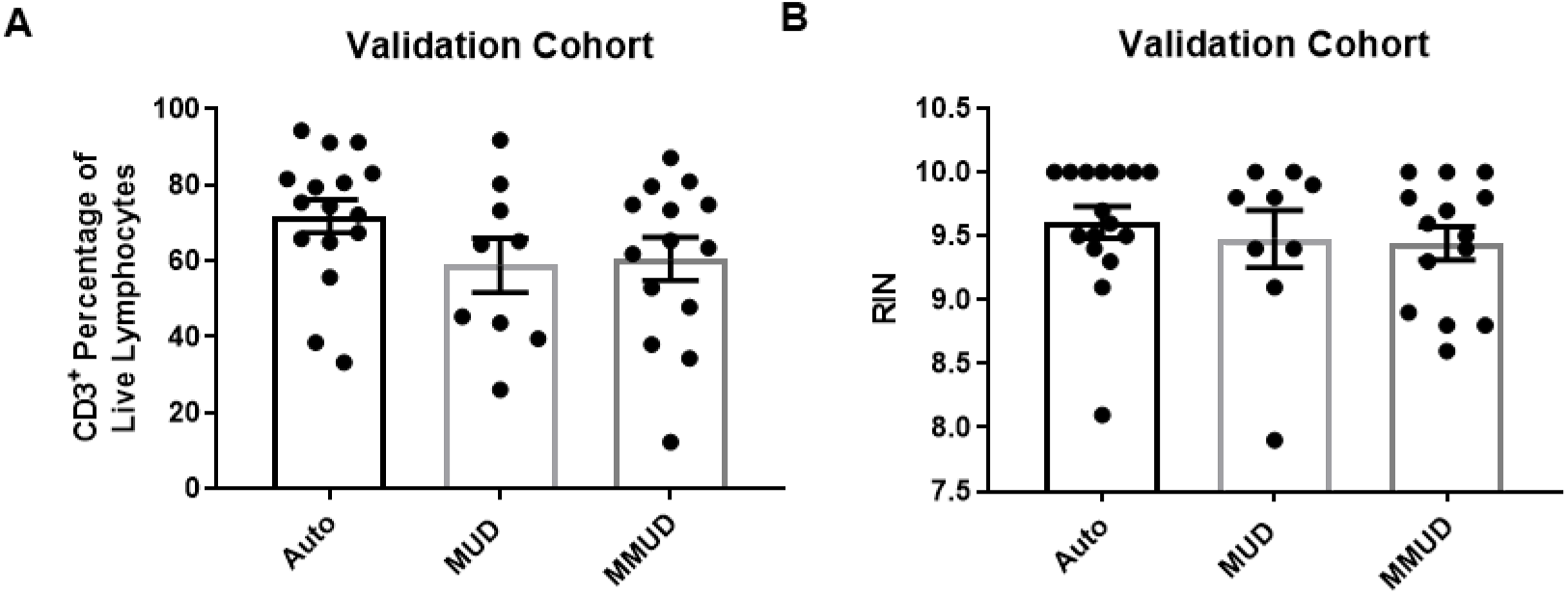
Validation patient cohort CD3^+^ purity and RNA integrity values. Following magnetically assisted CD3 positive selection of frozen human PBMCs, cells were stained with an anti-CD3 antibody and a dead cell marker. These were analyzed by flow cytometry and the CD3^+^ percentage of live lymphocytes is depicted in **A**. Following RNA isolation from the cells described in A, RNA integrity was assessed via Bioanalyzer analysis and presented in **B.** Error bars represent the mean +/− the SEM.

**Supplemental Figure 4:**
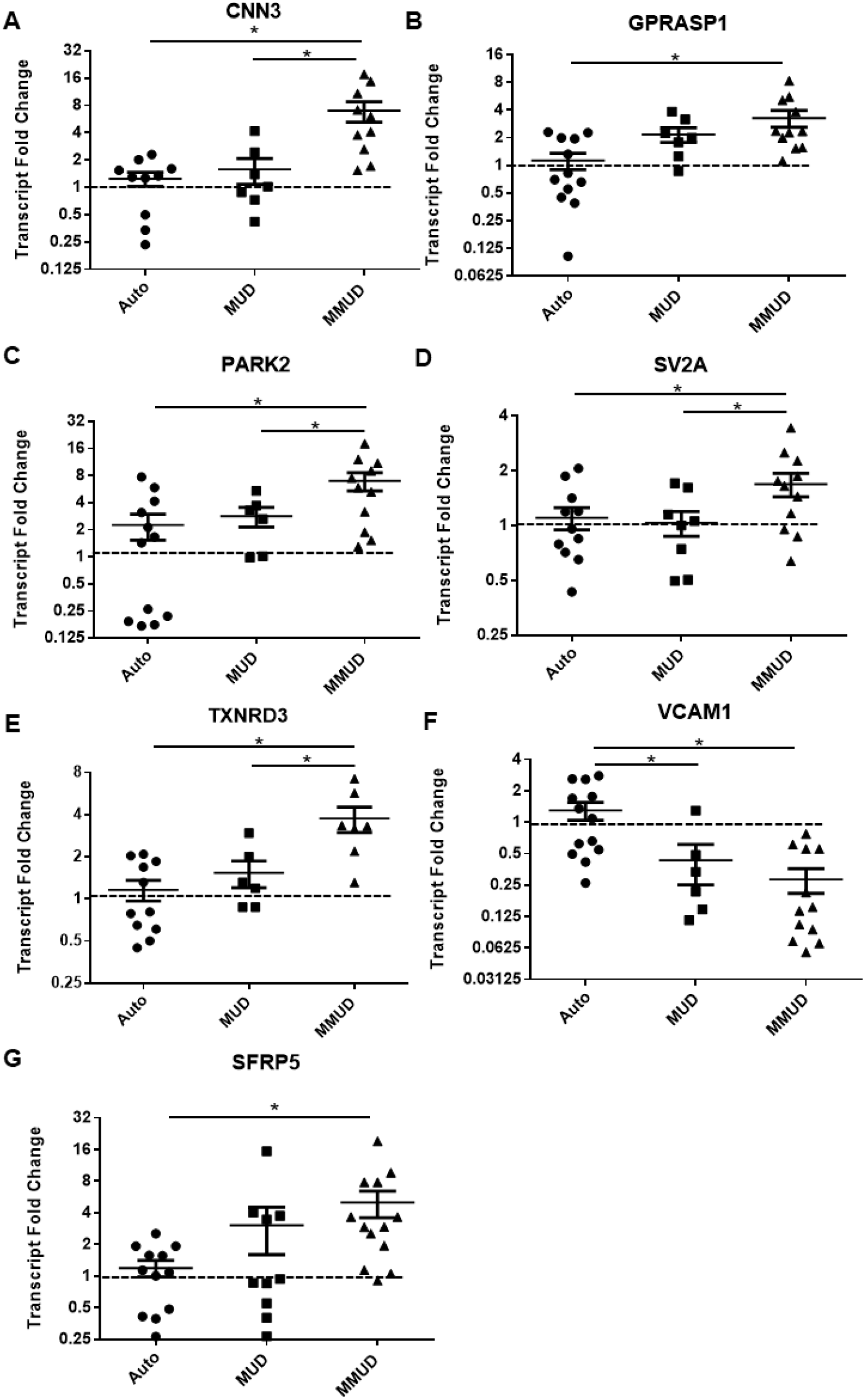
Expression of protein-coding genes in an independent patient cohort. qRT-PCR was performed on cryogenically preserved CD3^+^ peripheral blood T-cells from patients detailed in Table 2. The relative expression of CNN3 (**A**), GPRASP1 (**B**), PARK2 (**C**), SV2A (**D**), TXNRD3 (**E**), VCAM1 (**F**), and SFRP5 (**G**) are shown relative to β-actin. Error bars represent the mean +/− the SEM. \**p* < 0.05 (one-way ANOVA, original FDR method of Benjamini and Hochberg).

**Supplemental Figure 5:**
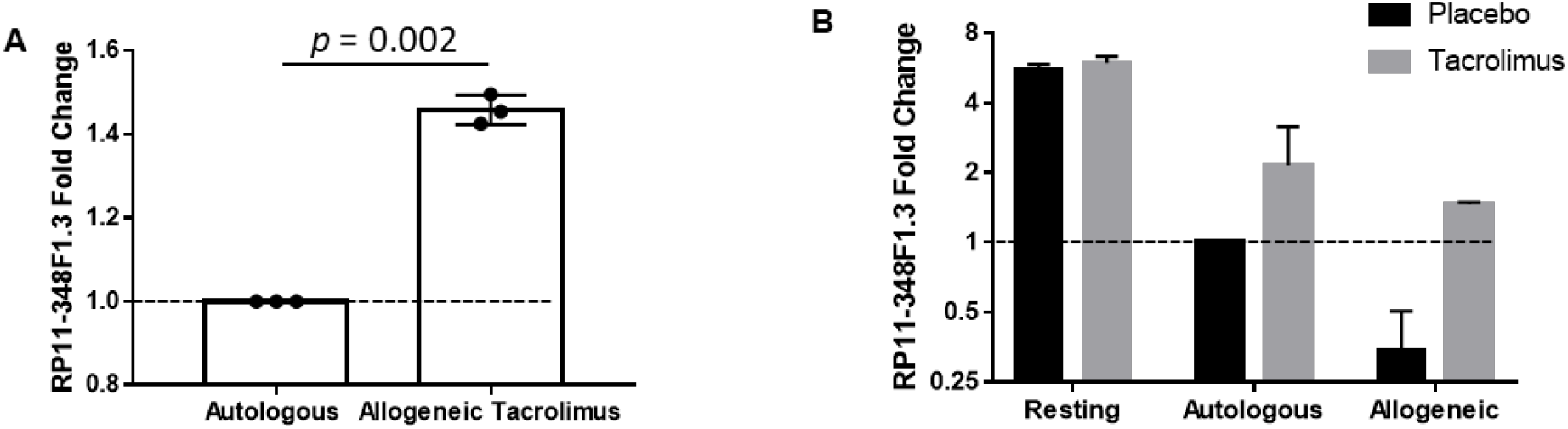
Treatment of activated T-cells with tacrolimus preserves RP11-348F1.3 expression. Human T-cells were pre-incubated with tacrolimus (30 ng/mL) or placebo for 1 hour prior to mixing them with irradiated (30 Gy) autologous or allogeneic monocytes or placebo. Seven days later, T-cells were negatively magnetically isolated and subjected to qRT-PCR analysis of RP11-348F1.3 with fold changes set relative to autologous placebo-treated controls (**A and B**) (n = 2-3 per condition, \**p*<0.05).

**Supplemental Figure 6:**
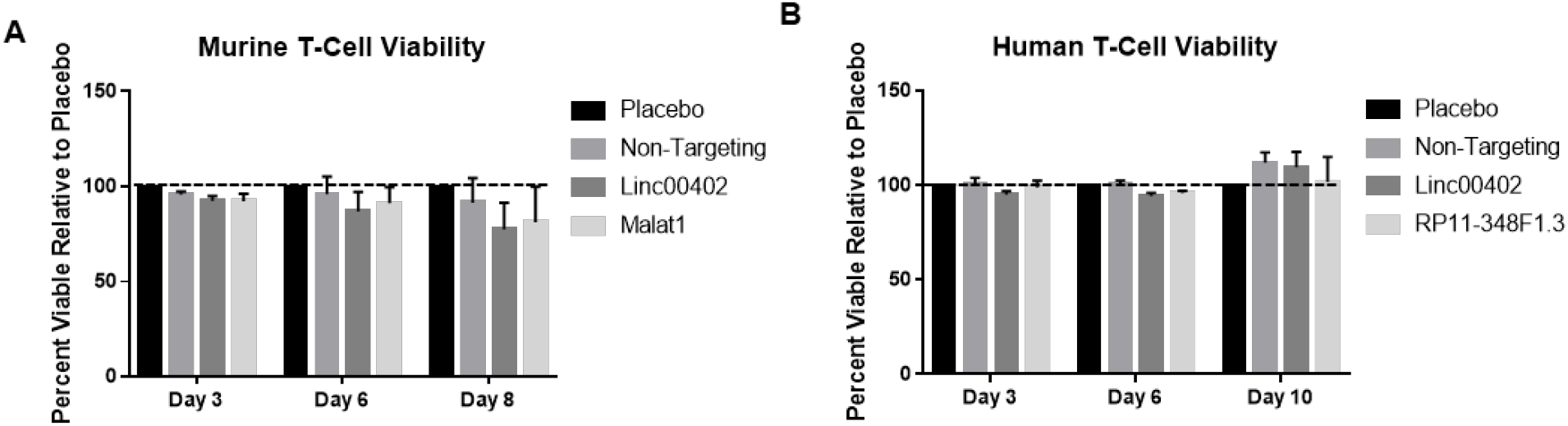
GapmeRs are not cytotoxic to T-cells. Mouse (**A**) or human (**B**) T-cells were incubated with 500 nM of the indicated GapmeRs described in figure 6 for the times indicated. Viability was assessed via the exclusion of 7-AAD and annexin V as measured by flow cytometry relative to placebo-treated control cells. Error bars represent the mean +/− the SEM (n=3 per condition).

**Supplemental Table 4.** Enriched GO terms for differentially express protein coding genes between the MMUD and Auto groups.

| GOBPID | P value | Odds Ratio | ExpCount | Count | Size | Term |
| --- | --- | --- | --- | --- | --- | --- |
| GO:0035924 | 7.50E-05 | 44.28571429 | 0.083586626 | 3 | 22 | cellular response to vascular endothelial growth factor stimulus |
| GO:0071363 | 0.000113 | 5.579545455 | 1.95668693 | 9 | 515 | cellular response to growth factor stimulus |
| GO:0070848 | 0.000127 | 5.48881323 | 1.987082067 | 9 | 523 | response to growth factor |
| GO:0070887 | 0.000265 | 3.372201783 | 6.363981763 | 16 | 1675 | cellular response to chemical stimulus |
| GO:0030154 | 0.000355 | 3.062607872 | 8.708206687 | 19 | 2292 | cell differentiation |
| GO:0048856 | 0.000535 | 2.817783219 | 12.21884498 | 23 | 3216 | anatomical structure development |
| GO:0048869 | 0.000919 | 2.794310963 | 9.36550152 | 19 | 2465 | cellular developmental process |
| GO:0048646 | 0.000989 | 4.075146628 | 2.625379939 | 9 | 691 | anatomical structure formation involved in morphogenesis |
| GO:0044767 | 0.001102 | 2.625888365 | 13.68541033 | 24 | 3602 | single-organism developmental process |
| GO:0007520 | 0.001248 | 45.68604651 | 0.053191489 | 2 | 14 | myoblast fusion |
| GO:0032502 | 0.001326 | 2.581133862 | 13.84878419 | 24 | 3645 | developmental process |
| GO:1990267 | 0.001367 | 15.25194805 | 0.220364742 | 3 | 58 | response to transition metal nanoparticle |
| GO:0061448 | 0.00163 | 8.757223265 | 0.509118541 | 4 | 134 | connective tissue development |
| GO:0014902 | 0.002072 | 13.09709821 | 0.254559271 | 3 | 67 | myotube differentiation |
| GO:0007612 | 0.002162 | 12.89450549 | 0.258358663 | 3 | 68 | learning |
| GO:0090090 | 0.002162 | 12.89450549 | 0.258358663 | 3 | 68 | negative regulation of canonical Wnt signaling pathway |
| GO:0001736 | 0.002317 | 32.23529412 | 0.07218845 | 2 | 19 | establishment of planar polarity |
| GO:0007164 | 0.002317 | 32.23529412 | 0.07218845 | 2 | 19 | establishment of tissue polarity |
| GO:0051146 | 0.00252 | 7.733200597 | 0.573708207 | 4 | 151 | striated muscle cell differentiation |
| GO:0000768 | 0.002568 | 30.44186047 | 0.075987842 | 2 | 20 | syncytium formation by plasma membrane fusion |
| GO:0044707 | 0.002802 | 2.397069733 | 15.45972644 | 25 | 4069 | single-multicellular organism process |
| GO:0060071 | 0.003107 | 27.39302326 | 0.083586626 | 2 | 22 | Wnt signaling pathway, planar cell polarity pathway |
| GO:0006949 | 0.003394 | 26.08637874 | 0.087386018 | 2 | 23 | syncytium formation |
| GO:0090175 | 0.003394 | 26.08637874 | 0.087386018 | 2 | 23 | regulation of establishment of planar polarity |
| GO:0022603 | 0.003424 | 4.003650745 | 1.998480243 | 7 | 526 | regulation of anatomical structure morphogenesis |
| GO:0009810 | 0.003799 | Inf | 0.003799392 | 1 | 1 | stilbene metabolic process |
| GO:0010636 | 0.003799 | Inf | 0.003799392 | 1 | 1 | positive regulation of mitochondrial fusion |
| GO:0035989 | 0.003799 | Inf | 0.003799392 | 1 | 1 | tendon development |
| GO:0036363 | 0.003799 | Inf | 0.003799392 | 1 | 1 | transforming growth factor beta activation |
| GO:0044333 | 0.003799 | Inf | 0.003799392 | 1 | 1 | Wnt signaling pathway involved in digestive tract morphogenesis |
| GO:0046271 | 0.003799 | Inf | 0.003799392 | 1 | 1 | phenylpropanoid catabolic process |
| GO:0046272 | 0.003799 | Inf | 0.003799392 | 1 | 1 | stilbene catabolic process |
| GO:0060302 | 0.003799 | Inf | 0.003799392 | 1 | 1 | negative regulation of cytokine activity |
| GO:0097168 | 0.003799 | Inf | 0.003799392 | 1 | 1 | mesenchymal stem cell proliferation |
| GO:1900673 | 0.003799 | Inf | 0.003799392 | 1 | 1 | olefin metabolic process |
| GO:1902302 | 0.003799 | Inf | 0.003799392 | 1 | 1 | regulation of potassium ion export |
| GO:1902303 | 0.003799 | Inf | 0.003799392 | 1 | 1 | negative regulation of potassium ion export |
| GO:1902460 | 0.003799 | Inf | 0.003799392 | 1 | 1 | regulation of mesenchymal stem cell proliferation |
| GO:19024<br>62 | 0.00379<br>9 | Inf | 0.0037993<br>92 | 1 | 1 | positive regulation of mesenchymal stem cell proliferation |
| GO:19032<br>25 | 0.00379<br>9 | Inf | 0.0037993<br>92 | 1 | 1 | negative regulation of endodermal cell differentiation |
| GO:19032<br>65 | 0.00379<br>9 | Inf | 0.0037993<br>92 | 1 | 1 | positive regulation of tumor necrosis factor-mediated signaling pathway |
| GO:19901<br>72 | 0.00379<br>9 | Inf | 0.0037993<br>92 | 1 | 1 | G-protein coupled receptor catabolic process |
| GO:20000<br>40 | 0.00379<br>9 | Inf | 0.0037993<br>92 | 1 | 1 | regulation of planar cell polarity pathway involved in axis elongation |
| GO:20000<br>41 | 0.00379<br>9 | Inf | 0.0037993<br>92 | 1 | 1 | negative regulation of planar cell polarity pathway involved in axis elongation |
| GO:20000<br>56 | 0.00379<br>9 | Inf | 0.0037993<br>92 | 1 | 1 | regulation of Wnt signaling pathway involved in digestive tract morphogenesis |
| GO:20000<br>57 | 0.00379<br>9 | Inf | 0.0037993<br>92 | 1 | 1 | negative regulation of Wnt signaling pathway involved in digestive tract morphogenesis |
| GO:00017<br>38 | 0.00400<br>6 | 23.813953<br>49 | 0.0949848<br>02 | 2 | 25 | morphogenesis of a polarized epithelium |
| GO:00713<br>10 | 0.00404<br>1 | 2.8028178<br>46 | 5.1937689<br>97 | 12 | 136<br>7 | cellular response to organic substance |
| GO:00071<br>55 | 0.00406<br>6 | 3.5189743<br>18 | 2.6253799<br>39 | 8 | 691 | cell adhesion |
| GO:00226<br>10 | 0.00413<br>8 | 3.5080686<br>53 | 2.6329787<br>23 | 8 | 693 | biological adhesion |
| GO:00100<br>43 | 0.00466<br>4 | 21.905116<br>28 | 0.1025835<br>87 | 2 | 27 | response to zinc ion |
| GO:00355<br>67 | 0.00466<br>4 | 21.905116<br>28 | 0.1025835<br>87 | 2 | 27 | non-canonical Wnt signaling pathway |
| GO:00325<br>01 | 0.00469<br>2 | 2.2750358<br>51 | 15.991641<br>34 | 25 | 420<br>9 | multicellular organismal process |
| GO:00083<br>33 | 0.00501<br>1 | 21.060822<br>9 | 0.1063829<br>79 | 2 | 28 | endosome to lysosome transport |
| GO:00301<br>78 | 0.00508<br>4 | 9.3980738<br>36 | 0.3495440<br>73 | 3 | 92 | negative regulation of Wnt signaling pathway |
| GO:00422<br>21 | 0.00514<br>4 | 2.3927225<br>41 | 9.1375379<br>94 | 17 | 240<br>5 | response to chemical |
| GO:0023057 | 0.005298 | 3.35680218 | 2.743161094 | 8 | 722 | negative regulation of signaling |
| GO:0010648 | 0.005387 | 3.346821682 | 2.750759878 | 8 | 724 | negative regulation of cell communication |
| GO:2000027 | 0.005398 | 9.189952904 | 0.357142857 | 3 | 94 | regulation of organ morphogenesis |
| GO:0007507 | 0.005841 | 4.774916944 | 1.162613982 | 5 | 306 | heart development |
| GO:0007275 | 0.006095 | 2.261693156 | 11.78951368 | 20 | 3103 | multicellular organismal development |
| GO:0061035 | 0.00612 | 18.87730553 | 0.117781155 | 2 | 31 | regulation of cartilage development |
| GO:0009653 | 0.00623 | 2.552604167 | 6.204407295 | 13 | 1633 | anatomical structure morphogenesis |
| GO:0006935 | 0.006533 | 3.962323391 | 1.698328267 | 6 | 447 | chemotaxis |
| GO:0042330 | 0.006533 | 3.962323391 | 1.698328267 | 6 | 447 | taxis |
| GO:0051216 | 0.006589 | 8.528425656 | 0.383738602 | 3 | 101 | cartilage development |
| GO:0007167 | 0.006854 | 3.203535976 | 2.864741641 | 8 | 754 | enzyme linked receptor protein signaling pathway |
| GO:0050793 | 0.006906 | 2.698896186 | 4.840425532 | 11 | 1274 | regulation of developmental process |
| GO:0072358 | 0.007067 | 3.474880383 | 2.28343465 | 7 | 601 | cardiovascular system development |
| GO:0072359 | 0.007067 | 3.474880383 | 2.28343465 | 7 | 601 | circulatory system development |
| GO:0042692 | 0.007335 | 5.65804878 | 0.775075988 | 4 | 204 | muscle cell differentiation |
| GO:0003064 | 0.007585 | 268.1363636 | 0.007598784 | 1 | 2 | regulation of heart rate by hormone |
| GO:0003402 | 0.007585 | 268.1363636 | 0.007598784 | 1 | 2 | planar cell polarity pathway involved in axis elongation |
| GO:0009698 | 0.007585 | 268.1363636 | 0.007598784 | 1 | 2 | phenylpropanoid metabolic process |
| GO:0010760 | 0.007585 | 268.1363636 | 0.007598784 | 1 | 2 | negative regulation of macrophage chemotaxis |
| GO:0036309 | 0.007585 | 268.1363636 | 0.007598784 | 1 | 2 | protein localization to M-band |
| GO:0038155 | 0.007585 | 268.1363636 | 0.007598784 | 1 | 2 | interleukin-23-mediated signaling pathway |
| GO:0060513 | 0.007585 | 268.1363636 | 0.007598784 | 1 | 2 | prostatic bud formation |
| GO:0090193 | 0.007585 | 268.1363636 | 0.007598784 | 1 | 2 | positive regulation of glomerulus development |
| GO:1900452 | 0.007585 | 268.1363636 | 0.007598784 | 1 | 2 | regulation of long term synaptic depression |
| GO:19025<br>28 | 0.00758<br>5 | 268.13636<br>36 | 0.0075987<br>84 | 1 | 2 | regulation of protein linear polyubiquitination |
| GO:19025<br>30 | 0.00758<br>5 | 268.13636<br>36 | 0.0075987<br>84 | 1 | 2 | positive regulation of protein linear polyubiquitination |
| GO:20007<br>41 | 0.00758<br>5 | 268.13636<br>36 | 0.0075987<br>84 | 1 | 2 | positive regulation of mesenchymal stem cell differentiation |
| GO:00017<br>06 | 0.00864 | 15.633222<br>59 | 0.1405775<br>08 | 2 | 37 | endoderm formation |
| GO:00447<br>08 | 0.00865<br>4 | 5.3839721<br>25 | 0.8130699<br>09 | 4 | 214 | single-organism behavior |
| GO:00610<br>61 | 0.00946<br>3 | 4.2256637<br>17 | 1.3069908<br>81 | 5 | 344 | muscle structure development |
| GO:00020<br>40 | 0.00956<br>6 | 14.785669<br>39 | 0.1481762<br>92 | 2 | 39 | sprouting angiogenesis |
| GO:00608<br>28 | 0.00986<br>1 | 7.3214285<br>71 | 0.4445288<br>75 | 3 | 117 | regulation of canonical Wnt signaling pathway |

**Supplemental Table 5.**
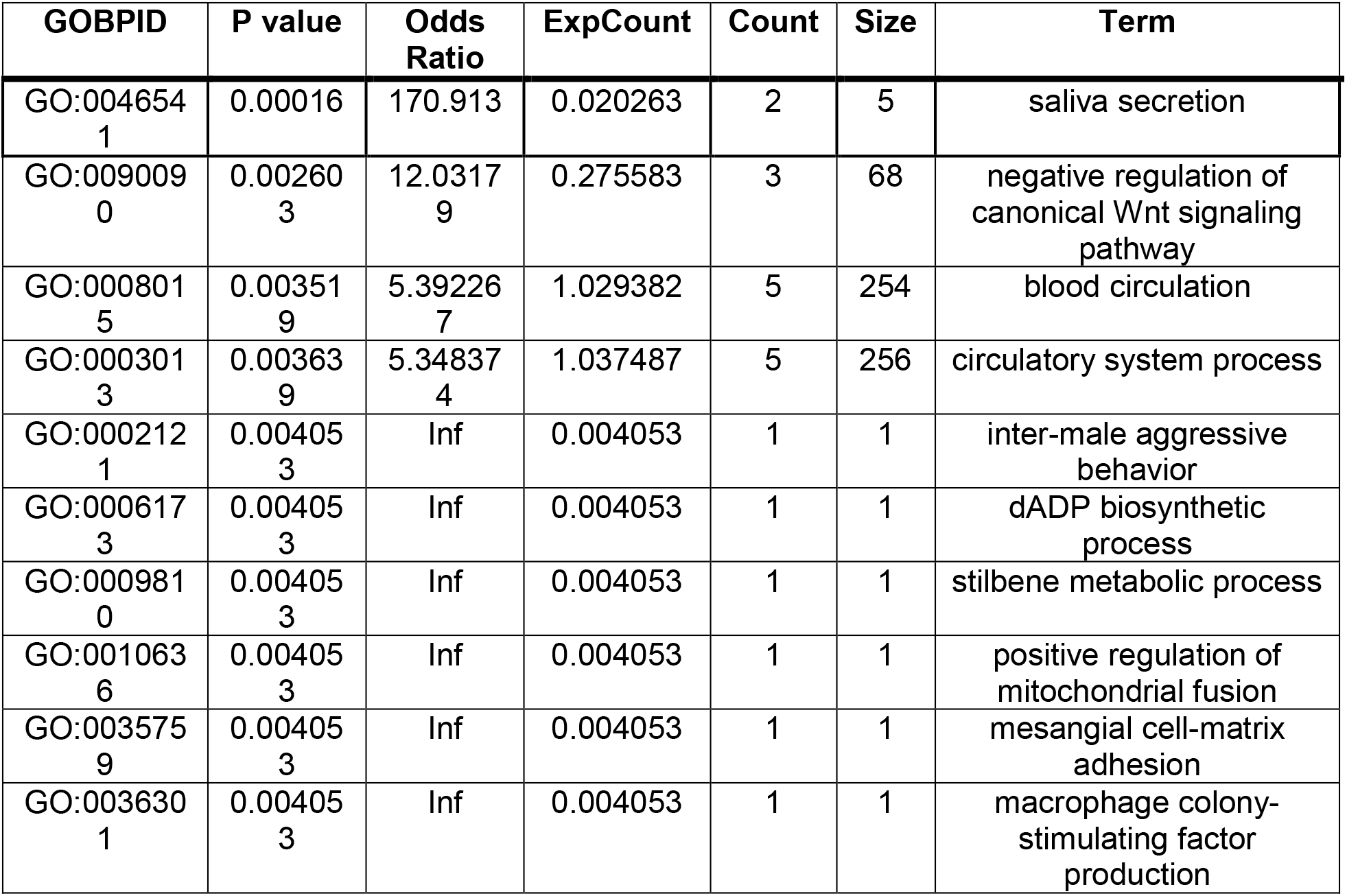
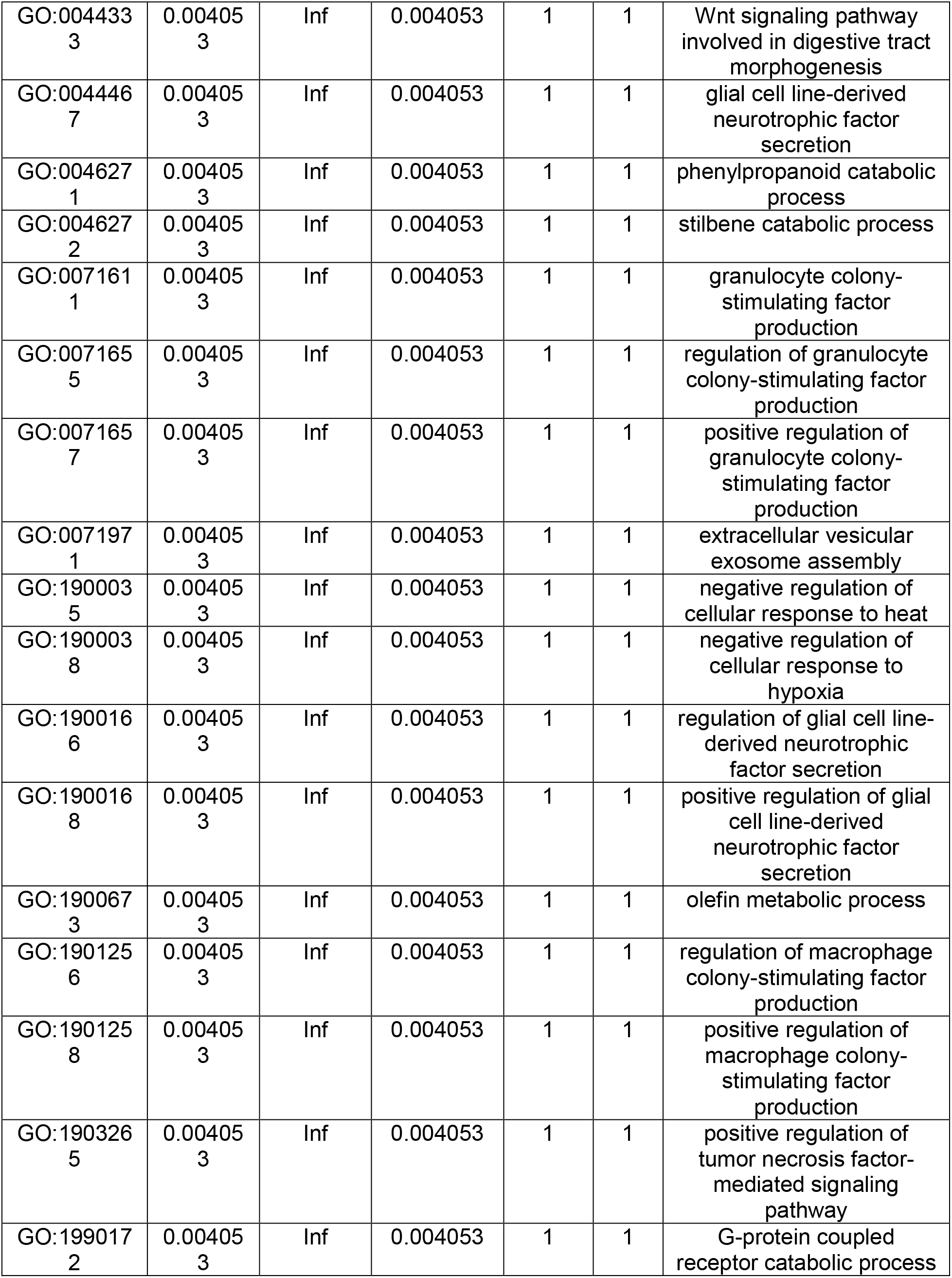
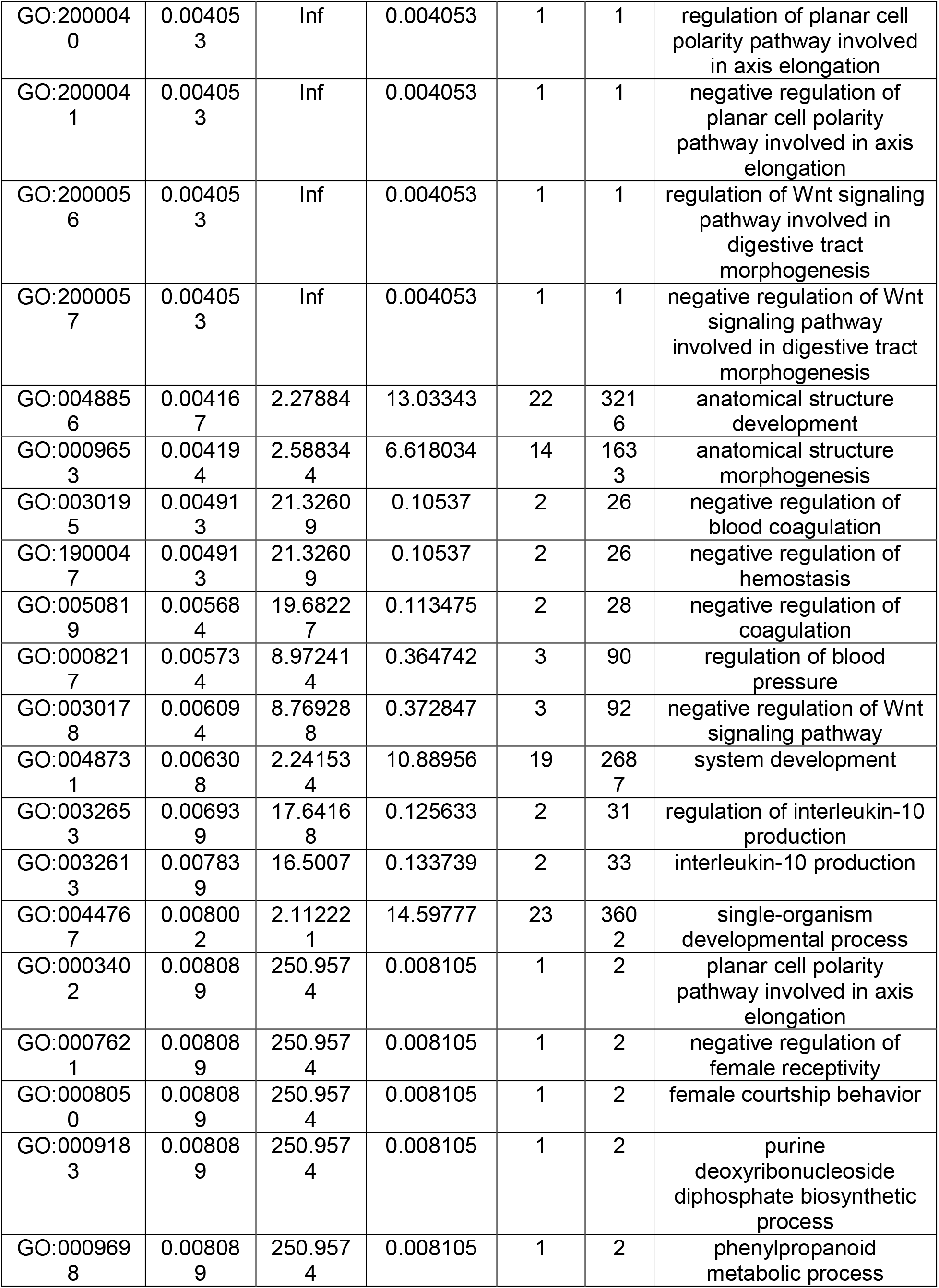
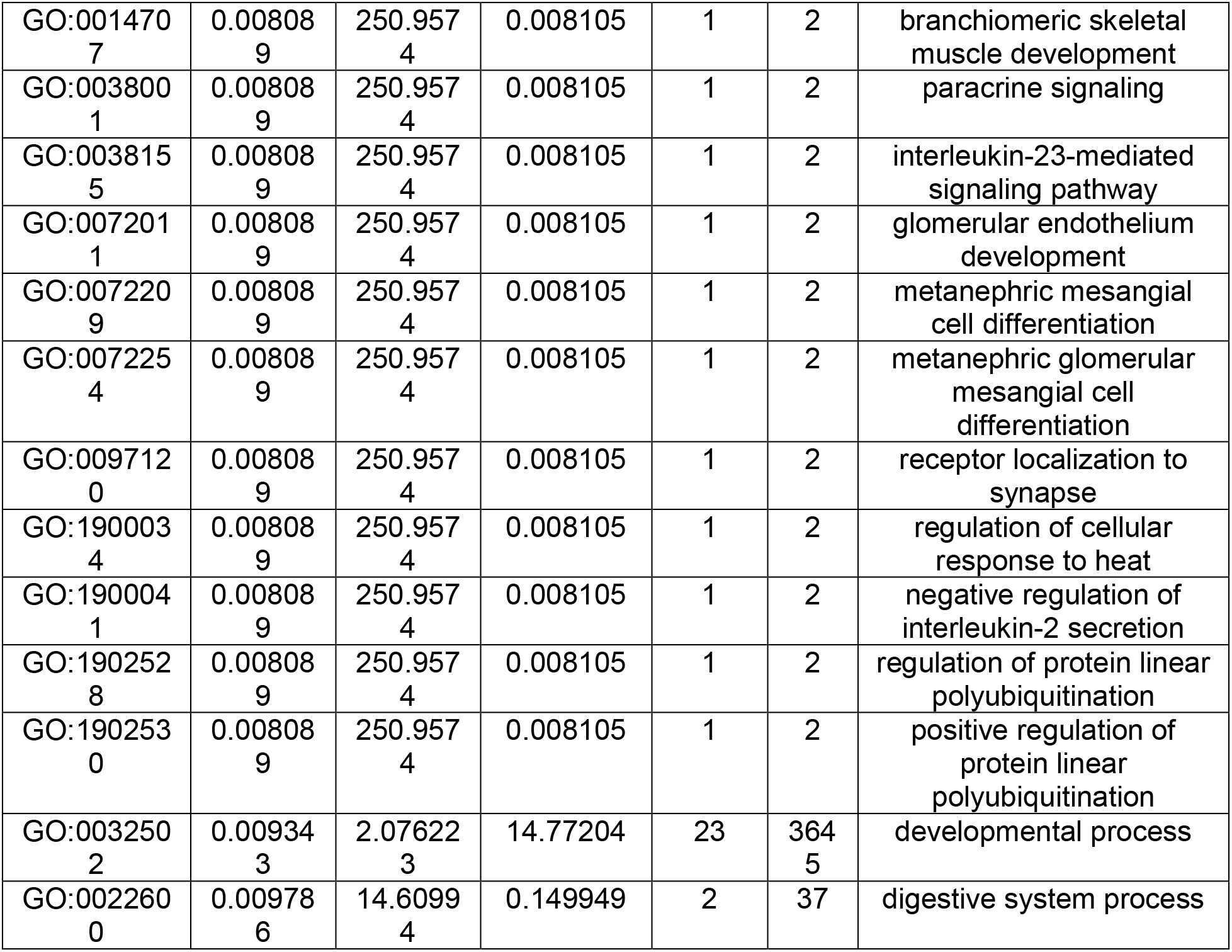
Enriched GO terms for differentially express protein coding genes between the MMUD and MUD groups.

**Supplemental Table 6.**
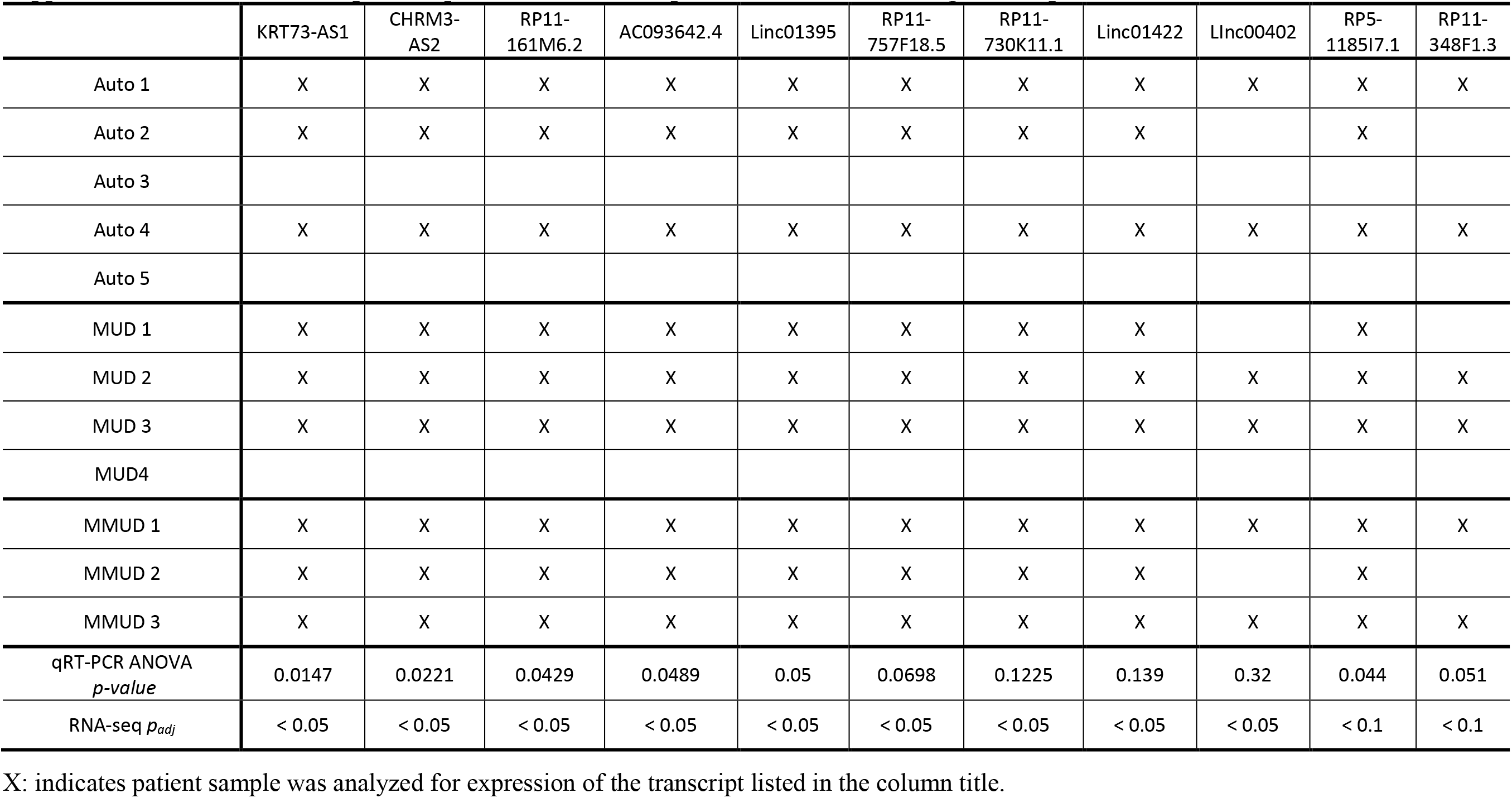
RNA-seq cohort patients tested for expression of each lncRNA gene via qRT-PCR.

**Supplemental Table 7.** RNA-seq cohort patients tested for expression of each protein-coding gene via qRT-PCR.

|  | MMP2<br>8 | ERAP2 | IL17-RE | ZFYVE9 | CNN3 | FLT4 | IL23-R | SFRP5 | NOG | PARK<br>2 | DKK | HTR2B | TRABD2A | CDH2 | KRT7<br>3 | RORC |
| --- | --- | --- | --- | --- | --- | --- | --- | --- | --- | --- | --- | --- | --- | --- | --- | --- |
| Auto 1 | X | X | X | X | X | X | X | X | X | X | X | X | X | X | X | X |
| Auto 2 | X | X | X | X |  |  | X | X | X |  |  | X | X |  | X | X |
| Auto 3 |  |  |  |  |  |  |  |  |  |  |  |  |  |  |  |  |
| Auto 4 | X | X | X | X | X | X | X | X | X | X | X | X | X | X | X | X |
| Auto 5 |  |  |  |  |  |  |  |  |  |  |  |  |  |  |  |  |
| MUD 1 | X | X | X | X |  |  | X | X | X |  | X | X | X | X | X | X |
| MUD 2 |  | X | X | X | X | X | X | X |  | X |  |  |  |  | X |  |
| MUD 3 | X | X | X | X | X | X | X | X | X | X | X | X | X | X | X | X |
| MUD4 |  |  |  |  |  |  |  |  |  |  |  |  |  |  |  |  |
| MMUD 1 |  | X |  |  | X | X | X | X |  | X |  |  |  |  |  |  |
| MMUD 2 | X | X | X | X |  |  |  | X | X |  | X | X | X | X | X | X |
| MMUD 3 |  | X | X | X | X | X | X | X | X | X | X | X | X |  | X | X |
| qRT-PCR ANOVA<br>*p-value* | 0.008 | 0.0143 | 0.029 | 0.0414 | 0.093 | 0.11 | 0.126 | 0.135 | 0.158<br>6 | 0.3 | 0.14 | 0.34 | 0.306 | 0.002<br>7 | 0.052 | 0.216 |
| RNA-seq *p<sub>adj</sub>* | < 0.05 | < 0.05 | < 0.05 | < 0.05 | <<br>0.05 | < 0.05 | < 0.05 | < 0.05 | < 0.05 | <<br>0.05 | < 0.1 | < 0.1 | < 0.1 | > 0.1 | > 0.1 | > 0.1 |

X: indicates patient sample was analyzed for expression of the transcript listed in the column title.

**Supplemental Table 8.** Confirmation cohort patients tested for expression of each lncRNA gene.

|  | Linc00402 | RP11-348F1.3 | RP5-1185I7.1 | Linc01395 | RP11-730K11.1 | Linc00824 | AC093642.4 | CHRM3-AS2 | KRT73-AS1 | Linc01422 | RP11-357H14.17 | RP11-489E7.4 | RP11-161M6.2 | RP11-757F18.5 |
| --- | --- | --- | --- | --- | --- | --- | --- | --- | --- | --- | --- | --- | --- | --- |
| C-Auto 1 | X | X | X | X | X | X | X | X | X | X | X | X | X | X |
| C-Auto 2 | X | X |  | X | X |  | X | X | X | X | X | X | X | X |
| C-Auto 3 | X | X | X | X | X | X | X | X | X | X | X | X | X | X |
| C-Auto 4 | X | X | X | X | X | X | X | X | X | X | X | X | X | X |
| C-Auto 5 | X | X | X | X | X |  | X | X | X | X | X | X | X | X |
| C-Auto 6 | X | X | X |  | X | X | X | X |  |  |  | X |  |  |
| C-Auto 7 | X | X | X | X | X | X | X | X | X | X | X | X | X | X |
| C-Auto 8 | X | X | X | X | X |  | X |  | X |  |  |  |  | X |
| C-Auto 9 | X | X | X | X | X | X | X | X | X | X | X |  | X | X |
| C-Auto 10 | X | X | X | X | X | X | X | X | X |  | X |  | X | X |
| C-Auto 11 | X | X | X | X |  | X | X | X |  | X | X | X | X | X |
| C-Auto 12 |  |  | X | X |  |  |  |  | X | X | X |  |  | X |
| C-Auto 12 |  |  |  | X |  |  | X | X | X |  |  |  | X | X |
| C-Auto 13 |  |  |  | X |  |  | X |  | X |  |  |  |  | X |
| C-Auto 14 |  |  |  |  | X | X |  |  | X | X |  |  | X | X |
| C-MUD 1 | X | X | X | X |  | X | X | X | X | X | X | X | X | X |
| C-MUD 2 | X | X | X | X | X | X | X | X |  |  | X | X | X | X |
| C-MUD 3 | X |  | X | X | X | X |  | X | X | X | X |  | X | X |
| C-MUD 4 | X | X | X | X | X | X | X | X | X | X | X | X | X | X |
| C-MUD 5 | X | X | X | X | X |  | X | X | X | X |  | X | X | X |
| C-MUD 6 | X | X | X | X | X | X | X | X | X | X | X | X | X | X |
| C-MUD 7 | X |  |  |  | X |  | X | X |  | X |  |  | X | X |
| C-MUD 8 | X | X | X | X |  | X | X | X | X |  | X | X | X | X |
| C-MUD 9 |  | X | X | X | X | X |  |  | X | X | X | X |  |  |
| C-MUD 10 |  |  | X | X | X |  | X |  |  |  |  |  |  |  |
| C-MUD 11 |  |  |  | X | X |  | X | X | X |  |  |  | X | X |
| C-MMUD 1 | X | X | X |  | X | X | X | X |  | X |  | X | X | X |
| C-MMUD 2 | X |  | X |  | X | X |  |  |  | X |  |  |  |  |
| C-MMUD 3 | X | X | X | X | X | X | X | X | X | X | X | X | X | X |
| C-MMUD 4 | X | X | X | X |  | X | X | X | X | X | X |  | X |  |
| C-MMUD 5 | X | X | X | X | X | X | X | X | X | X | X |  | X | X |
| C-MMUD 6 | X | X | X | X | X | X | X | X | X | X | X |  | X | X |
| C-MMUD 7 | X | X | X | X |  | X | X | X |  | X | X |  | X | X |
| C-MMUD 8 | X | X | X | X |  |  | X | X | X |  |  |  | X | X |
| C-MMUD 9 | X | X | X | X | X |  | X |  | X |  |  |  |  | X |
| C-MMUD 10 |  | X |  | X | X | X | X | X | X | X | X |  |  | X |
| C-MMUD 11 |  |  | X | X |  |  |  |  | X |  |  |  |  | X |
| C-MMUD 12 |  |  | X | X | X |  | X | X | X | X |  |  |  |  |
| C-MMUD 13 |  |  | X | X | X |  |  |  | X | X | X |  |  | X |
| C-MMUD 14 |  |  | X | X | X | X | X | X | X | X | X | X | X | X |
| C-MMUD 15 |  |  |  | X | X |  | X | X | X |  |  |  | X | X |
| C-MMUD 16 | X | X | X |  | X | X | X | X | X | X | X | X | X | X |
| C-MMUD 17 |  |  | X | X | X |  | X |  |  |  |  |  |  |  |
| ANOVA<br>p-value | 0.0084 | 0.0366 | 0.0237 | 0.0905 | 0.0039 | 0.072 | > 0.1 | > 0.1 | > 0.1 | > 0.1 | > 0.1 | > 0.1 | > 0.1 | > 0.1 |

X: indicates patient sample was analyzed for expression of the transcript listed in the column title.

**Supplemental Table 9.** Confirmation cohort patients tested for expression of each protein coding gene.

|  | CNN3 | GPRASP1 | PARK2 | SV2A | TXNRD3 | VCAM1 | SFRP5 | PRSS50 | IL23-R | NR3C2 | ERAP2 | FLT4 |
| --- | --- | --- | --- | --- | --- | --- | --- | --- | --- | --- | --- | --- |
| C-Auto 1 | X | X | X | X | X | X | X | X |  | X | X | X |
| C-Auto 2 | X | X | X | X | X | X | X | X |  | X | X | X |
| C-Auto 3 | X | X | X | X | X | X | X | X |  | X | X | X |
| C-Auto 4 | X | X | X | X |  | X | X | X |  | X |  | X |
| C-Auto 5 |  | X | X | X | X | X | X | X |  | X | X |  |
| C-Auto 6 | X |  | X |  | X | X |  | X |  | X | X | X |
| C-Auto 7 | X | X | X | X | X | X | X | X |  | X | X | X |
| C-Auto 8 | X | X | X | X | X |  | X | X |  | X | X | X |
| C-Auto 9 |  | X | X | X | X | X | X | X | X | X | X | X |
| C-Auto 10 | X | X | X | X | X |  | X |  | X | X |  |  |
| C-Auto 11 | X | X | X | X | X | X | X | X | X | X |  | X |
| C-Auto 12 |  |  |  |  |  | X | X |  |  |  | X |  |
| C-Auto 12 |  | X |  | X |  | X | X | X | X | X | X |  |
| C-Auto 13 |  | X |  | X |  | X |  |  |  | X | X |  |
| C-Auto 14 | X |  |  |  | X | X |  | X |  | X | X | X |
| C-MUD 1 | X | X | X | X |  | X | X | X |  | X | X | X |
| C-MUD 2 | X | X | X | X | X | X | X | X | X | X | X | X |
| C-MUD 3 | X |  |  | X | X | X | X |  |  | X | X | X |
| C-MUD 4 | X | X | X | X | X |  | X | X |  | X | X | X |
| C-MUD 5 | X | X | X | X | X |  | X | X |  | X | X | X |
| C-MUD 6 |  | X |  | X |  |  | X |  |  | X | X |  |
| C-MUD 7 |  |  |  |  |  | X |  | X |  | X | X | X |
| C-MUD 8 | X | X | X | X | X | X | X | X |  | X | X |  |
| C-MUD 9 | X | X | X | X | X | X | X | X |  | X |  | X |
| C-MUD 10 |  |  |  |  |  |  | X | X | X |  |  |  |
| C-MUD 11 |  |  |  |  |  |  | X | X | X |  | X |  |
| C-MMUD 1 | X | X | X | X | X | X |  |  |  | X | X | X |
| C-MMUD 2 | X |  | X | X | X | X |  |  |  |  | X | X |
| C-MMUD 3 | X | X | X | X |  |  | X | X |  | X | X | X |
| C-MMUD 4 | X | X | X | X | X | X | X | X | X | X | X |  |
| C-MMUD 5 | X | X | X | X | X | X | X | X | X | X | X | X |
| C-MMUD 6 | X | X | X | X | X | X | X | X | X | X | X | X |
| C-MMUD 7 | X | X | X | X |  | X | X | X | X | X | X | X |
| C-MMUD 8 | X | X | X |  | X | X | X | X |  | X |  |  |
| C-MMUD 9 | X | X | X | X |  | X | X | X |  | X | X | X |
| C-MMUD 10 |  | X |  | X |  | X | X | X |  | X | X |  |
| C-MMUD 11 |  |  |  |  |  |  |  | X |  |  |  |  |
| C-MMUD 12 |  |  |  |  |  |  | X |  | X |  | X |  |
| C-MMUD 13 |  |  |  |  |  | X | X | X |  |  |  |  |
| C-MMUD 14 |  | X |  | X |  |  | X | X |  | X | X |  |
| C-MMUD 15 |  |  |  |  |  |  | X | X |  |  | X |  |
| C-MMUD 16 | X | X | X | X | X | X | X | X |  | X | X | X |
| C-MMUD 17 |  |  |  |  |  | X |  | X | X | X |  | X |
| ANOVA<br>p-value | 0.0021 | 0.0087 | 0.0136 | 0.052 | 0.001 | 0.0013 | 0.0715 | > 0.1 | > 0.1 | > 0.1 | > 0.1 | > 0.1 |

X: indicates patient sample was analyzed for expression of the transcript listed in the column title.

**Supplemental Table 10.**
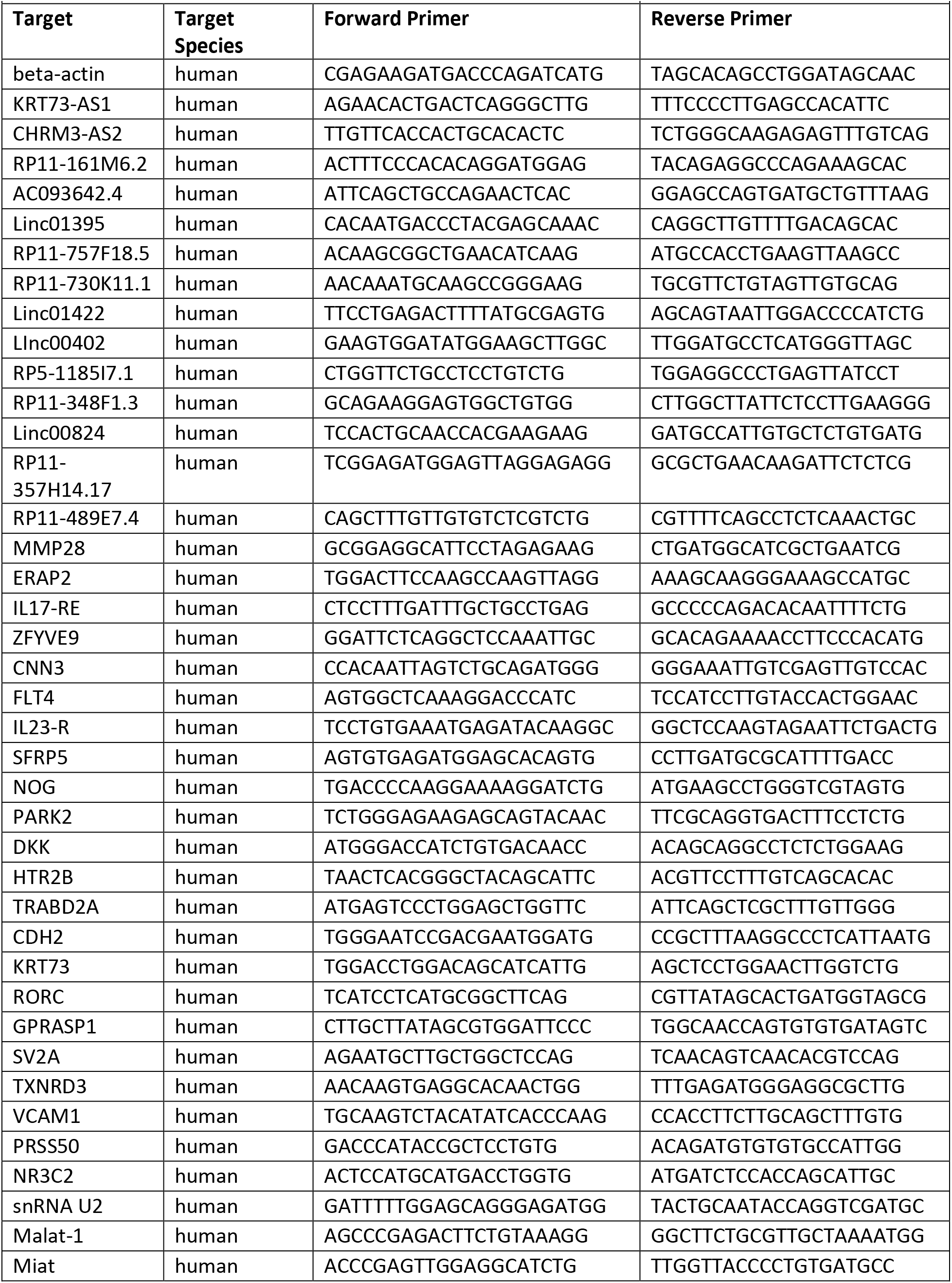
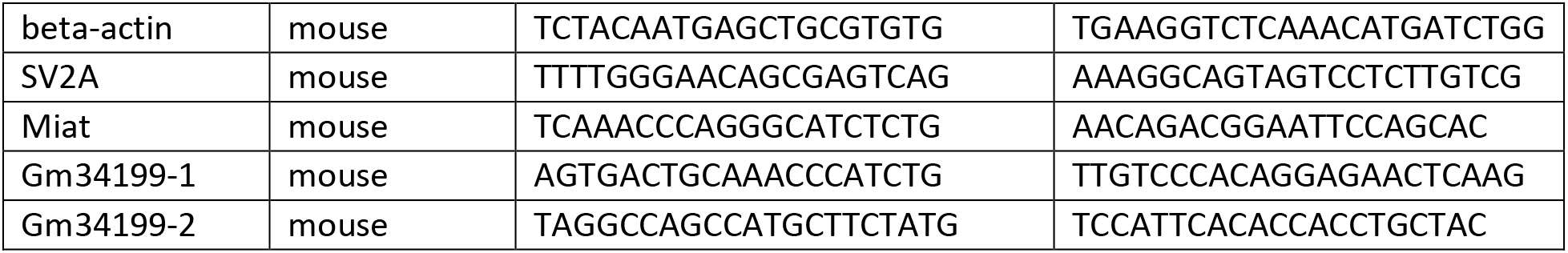
qPCR Primer Sequences

