## Supplemental Table 1 for "RNA-seq of Human T-Cells After Hematopoietic Stem Cell Transplantation Identifies Linc00402 as a Novel Regulator of T-Cell Alloimmunity"

| GOMFID | Pvalue | OddsRatio | ExpCount | Count | Size | Term |
|---|---|---|---|---|---|---|
| GO:00037 | 5.09E-21 | 5.318613 | 22.866 | 71 | 144 | structural constituent of ribosome |
| GO:00048 | 3.62E-11 | 1.849669 | 115.6003 | 182 | 728 | receptor activity |
| GO:00051 | 1.10E-09 | 2.05156 | 66.3749 | 114 | 418 | structural molecule activity |
| GO:00380 | 1.83E-08 | 1.801165 | 89.2409 | 139 | 562 | signaling receptor activity |
| GO:00048 | 2.21E-07 | 1.783106 | 77.33153 | 120 | 487 | transmembrane signaling receptor activity |
| GO:00083 | 3.59E-07 | 19.5349 | 2.223083 | 11 | 14 | signaling pattern recognition receptor activity |
| GO:00381 | 3.59E-07 | 19.5349 | 2.223083 | 11 | 14 | pattern recognition receptor activity |
| GO:00048 | 6.88E-06 | 1.50117 | 128.3036 | 174 | 808 | signal transducer activity |
| GO:00600 | 6.88E-06 | 1.50117 | 128.3036 | 174 | 808 | molecular transducer activity |
| GO:00055 | 6.97E-06 | 1.702798 | 68.43919 | 103 | 431 | calcium ion binding |
| GO:00198 | 2.45E-05 | 11.97304 | 2.064291 | 9 | 13 | immunoglobulin binding |
| GO:00380 | 4.22E-05 | 3.84197 | 6.82804 | 18 | 43 | cargo receptor activity |
| GO:00431 | 0.000231 | 1.217694 | 505.4338 | 568 | 3183 | cation binding |
| GO:00468 | 0.000321 | 1.21261 | 499.2409 | 560 | 3144 | metal ion binding |
| GO:00508 | 0.000367 | 3.729001 | 5.398916 | 14 | 34 | extracellular matrix binding |
| GO:00302 | 0.000459 | 9.302231 | 1.746708 | 7 | 11 | lipoprotein particle receptor activity |
| GO:00198 | 0.000865 | 10.62643 | 1.429125 | 6 | 9 | IgG binding |
| GO:00198 | 0.001301 | 3.548282 | 4.763749 | 12 | 30 | rRNA binding |
| GO:00047 | 0.001345 | 2.959016 | 6.669249 | 15 | 42 | non-membrane spanning protein tyrosine kinase activity |
| GO:00422 | 0.001471 | 1.937868 | 19.69016 | 33 | 124 | peptide binding |
| GO:00050 | 0.001591 | 13.27717 | 1.111541 | 5 | 7 | low-density lipoprotein receptor activity |
| GO:00018 | 0.002768 | 21.23411 | 0.793958 | 4 | 5 | lipopolysaccharide receptor activity |
| GO:00332 | 0.002954 | 1.835989 | 20.48412 | 33 | 129 | amide binding |
| GO:00432 | 0.002997 | 3.987745 | 3.334624 | 9 | 21 | laminin binding |
| GO:00055 | 0.003048 | 1.97362 | 15.87916 | 27 | 100 | glycosaminoglycan binding |
| GO:00301 | 0.003565 | 6.374551 | 1.746708 | 6 | 11 | low-density lipoprotein particle binding |
| GO:00055 | 0.003792 | 1.838809 | 19.21379 | 31 | 121 | calmodulin binding |
| GO:00049 | 0.003998 | Inf | 0.476375 | 3 | 3 | N-formyl peptide receptor activity |
| GO:00302 | 0.003998 | Inf | 0.476375 | 3 | 3 | very-low-density lipoprotein particle receptor activity |
| GO:00310 | 0.003998 | Inf | 0.476375 | 3 | 3 | troponin T binding |
| GO:00706 | 0.003998 | Inf | 0.476375 | 3 | 3 | vitamin D response element binding |
| GO:00302 | 0.004229 | 1.626014 | 30.64679 | 45 | 193 | carbohydrate binding |
| GO:00050 | 0.004729 | 2.200486 | 10.32146 | 19 | 65 | Rho guanyl-nucleotide exchange factor activity |
| GO:00150 | 0.004749 | 2.05728 | 12.54454 | 22 | 79 | calcium ion transmembrane transporter activity |
| GO:00323 | 0.007216 | 6.637228 | 1.429125 | 5 | 9 | MHC class II receptor activity |
| GO:00167 | 0.007258 | 10.61597 | 0.95275 | 4 | 6 | oxidoreductase activity, oxidizing metal ions, NAD or NADP as acceptor |
| GO:00428 | 0.007258 | 10.61597 | 0.95275 | 4 | 6 | peptidoglycan binding |
| GO:00718 | 0.007656 | 4.132209 | 2.540666 | 7 | 16 | lipoprotein particle binding |
| GO:00718 | 0.007656 | 4.132209 | 2.540666 | 7 | 16 | protein-lipid complex binding |
| GO:00038 | 0.007934 | 2.241317 | 8.574748 | 16 | 54 | antigen binding |
| GO:00611 | 0.008164 | 1.757697 | 18.57862 | 29 | 117 | peptidase regulator activity |
| GO:00228 | 0.008885 | 1.536423 | 32.71108 | 46 | 206 | substrate-specific channel activity |
| GO:00050 | 0.009292 | 1.777228 | 17.1495 | 27 | 108 | Ras guanyl-nucleotide exchange factor activity |

| GOCCID | Pvalue | OddsRatio | ExpCount | Count | Size | Term |
|---|---|---|---|---|---|---|
| GO:00226 | 3.04E-37 | 17.32993 | 14.34583 | 69 | 91 | cytosolic ribosome |
| GO:00443 | 7.50E-27 | 7.168395 | 20.65169 | 74 | 131 | ribosomal subunit |
| GO:00226 | 1.15E-23 | 21.79503 | 7.882325 | 40 | 50 | cytosolic large ribosomal subunit |
| GO:00444 | 3.73E-21 | 4.610226 | 28.06108 | 81 | 178 | cytosolic part |
| GO:00058 | 1.93E-18 | 3.931925 | 31.05636 | 82 | 197 | ribosome |
| GO:00226 | 1.71E-16 | 17.45937 | 5.990567 | 29 | 38 | cytosolic small ribosomal subunit |
| GO:00159 | 6.29E-16 | 7.436337 | 11.1929 | 41 | 71 | large ribosomal subunit |
| GO:00719 | 8.26E-13 | 1.479784 | 468.3677 | 594 | 2971 | cell periphery |
| GO:00058 | 3.09E-12 | 1.469368 | 456.7019 | 578 | 2897 | plasma membrane |
| GO:00159 | 5.27E-12 | 6.387626 | 9.616436 | 33 | 61 | small ribosomal subunit |
| GO:00056 | 9.88E-10 | 1.8057 | 105.3079 | 164 | 668 | extracellular space |
| GO:00312 | 8.97E-09 | 1.662501 | 131.4772 | 192 | 834 | intrinsic component of plasma membrane |
| GO:00059 | 9.76E-09 | 2.102645 | 53.44216 | 94 | 339 | focal adhesion |
| GO:00059 | 1.57E-08 | 2.076604 | 53.9151 | 94 | 342 | cell-substrate adherens junction |
| GO:00300 | 2.50E-08 | 2.051186 | 54.38804 | 94 | 345 | cell-substrate junction |
| GO:00058 | 2.53E-08 | 1.653162 | 125.6443 | 183 | 797 | integral component of plasma membrane |
| GO:00306 | 2.97E-08 | 3.354322 | 16.07994 | 39 | 102 | endocytic vesicle membrane |
| GO:00444 | 4.19E-08 | 1.470455 | 231.898 | 305 | 1471 | plasma membrane part |
| GO:00701 | 5.79E-07 | 1.835741 | 63.37389 | 101 | 402 | anchoring junction |
| GO:00059 | 5.79E-07 | 1.847989 | 61.79743 | 99 | 392 | adherens junction |
| GO:00003 | 6.57E-07 | 1.83542 | 62.7433 | 100 | 398 | lytic vacuole |
| GO:00057 | 6.57E-07 | 1.83542 | 62.7433 | 100 | 398 | lysosome |
| GO:00301 | 1.01E-06 | 2.326307 | 28.37637 | 54 | 180 | endocytic vesicle |
| GO:00057 | 1.13E-06 | 1.764487 | 69.8374 | 108 | 443 | vacuole |
| GO:00055 | 1.20E-06 | 1.307044 | 455.2831 | 538 | 2888 | extracellular region |
| GO:00099 | 2.45E-06 | 1.723274 | 71.72916 | 109 | 455 | cell surface |
| GO:00426 | 5.04E-06 | 13.42125 | 2.207051 | 10 | 14 | MHC class II protein complex |
| GO:00444 | 9.49E-06 | 1.288278 | 404.048 | 476 | 2563 | extracellular region part |
| GO:00312 | 1.14E-05 | 1.892535 | 42.7222 | 70 | 271 | cell leading edge |
| GO:00444 | 1.95E-05 | 1.81993 | 46.50572 | 74 | 295 | vacuolar part |
| GO:00300 | 2.27E-05 | 1.466189 | 127.2207 | 170 | 807 | cell junction |
| GO:00310 | 8.55E-05 | 1.883634 | 34.20929 | 56 | 217 | extracellular matrix |
| GO:00310 | 0.00011 | 3.457413 | 7.251739 | 18 | 46 | platelet alpha granule |
| GO:00319 | 0.000121 | 1.24027 | 422.335 | 485 | 2679 | vesicle |
| GO:00306 | 0.000129 | 4.65496 | 4.414102 | 13 | 28 | clathrin-coated endocytic vesicle membrane |
| GO:00453 | 0.000134 | 2.749792 | 11.1929 | 24 | 71 | phagocytic vesicle |
| GO:00312 | 0.000164 | 1.214861 | 546.4028 | 613 | 3466 | intrinsic component of membrane |
| GO:00325 | 0.000179 | 3.838236 | 5.675274 | 15 | 36 | trans-Golgi network membrane |
| GO:00319 | 0.000192 | 1.233975 | 411.9303 | 472 | 2613 | membrane-bounded vesicle |
| GO:00300 | 0.000218 | 2.134547 | 20.0211 | 36 | 127 | lamellipodium |
| GO:00017 | 0.00023 | 2.101749 | 20.80934 | 37 | 132 | ruffle |
| GO:00985 | 0.00027 | 1.782365 | 35.62811 | 56 | 226 | side of membrane |
| GO:00453 | 0.00034 | 3.759908 | 5.359981 | 14 | 34 | clathrin-coated endocytic vesicle |
| GO:00057 | 0.000467 | 1.707009 | 38.78104 | 59 | 246 | vacuolar membrane |
| GO:00312 | 0.000477 | 2.23177 | 15.607 | 29 | 99 | leading edge membrane |
| GO:00055 | 0.000649 | 1.85678 | 26.48461 | 43 | 168 | proteinaceous extracellular matrix |
| GO:00312 | 0.000727 | 2.319882 | 13.08466 | 25 | 83 | anchored component of membrane |
| GO:00160 | 0.000825 | 1.187641 | 535.9981 | 594 | 3400 | integral component of membrane |
| GO:00444 | 0.000914 | 1.513007 | 60.6939 | 84 | 385 | cytoplasmic vesicle part |
| GO:00055 | 0.000938 | 3.09864 | 6.463506 | 15 | 41 | collagen trimer |
| GO:00057 | 0.000989 | 1.694374 | 34.99752 | 53 | 222 | lysosomal membrane |
| GO:00444 | 0.00127 | 1.169244 | 662.1153 | 721 | 4200 | membrane part |
| GO:00426 | 0.001465 | 4.126088 | 3.625869 | 10 | 23 | MHC protein complex |
| GO:00306 | 0.001587 | 2.490527 | 9.45879 | 19 | 60 | clathrin-coated vesicle membrane |
| GO:00429 | 0.001674 | 1.277734 | 174.83 | 210 | 1109 | cell projection |
| GO:00098 | 0.002047 | 1.781463 | 24.7505 | 39 | 157 | external side of plasma membrane |
| GO:00314 | 0.002492 | 1.305074 | 131.3195 | 161 | 833 | cytoplasmic vesicle |
| GO:00160 | 0.002629 | 1.316359 | 120.5996 | 149 | 765 | cytoplasmic membrane-bounded vesicle |
| GO:00092 | 0.002692 | 21.41463 | 0.788232 | 4 | 5 | cell outer membrane |
| GO:00303 | 0.002692 | 21.41463 | 0.788232 | 4 | 5 | cell envelope |
| GO:00444 | 0.002692 | 21.41463 | 0.788232 | 4 | 5 | external encapsulating structure part |
| GO:00301 | 0.003269 | 1.642755 | 31.05636 | 46 | 197 | secretory granule |
| GO:00306 | 0.003356 | 1.496204 | 48.71277 | 67 | 309 | cytoplasmic vesicle membrane |
| GO:00303 | 0.003569 | 8.925606 | 1.261172 | 5 | 8 | external encapsulating structure |
| GO:00325 | 0.003615 | 2.268282 | 10.08938 | 19 | 64 | ruffle membrane |
| GO:00016 | 0.003913 | Inf | 0.472939 | 3 | 3 | granular component |
| GO:00125 | 0.003977 | 1.473845 | 50.76217 | 69 | 322 | vesicle membrane |

| | | | | | | |
|---|---|---|---|---|---|---|
| GO:00427 | 0.004326 | 3.105236 | 4.729395 | 11 | 30 | presynaptic membrane |
| GO:00432 | 0.00458 | 1.183429 | 332.3188 | 373 | 2108 | extracellular organelle |
| GO:00650 | 0.00458 | 1.183429 | 332.3188 | 373 | 2108 | extracellular membrane-bounded organelle |
| GO:00700 | 0.00458 | 1.183429 | 332.3188 | 373 | 2108 | extracellular vesicular exosome |
| GO:00310 | 0.005491 | 2.798758 | 5.517627 | 12 | 35 | platelet alpha granule lumen |
| GO:00432 | 0.006084 | 2.32134 | 8.355264 | 16 | 53 | lysosomal lumen |
| GO:00057 | 0.007067 | 10.7063 | 0.945879 | 4 | 6 | secondary lysosome |
| GO:00715 | 0.008121 | 3.216098 | 3.783516 | 9 | 24 | integral component of lumenal side of endoplasmic reticulum membrane |
| GO:00985 | 0.008121 | 3.216098 | 3.783516 | 9 | 24 | lumenal side of endoplasmic reticulum membrane |
| GO:00985 | 0.008121 | 3.216098 | 3.783516 | 9 | 24 | lumenal side of membrane |

| GOBPID | Pvalue | OddsRatio | ExpCount | Count | Size | Term |
|---|---|---|---|---|---|---|
| GO:00064 | 1.01E-36 | 16.03 | 14.9127 | 70 | 94 | translational termination |
| GO:00066 | 1.07E-28 | 9.66 | 16.6578 | 67 | 105 | SRP-dependent cotranslational protein targeting to membrane |
| GO:00066 | 5.62E-28 | 9.17 | 16.9751 | 67 | 107 | cotranslational protein targeting to membrane |
| GO:00450 | 1.25E-27 | 8.95 | 17.1337 | 67 | 108 | protein targeting to ER |
| GO:00725 | 2.76E-27 | 8.74 | 17.2924 | 67 | 109 | establishment of protein localization to endoplasmic reticulum |
| GO:00064 | 4.20E-27 | 7.99 | 18.7202 | 70 | 118 | translational elongation |
| GO:00001 | 6.20E-26 | 7.76 | 18.4029 | 68 | 116 | nuclear-transcribed mRNA catabolic process, nonsense-mediated decay |
| GO:00436 | 1.15E-23 | 5.30 | 26.1765 | 81 | 165 | cellular protein complex disassembly |
| GO:00709 | 1.29E-23 | 6.64 | 19.6721 | 68 | 124 | protein localization to endoplasmic reticulum |
| GO:00432 | 1.83E-21 | 4.53 | 29.5081 | 84 | 186 | protein complex disassembly |
| GO:00190 | 1.32E-20 | 4.94 | 24.7487 | 74 | 156 | viral transcription |
| GO:00329 | 2.72E-20 | 4.21 | 31.0946 | 85 | 196 | macromolecular complex disassembly |
| GO:00066 | 4.01E-19 | 4.46 | 26.4938 | 75 | 167 | protein targeting to membrane |
| GO:00440 | 5.96E-19 | 4.30 | 27.763 | 77 | 175 | multi-organism metabolic process |
| GO:00190 | 1.17E-18 | 4.40 | 26.3352 | 74 | 166 | viral gene expression |
| GO:00064 | 2.23E-18 | 4.39 | 26.0179 | 73 | 164 | translational initiation |
| GO:00009 | 1.82E-17 | 4.00 | 28.5562 | 76 | 180 | nuclear-transcribed mRNA catabolic process |
| GO:00064 | 6.90E-17 | 3.78 | 30.3013 | 78 | 191 | mRNA catabolic process |
| GO:00224 | 6.77E-16 | 2.68 | 57.2711 | 118 | 361 | cellular component disassembly |
| GO:00064 | 1.07E-14 | 3.23 | 34.5848 | 81 | 218 | RNA catabolic process |
| GO:00901 | 7.40E-12 | 2.57 | 44.5794 | 90 | 281 | establishment of protein localization to membrane |
| GO:00096 | 4.47E-11 | 1.85 | 114.225 | 180 | 720 | response to wounding |
| GO:00726 | 3.85E-10 | 2.22 | 54.8914 | 100 | 346 | protein localization to membrane |
| GO:00096 | 4.58E-10 | 1.57 | 215.917 | 297 | 1361 | response to external stimulus |
| GO:00069 | 9.83E-10 | 2.07 | 64.7275 | 112 | 408 | inflammatory response |
| GO:19025 | 1.40E-09 | 2.08 | 62.8237 | 109 | 396 | single-organism localization |
| GO:19025 | 1.40E-09 | 2.08 | 62.8237 | 109 | 396 | single-organism cellular localization |
| GO:00069 | 1.71E-09 | 1.61 | 171.02 | 242 | 1078 | defense response |
| GO:00190 | 3.27E-09 | 2.27 | 45.8486 | 85 | 289 | viral life cycle |
| GO:00022 | 6.10E-09 | 3.36 | 17.7683 | 43 | 112 | myeloid leukocyte activation |
| GO:00447 | 7.32E-09 | 1.34 | 645.529 | 754 | 4069 | single-multicellular organism process |
| GO:00096 | 8.56E-09 | 2.26 | 43.7862 | 81 | 276 | response to bacterium |
| GO:00427 | 3.51E-08 | 3.33 | 16.1819 | 39 | 102 | defense response to bacterium |
| GO:00023 | 4.47E-08 | 1.43 | 286.99 | 366 | 1809 | immune system process |
| GO:00325 | 5.62E-08 | 1.32 | 667.74 | 770 | 4209 | multicellular organismal process |
| GO:00517 | 9.69E-08 | 1.45 | 244.79 | 317 | 1543 | multi-organism process |
| GO:00022 | 1.41E-07 | 2.43 | 29.6668 | 58 | 187 | adaptive immune response |
| GO:19030 | 1.94E-07 | 2.19 | 38.5509 | 70 | 243 | regulation of response to wounding |
| GO:00420 | 2.00E-07 | 1.76 | 82.8131 | 127 | 522 | wound healing |
| GO:00903 | 2.99E-07 | 5.63 | 6.18718 | 20 | 39 | phagosome maturation |
| GO:00512 | 4.18E-07 | 1.42 | 241.142 | 309 | 1520 | regulation of multicellular organismal process |
| GO:00024 | 4.38E-07 | 2.46 | 26.3352 | 52 | 166 | adaptive immune response based on somatic recombination of immune receptors built from immunoglobulin superfamily domains |
| GO:00069 | 4.96E-07 | 3.20 | 14.4368 | 34 | 91 | humoral immune response |
| GO:00448 | 7.60E-07 | 1.67 | 92.9664 | 137 | 586 | single-organism membrane organization |
| GO:00507 | 9.17E-07 | 4.15 | 8.72552 | 24 | 55 | positive regulation of inflammatory response |
| GO:19030 | 1.32E-06 | 3.62 | 10.6293 | 27 | 67 | positive regulation of response to wounding |
| GO:00075 | 1.36E-06 | 1.77 | 68.535 | 106 | 432 | hemostasis |
| GO:00075 | 1.37E-06 | 1.77 | 67.7417 | 105 | 427 | blood coagulation |
| GO:00069 | 1.57E-06 | 1.47 | 169.592 | 225 | 1069 | immune response |
| GO:00321 | 1.68E-06 | 2.57 | 21.0999 | 43 | 133 | positive regulation of response to external stimulus |
| GO:00508 | 1.73E-06 | 1.76 | 68.059 | 105 | 429 | coagulation |
| GO:00447 | 1.83E-06 | 1.32 | 407.402 | 485 | 2568 | single-organism transport |
| GO:00325 | 2.11E-06 | 1.28 | 578.264 | 664 | 3645 | developmental process |
| GO:00400 | 2.29E-06 | 1.47 | 165.15 | 219 | 1041 | locomotion |
| GO:00017 | 2.71E-06 | 1.56 | 113.59 | 159 | 716 | cell activation |
| GO:00508 | 2.74E-06 | 1.68 | 80.1161 | 119 | 505 | regulation of body fluid levels |
| GO:00421 | 3.15E-06 | 5.06 | 5.86989 | 18 | 37 | macrophage activation |
| GO:00164 | 3.36E-06 | 1.55 | 117.398 | 163 | 740 | cell migration |
| GO:00725 | 3.49E-06 | 1.74 | 67.4244 | 103 | 425 | establishment of protein localization to organelle |
| GO:00488 | 3.51E-06 | 1.52 | 126.917 | 174 | 800 | cell motility |
| GO:00516 | 3.51E-06 | 1.52 | 126.917 | 174 | 800 | localization of cell |
| GO:00024 | 3.68E-06 | 5.34 | 5.39395 | 17 | 34 | humoral immune response mediated by circulating immunoglobulin |
| GO:00197 | 3.78E-06 | 3.02 | 13.6435 | 31 | 86 | B cell mediated immunity |
| GO:00450 | 3.88E-06 | 1.56 | 109.148 | 153 | 688 | innate immune response |
| GO:00160 | 5.06E-06 | 3.03 | 13.1676 | 30 | 83 | immunoglobulin mediated immune response |
| GO:00447 | 5.66E-06 | 1.27 | 571.442 | 653 | 3602 | single-organism developmental process |
| GO:00024 | 6.94E-06 | 2.11 | 31.5705 | 56 | 199 | leukocyte mediated immunity |
| GO:00507 | 8.56E-06 | 2.18 | 28.0803 | 51 | 177 | regulation of inflammatory response |
| GO:00488 | 1.27E-05 | 1.26 | 510.205 | 586 | 3216 | anatomical structure development |
| GO:00324 | 1.32E-05 | 21.30 | 1.58646 | 8 | 10 | response to peptidoglycan |
| GO:00096 | 1.46E-05 | 1.60 | 82.8131 | 119 | 522 | response to biotic stimulus |
| GO:00508 | 1.49E-05 | 1.24 | 856.687 | 940 | 5400 | response to stimulus |
| GO:00069 | 1.61E-05 | 2.24 | 24.2728 | 45 | 153 | phagocytosis |
| GO:00022 | 1.95E-05 | 1.62 | 76.6259 | 111 | 483 | immune effector process |
| GO:00022 | 2.09E-05 | 2.07 | 29.6668 | 52 | 187 | response to molecule of bacterial origin |
| GO:00313 | 2.14E-05 | 1.93 | 37.2817 | 62 | 235 | positive regulation of defense response |
| GO:00488 | 2.50E-05 | 1.28 | 391.062 | 458 | 2465 | cellular developmental process |
| GO:00082 | 2.68E-05 | 1.37 | 205.129 | 257 | 1293 | cell proliferation |
| GO:00432 | 2.70E-05 | 1.60 | 78.8469 | 113 | 497 | response to external biotic stimulus |
| GO:00517 | 2.70E-05 | 1.60 | 78.8469 | 113 | 497 | response to other organism |
| GO:00650 | 2.95E-05 | 1.24 | 1198.25 | 1275 | 7553 | biological regulation |
| GO:00610 | 2.97E-05 | 1.49 | 113.114 | 153 | 713 | membrane organization |
| GO:00024 | 3.21E-05 | 3.76 | 7.2977 | 19 | 46 | myeloid leukocyte mediated immunity |
| GO:00444 | 4.13E-05 | 1.48 | 111.211 | 150 | 701 | symbiosis, encompassing mutualism through parasitism |
| GO:00444 | 4.13E-05 | 1.48 | 111.211 | 150 | 701 | interspecies interaction between organisms |
| GO:00321 | 4.16E-05 | 1.63 | 67.9004 | 99 | 428 | regulation of response to external stimulus |
| GO:00985 | 4.41E-05 | 1.82 | 42.0411 | 67 | 265 | defense response to other organism |
| GO:00072 | 4.48E-05 | 1.25 | 492.278 | 562 | 3103 | multicellular organismal development |
| GO:00022 | 4.60E-05 | 3.63 | 7.45635 | 19 | 47 | myeloid cell activation involved in immune response |
| GO:00069 | 4.68E-05 | 1.26 | 421.839 | 488 | 2659 | response to stress |
| GO:00301 | 4.74E-05 | 1.89 | 36.6472 | 60 | 231 | extracellular matrix organization |
| GO:00430 | 4.74E-05 | 1.89 | 36.6472 | 60 | 231 | extracellular structure organization |
| GO:00324 | 5.34E-05 | 2.05 | 27.6044 | 48 | 174 | response to lipopolysaccharide |
| GO:00650 | 5.48E-05 | 1.27 | 352.511 | 414 | 2222 | regulation of biological quality |
| GO:00422 | 5.96E-05 | 5.82 | 3.64885 | 12 | 23 | ribosomal small subunit biogenesis |
| GO:00082 | 6.25E-05 | 1.55 | 82.0198 | 115 | 517 | positive regulation of cell proliferation |
| GO:00509 | 6.26E-05 | 1.86 | 37.7577 | 61 | 238 | leukocyte migration |
| GO:00022 | 6.69E-05 | 2.19 | 21.8931 | 40 | 138 | cell activation involved in immune response |
| GO:00023 | 6.69E-05 | 2.19 | 21.8931 | 40 | 138 | leukocyte activation involved in immune response |
| GO:00326 | 6.80E-05 | 3.34 | 8.24958 | 20 | 52 | chemokine production |
| GO:00069 | 8.25E-05 | 1.35 | 186.567 | 233 | 1176 | cellular component movement |
| GO:00301 | 8.60E-05 | 1.26 | 363.616 | 424 | 2292 | cell differentiation |
| GO:00068 | 9.17E-05 | 1.61 | 65.0447 | 94 | 410 | endocytosis |
| GO:00069 | 9.25E-05 | 1.58 | 70.9146 | 101 | 447 | chemotaxis |
| GO:00423 | 9.25E-05 | 1.58 | 70.9146 | 101 | 447 | taxis |
| GO:00718 | 9.53E-05 | 1.35 | 188.788 | 235 | 1190 | protein complex subunit organization |
| GO:00330 | 9.58E-05 | 31.92 | 1.11052 | 6 | 7 | positive regulation of mast cell activation involved in immune response |
| GO:00433 | 9.58E-05 | 31.92 | 1.11052 | 6 | 7 | positive regulation of mast cell degranulation |
| GO:00603 | 0.0001 | 2.12 | 23.0036 | 41 | 145 | cell chemotaxis |
| GO:00066 | 0.0001 | 1.56 | 73.6116 | 104 | 464 | protein targeting |
| GO:00421 | 0.0001 | 1.38 | 153.886 | 196 | 970 | regulation of cell proliferation |
| GO:00030 | 0.00011 | 2.50 | 14.4368 | 29 | 91 | vascular process in circulatory system |
| GO:00017 | 0.00011 | 10.65 | 1.90375 | 8 | 12 | microglial cell activation |
| GO:00712 | 0.00011 | 2.41 | 15.8646 | 31 | 100 | cellular response to molecule of bacterial origin |
| GO:00025 | 0.00012 | 5.86 | 3.33156 | 11 | 21 | production of molecular mediator involved in inflammatory response |
| GO:00028 | 0.00012 | 5.86 | 3.33156 | 11 | 21 | regulation of myeloid leukocyte mediated immunity |
| GO:00330 | 0.00012 | 5.86 | 3.33156 | 11 | 21 | regulation of mast cell activation |
| GO:00485 | 0.00012 | 1.27 | 309.518 | 365 | 1951 | organ development |
| GO:00441 | 0.00013 | 7.99 | 2.37969 | 9 | 15 | growth involved in symbiotic interaction |
| GO:00441 | 0.00013 | 7.99 | 2.37969 | 9 | 15 | growth of symbiont involved in interaction with host |
| GO:00441 | 0.00013 | 7.99 | 2.37969 | 9 | 15 | growth of symbiont in host |
| GO:00327 | 0.00014 | 4.62 | 4.44208 | 13 | 28 | positive regulation of interleukin-6 production |
| GO:00508 | 0.00015 | 3.71 | 6.18718 | 16 | 39 | defense response to Gram-positive bacterium |
| GO:00430 | 0.00016 | 1.66 | 52.6704 | 78 | 332 | positive regulation of programmed cell death |
| GO:00453 | 0.00016 | 1.50 | 84.7168 | 116 | 534 | leukocyte activation |
| GO:00485 | 0.00018 | 1.48 | 92.6491 | 125 | 584 | hematopoietic or lymphoid organ development |
| GO:00712 | 0.00018 | 2.22 | 18.4029 | 34 | 116 | cellular response to biotic stimulus |
| GO:00469 | 0.00018 | 1.45 | 103.12 | 137 | 650 | secretion |
| GO:00512 | 0.00019 | 1.56 | 67.1071 | 95 | 423 | positive regulation of multicellular organismal process |
| GO:00083 | 0.00019 | 2.49 | 13.4849 | 27 | 85 | regulation of cell shape |
| GO:00975 | 0.00019 | 2.49 | 13.4849 | 27 | 85 | myeloid leukocyte migration |
| GO:00330 | 0.00019 | 12.42 | 1.58646 | 7 | 10 | positive regulation of mast cell activation |
| GO:00433 | 0.00019 | 12.42 | 1.58646 | 7 | 10 | positive regulation of leukocyte degranulation |
| GO:19033 | 0.00019 | 12.42 | 1.58646 | 7 | 10 | positive regulation of regulated secretory pathway |

| | | | | | | |
|---|---|---|---|---|---|---|
| GO:00323 | 0.00019 | 2.18 | 19.1961 | 35 | 121 | positive regulation of Rho GTPase activity |
| GO:00712 | 0.0002 | 2.39 | 14.9127 | 29 | 94 | cellular response to lipopolysaccharide |
| GO:00024 | 0.00021 | 1.99 | 25.2247 | 43 | 159 | lymphocyte mediated immunity |
| GO:00326 | 0.00021 | 2.80 | 10.1533 | 22 | 64 | regulation of tumor necrosis factor production |
| GO:00026 | 0.00021 | 1.48 | 90.4281 | 122 | 570 | positive regulation of immune system process |
| GO:00422 | 0.00022 | 1.24 | 381.543 | 439 | 2405 | response to chemical |
| GO:00305 | 0.00022 | 2.25 | 17.1337 | 32 | 108 | leukocyte chemotaxis |
| GO:00716 | 0.00023 | 3.07 | 8.24958 | 19 | 52 | granulocyte chemotaxis |
| GO:00507 | 0.00024 | 1.20 | 1143.99 | 1212 | 7211 | regulation of biological process |
| GO:00441 | 0.00024 | 8.52 | 2.06239 | 8 | 13 | regulation of growth of symbiont in host |
| GO:00441 | 0.00024 | 8.52 | 2.06239 | 8 | 13 | negative regulation of growth of symbiont in host |
| GO:00441 | 0.00024 | 8.52 | 2.06239 | 8 | 13 | modulation of growth of symbiont involved in interaction with host |
| GO:00441 | 0.00024 | 8.52 | 2.06239 | 8 | 13 | negative regulation of growth of symbiont involved in interaction with host |
| GO:00430 | 0.00024 | 1.64 | 51.7185 | 76 | 326 | positive regulation of apoptotic process |
| GO:00226 | 0.00025 | 1.42 | 109.941 | 144 | 693 | biological adhesion |
| GO:00064 | 0.00025 | 1.51 | 76.1499 | 105 | 480 | translation |
| GO:00457 | 0.00026 | 2.67 | 10.9466 | 23 | 69 | positive regulation of angiogenesis |
| GO:00300 | 0.00027 | 1.47 | 89.1589 | 120 | 562 | hemopoiesis |
| GO:00326 | 0.00027 | 2.73 | 10.312 | 22 | 65 | tumor necrosis factor production |
| GO:00487 | 0.00027 | 1.23 | 426.281 | 485 | 2687 | system development |
| GO:00432 | 0.00028 | 3.64 | 5.86989 | 15 | 37 | leukocyte degranulation |
| GO:00703 | 0.00029 | 2.18 | 18.0856 | 33 | 114 | ERK1 and ERK2 cascade |
| GO:00025 | 0.00029 | 1.44 | 98.0431 | 130 | 618 | immune system development |
| GO:00703 | 0.0003 | 2.24 | 16.6578 | 31 | 105 | regulation of ERK1 and ERK2 cascade |
| GO:00506 | 0.0003 | 2.32 | 15.23 | 29 | 96 | cytokine secretion |
| GO:00305 | 0.0003 | 3.24 | 7.13906 | 17 | 45 | neutrophil chemotaxis |
| GO:00018 | 0.00031 | 1.53 | 70.5973 | 98 | 445 | cytokine production |
| GO:00071 | 0.00031 | 1.42 | 109.624 | 143 | 691 | cell adhesion |
| GO:00026 | 0.00033 | 2.26 | 16.0232 | 30 | 101 | positive regulation of immune effector process |
| GO:00000 | 0.00033 | 15.96 | 1.26917 | 6 | 8 | ribosomal small subunit assembly |
| GO:00329 | 0.00036 | 1.45 | 91.5386 | 122 | 577 | secretion by cell |
| GO:00333 | 0.00037 | 1.46 | 89.0003 | 119 | 561 | protein localization to organelle |
| GO:00450 | 0.00041 | 3.13 | 7.2977 | 17 | 46 | regulated secretory pathway |
| GO:19902 | 0.00041 | 3.13 | 7.2977 | 17 | 46 | neutrophil migration |
| GO:00066 | 0.00044 | 2.61 | 10.6293 | 22 | 67 | icosanoid metabolic process |
| GO:19015 | 0.00044 | 2.61 | 10.6293 | 22 | 67 | fatty acid derivative metabolic process |
| GO:00022 | 0.00046 | 9.31 | 1.7451 | 7 | 11 | neutrophil activation involved in immune response |
| GO:00018 | 0.00046 | 1.54 | 62.8237 | 88 | 396 | regulation of cytokine production |
| GO:00300 | 0.00047 | 1.68 | 41.2479 | 62 | 260 | myeloid cell differentiation |
| GO:00610 | 0.00047 | 2.47 | 12.0571 | 24 | 76 | regulation of wound healing |
| GO:00433 | 0.00048 | 7.10 | 2.22104 | 8 | 14 | regulation of mast cell degranulation |
| GO:00303 | 0.00052 | 1.79 | 31.7291 | 50 | 200 | positive regulation of cell migration |
| GO:00703 | 0.00052 | 26.58 | 0.95187 | 5 | 6 | response to lipoteichoic acid |
| GO:00712 | 0.00052 | 26.58 | 0.95187 | 5 | 6 | cellular response to lipoteichoic acid |
| GO:20011 | 0.00052 | 26.58 | 0.95187 | 5 | 6 | regulation of interleukin-10 secretion |
| GO:00717 | 0.00055 | 2.55 | 10.7879 | 22 | 68 | tumor necrosis factor superfamily cytokine production |
| GO:00507 | 0.00055 | 3.16 | 6.82177 | 16 | 43 | regulation of phagocytosis |
| GO:00512 | 0.00058 | 1.76 | 33.4742 | 52 | 211 | positive regulation of cellular component movement |
| GO:00351 | 0.00058 | 2.42 | 12.2157 | 24 | 77 | regulation of tube size |
| GO:00508 | 0.00058 | 2.42 | 12.2157 | 24 | 77 | regulation of blood vessel size |
| GO:00109 | 0.00059 | 1.56 | 55.6847 | 79 | 351 | positive regulation of cell death |
| GO:00071 | 0.00061 | 1.19 | 630.458 | 692 | 3974 | cell communication |
| GO:00075 | 0.00062 | 4.00 | 4.44208 | 12 | 28 | embryo implantation |
| GO:00507 | 0.00062 | 4.00 | 4.44208 | 12 | 28 | positive regulation of phagocytosis |
| GO:00508 | 0.00062 | 1.57 | 54.0982 | 77 | 341 | regulation of cell activation |
| GO:00023 | 0.00063 | Inf | 0.63458 | 4 | 4 | MHC class II protein complex assembly |
| GO:00025 | 0.00063 | Inf | 0.63458 | 4 | 4 | peptide antigen assembly with MHC class II protein complex |
| GO:00605 | 0.00063 | Inf | 0.63458 | 4 | 4 | cartilage morphogenesis |
| GO:00712 | 0.00063 | Inf | 0.63458 | 4 | 4 | cellular response to peptidoglycan |
| GO:00433 | 0.00064 | 4.84 | 3.33156 | 10 | 21 | regulation of leukocyte degranulation |
| GO:19033 | 0.00064 | 4.84 | 3.33156 | 10 | 21 | regulation of regulated secretory pathway |
| GO:00507 | 0.00065 | 1.29 | 202.115 | 243 | 1274 | regulation of developmental process |
| GO:00485 | 0.00066 | 1.22 | 382.495 | 435 | 2411 | regulation of response to stimulus |
| GO:00975 | 0.00067 | 2.74 | 8.88416 | 19 | 56 | granulocyte migration |
| GO:00442 | 0.00067 | 1.28 | 204.97 | 246 | 1292 | cellular nitrogen compound catabolic process |
| GO:00323 | 0.00068 | 1.94 | 22.6863 | 38 | 143 | regulation of Rho GTPase activity |
| GO:00511 | 0.00068 | 1.19 | 607.613 | 668 | 3830 | localization |
| GO:00069 | 0.00069 | 3.65 | 5.07666 | 13 | 32 | complement activation |
| GO:00423 | 0.00074 | 3.20 | 6.34583 | 15 | 40 | vasoconstriction |
| GO:00080 | 0.00074 | 1.66 | 40.296 | 60 | 254 | blood circulation |
| GO:19013 | 0.00076 | 1.28 | 208.143 | 249 | 1312 | organic cyclic compound catabolic process |
| GO:00467 | 0.00077 | 1.28 | 204.494 | 245 | 1289 | heterocycle catabolic process |
| GO:19004 | 0.00077 | 5.32 | 2.85562 | 9 | 18 | regulation of glutamate receptor signaling pathway |
| GO:00015 | 0.00085 | 1.62 | 43.7862 | 64 | 276 | angiogenesis |
| GO:00447 | 0.00085 | 1.20 | 1346.74 | 1403 | 8489 | single-organism cellular process |
| GO:00025 | 0.00086 | 2.45 | 11.1052 | 22 | 70 | platelet degranulation |
| GO:00072 | 0.00088 | 2.33 | 12.533 | 24 | 79 | integrin-mediated signaling pathway |
| GO:00330 | 0.00089 | 6.08 | 2.37969 | 8 | 15 | regulation of mast cell activation involved in immune response |
| GO:00422 | 0.00089 | 6.08 | 2.37969 | 8 | 15 | ribosomal large subunit biogenesis |
| GO:00230 | 0.0009 | 1.18 | 620.781 | 680 | 3913 | signaling |
| GO:00447 | 0.0009 | 1.18 | 620.781 | 680 | 3913 | single organism signaling |
| GO:00327 | 0.0009 | 3.76 | 4.60073 | 12 | 29 | positive regulation of chemokine production |
| GO:00327 | 0.0009 | 3.76 | 4.60073 | 12 | 29 | positive regulation of tumor necrosis factor production |
| GO:00030 | 0.00091 | 1.64 | 40.6133 | 60 | 256 | circulatory system process |
| GO:00326 | 0.00093 | 2.49 | 10.4706 | 21 | 66 | regulation of interferon-gamma production |
| GO:00708 | 0.00093 | 1.24 | 265.732 | 310 | 1675 | cellular response to chemical stimulus |
| GO:00071 | 0.00094 | 1.18 | 571.283 | 629 | 3601 | signal transduction |
| GO:20001 | 0.00094 | 1.73 | 32.5224 | 50 | 205 | positive regulation of cell motility |
| GO:00027 | 0.00095 | 7.45 | 1.90375 | 7 | 12 | positive regulation of B cell mediated immunity |
| GO:00028 | 0.00095 | 7.45 | 1.90375 | 7 | 12 | positive regulation of immunoglobulin mediated immune response |
| GO:00706 | 0.00096 | 1.76 | 30.1427 | 47 | 190 | leukocyte proliferation |
| GO:00194 | 0.00096 | 1.27 | 205.288 | 245 | 1294 | aromatic compound catabolic process |
| GO:00326 | 0.00098 | 2.94 | 7.13906 | 16 | 45 | regulation of chemokine production |
| GO:00024 | 0.00105 | 2.21 | 14.1195 | 26 | 89 | antigen processing and presentation of peptide antigen via MHC class II |
| GO:00510 | 0.00105 | 1.41 | 91.3799 | 119 | 576 | positive regulation of developmental process |
| GO:00329 | 0.00109 | 1.77 | 28.7149 | 45 | 181 | mononuclear cell proliferation |
| GO:00508 | 0.00114 | 1.68 | 35.2194 | 53 | 222 | positive regulation of cell activation |
| GO:00096 | 0.00115 | 1.24 | 259.068 | 302 | 1633 | anatomical structure morphogenesis |
| GO:00326 | 0.00118 | 2.32 | 12.0571 | 23 | 76 | interferon-gamma production |
| GO:00096 | 0.00118 | 3.91 | 4.12479 | 11 | 26 | response to fungus |
| GO:00457 | 0.00125 | 1.96 | 19.5134 | 33 | 123 | regulation of angiogenesis |
| GO:00025 | 0.00126 | 2.17 | 14.2781 | 26 | 90 | antigen processing and presentation of peptide or polysaccharide antigen via MHC class II |
| GO:00422 | 0.00126 | 4.79 | 3.01427 | 9 | 19 | ribosome assembly |
| GO:00400 | 0.00129 | 1.68 | 34.5848 | 52 | 218 | positive regulation of locomotion |
| GO:00615 | 0.00129 | 2.84 | 7.2977 | 16 | 46 | myeloid cell development |
| GO:00507 | 0.0013 | 2.25 | 12.8503 | 24 | 81 | regulation of cytokine secretion |
| GO:00507 | 0.0013 | 1.17 | 1092.59 | 1152 | 6887 | regulation of cellular process |
| GO:00323 | 0.00136 | 1.65 | 37.1231 | 55 | 234 | positive regulation of Ras GTPase activity |
| GO:00508 | 0.00137 | 2.53 | 9.3601 | 19 | 59 | regulation of coagulation |
| GO:00801 | 0.00142 | 1.34 | 122.475 | 153 | 772 | regulation of response to stress |
| GO:00485 | 0.00145 | 1.27 | 186.409 | 223 | 1175 | positive regulation of response to stimulus |
| GO:00069 | 0.00147 | 2.60 | 8.72552 | 18 | 55 | smooth muscle contraction |
| GO:00301 | 0.00147 | 2.60 | 8.72552 | 18 | 55 | regulation of blood coagulation |
| GO:19000 | 0.00147 | 2.60 | 8.72552 | 18 | 55 | regulation of hemostasis |
| GO:00726 | 0.00158 | 13.29 | 1.11052 | 5 | 7 | interleukin-10 secretion |
| GO:00703 | 0.00159 | 2.30 | 11.5811 | 22 | 73 | positive regulation of ERK1 and ERK2 cascade |
| GO:00026 | 0.00163 | 1.53 | 50.7666 | 71 | 320 | regulation of leukocyte activation |
| GO:00421 | 0.00168 | 2.15 | 13.8022 | 25 | 87 | cytokine metabolic process |
| GO:00335 | 0.00174 | 2.23 | 12.3744 | 23 | 78 | unsaturated fatty acid metabolic process |
| GO:00508 | 0.00174 | 2.23 | 12.3744 | 23 | 78 | regulation of B cell activation |
| GO:00464 | 0.00175 | 2.86 | 6.82177 | 15 | 43 | icosanoid biosynthetic process |
| GO:19015 | 0.00175 | 2.86 | 6.82177 | 15 | 43 | fatty acid derivative biosynthetic process |
| GO:00466 | 0.00176 | 1.73 | 28.5562 | 44 | 180 | lymphocyte proliferation |
| GO:00068 | 0.00177 | 1.83 | 23.0036 | 37 | 145 | receptor-mediated endocytosis |
| GO:00148 | 0.00177 | 6.21 | 2.06239 | 7 | 13 | vascular smooth muscle contraction |
| GO:00197 | 0.00177 | 6.21 | 2.06239 | 7 | 13 | antibacterial humoral response |
| GO:00450 | 0.00177 | 6.21 | 2.06239 | 7 | 13 | T-helper 1 cell differentiation |
| GO:00420 | 0.0018 | 3.36 | 4.91802 | 12 | 31 | T-helper 1 type immune response |
| GO:00326 | 0.00181 | 2.98 | 6.18718 | 14 | 39 | interleukin-12 production |
| GO:00455 | 0.00183 | 3.15 | 5.5526 | 13 | 35 | mast cell activation |
| GO:19017 | 0.00185 | 1.38 | 94.5529 | 121 | 596 | cellular response to oxygen-containing compound |
| GO:00329 | 0.00186 | 7.98 | 1.58646 | 6 | 10 | inositol phosphate biosynthetic process |
| GO:00719 | 0.00195 | 1.50 | 55.3674 | 76 | 349 | regulation of protein serine/threonine kinase activity |
| GO:00023 | 0.00198 | 4.36 | 3.17291 | 9 | 20 | mucosal immune response |
| GO:00024 | 0.00198 | 4.36 | 3.17291 | 9 | 20 | neutrophil mediated immunity |

| | | | | | | |
|---|---|---|---|---|---|---|
| GO:00026 | 0.00198 | 1.30 | 147.065 | 179 | 927 | regulation of immune system process |
| GO:00512 | 0.002 | 1.58 | 41.8825 | 60 | 264 | negative regulation of multicellular organismal process |
| GO:00025 | 0.002 | 1.83 | 22.369 | 36 | 141 | myeloid leukocyte differentiation |
| GO:00093 | 0.00202 | 1.77 | 25.542 | 40 | 161 | protein secretion |
| GO:00350 | 0.00202 | 1.77 | 25.542 | 40 | 161 | regulation of Rho protein signal transduction |
| GO:00713 | 0.00213 | 1.62 | 36.1712 | 53 | 228 | cellular response to lipid |
| GO:00030 | 0.00213 | 1.30 | 138.18 | 169 | 871 | system process |
| GO:00226 | 0.00217 | 1.39 | 83.4477 | 108 | 526 | regulation of anatomical structure morphogenesis |
| GO:00461 | 0.00227 | 3.80 | 3.8075 | 10 | 24 | polyol biosynthetic process |
| GO:00019 | 0.00227 | 2.76 | 6.98041 | 15 | 44 | positive regulation of endothelial cell proliferation |
| GO:00420 | 0.00233 | 2.22 | 11.8984 | 22 | 75 | regulation of cytokine biosynthetic process |
| GO:00454 | 0.00248 | 3.20 | 5.07666 | 12 | 32 | regulation of nitric oxide biosynthetic process |
| GO:00301 | 0.0026 | 1.65 | 32.3637 | 48 | 204 | platelet activation |
| GO:00420 | 0.00266 | 2.10 | 13.4849 | 24 | 85 | cytokine biosynthetic process |
| GO:00026 | 0.00266 | 1.61 | 35.6953 | 52 | 225 | regulation of immune effector process |
| GO:00023 | 0.00272 | 2.58 | 7.77364 | 16 | 49 | cytokine production involved in immune response |
| GO:00022 | 0.00276 | 21.26 | 0.79323 | 4 | 5 | T cell activation via T cell receptor contact with antigen bound to MHC molecule on antigen presenting cell |
| GO:00023 | 0.00276 | 21.26 | 0.79323 | 4 | 5 | MHC protein complex assembly |
| GO:00025 | 0.00276 | 21.26 | 0.79323 | 4 | 5 | peptide antigen assembly with MHC protein complex |
| GO:00510 | 0.00284 | 1.70 | 27.6044 | 42 | 174 | positive regulation of secretion |
| GO:00071 | 0.00284 | 1.40 | 74.4048 | 97 | 469 | G-protein coupled receptor signaling pathway |
| GO:19013 | 0.00291 | 1.84 | 20.4653 | 33 | 129 | regulation of vasculature development |
| GO:00439 | 0.00295 | 1.24 | 213.22 | 249 | 1344 | macromolecular complex subunit organization |
| GO:00512 | 0.00295 | 1.17 | 504.652 | 554 | 3181 | establishment of localization |
| GO:00022 | 0.00298 | 3.99 | 3.33156 | 9 | 21 | organ or tissue specific immune response |
| GO:00069 | 0.00298 | 3.99 | 3.33156 | 9 | 21 | complement activation, classical pathway |
| GO:00095 | 0.00298 | 3.99 | 3.33156 | 9 | 21 | detection of biotic stimulus |
| GO:00457 | 0.00298 | 3.99 | 3.33156 | 9 | 21 | respiratory burst |
| GO:00343 | 0.00301 | 1.98 | 15.8646 | 27 | 100 | response to interferon-gamma |
| GO:00026 | 0.00307 | 1.61 | 34.2675 | 50 | 216 | positive regulation of leukocyte activation |
| GO:00316 | 0.0031 | 2.76 | 6.50447 | 14 | 41 | lipopolysaccharide-mediated signaling pathway |
| GO:00198 | 0.00314 | 2.07 | 13.6435 | 24 | 86 | antigen processing and presentation of exogenous peptide antigen via MHC class II |
| GO:00440 | 0.00316 | 1.59 | 36.8058 | 53 | 232 | regulation of system process |
| GO:00000 | 0.00325 | 3.55 | 3.96614 | 10 | 25 | regulation of transcription involved in G1/S transition of mitotic cell cycle |
| GO:00507 | 0.00327 | 1.86 | 19.0375 | 31 | 120 | regulation of protein secretion |
| GO:00326 | 0.00334 | 3.04 | 5.23531 | 12 | 33 | interleukin-10 production |
| GO:00156 | 0.00335 | 3.25 | 4.60073 | 11 | 29 | ferric iron transport |
| GO:00725 | 0.00335 | 3.25 | 4.60073 | 11 | 29 | trivalent inorganic cation transport |
| GO:00346 | 0.00337 | 1.24 | 200.687 | 235 | 1265 | nucleobase-containing compound catabolic process |
| GO:00507 | 0.0034 | 1.98 | 15.23 | 26 | 96 | positive regulation of peptidyl-tyrosine phosphorylation |
| GO:00723 | 0.00348 | 1.35 | 95.3461 | 120 | 601 | cardiovascular system development |
| GO:00723 | 0.00348 | 1.35 | 95.3461 | 120 | 601 | circulatory system development |
| GO:00603 | 0.00348 | 2.22 | 10.7879 | 20 | 68 | interferon-gamma-mediated signaling pathway |
| GO:00713 | 0.00353 | 2.08 | 13.0089 | 23 | 82 | cellular response to interferon-gamma |
| GO:00071 | 0.00355 | 1.19 | 326.493 | 368 | 2058 | cell surface receptor signaling pathway |
| GO:00158 | 0.00355 | 6.38 | 1.7451 | 6 | 11 | norepinephrine transport |
| GO:00026 | 0.00367 | 8.86 | 1.26917 | 5 | 8 | regulation of germinal center formation |
| GO:00026 | 0.00367 | 8.86 | 1.26917 | 5 | 8 | respiratory burst involved in defense response |
| GO:00069 | 0.00367 | 8.86 | 1.26917 | 5 | 8 | phagocytosis, recognition |
| GO:00199 | 0.00367 | 8.86 | 1.26917 | 5 | 8 | cGMP-mediated signaling |
| GO:00517 | 0.00367 | 8.86 | 1.26917 | 5 | 8 | positive regulation of nitric-oxide synthase biosynthetic process |
| GO:00068 | 0.00376 | 1.52 | 42.993 | 60 | 271 | cellular metal ion homeostasis |
| GO:00329 | 0.00378 | 1.77 | 22.369 | 35 | 141 | regulation of mononuclear cell proliferation |
| GO:00506 | 0.00378 | 1.77 | 22.369 | 35 | 141 | regulation of lymphocyte proliferation |
| GO:00068 | 0.00386 | 1.16 | 492.595 | 540 | 3105 | transport |
| GO:00985 | 0.00387 | 4.26 | 2.85562 | 8 | 18 | detection of external biotic stimulus |
| GO:00517 | 0.00396 | 1.96 | 15.3886 | 26 | 97 | interaction with host |
| GO:00027 | 0.00398 | 2.66 | 6.66312 | 14 | 42 | regulation of cytokine production involved in immune response |
| GO:00017 | 0.00399 | Inf | 0.47594 | 3 | 3 | type IIa hypersensitivity |
| GO:00017 | 0.00399 | Inf | 0.47594 | 3 | 3 | regulation of type IIa hypersensitivity |
| GO:00017 | 0.00399 | Inf | 0.47594 | 3 | 3 | positive regulation of type IIa hypersensitivity |
| GO:00022 | 0.00399 | Inf | 0.47594 | 3 | 3 | innate immune response activating cell surface receptor signaling pathway |
| GO:00022 | 0.00399 | Inf | 0.47594 | 3 | 3 | response to molecule of fungal origin |
| GO:00023 | 0.00399 | Inf | 0.47594 | 3 | 3 | serotonin production involved in inflammatory response |
| GO:00024 | 0.00399 | Inf | 0.47594 | 3 | 3 | serotonin secretion involved in inflammatory response |
| GO:00024 | 0.00399 | Inf | 0.47594 | 3 | 3 | type II hypersensitivity |
| GO:00025 | 0.00399 | Inf | 0.47594 | 3 | 3 | serotonin secretion by platelet |
| GO:00027 | 0.00399 | Inf | 0.47594 | 3 | 3 | cell surface pattern recognition receptor signaling pathway |
| GO:00027 | 0.00399 | Inf | 0.47594 | 3 | 3 | Fc receptor mediated inhibitory signaling pathway |
| GO:00028 | 0.00399 | Inf | 0.47594 | 3 | 3 | regulation of type II hypersensitivity |
| GO:00028 | 0.00399 | Inf | 0.47594 | 3 | 3 | positive regulation of type II hypersensitivity |
| GO:00109 | 0.00399 | Inf | 0.47594 | 3 | 3 | regulation of isomerase activity |
| GO:00109 | 0.00399 | Inf | 0.47594 | 3 | 3 | positive regulation of isomerase activity |
| GO:00341 | 0.00399 | Inf | 0.47594 | 3 | 3 | toll-like receptor 7 signaling pathway |
| GO:00355 | 0.00399 | Inf | 0.47594 | 3 | 3 | sequestering of TGFbeta in extracellular matrix |
| GO:00604 | 0.00399 | Inf | 0.47594 | 3 | 3 | positive regulation of penile erection |
| GO:20011 | 0.00399 | Inf | 0.47594 | 3 | 3 | regulation of T cell activation via T cell receptor contact with antigen bound to MHC molecule on antigen presenting cell |
| GO:00550 | 0.00407 | 1.48 | 49.9734 | 68 | 315 | metal ion homeostasis |
| GO:00507 | 0.00408 | 1.41 | 66.4726 | 87 | 419 | positive regulation of immune response |
| GO:00517 | 0.00423 | 1.15 | 701.69 | 753 | 4423 | cellular response to stimulus |
| GO:00326 | 0.00423 | 2.77 | 6.02854 | 13 | 38 | regulation of interleukin-12 production |
| GO:00068 | 0.00425 | 1.27 | 145.954 | 175 | 920 | ion transport |
| GO:00027 | 0.00434 | 3.68 | 3.49021 | 9 | 22 | positive regulation of cytokine production involved in immune response |
| GO:20001 | 0.00439 | 1.42 | 63.141 | 83 | 398 | regulation of cell motility |
| GO:00507 | 0.00445 | 1.33 | 99.6295 | 124 | 628 | regulation of immune response |
| GO:00019 | 0.00454 | 3.33 | 4.12479 | 10 | 26 | blood vessel remodeling |
| GO:00339 | 0.00465 | 1.38 | 74.7221 | 96 | 471 | response to lipid |
| GO:00022 | 0.0047 | 1.44 | 56.3192 | 75 | 355 | activation of immune response |
| GO:00510 | 0.00488 | 1.43 | 57.2711 | 76 | 361 | regulation of secretion |
| GO:00197 | 0.00496 | 4.65 | 2.37969 | 7 | 15 | antimicrobial humoral response |
| GO:00985 | 0.00496 | 4.65 | 2.37969 | 7 | 15 | detection of other organism |
| GO:00313 | 0.00498 | 1.40 | 66.9485 | 87 | 422 | regulation of defense response |
| GO:19017 | 0.00524 | 1.27 | 140.243 | 168 | 884 | response to oxygen-containing compound |
| GO:00303 | 0.00524 | 1.42 | 59.1749 | 78 | 373 | regulation of cell migration |
| GO:00068 | 0.00526 | 1.47 | 47.911 | 65 | 302 | cellular ion homeostasis |
| GO:00421 | 0.00527 | 2.37 | 8.24958 | 16 | 52 | B cell proliferation |
| GO:00512 | 0.00558 | 1.38 | 70.756 | 91 | 446 | regulation of cellular component movement |
| GO:00485 | 0.00559 | 1.44 | 52.3531 | 70 | 330 | blood vessel morphogenesis |
| GO:00102 | 0.00571 | 1.35 | 80.592 | 102 | 508 | response to organonitrogen compound |
| GO:00301 | 0.00577 | 3.87 | 3.01427 | 8 | 19 | sphingolipid catabolic process |
| GO:00326 | 0.00577 | 3.87 | 3.01427 | 8 | 19 | negative regulation of interferon-gamma production |
| GO:00068 | 0.00578 | 2.78 | 5.5526 | 12 | 35 | superoxide metabolic process |
| GO:00072 | 0.00578 | 2.78 | 5.5526 | 12 | 35 | glutamate receptor signaling pathway |
| GO:00484 | 0.0058 | 2.42 | 7.61499 | 15 | 48 | oogenesis |
| GO:00516 | 0.00582 | 1.67 | 25.3833 | 38 | 160 | detection of stimulus |
| GO:00400 | 0.00601 | 1.38 | 69.1695 | 89 | 436 | regulation of locomotion |
| GO:00326 | 0.00605 | 2.93 | 4.91802 | 11 | 31 | regulation of interleukin-10 production |
| GO:00706 | 0.00607 | 1.70 | 23.0036 | 35 | 145 | regulation of leukocyte proliferation |
| GO:00433 | 0.00614 | 3.42 | 3.64885 | 9 | 23 | mast cell degranulation |
| GO:00022 | 0.00615 | 5.32 | 1.90375 | 6 | 12 | innate immune response in mucosa |
| GO:00022 | 0.00615 | 5.32 | 1.90375 | 6 | 12 | macrophage activation involved in immune response |
| GO:00362 | 0.00615 | 5.32 | 1.90375 | 6 | 12 | response to increased oxygen levels |
| GO:00550 | 0.00615 | 5.32 | 1.90375 | 6 | 12 | response to hyperoxia |
| GO:00027 | 0.0062 | 3.13 | 4.28343 | 10 | 27 | positive regulation of production of molecular mediator of immune response |
| GO:00107 | 0.0062 | 3.13 | 4.28343 | 10 | 27 | positive regulation of epithelial to mesenchymal transition |
| GO:00326 | 0.0063 | 2.18 | 9.83604 | 18 | 62 | regulation of interleukin-6 production |
| GO:00105 | 0.00633 | 1.31 | 102.485 | 126 | 646 | positive regulation of phosphorus metabolic process |
| GO:00459 | 0.00633 | 1.31 | 102.485 | 126 | 646 | positive regulation of phosphate metabolic process |
| GO:00450 | 0.00634 | 1.59 | 30.46 | 44 | 192 | positive regulation of innate immune response |
| GO:00308 | 0.00635 | 2.49 | 6.98041 | 14 | 44 | regulation of B cell proliferation |
| GO:00723 | 0.00635 | 2.49 | 6.98041 | 14 | 44 | protein activation cascade |
| GO:00435 | 0.00639 | 1.39 | 63.141 | 82 | 398 | positive regulation of GTPase activity |
| GO:00026 | 0.00647 | 2.30 | 8.40822 | 16 | 53 | positive regulation of leukocyte migration |
| GO:00462 | 0.00647 | 2.30 | 8.40822 | 16 | 53 | nitric oxide metabolic process |
| GO:00300 | 0.00657 | 1.46 | 46.6418 | 63 | 294 | cellular cation homeostasis |
| GO:00516 | 0.00681 | 1.19 | 283.976 | 320 | 1790 | establishment of localization in cell |
| GO:00068 | 0.00691 | 2.56 | 6.34583 | 13 | 40 | nitric oxide biosynthetic process |
| GO:00323 | 0.00703 | 1.47 | 45.0554 | 61 | 284 | regulation of Ras GTPase activity |
| GO:00454 | 0.0071 | 1.76 | 19.1961 | 30 | 121 | fat cell differentiation |
| GO:00455 | 0.00715 | 1.37 | 66.9485 | 86 | 422 | positive regulation of cell differentiation |
| GO:00508 | 0.00716 | 2.35 | 7.77364 | 15 | 49 | positive regulation of B cell activation |
| GO:00510 | 0.00717 | 1.36 | 71.3906 | 91 | 450 | positive regulation of transport |
| GO:00360 | 0.00719 | 6.64 | 1.42781 | 5 | 9 | osteoclast development |
| GO:00433 | 0.00719 | 6.64 | 1.42781 | 5 | 9 | neutrophil degranulation |
| GO:00610 | 0.00719 | 6.64 | 1.42781 | 5 | 9 | positive regulation of myeloid leukocyte cytokine production involved in immune response |

| | | | | | | |
|---|---|---|---|---|---|---|
| GO:00712 | 0.00719 | 6.64 | 1.42781 | 5 | 9 | cellular response to antibiotic |
| GO:00903 | 0.00719 | 6.64 | 1.42781 | 5 | 9 | regulation of platelet aggregation |
| GO:00066 | 0.00723 | 10.63 | 0.95187 | 4 | 6 | sphingomyelin catabolic process |
| GO:00109 | 0.00723 | 10.63 | 0.95187 | 4 | 6 | regulation of inositol phosphate biosynthetic process |
| GO:00316 | 0.00723 | 10.63 | 0.95187 | 4 | 6 | positive regulation of lipopolysaccharide-mediated signaling pathway |
| GO:00326 | 0.00723 | 10.63 | 0.95187 | 4 | 6 | interleukin-18 production |
| GO:00443 | 0.00723 | 10.63 | 0.95187 | 4 | 6 | cellular response to leptin stimulus |
| GO:00520 | 0.00723 | 10.63 | 0.95187 | 4 | 6 | pathogen-associated molecular pattern dependent induction by symbiont of host innate immune response |
| GO:00521 | 0.00723 | 10.63 | 0.95187 | 4 | 6 | positive regulation by symbiont of host innate immune response |
| GO:00521 | 0.00723 | 10.63 | 0.95187 | 4 | 6 | modulation by symbiont of host innate immune response |
| GO:00521 | 0.00723 | 10.63 | 0.95187 | 4 | 6 | pathogen-associated molecular pattern dependent modulation by symbiont of host innate immune response |
| GO:00522 | 0.00723 | 10.63 | 0.95187 | 4 | 6 | pathogen-associated molecular pattern dependent induction by organism of innate immune response of other organism involved in symbiotic interaction |
| GO:00523 | 0.00723 | 10.63 | 0.95187 | 4 | 6 | positive regulation by organism of innate immune response in other organism involved in symbiotic interaction |
| GO:00523 | 0.00723 | 10.63 | 0.95187 | 4 | 6 | modulation by organism of innate immune response in other organism involved in symbiotic interaction |
| GO:00523 | 0.00723 | 10.63 | 0.95187 | 4 | 6 | pathogen-associated molecular pattern dependent modulation by organism of innate immune response in other organism involved in symbiotic interaction |
| GO:00525 | 0.00723 | 10.63 | 0.95187 | 4 | 6 | positive regulation by organism of immune response of other organism involved in symbiotic interaction |
| GO:00525 | 0.00723 | 10.63 | 0.95187 | 4 | 6 | positive regulation by symbiont of host immune response |
| GO:00609 | 0.00723 | 10.63 | 0.95187 | 4 | 6 | positive regulation of macrophage cytokine production |
| GO:00061 | 0.00736 | 1.33 | 84.8755 | 106 | 535 | regulation of nucleotide metabolic process |
| GO:00027 | 0.00744 | 2.66 | 5.71125 | 12 | 36 | regulation of B cell mediated immunity |
| GO:00510 | 0.00755 | 1.38 | 62.6651 | 81 | 395 | regulation of small GTPase mediated signal transduction |
| GO:00330 | 0.00756 | 1.96 | 13.0089 | 22 | 82 | muscle cell proliferation |
| GO:20000 | 0.00759 | 1.25 | 148.968 | 176 | 939 | regulation of multicellular organismal development |
| GO:00513 | 0.0076 | 1.29 | 108.514 | 132 | 684 | positive regulation of hydrolase activity |
| GO:00425 | 0.00762 | 4.14 | 2.53833 | 7 | 16 | superoxide anion generation |
| GO:00140 | 0.00769 | 1.35 | 74.2462 | 94 | 468 | response to organic cyclic compound |
| GO:00028 | 0.00798 | 1.85 | 15.3886 | 25 | 97 | regulation of adaptive immune response |
| GO:00022 | 0.008 | 1.62 | 25.8593 | 38 | 163 | activation of innate immune response |
| GO:00442 | 0.00801 | 1.41 | 54.8914 | 72 | 346 | small molecule biosynthetic process |
| GO:00024 | 0.00805 | 1.83 | 16.1819 | 26 | 102 | production of molecular mediator of immune response |
| GO:00308 | 0.00829 | 2.96 | 4.44208 | 10 | 28 | positive regulation of B cell proliferation |
| GO:00335 | 0.00829 | 2.96 | 4.44208 | 10 | 28 | transferrin transport |
| GO:00459 | 0.00829 | 2.96 | 4.44208 | 10 | 28 | positive regulation of exocytosis |
| GO:00226 | 0.00829 | 2.01 | 11.5811 | 20 | 73 | extracellular matrix disassembly |
| GO:00105 | 0.0083 | 3.55 | 3.17291 | 8 | 20 | regulation of platelet activation |
| GO:00421 | 0.0083 | 3.55 | 3.17291 | 8 | 20 | neutrophil activation |
| GO:00464 | 0.0083 | 3.55 | 3.17291 | 8 | 20 | membrane lipid catabolic process |
| GO:00072 | 0.00836 | 1.28 | 112.48 | 136 | 709 | cell-cell signaling |
| GO:00423 | 0.00844 | 1.31 | 92.4905 | 114 | 583 | positive regulation of phosphorylation |
| GO:00022 | 0.00846 | 3.19 | 3.8075 | 9 | 24 | mast cell activation involved in immune response |
| GO:00024 | 0.00846 | 3.19 | 3.8075 | 9 | 24 | mast cell mediated immunity |
| GO:19005 | 0.00854 | 1.32 | 84.3995 | 105 | 532 | regulation of purine nucleotide metabolic process |
| GO:00015 | 0.00874 | 1.39 | 59.4921 | 77 | 375 | blood vessel development |
| GO:00327 | 0.00894 | 1.15 | 443.256 | 484 | 2794 | RNA biosynthetic process |
| GO:00163 | 0.00924 | 1.50 | 36.1712 | 50 | 228 | single organismal cell-cell adhesion |
| GO:00446 | 0.00944 | 1.16 | 1478.58 | 1517 | 9320 | single-organism process |
| GO:00326 | 0.00946 | 2.56 | 5.86989 | 12 | 37 | interleukin-1 production |
| GO:00508 | 0.0095 | 1.37 | 62.3478 | 80 | 393 | ion homeostasis |
| GO:00069 | 0.00974 | 2.33 | 7.2977 | 14 | 46 | cellular defense response |
| GO:19016 | 0.00985 | 1.31 | 86.6206 | 107 | 546 | response to nitrogen compound |
| GO:00021 | 0.00991 | 4.56 | 2.06239 | 6 | 13 | cytoplasmic translation |
| GO:00024 | 0.00991 | 4.56 | 2.06239 | 6 | 13 | germinal center formation |
| GO:00066 | 0.00991 | 4.56 | 2.06239 | 6 | 13 | sphingomyelin metabolic process |
| GO:00156 | 0.00991 | 4.56 | 2.06239 | 6 | 13 | organic cation transport |
| GO:00425 | 0.00991 | 4.56 | 2.06239 | 6 | 13 | tumor necrosis factor biosynthetic process |
| GO:00425 | 0.00991 | 4.56 | 2.06239 | 6 | 13 | regulation of tumor necrosis factor biosynthetic process |
| GO:00440 | 0.00991 | 4.56 | 2.06239 | 6 | 13 | positive regulation of cellular component biogenesis |
