## Supplemental Table 2 for "RNA-seq of Human T-Cells After Hematopoietic Stem Cell Transplantation Identifies Linc00402 as a Novel Regulator of T-Cell Alloimmunity"

| GOMFID | Pvalue | OddsRatio | ExpCount | Count | Size | Term |
|---|---|---|---|---|---|---|
| GO:00048 | 1.70E-17 | 2.039487 | 158.645 | 255 | 728 | receptor activity |
| GO:00380 | 1.14E-14 | 2.067085 | 122.4704 | 200 | 562 | signaling receptor activity |
| GO:00037 | 8.13E-13 | 3.462961 | 31.38033 | 70 | 144 | structural constituent of ribosome |
| GO:00048 | 9.44E-13 | 2.048975 | 106.1265 | 173 | 487 | transmembrane signaling receptor activity |
| GO:00048 | 2.74E-09 | 1.621941 | 176.0785 | 245 | 808 | signal transducer activity |
| GO:00600 | 2.74E-09 | 1.621941 | 176.0785 | 245 | 808 | molecular transducer activity |
| GO:00055 | 3.45E-07 | 1.73011 | 93.92306 | 138 | 431 | calcium ion binding |
| GO:00302 | 2.10E-06 | 36.02696 | 2.397108 | 10 | 11 | lipoprotein particle receptor activity |
| GO:00611 | 1.24E-05 | 2.349699 | 25.49651 | 46 | 117 | peptidase regulator activity |
| GO:00304 | 1.60E-05 | 2.660322 | 18.52311 | 36 | 85 | peptidase inhibitor activity |
| GO:00051 | 1.89E-05 | 1.596176 | 91.09011 | 127 | 418 | structural molecule activity |
| GO:00050 | 2.32E-05 | Inf | 1.525432 | 7 | 7 | low-density lipoprotein receptor activity |
| GO:00048 | 5.11E-05 | 3.610514 | 9.152595 | 21 | 42 | serine-type endopeptidase inhibitor activity |
| GO:00049 | 5.65E-05 | 1.772495 | 51.21095 | 77 | 235 | G-protein coupled receptor activity |
| GO:00152 | 6.20E-05 | 1.804945 | 47.28841 | 72 | 217 | channel activity |
| GO:00228 | 6.20E-05 | 1.804945 | 47.28841 | 72 | 217 | passive transmembrane transporter activity |
| GO:00048 | 6.21E-05 | 2.510515 | 18.08727 | 34 | 83 | endopeptidase inhibitor activity |
| GO:00083 | 9.92E-05 | 9.003767 | 3.050865 | 10 | 14 | signaling pattern recognition receptor activity |
| GO:00381 | 9.92E-05 | 9.003767 | 3.050865 | 10 | 14 | pattern recognition receptor activity |
| GO:00051 | 0.00011 | 3.82002 | 7.627162 | 18 | 35 | Rho GTPase activator activity |
| GO:00228 | 0.000117 | 1.789632 | 44.8913 | 68 | 206 | substrate-specific channel activity |
| GO:00508 | 0.000185 | 2.037168 | 27.23987 | 45 | 125 | cell adhesion molecule binding |
| GO:00611 | 0.000187 | 2.320015 | 18.95895 | 34 | 87 | endopeptidase regulator activity |
| GO:00082 | 0.000208 | 1.50099 | 90.43635 | 121 | 415 | lipid binding |
| GO:00380 | 0.000271 | 3.137635 | 9.370514 | 20 | 43 | cargo receptor activity |
| GO:00052 | 0.000366 | 1.721261 | 44.01962 | 65 | 202 | ion channel activity |
| GO:00055 | 0.000407 | 3.115346 | 8.934676 | 19 | 41 | collagen binding |
| GO:00038 | 0.000478 | 2.677946 | 11.76762 | 23 | 54 | antigen binding |
| GO:00018 | 0.00049 | Inf | 1.089595 | 5 | 5 | lipopolysaccharide receptor activity |
| GO:00433 | 0.000513 | 6.00119 | 3.486703 | 10 | 16 | proteoglycan binding |
| GO:00228 | 0.000547 | 1.380552 | 127.9184 | 161 | 587 | transmembrane transporter activity |
| GO:00228 | 0.000573 | 1.794287 | 34.86703 | 53 | 160 | gated channel activity |
| GO:00038 | 0.000607 | 21.58195 | 1.525432 | 6 | 7 | gamma-glutamyltransferase activity |
| GO:00052 | 0.000688 | 1.810435 | 32.68784 | 50 | 150 | cation channel activity |
| GO:00468 | 0.000872 | 1.609653 | 50.33927 | 71 | 231 | metal ion transmembrane transporter activity |
| GO:00508 | 0.000936 | 3.204028 | 7.409243 | 16 | 34 | extracellular matrix binding |
| GO:00055 | 0.001004 | 5.14331 | 3.704622 | 10 | 17 | 1-phosphatidylinositol binding |
| GO:00167 | 0.001004 | 5.14331 | 3.704622 | 10 | 17 | transferase activity, transferring amino-acyl groups |
| GO:00055 | 0.001265 | 1.586337 | 50.12135 | 70 | 230 | phospholipid binding |
| GO:00228 | 0.001339 | 1.365601 | 115.9329 | 145 | 532 | substrate-specific transmembrane transporter activity |
| GO:00341 | 0.001481 | 8.394455 | 2.179189 | 7 | 10 | apolipoprotein binding |
| GO:00302 | 0.001558 | 1.634061 | 42.05835 | 60 | 193 | carbohydrate binding |
| GO:00150 | 0.00175 | 1.37047 | 106.7803 | 134 | 490 | ion transmembrane transporter activity |
| GO:00047 | 0.001764 | 2.703759 | 9.152595 | 18 | 42 | transmembrane receptor protein tyrosine kinase activity |
| GO:00151 | 0.001764 | 2.703759 | 9.152595 | 18 | 42 | amino acid transmembrane transporter activity |
| GO:00355 | 0.001819 | 4.499901 | 3.922541 | 10 | 18 | purinergic receptor activity |
| GO:00051 | 0.001931 | 2.105731 | 16.56184 | 28 | 76 | integrin binding |
| GO:00198 | 0.002234 | 5.757211 | 2.832946 | 8 | 13 | immunoglobulin binding |
| GO:00169 | 0.002251 | Inf | 0.871676 | 4 | 4 | galactoside binding |
| GO:00422 | 0.002367 | 1.785532 | 27.02195 | 41 | 124 | peptide binding |
| GO:00332 | 0.002919 | 1.744911 | 28.11154 | 42 | 129 | amide binding |
| GO:00016 | 0.00292 | 2.266864 | 12.42138 | 22 | 57 | peptide receptor activity |
| GO:00171 | 0.003096 | 3.999471 | 4.14046 | 10 | 19 | Wnt-protein binding |
| GO:00037 | 0.00318 | 1.272149 | 167.7976 | 199 | 770 | sequence-specific DNA binding transcription factor activity |
| GO:00050 | 0.003232 | 1.860361 | 21.79189 | 34 | 100 | Ras GTPase activator activity |
| GO:00151 | 0.00324 | 3.121239 | 6.10173 | 13 | 28 | L-amino acid transmembrane transporter activity |
| GO:00010 | 0.003384 | 1.269776 | 168.0155 | 199 | 771 | nucleic acid binding transcription factor activity |
| GO:00052 | 0.003643 | 1.861738 | 21.13814 | 33 | 97 | voltage-gated ion channel activity |
| GO:00228 | 0.003643 | 1.861738 | 21.13814 | 33 | 97 | voltage-gated channel activity |
| GO:00048 | 0.003752 | 2.203652 | 12.6393 | 22 | 58 | cytokine receptor activity |

| | | | | | | |
|---|---|---|---|---|---|---|
| GO:00016 | 0.004221 | 4.797147 | 3.050865 | 8 | 14 | purinergic nucleotide receptor activity |
| GO:00165 | 0.004221 | 4.797147 | 3.050865 | 8 | 14 | nucleotide receptor activity |
| GO:00228 | 0.004496 | 1.286328 | 138.5964 | 166 | 636 | substrate-specific transporter activity |
| GO:00019 | 0.004654 | 2.252936 | 11.33178 | 20 | 52 | glycoprotein binding |
| GO:00151 | 0.004754 | 4.048306 | 3.704622 | 9 | 17 | neutral amino acid transmembrane transporter activity |
| GO:00053 | 0.004838 | 7.192399 | 1.96127 | 6 | 9 | glucose transmembrane transporter activity |
| GO:00151 | 0.004838 | 7.192399 | 1.96127 | 6 | 9 | hexose transmembrane transporter activity |
| GO:00323 | 0.004838 | 7.192399 | 1.96127 | 6 | 9 | MHC class II receptor activity |
| GO:00426 | 0.005025 | 3.299782 | 5.012135 | 11 | 23 | peptide antigen binding |
| GO:00053 | 0.005267 | 2.003723 | 15.25432 | 25 | 70 | organic acid transmembrane transporter activity |
| GO:00085 | 0.005341 | 2.162963 | 12.20346 | 21 | 56 | G-protein coupled peptide receptor activity |
| GO:00191 | 0.005341 | 2.162963 | 12.20346 | 21 | 56 | transmembrane receptor protein kinase activity |
| GO:00052 | 0.005391 | 1.250997 | 170.6305 | 200 | 783 | transporter activity |
| GO:00048 | 0.005653 | 1.494348 | 47.72424 | 64 | 219 | enzyme inhibitor activity |
| GO:00167 | 0.005735 | 2.572791 | 7.845081 | 15 | 36 | transferase activity, transferring sulfur-containing groups |
| GO:00015 | 0.00645 | 5.035564 | 2.615027 | 7 | 12 | lipopolysaccharide binding |
| GO:00152 | 0.006639 | 1.67668 | 26.80403 | 39 | 123 | secondary active transmembrane transporter activity |
| GO:00452 | 0.006905 | 8.98793 | 1.525432 | 5 | 7 | alpha-catenin binding |
| GO:00047 | 0.007003 | 1.710084 | 24.40692 | 36 | 112 | protein tyrosine kinase activity |
| GO:00048 | 0.00712 | 2.88 | 5.883811 | 12 | 27 | cysteine-type endopeptidase inhibitor activity |
| GO:00469 | 0.007307 | 1.966725 | 14.81849 | 24 | 68 | carboxylic acid transmembrane transporter activity |
| GO:00150 | 0.00752 | 1.872754 | 17.2156 | 27 | 79 | calcium ion transmembrane transporter activity |
| GO:00432 | 0.007694 | 3.271574 | 4.576297 | 10 | 21 | laminin binding |
| GO:00050 | 0.008242 | 1.974179 | 14.16473 | 23 | 65 | Rho guanyl-nucleotide exchange factor activity |
| GO:00162 | 0.008938 | 1.922807 | 15.03641 | 24 | 69 | lipase activity |
| GO:00301 | 0.008938 | 1.922807 | 15.03641 | 24 | 69 | PDZ domain binding |
| GO:00049 | 0.009295 | 14.37658 | 1.089595 | 4 | 5 | extracellular ATP-gated cation channel activity |
| GO:00353 | 0.009295 | 14.37658 | 1.089595 | 4 | 5 | ATP-gated ion channel activity |
| GO:00198 | 0.009565 | 1.803315 | 18.30519 | 28 | 84 | growth factor binding |

| GOCCID | Pvalue | OddsRatio | ExpCount | Count | Size | Term |
|---|---|---|---|---|---|---|
| GO:00226 | 5.12E-27 | 10.84058 | 19.85283 | 68 | 91 | cytosolic ribosome |
| GO:00719 | 8.18E-25 | 1.648673 | 648.162 | 854 | 2971 | cell periphery |
| GO:00058 | 4.12E-24 | 1.642439 | 632.018 | 833 | 2897 | plasma membrane |
| GO:00443 | 9.63E-19 | 4.907014 | 28.57934 | 75 | 131 | ribosomal subunit |
| GO:00226 | 2.75E-18 | 14.53328 | 10.90815 | 40 | 50 | cytosolic large ribosomal subunit |
| GO:00444 | 1.20E-17 | 1.706691 | 320.9177 | 452 | 1471 | plasma membrane part |
| GO:00312 | 2.84E-14 | 1.82665 | 181.9479 | 273 | 834 | intrinsic component of plasma membrane |
| GO:00058 | 3.53E-13 | 1.801302 | 173.8758 | 259 | 797 | integral component of plasma membrane |
| GO:00159 | 9.62E-12 | 5.255727 | 15.48957 | 42 | 71 | large ribosomal subunit |
| GO:00226 | 1.25E-11 | 10.1281 | 8.290191 | 28 | 38 | cytosolic small ribosomal subunit |
| GO:00444 | 3.44E-11 | 2.848133 | 38.833 | 78 | 178 | cytosolic part |
| GO:00312 | 4.86E-11 | 1.359382 | 756.1527 | 892 | 3466 | intrinsic component of membrane |
| GO:00056 | 1.14E-10 | 1.760828 | 145.7328 | 215 | 668 | extracellular space |
| GO:00160 | 2.94E-10 | 1.344069 | 741.7539 | 871 | 3400 | integral component of membrane |
| GO:00055 | 5.27E-10 | 1.357916 | 630.0545 | 751 | 2888 | extracellular region |
| GO:00058 | 1.67E-09 | 2.494216 | 42.9781 | 80 | 197 | ribosome |
| GO:00003 | 5.77E-09 | 1.905376 | 86.82884 | 136 | 398 | lytic vacuole |
| GO:00057 | 5.77E-09 | 1.905376 | 86.82884 | 136 | 398 | lysosome |
| GO:00444 | 6.00E-09 | 1.295179 | 916.2843 | 1042 | 4200 | membrane part |
| GO:00057 | 1.27E-08 | 1.824152 | 96.64617 | 147 | 443 | vacuole |
| GO:00059 | 1.64E-08 | 1.953994 | 74.61172 | 119 | 342 | cell-substrate adherens junction |
| GO:00057 | 1.69E-08 | 4.629737 | 12.43529 | 32 | 57 | vacuolar lumen |
| GO:00059 | 1.84E-08 | 1.954768 | 73.95723 | 118 | 339 | focal adhesion |
| GO:00444 | 2.34E-08 | 1.330932 | 559.1516 | 663 | 2563 | extracellular region part |
| GO:00300 | 2.87E-08 | 1.927451 | 75.26621 | 119 | 345 | cell-substrate junction |
| GO:00159 | 3.48E-08 | 4.263141 | 13.30794 | 33 | 61 | small ribosomal subunit |
| GO:00432 | 3.67E-08 | 4.715273 | 11.56263 | 30 | 53 | lysosomal lumen |
| GO:00059 | 9.26E-08 | 1.817111 | 85.51987 | 130 | 392 | adherens junction |
| GO:00306 | 1.27E-07 | 2.977105 | 22.25262 | 46 | 102 | endocytic vesicle membrane |
| GO:00701 | 1.41E-07 | 1.790271 | 87.70149 | 132 | 402 | anchoring junction |
| GO:00444 | 2.45E-07 | 1.928704 | 64.35806 | 102 | 295 | vacuolar part |
| GO:00300 | 2.98E-07 | 1.516843 | 176.0575 | 235 | 807 | cell junction |
| GO:00099 | 4.60E-07 | 1.696077 | 99.26413 | 144 | 455 | cell surface |
| GO:00319 | 1.03E-06 | 1.278465 | 584.4585 | 676 | 2679 | vesicle |
| GO:00319 | 1.63E-06 | 1.275059 | 570.0597 | 659 | 2613 | membrane-bounded vesicle |
| GO:00160 | 2.30E-06 | 1.220725 | 1329.703 | 1436 | 6095 | membrane |
| GO:00429 | 9.01E-06 | 1.369576 | 241.9427 | 300 | 1109 | cell projection |
| GO:00306 | 9.61E-06 | 5.566942 | 6.108562 | 17 | 28 | clathrin-coated endocytic vesicle membrane |
| GO:00426 | 9.81E-06 | 13.18948 | 3.054281 | 11 | 14 | MHC class II protein complex |
| GO:00301 | 1.51E-05 | 2.000699 | 39.26933 | 64 | 180 | endocytic vesicle |
| GO:00432 | 3.51E-05 | 1.252747 | 459.8874 | 530 | 2108 | extracellular organelle |
| GO:00650 | 3.51E-05 | 1.252747 | 459.8874 | 530 | 2108 | extracellular membrane-bounded organelle |
| GO:00700 | 3.51E-05 | 1.252747 | 459.8874 | 530 | 2108 | extracellular vesicular exosome |
| GO:00325 | 4.53E-05 | 4.026416 | 7.853865 | 19 | 36 | trans-Golgi network membrane |
| GO:00098 | 4.54E-05 | 2.007697 | 34.25158 | 56 | 157 | external side of plasma membrane |
| GO:00453 | 6.80E-05 | 4.051826 | 7.417539 | 18 | 34 | clathrin-coated endocytic vesicle |
| GO:00055 | 0.000186 | 1.858219 | 36.65137 | 57 | 168 | proteinaceous extracellular matrix |
| GO:00310 | 0.000214 | 1.725138 | 47.34135 | 70 | 217 | extracellular matrix |
| GO:00985 | 0.000266 | 1.693815 | 49.30482 | 72 | 226 | side of membrane |
| GO:00312 | 0.000389 | 2.26343 | 18.10752 | 32 | 83 | anchored component of membrane |
| GO:00057 | 0.000448 | 1.667524 | 48.43217 | 70 | 222 | lysosomal membrane |
| GO:00057 | 0.000518 | 1.619388 | 53.66808 | 76 | 246 | vacuolar membrane |
| GO:00985 | 0.000661 | 1.566445 | 59.9948 | 83 | 275 | plasma membrane region |
| GO:00312 | 0.001048 | 1.543946 | 59.12215 | 81 | 271 | cell leading edge |
| GO:00306 | 0.001097 | 2.401479 | 13.08978 | 24 | 60 | clathrin-coated vesicle membrane |
| GO:00312 | 0.00136 | 1.972323 | 21.59813 | 35 | 99 | leading edge membrane |
| GO:00301 | 0.002017 | 1.912162 | 22.03446 | 35 | 101 | clathrin-coated vesicle |
| GO:00452 | 0.002302 | 1.424087 | 78.10233 | 101 | 358 | synapse |
| GO:00430 | 0.002368 | 1.343086 | 114.7537 | 142 | 526 | neuron projection |
| GO:00058 | 0.002459 | 2.590778 | 9.381006 | 18 | 43 | kinesin complex |

| | | | | | | |
|---|---|---|---|---|---|---|
| GO:00444 | 0.00309 | 2.080898 | 15.48957 | 26 | 71 | extracellular matrix part |
| GO:00466 | 0.003277 | 3.115979 | 6.108562 | 13 | 28 | anchored component of plasma membrane |
| GO:00163 | 0.003369 | 1.739737 | 27.48853 | 41 | 126 | basolateral plasma membrane |
| GO:00325 | 0.005074 | 3.294334 | 5.017747 | 11 | 23 | neuron projection membrane |
| GO:00426 | 0.005074 | 3.294334 | 5.017747 | 11 | 23 | MHC protein complex |
| GO:00974 | 0.005522 | 1.277584 | 138.3153 | 165 | 634 | neuron part |
| GO:00432 | 0.006144 | 1.538806 | 40.14198 | 55 | 184 | receptor complex |
| GO:00453 | 0.006588 | 1.956441 | 15.48957 | 25 | 71 | phagocytic vesicle |
| GO:00347 | 0.00718 | 1.642237 | 28.57934 | 41 | 131 | ion channel complex |
| GO:00312 | 0.007387 | 4.104793 | 3.272444 | 8 | 15 | intrinsic component of external side of plasma membrane |
| GO:00059 | 0.007686 | 2.116257 | 11.7808 | 20 | 54 | caveola |
| GO:00056 | 0.008579 | 2.042028 | 12.65345 | 21 | 58 | basement membrane |

| GOBPID | Pvalue | OddsRatio | ExpCount | Count | Size | Term |
|---|---|---|---|---|---|---|
| GO:00064 | 2.60E-26 | 9.99 | 20.7381 | 69 | 94 | translational termination |
| GO:00066 | 2.13E-19 | 6.11 | 23.16489 | 66 | 105 | SRP-dependent cotranslational protein targeting to membrane |
| GO:00066 | 9.14E-19 | 5.81 | 23.60613 | 66 | 107 | cotranslational protein targeting to membrane |
| GO:00450 | 1.85E-18 | 5.67 | 23.82675 | 66 | 108 | protein targeting to ER |
| GO:00725 | 3.69E-18 | 5.54 | 24.04737 | 66 | 109 | establishment of protein localization to endoplasmic reticulum |
| GO:00436 | 4.56E-18 | 4.04 | 36.40198 | 87 | 165 | cellular protein complex disassembly |
| GO:00064 | 9.08E-18 | 5.08 | 26.03293 | 69 | 118 | translational elongation |
| GO:00001 | 1.31E-17 | 5.11 | 25.59169 | 68 | 116 | nuclear-transcribed mRNA catabolic process, nonsense-mediated decay |
| GO:00432 | 4.04E-16 | 3.47 | 41.03495 | 91 | 186 | protein complex disassembly |
| GO:00329 | 7.27E-15 | 3.20 | 43.24113 | 92 | 196 | macromolecular complex disassembly |
| GO:00709 | 2.72E-14 | 4.10 | 27.35664 | 66 | 124 | protein localization to endoplasmic reticulum |
| GO:00096 | 6.66E-13 | 1.59 | 300.2611 | 406 | 1361 | response to external stimulus |
| GO:00066 | 3.61E-12 | 3.08 | 36.84321 | 77 | 167 | protein targeting to membrane |
| GO:00064 | 3.64E-12 | 3.11 | 36.18136 | 76 | 164 | translational initiation |
| GO:00096 | 1.24E-11 | 2.40 | 60.89058 | 110 | 276 | response to bacterium |
| GO:00447 | 1.39E-11 | 1.36 | 897.6948 | 1042 | 4069 | single-multicellular organism process |
| GO:00190 | 1.71E-11 | 3.09 | 34.41641 | 72 | 156 | viral transcription |
| GO:00069 | 2.01E-11 | 2.07 | 90.01216 | 148 | 408 | inflammatory response |
| GO:00440 | 2.15E-11 | 2.90 | 38.60816 | 78 | 175 | multi-organism metabolic process |
| GO:00190 | 2.34E-11 | 2.97 | 36.62259 | 75 | 166 | viral gene expression |
| GO:00224 | 6.40E-11 | 2.12 | 79.64311 | 133 | 361 | cellular component disassembly |
| GO:00325 | 2.04E-10 | 1.33 | 928.5813 | 1065 | 4209 | multicellular organismal process |
| GO:00508 | 2.38E-10 | 1.32 | 1191.337 | 1332 | 5400 | response to stimulus |
| GO:00324 | 3.56E-10 | 2.73 | 38.38754 | 75 | 174 | response to lipopolysaccharide |
| GO:00022 | 3.90E-10 | 2.63 | 41.25557 | 79 | 187 | response to molecule of bacterial origin |
| GO:00096 | 4.66E-10 | 1.70 | 158.845 | 228 | 720 | response to wounding |
| GO:00069 | 5.32E-10 | 1.56 | 237.8262 | 320 | 1078 | defense response |
| GO:00064 | 1.29E-09 | 2.54 | 42.13804 | 79 | 191 | mRNA catabolic process |
| GO:00650 | 1.30E-09 | 1.32 | 1666.328 | 1795 | 7553 | biological regulation |
| GO:00023 | 2.09E-09 | 1.42 | 399.098 | 497 | 1809 | immune system process |
| GO:00017 | 2.23E-09 | 1.67 | 157.9625 | 224 | 716 | cell activation |
| GO:00009 | 2.28E-09 | 2.57 | 39.71125 | 75 | 180 | nuclear-transcribed mRNA catabolic process |
| GO:00164 | 7.02E-09 | 1.63 | 163.2573 | 228 | 740 | cell migration |
| GO:00488 | 8.96E-09 | 1.60 | 176.4944 | 243 | 800 | cell motility |
| GO:00516 | 8.96E-09 | 1.60 | 176.4944 | 243 | 800 | localization of cell |
| GO:00069 | 1.02E-08 | 1.51 | 235.8407 | 311 | 1069 | immune response |
| GO:00507 | 1.07E-08 | 1.30 | 1590.877 | 1714 | 7211 | regulation of biological process |
| GO:00603 | 1.72E-08 | 2.68 | 31.98961 | 62 | 145 | cell chemotaxis |
| GO:00512 | 1.72E-08 | 1.42 | 335.3394 | 421 | 1520 | regulation of multicellular organismal process |
| GO:00453 | 1.91E-08 | 1.73 | 117.81 | 172 | 534 | leukocyte activation |
| GO:00325 | 1.99E-08 | 1.30 | 804.1527 | 920 | 3645 | developmental process |
| GO:00071 | 2.19E-08 | 1.63 | 152.4471 | 213 | 691 | cell adhesion |
| GO:00400 | 2.33E-08 | 1.50 | 229.6634 | 302 | 1041 | locomotion |
| GO:00022 | 2.63E-08 | 2.99 | 24.70922 | 51 | 112 | myeloid leukocyte activation |
| GO:00230 | 2.71E-08 | 1.29 | 863.2784 | 980 | 3913 | signaling |
| GO:00447 | 2.71E-08 | 1.29 | 863.2784 | 980 | 3913 | single organism signaling |
| GO:00226 | 2.83E-08 | 1.62 | 152.8883 | 213 | 693 | biological adhesion |
| GO:00447 | 3.53E-08 | 1.29 | 794.6662 | 908 | 3602 | single-organism developmental process |
| GO:00064 | 4.83E-08 | 2.21 | 48.09473 | 83 | 218 | RNA catabolic process |
| GO:00096 | 5.19E-08 | 1.71 | 115.1626 | 167 | 522 | response to biotic stimulus |
| GO:00071 | 5.68E-08 | 1.28 | 876.7361 | 991 | 3974 | cell communication |
| GO:00432 | 6.30E-08 | 1.72 | 109.6472 | 160 | 497 | response to external biotic stimulus |
| GO:00517 | 6.30E-08 | 1.72 | 109.6472 | 160 | 497 | response to other organism |
| GO:00508 | 1.84E-07 | 1.86 | 75.23075 | 116 | 341 | regulation of cell activation |
| GO:00488 | 1.86E-07 | 1.28 | 709.5076 | 813 | 3216 | anatomical structure development |
| GO:00071 | 2.00E-07 | 1.27 | 794.4455 | 901 | 3601 | signal transduction |
| GO:00901 | 2.28E-07 | 1.96 | 61.99367 | 99 | 281 | establishment of protein localization to membrane |
| GO:00487 | 4.80E-07 | 1.29 | 592.8007 | 687 | 2687 | system development |
| GO:00506 | 5.62E-07 | 2.90 | 21.17933 | 43 | 96 | cytokine secretion |
| GO:00329 | 6.07E-07 | 2.21 | 39.93186 | 69 | 181 | mononuclear cell proliferation |
| GO:00485 | 6.99E-07 | 1.32 | 430.4258 | 513 | 1951 | organ development |
| GO:00026 | 7.53E-07 | 1.83 | 70.59777 | 108 | 320 | regulation of leukocyte activation |
| GO:00712 | 7.67E-07 | 2.81 | 22.0618 | 44 | 100 | cellular response to molecule of bacterial origin |
| GO:00726 | 7.98E-07 | 1.79 | 76.33384 | 115 | 346 | protein localization to membrane |
| GO:00712 | 8.23E-07 | 2.88 | 20.7381 | 42 | 94 | cellular response to lipopolysaccharide |
| GO:00072 | 8.42E-07 | 1.27 | 684.5778 | 781 | 3103 | multicellular organismal development |
| GO:00422 | 8.76E-07 | 1.29 | 530.5864 | 619 | 2405 | response to chemical |
| GO:00466 | 1.07E-06 | 2.18 | 39.71125 | 68 | 180 | lymphocyte proliferation |
| GO:00030 | 1.09E-06 | 1.47 | 192.1583 | 250 | 871 | system process |
| GO:00022 | 1.11E-06 | 2.14 | 41.25557 | 70 | 187 | adaptive immune response |
| GO:00469 | 1.19E-06 | 1.54 | 143.4017 | 194 | 650 | secretion |
| GO:00650 | 1.19E-06 | 1.30 | 490.2133 | 575 | 2222 | regulation of biological quality |
| GO:00305 | 1.33E-06 | 2.65 | 23.82675 | 46 | 108 | leukocyte chemotaxis |
| GO:00421 | 1.39E-06 | 1.44 | 213.9995 | 274 | 970 | regulation of cell proliferation |
| GO:00507 | 1.68E-06 | 1.24 | 1519.396 | 1623 | 6887 | regulation of cellular process |
| GO:19025 | 1.83E-06 | 1.70 | 87.36474 | 127 | 396 | single-organism localization |
| GO:19025 | 1.83E-06 | 1.70 | 87.36474 | 127 | 396 | single-organism cellular localization |
| GO:00082 | 2.02E-06 | 1.37 | 285.2591 | 352 | 1293 | cell proliferation |
| GO:00706 | 2.17E-06 | 2.09 | 41.91743 | 70 | 190 | leukocyte proliferation |
| GO:00716 | 2.23E-06 | 3.84 | 11.47214 | 27 | 52 | granulocyte chemotaxis |
| GO:00069 | 2.44E-06 | 1.39 | 259.4468 | 323 | 1176 | cellular component movement |
| GO:00022 | 2.98E-06 | 4.04 | 10.36905 | 25 | 47 | myeloid cell activation involved in immune response |
| GO:00975 | 3.84E-06 | 3.56 | 12.35461 | 28 | 56 | granulocyte migration |
| GO:00488 | 4.10E-06 | 1.27 | 543.8235 | 627 | 2465 | cellular developmental process |
| GO:00305 | 4.42E-06 | 4.07 | 9.927812 | 24 | 45 | neutrophil chemotaxis |
| GO:00339 | 4.52E-06 | 1.60 | 103.9111 | 145 | 471 | response to lipid |
| GO:00488 | 4.61E-06 | 1.54 | 125.311 | 170 | 568 | chemical homeostasis |
| GO:00024 | 4.87E-06 | 2.13 | 36.62259 | 62 | 166 | adaptive immune response based on somatic recombination of immune receptors built from immunoglobulin superfamily domains |
| GO:00450 | 5.68E-06 | 1.48 | 151.7852 | 200 | 688 | innate immune response |
| GO:00507 | 5.73E-06 | 2.85 | 17.87006 | 36 | 81 | regulation of cytokine secretion |
| GO:00508 | 5.99E-06 | 1.66 | 86.70289 | 124 | 393 | ion homeostasis |
| GO:00190 | 6.57E-06 | 1.79 | 63.75861 | 96 | 289 | viral life cycle |
| GO:00069 | 6.97E-06 | 2.67 | 20.07624 | 39 | 91 | humoral immune response |
| GO:00517 | 7.11E-06 | 1.32 | 340.4136 | 408 | 1543 | multi-organism process |
| GO:19902 | 7.31E-06 | 3.88 | 10.14843 | 24 | 46 | neutrophil migration |
| GO:00329 | 8.39E-06 | 2.22 | 31.10714 | 54 | 141 | regulation of mononuclear cell proliferation |
| GO:00506 | 8.39E-06 | 2.22 | 31.10714 | 54 | 141 | regulation of lymphocyte proliferation |

| | | | | | | |
|---|---|---|---|---|---|---|
| GO:00022 | 9.22E-06 | 2.23 | 30.44529 | 53 | 138 | cell activation involved in immune response |
| GO:00023 | 9.22E-06 | 2.23 | 30.44529 | 53 | 138 | leukocyte activation involved in immune response |
| GO:00485 | 9.54E-06 | 1.26 | 531.9101 | 611 | 2411 | regulation of response to stimulus |
| GO:00706 | 9.69E-06 | 2.18 | 31.98961 | 55 | 145 | regulation of leukocyte proliferation |
| GO:00324 | 9.82E-06 | 31.90 | 2.20618 | 9 | 10 | response to peptidoglycan |
| GO:00427 | 1.06E-05 | 2.50 | 22.50304 | 42 | 102 | defense response to bacterium |
| GO:00069 | 1.06E-05 | 1.59 | 98.61626 | 137 | 447 | chemotaxis |
| GO:00423 | 1.06E-05 | 1.59 | 98.61626 | 137 | 447 | taxis |
| GO:00327 | 1.13E-05 | 5.49 | 6.177305 | 17 | 28 | positive regulation of interleukin-6 production |
| GO:00510 | 1.14E-05 | 1.51 | 127.076 | 170 | 576 | positive regulation of developmental process |
| GO:00485 | 1.15E-05 | 1.51 | 128.8409 | 172 | 584 | hematopoietic or lymphoid organ development |
| GO:00517 | 1.17E-05 | 1.21 | 975.7936 | 1069 | 4423 | cellular response to stimulus |
| GO:00351 | 1.23E-05 | 2.82 | 16.98759 | 34 | 77 | regulation of tube size |
| GO:00508 | 1.23E-05 | 2.82 | 16.98759 | 34 | 77 | regulation of blood vessel size |
| GO:00509 | 1.25E-05 | 1.85 | 52.50709 | 81 | 238 | leukocyte migration |
| GO:00329 | 1.27E-05 | 1.51 | 127.2966 | 170 | 577 | secretion by cell |
| GO:00466 | 1.29E-05 | 1.57 | 102.3668 | 141 | 464 | lymphocyte activation |
| GO:00712 | 1.34E-05 | 2.35 | 25.59169 | 46 | 116 | cellular response to biotic stimulus |
| GO:00028 | 1.48E-05 | 7.10 | 4.632979 | 14 | 21 | regulation of myeloid leukocyte mediated immunity |
| GO:00343 | 1.52E-05 | 2.48 | 22.0618 | 41 | 100 | response to interferon-gamma |
| GO:19030 | 1.59E-05 | 1.83 | 53.61018 | 82 | 243 | regulation of response to wounding |
| GO:00068 | 1.59E-05 | 1.72 | 66.62665 | 98 | 302 | cellular ion homeostasis |
| GO:00550 | 1.60E-05 | 1.65 | 80.7462 | 115 | 366 | cellular chemical homeostasis |
| GO:00708 | 1.81E-05 | 1.29 | 369.5352 | 436 | 1675 | cellular response to chemical stimulus |
| GO:00550 | 1.82E-05 | 1.66 | 76.77508 | 110 | 348 | cation homeostasis |
| GO:00507 | 2.05E-05 | 1.33 | 281.0674 | 340 | 1274 | regulation of developmental process |
| GO:00447 | 2.10E-05 | 1.24 | 566.5471 | 644 | 2568 | single-organism transport |
| GO:00975 | 2.13E-05 | 2.62 | 18.75253 | 36 | 85 | myeloid leukocyte migration |
| GO:00300 | 2.57E-05 | 1.71 | 64.8617 | 95 | 294 | cellular cation homeostasis |
| GO:00447 | 2.57E-05 | 1.23 | 1872.826 | 1955 | 8489 | single-organism cellular process |
| GO:00420 | 3.01E-05 | 1.51 | 115.1626 | 154 | 522 | wound healing |
| GO:00022 | 3.07E-05 | 1.53 | 106.5585 | 144 | 483 | immune effector process |
| GO:00300 | 3.20E-05 | 1.49 | 123.9873 | 164 | 562 | hemopoiesis |
| GO:00301 | 3.38E-05 | 1.25 | 505.6565 | 578 | 2292 | cell differentiation |
| GO:20000 | 3.77E-05 | 1.37 | 207.1603 | 257 | 939 | regulation of multicellular organismal development |
| GO:00986 | 3.77E-05 | 1.74 | 58.02254 | 86 | 263 | single organism cell adhesion |
| GO:00510 | 3.78E-05 | 1.62 | 79.64311 | 112 | 361 | regulation of secretion |
| GO:00316 | 3.91E-05 | 3.73 | 9.045339 | 21 | 41 | lipopolysaccharide-mediated signaling pathway |
| GO:00104 | 4.01E-05 | 2.12 | 29.56282 | 50 | 134 | negative regulation of peptidase activity |
| GO:00512 | 4.81E-05 | 1.69 | 63.31738 | 92 | 287 | regulation of lymphocyte activation |
| GO:00030 | 4.83E-05 | 2.44 | 20.07624 | 37 | 91 | vascular process in circulatory system |
| GO:00420 | 5.05E-05 | 2.65 | 16.54635 | 32 | 75 | regulation of cytokine biosynthetic process |
| GO:00163 | 5.11E-05 | 1.79 | 50.30091 | 76 | 228 | single organismal cell-cell adhesion |
| GO:00713 | 5.11E-05 | 1.79 | 50.30091 | 76 | 228 | cellular response to lipid |
| GO:00024 | 5.25E-05 | 1.86 | 43.90299 | 68 | 199 | leukocyte mediated immunity |
| GO:00080 | 5.25E-05 | 1.74 | 56.03698 | 83 | 254 | blood circulation |
| GO:00516 | 5.62E-05 | 1.98 | 35.29889 | 57 | 160 | detection of stimulus |
| GO:00326 | 5.88E-05 | 2.79 | 14.56079 | 29 | 66 | regulation of interferon-gamma production |
| GO:00550 | 6.03E-05 | 1.64 | 69.49468 | 99 | 315 | metal ion homeostasis |
| GO:00525 | 6.13E-05 | 1.73 | 56.2576 | 83 | 255 | regulation of peptidase activity |
| GO:00069 | 6.28E-05 | 1.22 | 586.6234 | 660 | 2659 | response to stress |
| GO:00018 | 6.44E-05 | 1.56 | 87.36474 | 120 | 396 | regulation of cytokine production |
| GO:19025 | 6.62E-05 | 1.32 | 259.0056 | 312 | 1174 | regulation of intracellular signal transduction |
| GO:00321 | 6.95E-05 | 2.08 | 29.3422 | 49 | 133 | positive regulation of response to external stimulus |
| GO:00030 | 7.14E-05 | 1.72 | 56.47822 | 83 | 256 | circulatory system process |
| GO:00321 | 7.19E-05 | 1.53 | 94.42452 | 128 | 428 | regulation of response to external stimulus |
| GO:00026 | 7.23E-05 | 1.46 | 125.7523 | 164 | 570 | positive regulation of immune system process |
| GO:00025 | 7.27E-05 | 1.44 | 136.3419 | 176 | 618 | immune system development |
| GO:00024 | 7.94E-05 | 3.99 | 7.501013 | 18 | 34 | humoral immune response mediated by circulating immunoglobulin |
| GO:00066 | 8.20E-05 | 2.72 | 14.78141 | 29 | 67 | icosanoid metabolic process |
| GO:19015 | 8.20E-05 | 2.72 | 14.78141 | 29 | 67 | fatty acid derivative metabolic process |
| GO:19030 | 8.20E-05 | 2.72 | 14.78141 | 29 | 67 | positive regulation of response to wounding |
| GO:00018 | 8.40E-05 | 1.52 | 98.17503 | 132 | 445 | cytokine production |
| GO:00001 | 8.76E-05 | 1.50 | 102.5874 | 137 | 465 | MAPK cascade |
| GO:00511 | 8.89E-05 | 1.19 | 844.9671 | 925 | 3830 | localization |
| GO:00985 | 8.97E-05 | 1.69 | 58.46378 | 85 | 265 | defense response to other organism |
| GO:00230 | 9.06E-05 | 1.49 | 106.9997 | 142 | 485 | signal transduction by phosphorylation |
| GO:00082 | 9.42E-05 | 1.47 | 114.0595 | 150 | 517 | positive regulation of cell proliferation |
| GO:00507 | 9.48E-05 | 1.88 | 39.04939 | 61 | 177 | regulation of inflammatory response |
| GO:00330 | 9.48E-05 | 5.76 | 4.632979 | 13 | 21 | regulation of mast cell activation |
| GO:00433 | 9.48E-05 | 5.76 | 4.632979 | 13 | 21 | regulation of leukocyte degranulation |
| GO:19033 | 9.48E-05 | 5.76 | 4.632979 | 13 | 21 | regulation of regulated secretory pathway |
| GO:00507 | 9.57E-05 | 2.96 | 12.13399 | 25 | 55 | positive regulation of inflammatory response |
| GO:00109 | 9.59E-05 | 2.06 | 28.90096 | 48 | 131 | negative regulation of endopeptidase activity |
| GO:00464 | 9.61E-05 | 3.39 | 9.486575 | 21 | 43 | icosanoid biosynthetic process |
| GO:19015 | 9.61E-05 | 3.39 | 9.486575 | 21 | 43 | fatty acid derivative biosynthetic process |
| GO:00326 | 9.76E-05 | 3.05 | 11.47214 | 24 | 52 | chemokine production |
| GO:00024 | 9.80E-05 | 3.26 | 10.14843 | 22 | 46 | myeloid leukocyte mediated immunity |
| GO:00421 | 9.88E-05 | 2.40 | 19.19377 | 35 | 87 | cytokine metabolic process |
| GO:00703 | 0.000102 | 2.16 | 25.15046 | 43 | 114 | ERK1 and ERK2 cascade |
| GO:00434 | 0.000107 | 1.67 | 59.56687 | 86 | 270 | positive regulation of MAPK cascade |
| GO:00433 | 0.000111 | 8.86 | 3.088652 | 10 | 14 | regulation of mast cell degranulation |
| GO:20011 | 0.000115 | Inf | 1.323708 | 6 | 6 | regulation of interleukin-10 secretion |
| GO:00508 | 0.000116 | 1.75 | 48.9772 | 73 | 222 | positive regulation of cell activation |
| GO:00140 | 0.000121 | 1.49 | 103.2492 | 137 | 468 | response to organic cyclic compound |
| GO:00071 | 0.000121 | 1.23 | 454.0319 | 518 | 2058 | cell surface receptor signaling pathway |
| GO:00068 | 0.000124 | 1.66 | 59.78749 | 86 | 271 | cellular metal ion homeostasis |
| GO:00508 | 0.000125 | 2.48 | 17.20821 | 32 | 78 | regulation of B cell activation |
| GO:00068 | 0.000133 | 1.52 | 90.45339 | 122 | 410 | endocytosis |
| GO:00703 | 0.000136 | 2.19 | 23.16489 | 40 | 105 | regulation of ERK1 and ERK2 cascade |
| GO:00328 | 0.000139 | 1.28 | 303.1292 | 357 | 1374 | regulation of localization |
| GO:00725 | 0.000139 | 1.79 | 44.34422 | 67 | 201 | cellular divalent inorganic cation homeostasis |
| GO:00420 | 0.000141 | 2.37 | 18.75253 | 34 | 85 | cytokine biosynthetic process |
| GO:00525 | 0.00015 | 1.69 | 54.27204 | 79 | 246 | regulation of endopeptidase activity |
| GO:00068 | 0.000151 | 1.34 | 202.9686 | 248 | 920 | ion transport |
| GO:00512 | 0.000153 | 1.51 | 93.32143 | 125 | 423 | positive regulation of multicellular organismal process |
| GO:00075 | 0.000155 | 1.50 | 94.2039 | 126 | 427 | blood coagulation |
| GO:00725 | 0.000162 | 1.75 | 46.99164 | 70 | 213 | divalent inorganic cation homeostasis |

| | | | | | | |
|---|---|---|---|---|---|---|
| GO:00440 | 0.000173 | 1.71 | 51.18338 | 75 | 232 | regulation of system process |
| GO:00075 | 0.000174 | 1.50 | 95.30699 | 127 | 432 | hemostasis |
| GO:00326 | 0.00018 | 2.45 | 16.76697 | 31 | 76 | interferon-gamma production |
| GO:00434 | 0.000185 | 1.51 | 90.23278 | 121 | 409 | regulation of MAPK cascade |
| GO:00725 | 0.000191 | 1.50 | 93.76266 | 125 | 425 | establishment of protein localization to organelle |
| GO:00508 | 0.000192 | 1.49 | 94.64514 | 126 | 429 | coagulation |
| GO:00326 | 0.000194 | 3.77 | 7.280395 | 17 | 33 | interleukin-10 production |
| GO:00327 | 0.000206 | 3.55 | 7.942249 | 18 | 36 | positive regulation of interferon-gamma production |
| GO:00423 | 0.000206 | 3.55 | 7.942249 | 18 | 36 | vasodilation |
| GO:00508 | 0.000208 | 1.45 | 111.4121 | 145 | 505 | regulation of body fluid levels |
| GO:00801 | 0.000208 | 1.36 | 170.3171 | 211 | 772 | regulation of response to stress |
| GO:00903 | 0.000214 | 3.37 | 8.604103 | 19 | 39 | phagosome maturation |
| GO:00068 | 0.000216 | 1.79 | 41.69681 | 63 | 189 | cellular calcium ion homeostasis |
| GO:00512 | 0.000219 | 1.48 | 98.39564 | 130 | 446 | regulation of cellular component movement |
| GO:00072 | 0.000224 | 2.00 | 28.23911 | 46 | 128 | positive regulation of cytosolic calcium ion concentration |
| GO:00069 | 0.000229 | 1.89 | 33.75456 | 53 | 153 | phagocytosis |
| GO:00432 | 0.000233 | 1.58 | 68.39159 | 95 | 310 | regulation of ion transport |
| GO:00550 | 0.00024 | 1.76 | 43.46175 | 65 | 197 | calcium ion homeostasis |
| GO:00718 | 0.000253 | 1.28 | 262.5355 | 311 | 1190 | protein complex subunit organization |
| GO:00706 | 0.000255 | 2.21 | 20.7381 | 36 | 94 | positive regulation of leukocyte proliferation |
| GO:00026 | 0.000258 | 1.71 | 47.6535 | 70 | 216 | positive regulation of leukocyte activation |
| GO:00093 | 0.000264 | 1.85 | 35.5195 | 55 | 161 | protein secretion |
| GO:00026 | 0.000266 | 2.15 | 22.28242 | 38 | 101 | positive regulation of immune effector process |
| GO:00330 | 0.000267 | 7.09 | 3.309271 | 10 | 15 | regulation of mast cell activation involved in immune response |
| GO:00313 | 0.000269 | 1.68 | 51.84524 | 75 | 235 | positive regulation of defense response |
| GO:00025 | 0.00027 | 1.54 | 76.33384 | 104 | 346 | leukocyte differentiation |
| GO:00326 | 0.000291 | 3.79 | 6.839159 | 16 | 31 | regulation of interleukin-10 production |
| GO:00421 | 0.000297 | 2.82 | 11.47214 | 23 | 52 | B cell proliferation |
| GO:00507 | 0.000297 | 2.82 | 11.47214 | 23 | 52 | positive regulation of cytokine secretion |
| GO:00603 | 0.000301 | 2.49 | 15.00203 | 28 | 68 | interferon-gamma-mediated signaling pathway |
| GO:00450 | 0.000317 | 2.98 | 10.14843 | 21 | 46 | regulated secretory pathway |
| GO:00326 | 0.00032 | 3.36 | 8.162867 | 18 | 37 | interleukin-1 production |
| GO:00421 | 0.00032 | 3.36 | 8.162867 | 18 | 37 | macrophage activation |
| GO:00432 | 0.00032 | 3.36 | 8.162867 | 18 | 37 | leukocyte degranulation |
| GO:00226 | 0.000335 | 1.42 | 116.0451 | 149 | 526 | regulation of anatomical structure morphogenesis |
| GO:00340 | 0.000335 | 1.44 | 106.3379 | 138 | 482 | response to cytokine |
| GO:00026 | 0.000337 | 1.31 | 204.5129 | 247 | 927 | regulation of immune system process |
| GO:00083 | 0.000339 | 2.26 | 18.75253 | 33 | 85 | regulation of cell shape |
| GO:00329 | 0.000358 | 2.19 | 20.29686 | 35 | 92 | positive regulation of mononuclear cell proliferation |
| GO:00506 | 0.000358 | 2.19 | 20.29686 | 35 | 92 | positive regulation of lymphocyte proliferation |
| GO:00441 | 0.000364 | 7.97 | 2.868034 | 9 | 13 | regulation of growth of symbiont in host |
| GO:00441 | 0.000364 | 7.97 | 2.868034 | 9 | 13 | negative regulation of growth of symbiont in host |
| GO:00441 | 0.000364 | 7.97 | 2.868034 | 9 | 13 | modulation of growth of symbiont involved in interaction with host |
| GO:00441 | 0.000364 | 7.97 | 2.868034 | 9 | 13 | negative regulation of growth of symbiont involved in interaction with host |
| GO:00096 | 0.000365 | 1.24 | 360.2693 | 414 | 1633 | anatomical structure morphogenesis |
| GO:00713 | 0.000372 | 2.28 | 18.09068 | 32 | 82 | cellular response to interferon-gamma |
| GO:00507 | 0.000379 | 1.99 | 26.47416 | 43 | 120 | regulation of protein secretion |
| GO:00066 | 0.000395 | 4.14 | 5.736069 | 14 | 26 | leukotriene metabolic process |
| GO:00725 | 0.0004 | 1.65 | 51.62462 | 74 | 234 | divalent inorganic cation transport |
| GO:19017 | 0.000406 | 1.32 | 195.0263 | 236 | 884 | response to oxygen-containing compound |
| GO:00098 | 0.000413 | 1.28 | 242.018 | 287 | 1097 | tissue development |
| GO:00421 | 0.000415 | 1.53 | 74.5689 | 101 | 338 | T cell activation |
| GO:00197 | 0.000435 | 2.21 | 18.97315 | 33 | 86 | B cell mediated immunity |
| GO:00327 | 0.000436 | 3.80 | 6.397923 | 15 | 29 | positive regulation of chemokine production |
| GO:00708 | 0.000446 | 1.65 | 50.96277 | 73 | 231 | divalent metal ion transport |
| GO:00323 | 0.000464 | 1.96 | 26.69478 | 43 | 121 | positive regulation of Rho GTPase activity |
| GO:00308 | 0.00047 | 2.96 | 9.707194 | 20 | 44 | regulation of B cell proliferation |
| GO:00455 | 0.000478 | 3.35 | 7.721631 | 17 | 35 | mast cell activation |
| GO:00160 | 0.000479 | 2.23 | 18.3113 | 32 | 83 | immunoglobulin mediated immune response |
| GO:00022 | 0.000479 | 9.45 | 2.426798 | 8 | 11 | neutrophil activation involved in immune response |
| GO:00703 | 0.000486 | 2.34 | 16.10512 | 29 | 73 | positive regulation of ERK1 and ERK2 cascade |
| GO:00514 | 0.000534 | 1.88 | 30.00405 | 47 | 136 | cytosolic calcium ion homeostasis |
| GO:00193 | 0.00057 | 5.91 | 3.529889 | 10 | 16 | leukotriene biosynthetic process |
| GO:00508 | 0.000581 | 2.06 | 22.28242 | 37 | 101 | negative regulation of cell activation |
| GO:00075 | 0.000582 | 2.01 | 23.82675 | 39 | 108 | female pregnancy |
| GO:00461 | 0.000592 | 4.19 | 5.294833 | 13 | 24 | polyol biosynthetic process |
| GO:00719 | 0.000593 | 1.50 | 76.99569 | 103 | 349 | regulation of protein serine/threonine kinase activity |
| GO:00447 | 0.000597 | 1.88 | 29.3422 | 46 | 133 | multi-multicellular organism process |
| GO:00512 | 0.000605 | 1.68 | 44.78546 | 65 | 203 | positive regulation of lymphocyte activation |
| GO:00303 | 0.000619 | 1.48 | 82.29053 | 109 | 373 | regulation of cell migration |
| GO:00224 | 0.000648 | 1.33 | 166.346 | 203 | 754 | reproductive process |
| GO:00726 | 0.000652 | 21.24 | 1.544326 | 6 | 7 | interleukin-10 secretion |
| GO:00071 | 0.000653 | 1.42 | 103.4699 | 133 | 469 | G-protein coupled receptor signaling pathway |
| GO:00713 | 0.000665 | 1.24 | 301.5849 | 349 | 1367 | cellular response to organic substance |
| GO:00068 | 0.00067 | 1.65 | 47.43288 | 68 | 215 | calcium ion transport |
| GO:00326 | 0.000672 | 2.84 | 9.927812 | 20 | 45 | regulation of chemokine production |
| GO:00100 | 0.000695 | 1.22 | 390.4939 | 443 | 1770 | response to organic substance |
| GO:00015 | 0.000696 | 3.55 | 6.618541 | 15 | 30 | retinoid metabolic process |
| GO:00469 | 0.000696 | 3.55 | 6.618541 | 15 | 30 | regulation of neurotransmitter secretion |
| GO:20001 | 0.000706 | 1.46 | 87.80598 | 115 | 398 | regulation of cell motility |
| GO:00508 | 0.000714 | 3.04 | 8.604103 | 18 | 39 | defense response to Gram-positive bacterium |
| GO:00096 | 0.000717 | 1.98 | 24.04737 | 39 | 109 | response to toxic substance |
| GO:00335 | 0.000743 | 2.22 | 17.20821 | 30 | 78 | unsaturated fatty acid metabolic process |
| GO:00519 | 0.000819 | 6.38 | 3.088652 | 9 | 14 | negative regulation of calcium ion transport |
| GO:00432 | 0.000827 | 2.37 | 14.34017 | 26 | 65 | response to alkaloid |
| GO:00022 | 0.00084 | 2.60 | 11.47214 | 22 | 52 | T cell activation involved in immune response |
| GO:00400 | 0.00087 | 1.42 | 96.18946 | 124 | 436 | regulation of locomotion |
| GO:00024 | 0.000883 | 2.10 | 19.63501 | 33 | 89 | antigen processing and presentation of peptide antigen via MHC class II |
| GO:00066 | 0.000894 | 2.66 | 10.81028 | 21 | 49 | unsaturated fatty acid biosynthetic process |
| GO:00550 | 0.000907 | 1.30 | 182.0099 | 219 | 825 | transmembrane transport |
| GO:00457 | 0.000987 | 2.28 | 15.22264 | 27 | 69 | positive regulation of angiogenesis |
| GO:00507 | 0.000993 | 2.81 | 9.486575 | 19 | 43 | regulation of phagocytosis |
| GO:00485 | 0.001005 | 1.25 | 259.2262 | 302 | 1175 | positive regulation of response to stimulus |
| GO:00604 | 0.00101 | 1.34 | 144.946 | 178 | 657 | epithelium development |
| GO:00423 | 0.001033 | 2.90 | 8.824721 | 18 | 40 | vasoconstriction |
| GO:00075 | 0.001046 | 3.55 | 6.177305 | 14 | 28 | embryo implantation |
| GO:00326 | 0.001075 | 3.33 | 6.839159 | 15 | 31 | regulation of interleukin-1 production |
| GO:00420 | 0.001075 | 3.33 | 6.839159 | 15 | 31 | T-helper 1 type immune response |

| | | | | | | |
|---|---|---|---|---|---|---|
| GO:00326 | 0.001079 | 3.15 | 7.501013 | 16 | 34 | interleukin-1 beta production |
| GO:00456 | 0.001093 | 1.96 | 22.94428 | 37 | 104 | regulation of lymphocyte differentiation |
| GO:00512 | 0.001101 | 1.62 | 46.55041 | 66 | 211 | positive regulation of cellular component movement |
| GO:00025 | 0.001104 | 2.06 | 19.85562 | 33 | 90 | antigen processing and presentation of peptide or polysaccharide antigen via MHC class II |
| GO:00507 | 0.001157 | 1.42 | 92.43896 | 119 | 419 | positive regulation of immune response |
| GO:00022 | 0.001158 | 7.08 | 2.647416 | 8 | 12 | macrophage activation involved in immune response |
| GO:00303 | 0.00116 | 1.64 | 44.12361 | 63 | 200 | positive regulation of cell migration |
| GO:00024 | 0.001201 | 1.73 | 35.07827 | 52 | 159 | lymphocyte mediated immunity |
| GO:00421 | 0.001224 | 2.57 | 11.0309 | 21 | 50 | positive regulation of cytokine biosynthetic process |
| GO:00026 | 0.001231 | 2.02 | 20.7381 | 34 | 94 | negative regulation of leukocyte activation |
| GO:00324 | 0.001335 | 1.94 | 23.16489 | 37 | 105 | regulation of transporter activity |
| GO:00421 | 0.001335 | 4.33 | 4.412361 | 11 | 20 | neutrophil activation |
| GO:00423 | 0.001335 | 4.33 | 4.412361 | 11 | 20 | regulation of vasodilation |
| GO:00486 | 0.00135 | 1.52 | 62.21429 | 84 | 282 | reproductive structure development |
| GO:00422 | 0.001487 | 3.87 | 5.074215 | 12 | 23 | ribosomal small subunit biogenesis |
| GO:00433 | 0.001487 | 3.87 | 5.074215 | 12 | 23 | mast cell degranulation |
| GO:00313 | 0.001529 | 1.41 | 93.10081 | 119 | 422 | regulation of defense response |
| GO:00019 | 0.001577 | 3.55 | 5.736069 | 13 | 26 | blood vessel remodeling |
| GO:00096 | 0.001577 | 3.55 | 5.736069 | 13 | 26 | response to fungus |
| GO:00072 | 0.001585 | 2.99 | 7.721631 | 16 | 35 | glutamate receptor signaling pathway |
| GO:00329 | 0.001601 | 8.26 | 2.20618 | 7 | 10 | inositol phosphate biosynthetic process |
| GO:00357 | 0.001601 | 8.26 | 2.20618 | 7 | 10 | endothelial cell chemotaxis |
| GO:00512 | 0.001603 | 1.52 | 58.24316 | 79 | 264 | negative regulation of multicellular organismal process |
| GO:00970 | 0.001616 | 3.31 | 6.397923 | 14 | 29 | dendritic cell differentiation |
| GO:00441 | 0.001646 | 5.31 | 3.309271 | 9 | 15 | growth involved in symbiotic interaction |
| GO:00441 | 0.001646 | 5.31 | 3.309271 | 9 | 15 | growth of symbiont involved in interaction with host |
| GO:00441 | 0.001646 | 5.31 | 3.309271 | 9 | 15 | growth of symbiont in host |
| GO:19021 | 0.001665 | 1.94 | 21.84119 | 35 | 99 | positive regulation of leukocyte differentiation |
| GO:00022 | 0.001679 | 1.44 | 78.3194 | 102 | 355 | activation of immune response |
| GO:00300 | 0.00178 | 1.56 | 50.74215 | 70 | 230 | lymphocyte differentiation |
| GO:00323 | 0.001795 | 1.74 | 31.54838 | 47 | 143 | regulation of Rho GTPase activity |
| GO:00106 | 0.001796 | 1.18 | 451.3845 | 502 | 2046 | regulation of cell communication |
| GO:00072 | 0.001805 | 1.39 | 97.95441 | 124 | 444 | small GTPase mediated signal transduction |
| GO:00310 | 0.001892 | 2.05 | 18.09068 | 30 | 82 | regeneration |
| GO:00421 | 0.001904 | 1.68 | 35.74012 | 52 | 162 | B cell activation |
| GO:00347 | 0.001948 | 1.60 | 44.12361 | 62 | 200 | regulation of transmembrane transport |
| GO:19017 | 0.001972 | 1.33 | 131.4883 | 161 | 596 | cellular response to oxygen-containing compound |
| GO:00302 | 0.001975 | 1.67 | 36.62259 | 53 | 166 | T cell differentiation |
| GO:00456 | 0.001987 | 2.22 | 14.34017 | 25 | 65 | positive regulation of lymphocyte differentiation |
| GO:00717 | 0.002009 | 4.43 | 3.971125 | 10 | 18 | icosanoid transport |
| GO:19004 | 0.002009 | 4.43 | 3.971125 | 10 | 18 | regulation of glutamate receptor signaling pathway |
| GO:19015 | 0.002009 | 4.43 | 3.971125 | 10 | 18 | fatty acid derivative transport |
| GO:00301 | 0.002013 | 1.55 | 50.96277 | 70 | 231 | extracellular matrix organization |
| GO:00430 | 0.002013 | 1.55 | 50.96277 | 70 | 231 | extracellular structure organization |
| GO:00510 | 0.002028 | 1.91 | 22.0618 | 35 | 100 | negative regulation of secretion |
| GO:00308 | 0.002031 | 1.44 | 75.23075 | 98 | 341 | epithelial cell differentiation |
| GO:00068 | 0.002077 | 1.54 | 52.72771 | 72 | 239 | exocytosis |
| GO:00198 | 0.002105 | 2.00 | 18.97315 | 31 | 86 | antigen processing and presentation of exogenous peptide antigen via MHC class II |
| GO:00614 | 0.00211 | 1.48 | 63.09676 | 84 | 286 | reproductive system development |
| GO:00510 | 0.002112 | 1.65 | 38.38754 | 55 | 174 | positive regulation of secretion |
| GO:00000 | 0.002118 | 10.62 | 1.764944 | 6 | 8 | ribosomal small subunit assembly |
| GO:00028 | 0.002118 | 10.62 | 1.764944 | 6 | 8 | positive regulation of inflammatory response to antigenic stimulus |
| GO:00353 | 0.002118 | 10.62 | 1.764944 | 6 | 8 | wound healing, spreading of epidermal cells |
| GO:00601 | 0.002118 | 10.62 | 1.764944 | 6 | 8 | regulation of posttranscriptional gene silencing |
| GO:00609 | 0.002118 | 10.62 | 1.764944 | 6 | 8 | regulation of gene silencing by miRNA |
| GO:00609 | 0.002118 | 10.62 | 1.764944 | 6 | 8 | regulation of gene silencing by RNA |
| GO:00447 | 0.002138 | 1.30 | 151.7852 | 183 | 688 | single organism reproductive process |
| GO:00446 | 0.00216 | 1.17 | 2056.16 | 2109 | 9320 | single-organism process |
| GO:00020 | 0.002166 | 2.74 | 8.604103 | 17 | 39 | sprouting angiogenesis |
| GO:00140 | 0.002166 | 2.74 | 8.604103 | 17 | 39 | regulation of gliogenesis |
| GO:00072 | 0.002174 | 1.30 | 156.4182 | 188 | 709 | cell-cell signaling |
| GO:00326 | 0.002191 | 2.24 | 13.67832 | 24 | 62 | regulation of interleukin-6 production |
| GO:00455 | 0.002197 | 1.39 | 93.10081 | 118 | 422 | positive regulation of cell differentiation |
| GO:00513 | 0.002238 | 1.55 | 50.30091 | 69 | 228 | negative regulation of hydrolase activity |
| GO:00095 | 0.002248 | 3.90 | 4.632979 | 11 | 21 | detection of biotic stimulus |
| GO:20001 | 0.002271 | 1.58 | 45.2267 | 63 | 205 | positive regulation of cell motility |
| GO:00508 | 0.002275 | 2.84 | 7.942249 | 16 | 36 | positive regulation of synaptic transmission |
| GO:00171 | 0.002293 | 2.15 | 15.22264 | 26 | 69 | regulation of exocytosis |
| GO:00030 | 0.002318 | 1.57 | 46.10917 | 64 | 209 | muscle system process |
| GO:00082 | 0.00232 | 1.96 | 19.85562 | 32 | 90 | regulation of blood pressure |
| GO:00102 | 0.002333 | 1.35 | 112.074 | 139 | 508 | response to organonitrogen compound |
| GO:00347 | 0.002363 | 1.60 | 41.91743 | 59 | 190 | regulation of ion transmembrane transport |
| GO:00023 | 0.002365 | Inf | 0.882472 | 4 | 4 | MHC class II protein complex assembly |
| GO:00025 | 0.002365 | Inf | 0.882472 | 4 | 4 | peptide antigen assembly with MHC class II protein complex |
| GO:00148 | 0.002365 | Inf | 0.882472 | 4 | 4 | artery smooth muscle contraction |
| GO:00324 | 0.002365 | Inf | 0.882472 | 4 | 4 | positive regulation of sodium:proton antiporter activity |
| GO:00343 | 0.002365 | Inf | 0.882472 | 4 | 4 | 'de novo' NAD biosynthetic process from tryptophan |
| GO:00346 | 0.002365 | Inf | 0.882472 | 4 | 4 | 'de novo' NAD biosynthetic process |
| GO:00432 | 0.002365 | Inf | 0.882472 | 4 | 4 | sodium-independent organic anion transport |
| GO:00607 | 0.002365 | Inf | 0.882472 | 4 | 4 | positive regulation of inositol phosphate biosynthetic process |
| GO:00712 | 0.002365 | Inf | 0.882472 | 4 | 4 | cellular response to peptidoglycan |
| GO:00022 | 0.002382 | 3.54 | 5.294833 | 12 | 24 | mast cell activation involved in immune response |
| GO:00024 | 0.002382 | 3.54 | 5.294833 | 12 | 24 | mast cell mediated immunity |
| GO:00508 | 0.002382 | 3.54 | 5.294833 | 12 | 24 | negative regulation of B cell activation |
| GO:00023 | 0.002398 | 2.45 | 10.81028 | 20 | 49 | cytokine production involved in immune response |
| GO:00713 | 0.00243 | 1.39 | 89.79154 | 114 | 407 | cellular response to cytokine stimulus |
| GO:00094 | 0.00243 | 5.67 | 2.868034 | 8 | 13 | NAD biosynthetic process |
| GO:00156 | 0.00243 | 5.67 | 2.868034 | 8 | 13 | organic cation transport |
| GO:00440 | 0.00243 | 5.67 | 2.868034 | 8 | 13 | positive regulation of cellular component biogenesis |
| GO:00469 | 0.002432 | 1.63 | 38.60816 | 55 | 175 | carboxylic acid transport |
| GO:00030 | 0.002437 | 1.39 | 88.90907 | 113 | 403 | developmental process involved in reproduction |
| GO:00300 | 0.002467 | 1.36 | 104.1317 | 130 | 472 | metal ion transport |
| GO:00507 | 0.00248 | 1.31 | 138.5481 | 168 | 628 | regulation of immune response |
| GO:00026 | 0.002489 | 1.54 | 49.63906 | 68 | 225 | regulation of immune effector process |
| GO:00066 | 0.002495 | 1.36 | 102.3668 | 128 | 464 | protein targeting |
| GO:00487 | 0.002535 | 1.92 | 20.7381 | 33 | 94 | tissue remodeling |
| GO:00192 | 0.002548 | 1.21 | 313.9395 | 356 | 1423 | regulation of phosphate metabolic process |
| GO:00109 | 0.002553 | 17.70 | 1.323708 | 5 | 6 | regulation of inositol phosphate biosynthetic process |

| | | | | | | |
|---|---|---|---|---|---|---|
| GO:00326 | 0.002553 | 17.70 | 1.323708 | 5 | 6 | interleukin-18 production |
| GO:00703 | 0.002553 | 17.70 | 1.323708 | 5 | 6 | response to lipoteichoic acid |
| GO:00712 | 0.002553 | 17.70 | 1.323708 | 5 | 6 | cellular response to lipoteichoic acid |
| GO:00970 | 0.002553 | 17.70 | 1.323708 | 5 | 6 | dendritic cell apoptotic process |
| GO:20006 | 0.002553 | 17.70 | 1.323708 | 5 | 6 | regulation of dendritic cell apoptotic process |
| GO:00355 | 0.002562 | 1.19 | 392.9207 | 439 | 1781 | intracellular signal transduction |
| GO:00072 | 0.002604 | 2.08 | 16.10512 | 27 | 73 | neurotransmitter secretion |
| GO:00109 | 0.002636 | 1.64 | 37.06383 | 53 | 168 | regulation of metal ion transport |
| GO:19025 | 0.002754 | 1.33 | 120.6781 | 148 | 547 | positive regulation of intracellular signal transduction |
| GO:00158 | 0.002793 | 1.62 | 38.82877 | 55 | 176 | organic acid transport |
| GO:00519 | 0.002829 | 1.83 | 24.04737 | 37 | 109 | regulation of calcium ion transport |
| GO:00323 | 0.002884 | 1.52 | 51.62462 | 70 | 234 | positive regulation of Ras GTPase activity |
| GO:00513 | 0.002995 | 1.25 | 205.616 | 240 | 932 | regulation of hydrolase activity |
| GO:00230 | 0.002998 | 1.17 | 450.2814 | 498 | 2041 | regulation of signaling |
| GO:00193 | 0.00303 | 4.55 | 3.529889 | 9 | 16 | nicotinamide nucleotide biosynthetic process |
| GO:00193 | 0.00303 | 4.55 | 3.529889 | 9 | 16 | pyridine nucleotide biosynthetic process |
| GO:00323 | 0.00303 | 4.55 | 3.529889 | 9 | 16 | icosanoid secretion |
| GO:00425 | 0.00303 | 4.55 | 3.529889 | 9 | 16 | superoxide anion generation |
| GO:00421 | 0.003169 | 1.83 | 23.38551 | 36 | 106 | regulation of T cell proliferation |
| GO:00095 | 0.003181 | 2.37 | 11.0309 | 20 | 50 | detection of chemical stimulus |
| GO:00311 | 0.003198 | 2.70 | 8.162867 | 16 | 37 | organ regeneration |
| GO:00512 | 0.003234 | 1.98 | 17.87006 | 29 | 81 | negative regulation of lymphocyte activation |
| GO:00420 | 0.003305 | 1.72 | 29.12158 | 43 | 132 | T cell proliferation |
| GO:00161 | 0.003378 | 2.80 | 7.501013 | 15 | 34 | diterpenoid metabolic process |
| GO:00301 | 0.003409 | 3.94 | 4.191743 | 10 | 19 | sphingolipid catabolic process |
| GO:00327 | 0.003409 | 3.94 | 4.191743 | 10 | 19 | positive regulation of interleukin-1 production |
| GO:00714 | 0.003417 | 1.53 | 47.6535 | 65 | 216 | cellular response to organic cyclic compound |
| GO:00018 | 0.003554 | 1.51 | 50.30091 | 68 | 228 | positive regulation of cytokine production |
| GO:00343 | 0.003564 | 6.20 | 2.426798 | 7 | 11 | low-density lipoprotein particle clearance |
| GO:00443 | 0.003564 | 6.20 | 2.426798 | 7 | 11 | wound healing, spreading of cells |
| GO:00465 | 0.003564 | 6.20 | 2.426798 | 7 | 11 | ceramide catabolic process |
| GO:00519 | 0.003564 | 6.20 | 2.426798 | 7 | 11 | positive regulation of nervous system development |
| GO:00519 | 0.003564 | 6.20 | 2.426798 | 7 | 11 | positive regulation of synapse assembly |
| GO:00905 | 0.003564 | 6.20 | 2.426798 | 7 | 11 | epiboly involved in wound healing |
| GO:00099 | 0.00358 | 1.18 | 411.8939 | 457 | 1867 | regulation of signal transduction |
| GO:00508 | 0.003591 | 1.69 | 30.88652 | 45 | 140 | regulation of synaptic transmission |
| GO:00355 | 0.003608 | 3.54 | 4.853597 | 11 | 22 | purinergic receptor signaling pathway |
| GO:00362 | 0.003608 | 3.54 | 4.853597 | 11 | 22 | granulocyte activation |
| GO:00518 | 0.003612 | 1.99 | 17.20821 | 28 | 78 | membrane depolarization |
| GO:00507 | 0.003634 | 3.07 | 6.177305 | 13 | 28 | positive regulation of phagocytosis |
| GO:00514 | 0.003634 | 3.07 | 6.177305 | 13 | 28 | regulation of intracellular pH |
| GO:19021 | 0.003656 | 1.59 | 39.27001 | 55 | 178 | regulation of leukocyte differentiation |
| GO:00511 | 0.00366 | 1.20 | 316.5869 | 357 | 1435 | regulation of phosphorus metabolic process |
| GO:00000 | 0.00367 | 3.27 | 5.515451 | 12 | 25 | regulation of transcription involved in G1/S transition of mitotic cell cycle |
| GO:00062 | 0.00367 | 3.27 | 5.515451 | 12 | 25 | pyrimidine nucleobase metabolic process |
| GO:00157 | 0.003764 | 1.48 | 55.59574 | 74 | 252 | organic anion transport |
| GO:00508 | 0.00384 | 1.33 | 109.8678 | 135 | 498 | neurological system process |
| GO:00026 | 0.004168 | 1.58 | 39.49063 | 55 | 179 | negative regulation of immune system process |
| GO:00400 | 0.004328 | 1.51 | 48.09473 | 65 | 218 | positive regulation of locomotion |
| GO:00510 | 0.004352 | 1.23 | 223.9273 | 258 | 1015 | regulation of transport |
| GO:00158 | 0.004409 | 2.58 | 8.383485 | 16 | 38 | L-amino acid transport |
| GO:00352 | 0.004409 | 2.58 | 8.383485 | 16 | 38 | synaptic transmission, glutamatergic |
| GO:00028 | 0.004461 | 1.83 | 21.39995 | 33 | 97 | regulation of adaptive immune response |
| GO:00326 | 0.0045 | 2.08 | 14.34017 | 24 | 65 | interleukin-6 production |
| GO:00019 | 0.004552 | 1.37 | 86.92351 | 109 | 394 | vasculature development |
| GO:00467 | 0.004577 | 2.32 | 10.58967 | 19 | 48 | acid secretion |
| GO:00028 | 0.004578 | 4.72 | 3.088652 | 8 | 14 | regulation of inflammatory response to antigenic stimulus |
| GO:00355 | 0.004578 | 4.72 | 3.088652 | 8 | 14 | purinergic nucleotide receptor signaling pathway |
| GO:00430 | 0.004578 | 4.72 | 3.088652 | 8 | 14 | myeloid dendritic cell differentiation |
| GO:00301 | 0.004636 | 1.37 | 83.39362 | 105 | 378 | regulation of proteolysis |
| GO:00095 | 0.004665 | 1.87 | 19.85562 | 31 | 90 | detection of external stimulus |
| GO:19016 | 0.004689 | 1.31 | 120.4574 | 146 | 546 | response to nitrogen compound |
| GO:00068 | 0.004722 | 2.66 | 7.721631 | 15 | 35 | superoxide metabolic process |
| GO:00508 | 0.004722 | 2.66 | 7.721631 | 15 | 35 | neuromuscular process controlling balance |
| GO:00515 | 0.004722 | 2.66 | 7.721631 | 15 | 35 | regulation of neurotransmitter transport |
| GO:00902 | 0.004737 | 1.75 | 24.70922 | 37 | 112 | regulation of muscle system process |
| GO:00105 | 0.00481 | 1.28 | 142.5193 | 170 | 646 | positive regulation of phosphorus metabolic process |
| GO:00459 | 0.00481 | 1.28 | 142.5193 | 170 | 646 | positive regulation of phosphate metabolic process |
| GO:00192 | 0.004828 | 1.41 | 69.27406 | 89 | 314 | cytokine-mediated signaling pathway |
| GO:00508 | 0.004857 | 1.50 | 48.31535 | 65 | 219 | regulation of T cell activation |
| GO:00301 | 0.00487 | 2.19 | 12.13399 | 21 | 55 | regulation of blood coagulation |
| GO:19000 | 0.00487 | 2.19 | 12.13399 | 21 | 55 | regulation of hemostasis |
| GO:00718 | 0.004967 | 1.96 | 16.76697 | 27 | 76 | leukocyte apoptotic process |
| GO:00022 | 0.005017 | 2.76 | 7.059777 | 14 | 32 | CD4-positive, alpha-beta T cell differentiation involved in immune response |
| GO:00420 | 0.005017 | 2.76 | 7.059777 | 14 | 32 | T-helper cell differentiation |
| GO:00507 | 0.005017 | 2.76 | 7.059777 | 14 | 32 | negative regulation of protein secretion |
| GO:00027 | 0.005036 | 1.78 | 23.16489 | 35 | 105 | regulation of leukocyte mediated immunity |
| GO:00015 | 0.005075 | 1.37 | 82.73176 | 104 | 375 | blood vessel development |
| GO:00433 | 0.005165 | 7.08 | 1.985562 | 6 | 9 | neutrophil degranulation |
| GO:00454 | 0.005165 | 7.08 | 1.985562 | 6 | 9 | regulation of interleukin-8 biosynthetic process |
| GO:00516 | 0.005165 | 7.08 | 1.985562 | 6 | 9 | establishment of mitochondrion localization |
| GO:00860 | 0.005165 | 7.08 | 1.985562 | 6 | 9 | membrane repolarization during action potential |
| GO:00860 | 0.005165 | 7.08 | 1.985562 | 6 | 9 | membrane repolarization during cardiac muscle cell action potential |
| GO:20000 | 0.005165 | 7.08 | 1.985562 | 6 | 9 | regulation of Wnt signaling pathway, planar cell polarity pathway |
| GO:00422 | 0.005188 | 3.98 | 3.750507 | 9 | 17 | response to cocaine |
| GO:00610 | 0.005188 | 3.98 | 3.750507 | 9 | 17 | myeloid leukocyte cytokine production |
| GO:00436 | 0.005222 | 1.66 | 29.78343 | 43 | 135 | cellular amide metabolic process |
| GO:00343 | 0.00527 | 1.61 | 33.97518 | 48 | 154 | cell junction assembly |
| GO:00326 | 0.005274 | 2.88 | 6.397923 | 13 | 29 | regulation of interleukin-1 beta production |
| GO:00072 | 0.005465 | 3.04 | 5.736069 | 12 | 26 | phospholipase C-activating G-protein coupled receptor signaling pathway |
| GO:00432 | 0.005465 | 3.04 | 5.736069 | 12 | 26 | apoptotic cell clearance |
| GO:00027 | 0.005483 | 2.41 | 9.265957 | 17 | 42 | regulation of cytokine production involved in immune response |
| GO:00017 | 0.005483 | 3.54 | 4.412361 | 10 | 20 | myeloid dendritic cell activation |
| GO:00024 | 0.005483 | 3.54 | 4.412361 | 10 | 20 | neutrophil mediated immunity |
| GO:00105 | 0.005483 | 3.54 | 4.412361 | 10 | 20 | regulation of platelet activation |
| GO:00158 | 0.005483 | 3.54 | 4.412361 | 10 | 20 | neutral amino acid transport |
| GO:00464 | 0.005483 | 3.54 | 4.412361 | 10 | 20 | membrane lipid catabolic process |
| GO:00507 | 0.005483 | 3.54 | 4.412361 | 10 | 20 | interleukin-1 secretion |

| | | | | | | |
|---|---|---|---|---|---|---|
| GO:00454 | 0.005552 | 3.25 | 5.074215 | 11 | 23 | positive regulation of nitric oxide biosynthetic process |
| GO:00508 | 0.005576 | 2.11 | 13.01646 | 22 | 59 | regulation of coagulation |
| GO:20001 | 0.005576 | 2.11 | 13.01646 | 22 | 59 | regulation of leukocyte apoptotic process |
| GO:00095 | 0.00561 | 1.84 | 20.07624 | 31 | 91 | detection of abiotic stimulus |
| GO:00330 | 0.00562 | 2.03 | 14.56079 | 24 | 66 | T cell differentiation in thymus |
| GO:00015 | 0.005832 | 1.43 | 60.89058 | 79 | 276 | angiogenesis |
| GO:00508 | 0.005957 | 2.25 | 10.81028 | 19 | 49 | positive regulation of B cell activation |
| GO:00705 | 0.005957 | 1.75 | 23.38551 | 35 | 106 | calcium ion transmembrane transport |
| GO:00326 | 0.00597 | 2.47 | 8.604103 | 16 | 39 | interleukin-12 production |
| GO:00507 | 0.006073 | 1.92 | 16.98759 | 27 | 77 | positive regulation of protein secretion |
| GO:00423 | 0.006112 | 1.28 | 128.6203 | 154 | 583 | positive regulation of phosphorylation |
| GO:00342 | 0.006527 | 1.30 | 116.9276 | 141 | 530 | ion transmembrane transport |
| GO:00068 | 0.006534 | 1.13 | 685.019 | 735 | 3105 | transport |
| GO:00026 | 0.006572 | 2.28 | 10.14843 | 18 | 46 | regulation of leukocyte chemotaxis |
| GO:00069 | 0.006572 | 2.28 | 10.14843 | 18 | 46 | cellular defense response |
| GO:00723 | 0.00668 | 1.28 | 132.5914 | 158 | 601 | cardiovascular system development |
| GO:00723 | 0.00668 | 1.28 | 132.5914 | 158 | 601 | circulatory system development |
| GO:00425 | 0.006725 | 1.22 | 209.8078 | 241 | 951 | homeostatic process |
| GO:00462 | 0.006895 | 2.15 | 11.69276 | 20 | 53 | nitric oxide metabolic process |
| GO:00027 | 0.006932 | 4.96 | 2.647416 | 7 | 12 | positive regulation of B cell mediated immunity |
| GO:00028 | 0.006932 | 4.96 | 2.647416 | 7 | 12 | positive regulation of immunoglobulin mediated immune response |
| GO:00312 | 0.006932 | 4.96 | 2.647416 | 7 | 12 | retinal ganglion cell axon guidance |
| GO:00517 | 0.006932 | 4.96 | 2.647416 | 7 | 12 | nitric-oxide synthase biosynthetic process |
| GO:00517 | 0.006932 | 4.96 | 2.647416 | 7 | 12 | regulation of nitric-oxide synthase biosynthetic process |
| GO:00905 | 0.006932 | 4.96 | 2.647416 | 7 | 12 | epiboly |
| GO:00970 | 0.006972 | 2.61 | 7.280395 | 14 | 33 | regulation of plasma lipoprotein particle levels |
| GO:00015 | 0.007112 | 1.78 | 21.17933 | 32 | 96 | regulation of neurotransmitter levels |
| GO:00228 | 0.007112 | 1.78 | 21.17933 | 32 | 96 | regulation of transmembrane transporter activity |
| GO:00326 | 0.007304 | 8.85 | 1.544326 | 5 | 7 | granulocyte macrophage colony-stimulating factor production |
| GO:00326 | 0.007304 | 8.85 | 1.544326 | 5 | 7 | regulation of granulocyte macrophage colony-stimulating factor production |
| GO:00329 | 0.007304 | 8.85 | 1.544326 | 5 | 7 | inositol trisphosphate metabolic process |
| GO:00330 | 0.007304 | 8.85 | 1.544326 | 5 | 7 | positive regulation of mast cell activation involved in immune response |
| GO:00433 | 0.007304 | 8.85 | 1.544326 | 5 | 7 | positive regulation of mast cell degranulation |
| GO:00444 | 0.007304 | 8.85 | 1.544326 | 5 | 7 | adhesion of symbiont to host |
| GO:00487 | 0.007304 | 8.85 | 1.544326 | 5 | 7 | embryonic viscerocranium morphogenesis |
| GO:19024 | 0.007304 | 8.85 | 1.544326 | 5 | 7 | L-alpha-amino acid transmembrane transport |
| GO:00333 | 0.007448 | 1.28 | 123.7667 | 148 | 561 | protein localization to organelle |
| GO:00306 | 0.007455 | 2.71 | 6.618541 | 13 | 30 | regulation of cellular pH |
| GO:00068 | 0.007499 | 1.75 | 22.0618 | 33 | 100 | neurotransmitter transport |
| GO:00027 | 0.007634 | 1.38 | 70.37715 | 89 | 319 | immune response-activating signal transduction |
| GO:00512 | 0.007647 | 1.13 | 701.786 | 751 | 3181 | establishment of localization |
| GO:00181 | 0.007675 | 1.45 | 51.84524 | 68 | 235 | peptidyl-tyrosine phosphorylation |
| GO:00326 | 0.007792 | 1.99 | 14.11955 | 23 | 64 | regulation of tumor necrosis factor production |
| GO:00024 | 0.007891 | 2.83 | 5.956687 | 12 | 27 | inflammatory response to antigenic stimulus |
| GO:00455 | 0.007927 | 1.84 | 18.09068 | 28 | 82 | regulation of T cell differentiation |
| GO:00015 | 0.007929 | 4.05 | 3.309271 | 8 | 15 | response to protozoan |
| GO:00422 | 0.007929 | 4.05 | 3.309271 | 8 | 15 | ribosomal large subunit biogenesis |
| GO:19010 | 0.007929 | 4.05 | 3.309271 | 8 | 15 | regulation of response to reactive oxygen species |
| GO:00068 | 0.007948 | 2.36 | 8.824721 | 16 | 40 | nitric oxide biosynthetic process |
| GO:00300 | 0.007948 | 2.36 | 8.824721 | 16 | 40 | cellular monovalent inorganic cation homeostasis |
| GO:00324 | 0.007984 | 1.78 | 20.51748 | 31 | 93 | regulation of ion transmembrane transporter activity |
| GO:00350 | 0.007999 | 1.56 | 35.5195 | 49 | 161 | regulation of Rho protein signal transduction |
| GO:00451 | 0.008235 | 3.00 | 5.294833 | 11 | 24 | regulation of bone resorption |
| GO:00105 | 0.008367 | 3.54 | 3.971125 | 9 | 18 | regulation of vascular endothelial growth factor production |
| GO:00327 | 0.008367 | 3.54 | 3.971125 | 9 | 18 | positive regulation of interleukin-1 beta production |
| GO:00423 | 0.008367 | 3.54 | 3.971125 | 9 | 18 | positive regulation of NF-kappaB import into nucleus |
| GO:00456 | 0.008367 | 3.54 | 3.971125 | 9 | 18 | regulation of T-helper cell differentiation |
| GO:00507 | 0.008367 | 3.54 | 3.971125 | 9 | 18 | interleukin-1 beta secretion |
| GO:00985 | 0.008367 | 3.54 | 3.971125 | 9 | 18 | detection of external biotic stimulus |
| GO:00434 | 0.008413 | 1.46 | 49.41844 | 65 | 224 | regulation of MAP kinase activity |
| GO:00025 | 0.008424 | 3.22 | 4.632979 | 10 | 21 | production of molecular mediator involved in inflammatory response |
| GO:00514 | 0.008424 | 3.22 | 4.632979 | 10 | 21 | intracellular pH reduction |
| GO:00516 | 0.008424 | 3.22 | 4.632979 | 10 | 21 | mitochondrion localization |
| GO:00070 | 0.008497 | 2.20 | 10.36905 | 18 | 47 | vacuolar transport |
| GO:00717 | 0.008555 | 1.94 | 15.00203 | 24 | 68 | tumor necrosis factor superfamily cytokine production |
| GO:00197 | 0.008783 | 1.30 | 103.2492 | 125 | 468 | cellular homeostasis |
| GO:00457 | 0.008831 | 1.72 | 22.28242 | 33 | 101 | positive regulation of cell cycle |
| GO:00026 | 0.008901 | 1.84 | 17.42882 | 27 | 79 | regulation of leukocyte migration |
| GO:00551 | 0.008901 | 1.84 | 17.42882 | 27 | 79 | digestive system development |
| GO:00423 | 0.00896 | 1.21 | 201.6449 | 231 | 914 | regulation of phosphorylation |
| GO:00719 | 0.009215 | 1.46 | 46.99164 | 62 | 213 | positive regulation of protein serine/threonine kinase activity |
| GO:00182 | 0.00942 | 1.43 | 52.28647 | 68 | 237 | peptidyl-tyrosine modification |
| GO:00450 | 0.00942 | 1.43 | 52.28647 | 68 | 237 | regulation of innate immune response |
| GO:00022 | 0.009489 | 2.48 | 7.501013 | 14 | 34 | alpha-beta T cell activation involved in immune response |
| GO:00022 | 0.009489 | 2.48 | 7.501013 | 14 | 34 | alpha-beta T cell differentiation involved in immune response |
| GO:00510 | 0.009547 | 1.33 | 87.14412 | 107 | 395 | regulation of small GTPase mediated signal transduction |
| GO:00442 | 0.009587 | 1.35 | 76.33384 | 95 | 346 | small molecule biosynthetic process |
| GO:00326 | 0.009594 | 1.94 | 14.34017 | 23 | 65 | tumor necrosis factor production |
| GO:00095 | 0.009731 | 2.11 | 11.25152 | 19 | 51 | detection of visible light |
| GO:00022 | 0.009739 | 14.15 | 1.10309 | 4 | 5 | T cell activation via T cell receptor contact with antigen bound to MHC molecule on antigen presenting cell |
| GO:00023 | 0.009739 | 14.15 | 1.10309 | 4 | 5 | MHC protein complex assembly |
| GO:00025 | 0.009739 | 14.15 | 1.10309 | 4 | 5 | peptide antigen assembly with MHC protein complex |
| GO:00108 | 0.009739 | 14.15 | 1.10309 | 4 | 5 | positive regulation of cholesterol storage |
| GO:00140 | 0.009739 | 14.15 | 1.10309 | 4 | 5 | regulation of neuron maturation |
| GO:00148 | 0.009739 | 14.15 | 1.10309 | 4 | 5 | tonic smooth muscle contraction |
| GO:00190 | 0.009739 | 14.15 | 1.10309 | 4 | 5 | virion attachment to host cell |
| GO:00324 | 0.009739 | 14.15 | 1.10309 | 4 | 5 | regulation of sodium:proton antiporter activity |
| GO:00326 | 0.009739 | 14.15 | 1.10309 | 4 | 5 | regulation of interleukin-18 production |
| GO:00446 | 0.009739 | 14.15 | 1.10309 | 4 | 5 | adhesion of symbiont to host cell |
| GO:00468 | 0.009739 | 14.15 | 1.10309 | 4 | 5 | quinolinate metabolic process |
| GO:00608 | 0.009739 | 14.15 | 1.10309 | 4 | 5 | establishment of blood-brain barrier |
| GO:19025 | 0.009739 | 14.15 | 1.10309 | 4 | 5 | regulation of neutrophil activation |
| GO:20012 | 0.009739 | 14.15 | 1.10309 | 4 | 5 | regulation of store-operated calcium entry |
| GO:00550 | 0.009762 | 2.01 | 12.79585 | 21 | 58 | monovalent inorganic cation homeostasis |
| GO:00082 | 0.009893 | 1.30 | 99.0575 | 120 | 449 | negative regulation of cell proliferation |
| GO:00069 | 0.009999 | 1.85 | 16.76697 | 26 | 76 | activation of cysteine-type endopeptidase activity involved in apoptotic process |
