## Supplemental Table 3 for "RNA-seq of Human T-Cells After Hematopoietic Stem Cell Transplantation Identifies Linc00402 as a Novel Regulator of T-Cell Alloimmunity"

| GOBPID | Pvalue | OddsRatio | ExpCount | Count | Size | Term |
|---|---|---|---|---|---|---|
| GO:00064 | 4.47E-26 | 9.81 | 20.1667 | 68 | 94 | translational termination |
| GO:00066 | 2.76E-19 | 6.08 | 22.5266 | 65 | 105 | SRP-dependent cotranslational protein targeting to membrane |
| GO:00066 | 1.14E-18 | 5.79 | 22.9557 | 65 | 107 | cotranslational protein targeting to membrane |
| GO:00450 | 2.28E-18 | 5.65 | 23.1702 | 65 | 108 | protein targeting to ER |
| GO:00725 | 4.47E-18 | 5.52 | 23.3848 | 65 | 109 | establishment of protein localization to endoplasmic reticulum |
| GO:00064 | 9.73E-18 | 5.09 | 25.3156 | 68 | 118 | translational elongation |
| GO:00001 | 1.44E-17 | 5.11 | 24.8865 | 67 | 116 | nuclear-transcribed mRNA catabolic process, nonsense-mediated decay |
| GO:00436 | 4.71E-17 | 3.89 | 35.3989 | 84 | 165 | cellular protein complex disassembly |
| GO:00432 | 9.17E-15 | 3.30 | 39.9043 | 87 | 186 | protein complex disassembly |
| GO:00709 | 2.75E-14 | 4.11 | 26.6028 | 65 | 124 | protein localization to endoplasmic reticulum |
| GO:00447 | 4.56E-14 | 1.41 | 872.959 | 1033 | 4069 | single-multicellular organism process |
| GO:00329 | 1.26E-13 | 3.05 | 42.0496 | 88 | 196 | macromolecular complex disassembly |
| GO:00224 | 1.30E-13 | 2.34 | 77.4486 | 138 | 361 | cellular component disassembly |
| GO:00190 | 3.81E-13 | 3.37 | 33.4681 | 74 | 156 | viral transcription |
| GO:00440 | 5.09E-13 | 3.15 | 37.5443 | 80 | 175 | multi-organism metabolic process |
| GO:00190 | 5.48E-13 | 3.24 | 35.6135 | 77 | 166 | viral gene expression |
| GO:00325 | 7.36E-13 | 1.39 | 902.995 | 1056 | 4209 | multicellular organismal process |
| GO:00066 | 8.07E-13 | 3.20 | 35.828 | 77 | 167 | protein targeting to membrane |
| GO:00064 | 8.37E-12 | 3.07 | 35.1844 | 74 | 164 | translational initiation |
| GO:00069 | 9.86E-12 | 2.10 | 87.5319 | 146 | 408 | inflammatory response |
| GO:00096 | 1.54E-11 | 1.56 | 291.988 | 390 | 1361 | response to external stimulus |
| GO:00508 | 6.97E-11 | 1.33 | 1158.51 | 1302 | 5400 | response to stimulus |
| GO:00512 | 7.18E-11 | 1.51 | 326.099 | 425 | 1520 | regulation of multicellular organismal process |
| GO:00325 | 1.84E-10 | 1.35 | 781.995 | 913 | 3645 | developmental process |
| GO:00096 | 4.72E-10 | 1.70 | 154.468 | 223 | 720 | response to wounding |
| GO:00650 | 6.79E-10 | 1.33 | 1620.41 | 1750 | 7553 | biological regulation |
| GO:00447 | 6.86E-10 | 1.34 | 772.77 | 899 | 3602 | single-organism developmental process |
| GO:00022 | 9.22E-10 | 3.34 | 24.0284 | 53 | 112 | myeloid leukocyte activation |
| GO:00072 | 1.16E-09 | 1.35 | 665.715 | 785 | 3103 | multicellular organismal development |
| GO:00009 | 1.62E-09 | 2.60 | 38.617 | 74 | 180 | nuclear-transcribed mRNA catabolic process |
| GO:00400 | 1.72E-09 | 1.55 | 223.335 | 301 | 1041 | locomotion |
| GO:00488 | 4.45E-09 | 1.33 | 689.957 | 806 | 3216 | anatomical structure development |
| GO:00901 | 5.80E-09 | 2.13 | 60.2855 | 102 | 281 | establishment of protein localization to membrane |
| GO:00164 | 6.79E-09 | 1.63 | 158.759 | 223 | 740 | cell migration |
| GO:00507 | 8.01E-09 | 1.30 | 1547.04 | 1670 | 7211 | regulation of biological process |
| GO:00487 | 8.45E-09 | 1.34 | 576.466 | 684 | 2687 | system development |
| GO:00082 | 1.07E-08 | 1.47 | 277.399 | 358 | 1293 | cell proliferation |
| GO:00488 | 1.17E-08 | 1.60 | 171.631 | 237 | 800 | cell motility |
| GO:00516 | 1.17E-08 | 1.60 | 171.631 | 237 | 800 | localization of cell |
| GO:00064 | 1.38E-08 | 2.41 | 40.977 | 75 | 191 | mRNA catabolic process |
| GO:00650 | 1.75E-08 | 1.36 | 476.706 | 575 | 2222 | regulation of biological quality |
| GO:19025 | 1.78E-08 | 1.88 | 84.9574 | 132 | 396 | single-organism localization |
| GO:19025 | 1.78E-08 | 1.88 | 84.9574 | 132 | 396 | single-organism cellular localization |
| GO:00726 | 2.45E-08 | 1.94 | 74.2305 | 118 | 346 | protein localization to membrane |
| GO:00422 | 3.33E-08 | 1.34 | 515.966 | 615 | 2405 | response to chemical |
| GO:00017 | 7.67E-08 | 1.59 | 153.61 | 212 | 716 | cell activation |
| GO:00069 | 1.04E-07 | 1.45 | 252.298 | 324 | 1176 | cellular component movement |
| GO:00453 | 1.16E-07 | 1.68 | 114.564 | 165 | 534 | leukocyte activation |
| GO:00230 | 1.29E-07 | 1.27 | 839.491 | 949 | 3913 | signaling |
| GO:00447 | 1.29E-07 | 1.27 | 839.491 | 949 | 3913 | single organism signaling |
| GO:00064 | 1.49E-07 | 2.16 | 46.7695 | 80 | 218 | RNA catabolic process |
| GO:00022 | 1.62E-07 | 2.28 | 40.1188 | 71 | 187 | adaptive immune response |
| GO:00421 | 1.64E-07 | 1.49 | 208.103 | 273 | 970 | regulation of cell proliferation |
| GO:00069 | 1.77E-07 | 1.46 | 231.273 | 299 | 1078 | defense response |
| GO:00071 | 1.86E-07 | 1.27 | 852.578 | 961 | 3974 | cell communication |
| GO:00507 | 2.73E-07 | 1.26 | 1477.53 | 1588 | 6887 | regulation of cellular process |
| GO:00071 | 2.78E-07 | 1.57 | 148.246 | 203 | 691 | cell adhesion |
| GO:00517 | 2.85E-07 | 1.26 | 948.906 | 1058 | 4423 | cellular response to stimulus |
| GO:00507 | 3.32E-07 | 4.12 | 11.7996 | 29 | 55 | positive regulation of inflammatory response |
| GO:00024 | 3.43E-07 | 2.33 | 35.6135 | 64 | 166 | adaptive immune response based on somatic recombination of immune receptors built from immunoglobulin superfamily domains |
| GO:00226 | 3.50E-07 | 1.56 | 148.676 | 203 | 693 | biological adhesion |
| GO:00488 | 4.07E-07 | 1.30 | 528.839 | 620 | 2465 | cellular developmental process |
| GO:00023 | 4.47E-07 | 1.35 | 388.101 | 469 | 1809 | immune system process |
| GO:00447 | 5.23E-07 | 1.28 | 1821.22 | 1919 | 8489 | single-organism cellular process |
| GO:00301 | 5.26E-07 | 2.05 | 49.5585 | 82 | 231 | extracellular matrix organization |
| GO:00430 | 5.26E-07 | 2.05 | 49.5585 | 82 | 231 | extracellular structure organization |
| GO:19030 | 6.69E-07 | 2.00 | 52.133 | 85 | 243 | regulation of response to wounding |
| GO:00096 | 6.93E-07 | 1.93 | 59.2128 | 94 | 276 | response to bacterium |
| GO:00071 | 6.97E-07 | 1.26 | 772.555 | 873 | 3601 | signal transduction |
| GO:00603 | 7.07E-07 | 2.40 | 31.1082 | 57 | 145 | cell chemotaxis |
| GO:00069 | 7.59E-07 | 1.43 | 229.342 | 293 | 1069 | immune response |
| GO:00507 | 9.31E-07 | 2.21 | 37.9734 | 66 | 177 | regulation of inflammatory response |
| GO:00485 | 1.04E-06 | 1.32 | 418.566 | 499 | 1951 | organ development |
| GO:00301 | 1.42E-06 | 1.30 | 491.723 | 576 | 2292 | cell differentiation |
| GO:20000 | 1.43E-06 | 1.45 | 201.452 | 260 | 939 | regulation of multicellular organismal development |
| GO:19030 | 1.50E-06 | 3.38 | 14.3741 | 32 | 67 | positive regulation of response to wounding |
| GO:00022 | 1.63E-06 | 2.38 | 29.6064 | 54 | 138 | cell activation involved in immune response |
| GO:00023 | 1.63E-06 | 2.38 | 29.6064 | 54 | 138 | leukocyte activation involved in immune response |
| GO:00508 | 1.69E-06 | 1.78 | 73.1578 | 110 | 341 | regulation of cell activation |
| GO:00903 | 1.97E-06 | 4.77 | 8.36702 | 22 | 39 | phagosome maturation |
| GO:00329 | 2.31E-06 | 2.13 | 38.8316 | 66 | 181 | mononuclear cell proliferation |
| GO:00485 | 2.45E-06 | 1.28 | 517.254 | 601 | 2411 | regulation of response to stimulus |
| GO:00069 | 2.52E-06 | 1.27 | 570.459 | 657 | 2659 | response to stress |
| GO:00708 | 2.64E-06 | 1.33 | 359.353 | 432 | 1675 | cellular response to chemical stimulus |
| GO:00096 | 2.84E-06 | 1.33 | 350.342 | 422 | 1633 | anatomical structure morphogenesis |
| GO:00190 | 3.50E-06 | 1.82 | 62.0018 | 95 | 289 | viral life cycle |
| GO:00507 | 3.94E-06 | 1.37 | 273.323 | 337 | 1274 | regulation of developmental process |
| GO:00026 | 4.00E-06 | 1.77 | 68.6525 | 103 | 320 | regulation of leukocyte activation |
| GO:00466 | 4.01E-06 | 2.10 | 38.617 | 65 | 180 | lymphocyte proliferation |
| GO:00718 | 4.16E-06 | 1.38 | 255.301 | 317 | 1190 | protein complex subunit organization |
| GO:00305 | 4.25E-06 | 2.54 | 23.1702 | 44 | 108 | leukocyte chemotaxis |
| GO:00716 | 4.94E-06 | 3.69 | 11.156 | 26 | 52 | granulocyte chemotaxis |
| GO:00321 | 6.22E-06 | 2.30 | 28.5337 | 51 | 133 | positive regulation of response to external stimulus |
| GO:00339 | 6.62E-06 | 1.60 | 101.048 | 141 | 471 | response to lipid |
| GO:00485 | 6.71E-06 | 1.37 | 252.083 | 312 | 1175 | positive regulation of response to stimulus |
| GO:00022 | 7.13E-06 | 3.85 | 10.0833 | 24 | 47 | myeloid cell activation involved in immune response |
| GO:00706 | 7.27E-06 | 2.02 | 40.7624 | 67 | 190 | leukocyte proliferation |
| GO:00510 | 7.71E-06 | 1.53 | 123.574 | 167 | 576 | positive regulation of developmental process |
| GO:00975 | 7.83E-06 | 3.43 | 12.0142 | 27 | 56 | granulocyte migration |
| GO:00024 | 1.03E-05 | 1.97 | 42.6933 | 69 | 199 | leukocyte mediated immunity |
| GO:00069 | 1.04E-05 | 1.60 | 95.8989 | 134 | 447 | chemotaxis |
| GO:00423 | 1.04E-05 | 1.60 | 95.8989 | 134 | 447 | taxis |
| GO:00028 | 1.05E-05 | 7.36 | 4.50532 | 14 | 21 | regulation of myeloid leukocyte mediated immunity |
| GO:00330 | 1.05E-05 | 7.36 | 4.50532 | 14 | 21 | regulation of mast cell activation |
| GO:00975 | 1.11E-05 | 2.71 | 18.2358 | 36 | 85 | myeloid leukocyte migration |
| GO:00030 | 1.15E-05 | 1.42 | 186.863 | 238 | 871 | system process |
| GO:00517 | 1.26E-05 | 1.31 | 331.034 | 396 | 1543 | multi-organism process |
| GO:00427 | 1.32E-05 | 2.48 | 21.883 | 41 | 102 | defense response to bacterium |
| GO:00421 | 1.38E-05 | 4.33 | 7.93794 | 20 | 37 | macrophage activation |
| GO:00096 | 1.43E-05 | 2.41 | 23.3848 | 43 | 109 | response to toxic substance |
| GO:00448 | 1.46E-05 | 1.50 | 125.72 | 168 | 586 | single-organism membrane organization |
| GO:00511 | 1.47E-05 | 1.22 | 821.684 | 910 | 3830 | localization |
| GO:00197 | 1.52E-05 | 2.66 | 18.4504 | 36 | 86 | B cell mediated immunity |
| GO:00098 | 1.55E-05 | 1.36 | 235.349 | 291 | 1097 | tissue development |
| GO:00506 | 1.59E-05 | 2.53 | 20.5957 | 39 | 96 | cytokine secretion |
| GO:00160 | 1.64E-05 | 2.69 | 17.8067 | 35 | 83 | immunoglobulin mediated immune response |
| GO:19017 | 1.72E-05 | 1.40 | 189.652 | 240 | 884 | response to oxygen-containing compound |
| GO:00329 | 1.84E-05 | 2.16 | 30.25 | 52 | 141 | regulation of mononuclear cell proliferation |
| GO:00506 | 1.84E-05 | 2.16 | 30.25 | 52 | 141 | regulation of lymphocyte proliferation |
| GO:00321 | 1.85E-05 | 1.59 | 91.8227 | 128 | 428 | regulation of response to external stimulus |
| GO:00025 | 1.89E-05 | 2.58 | 19.3085 | 37 | 90 | antigen processing and presentation of peptide or polysaccharide antigen via MHC class II |
| GO:00447 | 1.94E-05 | 1.25 | 550.936 | 628 | 2568 | single-organism transport |
| GO:00706 | 2.05E-05 | 2.13 | 31.1082 | 53 | 145 | regulation of leukocyte proliferation |
| GO:00725 | 2.08E-05 | 1.59 | 91.1791 | 127 | 425 | establishment of protein localization to organelle |
| GO:00507 | 2.43E-05 | 2.67 | 17.3777 | 34 | 81 | regulation of cytokine secretion |
| GO:00022 | 2.72E-05 | 1.54 | 103.622 | 141 | 483 | immune effector process |
| GO:00420 | 2.82E-05 | 2.75 | 16.0904 | 32 | 75 | regulation of cytokine biosynthetic process |
| GO:00022 | 3.37E-05 | 1.93 | 40.1188 | 64 | 187 | response to molecule of bacterial origin |
| GO:00226 | 3.68E-05 | 1.50 | 112.848 | 151 | 526 | regulation of anatomical structure morphogenesis |
| GO:00024 | 3.71E-05 | 2.51 | 19.094 | 36 | 89 | antigen processing and presentation of peptide antigen via MHC class II |
| GO:00305 | 4.11E-05 | 3.52 | 9.65426 | 22 | 45 | neutrophil chemotaxis |
| GO:00469 | 4.36E-05 | 1.44 | 139.45 | 181 | 650 | secretion |
| GO:00508 | 4.52E-05 | 1.82 | 47.6277 | 73 | 222 | positive regulation of cell activation |
| GO:00343 | 4.77E-05 | 2.36 | 21.4539 | 39 | 100 | response to interferon-gamma |

| | | | | | | |
|---|---|---|---|---|---|---|
| GO:00420 | 4.79E-05 | 4.47 | 6.65071 | 17 | 31 | T-helper 1 type immune response |
| GO:00100 | 5.12E-05 | 1.27 | 379.734 | 443 | 1770 | response to organic substance |
| GO:00421 | 5.42E-05 | 2.48 | 18.6649 | 35 | 87 | cytokine metabolic process |
| GO:19030 | 6.19E-05 | 2.70 | 15.2323 | 30 | 71 | negative regulation of response to wounding |
| GO:00024 | 6.29E-05 | 3.38 | 9.86879 | 22 | 46 | myeloid leukocyte mediated immunity |
| GO:19902 | 6.29E-05 | 3.38 | 9.86879 | 22 | 46 | neutrophil migration |
| GO:00069 | 6.48E-05 | 2.42 | 19.523 | 36 | 91 | humoral immune response |
| GO:00723 | 6.58E-05 | 1.45 | 128.938 | 168 | 601 | cardiovascular system development |
| GO:00723 | 6.58E-05 | 1.45 | 128.938 | 168 | 601 | circulatory system development |
| GO:00433 | 6.97E-05 | 5.97 | 4.50532 | 13 | 21 | regulation of leukocyte degranulation |
| GO:19033 | 6.97E-05 | 5.97 | 4.50532 | 13 | 21 | regulation of regulated secretory pathway |
| GO:00082 | 7.78E-05 | 1.48 | 110.917 | 147 | 517 | positive regulation of cell proliferation |
| GO:00420 | 7.89E-05 | 2.46 | 18.2358 | 34 | 85 | cytokine biosynthetic process |
| GO:00420 | 8.55E-05 | 1.48 | 111.989 | 148 | 522 | wound healing |
| GO:00433 | 8.61E-05 | 9.19 | 3.00355 | 10 | 14 | regulation of mast cell degranulation |
| GO:00192 | 9.21E-05 | 1.29 | 305.289 | 361 | 1423 | regulation of phosphate metabolic process |
| GO:00512 | 9.35E-05 | 1.52 | 95.6844 | 129 | 446 | regulation of cellular component movement |
| GO:00071 | 9.41E-05 | 1.24 | 441.521 | 506 | 2058 | cell surface receptor signaling pathway |
| GO:00455 | 9.43E-05 | 1.53 | 90.5355 | 123 | 422 | positive regulation of cell differentiation |
| GO:19025 | 9.83E-05 | 1.31 | 251.869 | 303 | 1174 | regulation of intracellular signal transduction |
| GO:00610 | 9.94E-05 | 1.40 | 152.966 | 194 | 713 | membrane organization |
| GO:00198 | 0.0001 | 2.41 | 18.4504 | 34 | 86 | antigen processing and presentation of exogenous peptide antigen via MHC class II |
| GO:00801 | 0.0001 | 1.38 | 165.624 | 208 | 772 | regulation of response to stress |
| GO:00512 | 0.00011 | 1.53 | 90.75 | 123 | 423 | positive regulation of multicellular organismal process |
| GO:00026 | 0.00011 | 1.78 | 46.3404 | 70 | 216 | positive regulation of leukocyte activation |
| GO:00511 | 0.00011 | 1.28 | 307.863 | 363 | 1435 | regulation of phosphorus metabolic process |
| GO:00226 | 0.00011 | 2.57 | 15.6613 | 30 | 73 | extracellular matrix disassembly |
| GO:00163 | 0.00012 | 1.75 | 48.9149 | 73 | 228 | single organismal cell-cell adhesion |
| GO:00018 | 0.00012 | 1.54 | 84.9574 | 116 | 396 | regulation of cytokine production |
| GO:00147 | 0.00012 | 1.78 | 45.6968 | 69 | 213 | striated muscle tissue development |
| GO:00432 | 0.00012 | 1.48 | 106.626 | 141 | 497 | response to external biotic stimulus |
| GO:00517 | 0.00012 | 1.48 | 106.626 | 141 | 497 | response to other organism |
| GO:00070 | 0.00012 | 1.92 | 34.5408 | 55 | 161 | chromosome segregation |
| GO:00986 | 0.00013 | 1.68 | 56.4238 | 82 | 263 | single organism cell adhesion |
| GO:00068 | 0.00013 | 1.63 | 64.7908 | 92 | 302 | cellular ion homeostasis |
| GO:00324 | 0.00013 | 14.69 | 2.14539 | 8 | 10 | response to peptidoglycan |
| GO:00330 | 0.00013 | 14.69 | 2.14539 | 8 | 10 | positive regulation of mast cell activation |
| GO:00321 | 0.00014 | 2.08 | 26.1738 | 44 | 122 | negative regulation of response to external stimulus |
| GO:00026 | 0.00015 | 2.23 | 21.6684 | 38 | 101 | positive regulation of immune effector process |
| GO:00550 | 0.00015 | 1.56 | 78.5213 | 108 | 366 | cellular chemical homeostasis |
| GO:00446 | 0.00017 | 1.22 | 1999.5 | 2065 | 9320 | single-organism process |
| GO:00509 | 0.00017 | 1.71 | 51.0603 | 75 | 238 | leukocyte migration |
| GO:00324 | 0.00017 | 1.85 | 37.3298 | 58 | 174 | response to lipopolysaccharide |
| GO:00508 | 0.00018 | 1.46 | 108.342 | 142 | 505 | regulation of body fluid levels |
| GO:00400 | 0.00018 | 1.50 | 93.539 | 125 | 436 | regulation of locomotion |
| GO:00508 | 0.00018 | 2.43 | 16.734 | 31 | 78 | regulation of B cell activation |
| GO:00327 | 0.00019 | 4.24 | 6.00709 | 15 | 28 | positive regulation of interleukin-6 production |
| GO:00096 | 0.00019 | 1.45 | 111.989 | 146 | 522 | response to biotic stimulus |
| GO:00326 | 0.00019 | 2.92 | 11.156 | 23 | 52 | chemokine production |
| GO:00507 | 0.0002 | 2.06 | 25.7447 | 43 | 120 | regulation of protein secretion |
| GO:00450 | 0.0002 | 1.38 | 147.603 | 186 | 688 | innate immune response |
| GO:00308 | 0.0002 | 1.56 | 73.1578 | 101 | 341 | epithelial cell differentiation |
| GO:00488 | 0.00021 | 1.42 | 121.858 | 157 | 568 | chemical homeostasis |
| GO:00719 | 0.00021 | 1.56 | 74.8741 | 103 | 349 | regulation of protein serine/threonine kinase activity |
| GO:00303 | 0.00021 | 1.53 | 80.023 | 109 | 373 | regulation of cell migration |
| GO:00330 | 0.00021 | 7.35 | 3.21809 | 10 | 15 | regulation of mast cell activation involved in immune response |
| GO:00450 | 0.00021 | 3.09 | 9.86879 | 21 | 46 | regulated secretory pathway |
| GO:00300 | 0.00021 | 1.61 | 63.0745 | 89 | 294 | cellular cation homeostasis |
| GO:00326 | 0.00022 | 2.66 | 13.3014 | 26 | 62 | regulation of interleukin-6 production |
| GO:00106 | 0.00022 | 1.34 | 186.649 | 229 | 870 | positive regulation of cell communication |
| GO:00329 | 0.00022 | 1.42 | 123.789 | 159 | 577 | secretion by cell |
| GO:00432 | 0.00022 | 3.49 | 7.93794 | 18 | 37 | leukocyte degranulation |
| GO:20001 | 0.00023 | 1.51 | 85.3865 | 115 | 398 | regulation of cell motility |
| GO:00075 | 0.00023 | 1.72 | 47.4131 | 70 | 221 | muscle organ development |
| GO:00102 | 0.00024 | 1.45 | 108.986 | 142 | 508 | response to organonitrogen compound |
| GO:00469 | 0.00024 | 2.38 | 16.9486 | 31 | 79 | cellular transition metal ion homeostasis |
| GO:00106 | 0.00024 | 1.23 | 438.947 | 499 | 2046 | regulation of cell communication |
| GO:00015 | 0.00026 | 1.52 | 80.4521 | 109 | 375 | blood vessel development |
| GO:00140 | 0.00026 | 1.46 | 100.404 | 132 | 468 | response to organic cyclic compound |
| GO:00015 | 0.00026 | 1.62 | 59.2128 | 84 | 276 | angiogenesis |
| GO:00066 | 0.00026 | 1.46 | 99.5461 | 131 | 464 | protein targeting |
| GO:00713 | 0.00027 | 1.27 | 293.275 | 344 | 1367 | cellular response to organic substance |
| GO:00605 | 0.00027 | 1.71 | 47.6277 | 70 | 222 | muscle tissue development |
| GO:00421 | 0.00029 | 2.89 | 10.727 | 22 | 50 | positive regulation of cytokine biosynthetic process |
| GO:00425 | 0.00029 | 8.26 | 2.78901 | 9 | 13 | tumor necrosis factor biosynthetic process |
| GO:00425 | 0.00029 | 8.26 | 2.78901 | 9 | 13 | regulation of tumor necrosis factor biosynthetic process |
| GO:00018 | 0.0003 | 1.47 | 95.4699 | 126 | 445 | cytokine production |
| GO:00024 | 0.00033 | 1.85 | 34.1117 | 53 | 159 | lymphocyte mediated immunity |
| GO:00706 | 0.00033 | 2.19 | 20.1667 | 35 | 94 | positive regulation of leukocyte proliferation |
| GO:00313 | 0.00033 | 1.67 | 50.4167 | 73 | 235 | positive regulation of defense response |
| GO:00455 | 0.00034 | 3.47 | 7.50887 | 17 | 35 | mast cell activation |
| GO:00026 | 0.00035 | 1.40 | 122.287 | 156 | 570 | positive regulation of immune system process |
| GO:00019 | 0.00036 | 1.49 | 84.5284 | 113 | 394 | vasculature development |
| GO:00068 | 0.00038 | 1.61 | 58.1401 | 82 | 271 | cellular metal ion homeostasis |
| GO:00466 | 0.00039 | 1.45 | 99.5461 | 130 | 464 | lymphocyte activation |
| GO:00022 | 0.00039 | 9.79 | 2.35993 | 8 | 11 | neutrophil activation involved in immune response |
| GO:00230 | 0.00041 | 1.32 | 185.791 | 226 | 866 | positive regulation of signaling |
| GO:00026 | 0.00045 | 1.31 | 198.878 | 240 | 927 | regulation of immune system process |
| GO:00230 | 0.00045 | 1.21 | 437.874 | 495 | 2041 | regulation of signaling |
| GO:00466 | 0.00045 | 6.12 | 3.43262 | 10 | 16 | gamma-delta T cell activation |
| GO:00093 | 0.00046 | 1.81 | 34.5408 | 53 | 161 | protein secretion |
| GO:00442 | 0.00046 | 2.94 | 9.65426 | 20 | 45 | multicellular organismal catabolic process |
| GO:00329 | 0.00047 | 2.16 | 19.7376 | 34 | 92 | positive regulation of mononuclear cell proliferation |
| GO:00506 | 0.00047 | 2.16 | 19.7376 | 34 | 92 | positive regulation of lymphocyte proliferation |
| GO:00333 | 0.00048 | 1.40 | 120.356 | 153 | 561 | protein localization to organelle |
| GO:00717 | 0.00048 | 2.43 | 14.5887 | 27 | 68 | tumor necrosis factor superfamily cytokine production |
| GO:00020 | 0.0005 | 3.15 | 8.36702 | 18 | 39 | sprouting angiogenesis |
| GO:00604 | 0.00051 | 3.68 | 6.43617 | 15 | 30 | heart growth |
| GO:00439 | 0.00052 | 1.25 | 288.34 | 336 | 1344 | macromolecular complex subunit organization |
| GO:00713 | 0.00052 | 2.24 | 17.5922 | 31 | 82 | cellular response to interferon-gamma |
| GO:00099 | 0.00052 | 1.22 | 400.544 | 455 | 1867 | regulation of signal transduction |
| GO:00326 | 0.00053 | 2.46 | 13.945 | 26 | 65 | interleukin-6 production |
| GO:00507 | 0.00053 | 2.46 | 13.945 | 26 | 65 | negative regulation of inflammatory response |
| GO:00330 | 0.00055 | 22.02 | 1.50177 | 6 | 7 | positive regulation of mast cell activation involved in immune response |
| GO:00433 | 0.00055 | 22.02 | 1.50177 | 6 | 7 | positive regulation of mast cell degranulation |
| GO:00421 | 0.00057 | 1.51 | 72.5142 | 98 | 338 | T cell activation |
| GO:00485 | 0.00057 | 1.52 | 70.7979 | 96 | 330 | blood vessel morphogenesis |
| GO:00712 | 0.00058 | 2.07 | 21.4539 | 36 | 100 | cellular response to molecule of bacterial origin |
| GO:00327 | 0.00058 | 5.05 | 4.07624 | 11 | 19 | positive regulation of interleukin-1 production |
| GO:00352 | 0.00058 | 2.49 | 13.3014 | 25 | 62 | organ growth |
| GO:00016 | 0.00063 | 2.37 | 14.8032 | 27 | 69 | long-chain fatty acid metabolic process |
| GO:00440 | 0.00063 | 1.63 | 49.773 | 71 | 232 | regulation of system process |
| GO:00328 | 0.00064 | 1.25 | 294.777 | 342 | 1374 | regulation of localization |
| GO:00340 | 0.00066 | 1.42 | 103.408 | 133 | 482 | response to cytokine |
| GO:00507 | 0.00069 | 1.36 | 134.73 | 168 | 628 | regulation of immune response |
| GO:00305 | 0.00069 | 2.91 | 9.22518 | 19 | 43 | collagen catabolic process |
| GO:00326 | 0.0007 | 2.39 | 14.1596 | 26 | 66 | regulation of interferon-gamma production |
| GO:00326 | 0.00076 | 3.13 | 7.93794 | 17 | 37 | interleukin-1 production |
| GO:00985 | 0.00078 | 1.57 | 56.8528 | 79 | 265 | defense response to other organism |
| GO:00335 | 0.00078 | 3.68 | 6.00709 | 14 | 28 | transferrin transport |
| GO:00024 | 0.00078 | 3.27 | 7.29433 | 16 | 34 | humoral immune response mediated by circulating immunoglobulin |
| GO:00326 | 0.00078 | 3.27 | 7.29433 | 16 | 34 | interleukin-1 beta production |
| GO:00326 | 0.00079 | 3.45 | 6.65071 | 15 | 31 | regulation of interleukin-1 production |
| GO:00030 | 0.00083 | 2.10 | 19.523 | 33 | 91 | vascular process in circulatory system |
| GO:00454 | 0.00084 | 2.66 | 10.727 | 21 | 50 | endothelial cell differentiation |
| GO:00080 | 0.00086 | 1.58 | 54.4929 | 76 | 254 | blood circulation |
| GO:00302 | 0.00088 | 5.25 | 3.64716 | 10 | 17 | negative regulation of ossification |
| GO:00422 | 0.00088 | 5.25 | 3.64716 | 10 | 17 | response to cocaine |
| GO:00072 | 0.00089 | 1.42 | 95.2553 | 123 | 444 | small GTPase mediated signal transduction |
| GO:00604 | 0.00092 | 1.34 | 140.952 | 174 | 657 | epithelium development |
| GO:00421 | 0.00095 | 1.98 | 22.7411 | 37 | 106 | regulation of T cell proliferation |
| GO:00022 | 0.00095 | 7.34 | 2.57447 | 8 | 12 | macrophage activation involved in immune response |
| GO:00215 | 0.00095 | 7.34 | 2.57447 | 8 | 12 | cell migration in hindbrain |
| GO:19017 | 0.00095 | 7.34 | 2.57447 | 8 | 12 | positive regulation of signal transduction by p53 class mediator |
| GO:00068 | 0.00097 | 2.80 | 9.43972 | 19 | 44 | iron ion transport |

| | | | | | | |
|---|---|---|---|---|---|---|
| GO:00512 | 0.001 | 1.53 | 61.5727 | 84 | 287 | regulation of lymphocyte activation |
| GO:00326 | 0.00101 | 2.36 | 13.7305 | 25 | 64 | regulation of tumor necrosis factor production |
| GO:00323 | 0.00102 | 1.89 | 25.9592 | 41 | 121 | positive regulation of Rho GTPase activity |
| GO:00359 | 0.00104 | 2.88 | 8.7961 | 18 | 41 | skeletal muscle cell differentiation |
| GO:00071 | 0.00105 | 2.12 | 18.2358 | 31 | 85 | negative regulation of cell adhesion |
| GO:19017 | 0.00108 | 1.35 | 127.865 | 159 | 596 | cellular response to oxygen-containing compound |
| GO:00098 | 0.00109 | 1.38 | 112.633 | 142 | 525 | organ morphogenesis |
| GO:00508 | 0.0011 | 1.44 | 84.3138 | 110 | 393 | ion homeostasis |
| GO:00030 | 0.0011 | 1.56 | 54.922 | 76 | 256 | circulatory system process |
| GO:00198 | 0.00112 | 1.63 | 44.8387 | 64 | 209 | antigen processing and presentation |
| GO:00313 | 0.00113 | 1.42 | 90.5355 | 117 | 422 | regulation of defense response |
| GO:00100 | 0.00114 | 1.68 | 39.0461 | 57 | 182 | response to metal ion |
| GO:00422 | 0.00115 | 4.01 | 4.9344 | 12 | 23 | ribosomal small subunit biogenesis |
| GO:00603 | 0.00118 | 2.28 | 14.5887 | 26 | 68 | interferon-gamma-mediated signaling pathway |
| GO:00019 | 0.0012 | 3.67 | 5.57801 | 13 | 26 | blood vessel remodeling |
| GO:00018 | 0.00121 | 3.43 | 6.22163 | 14 | 29 | endothelial cell development |
| GO:00156 | 0.00121 | 3.43 | 6.22163 | 14 | 29 | ferric iron transport |
| GO:00326 | 0.00121 | 3.43 | 6.22163 | 14 | 29 | regulation of interleukin-1 beta production |
| GO:00327 | 0.00121 | 3.43 | 6.22163 | 14 | 29 | positive regulation of chemokine production |
| GO:00519 | 0.00121 | 3.43 | 6.22163 | 14 | 29 | regulation of chromosome segregation |
| GO:00725 | 0.00121 | 3.43 | 6.22163 | 14 | 29 | trivalent inorganic cation transport |
| GO:00099 | 0.00128 | 1.29 | 179.355 | 215 | 836 | positive regulation of signal transduction |
| GO:00025 | 0.0013 | 1.79 | 30.25 | 46 | 141 | myeloid leukocyte differentiation |
| GO:00326 | 0.00131 | 2.30 | 13.945 | 25 | 65 | tumor necrosis factor production |
| GO:00009 | 0.00132 | 1.29 | 177.638 | 213 | 828 | cell morphogenesis |
| GO:00985 | 0.00133 | 5.51 | 3.21809 | 9 | 15 | detection of other organism |
| GO:00320 | 0.00134 | 8.56 | 2.14539 | 7 | 10 | response to magnesium ion |
| GO:00433 | 0.00134 | 8.56 | 2.14539 | 7 | 10 | positive regulation of leukocyte degranulation |
| GO:19033 | 0.00134 | 8.56 | 2.14539 | 7 | 10 | positive regulation of regulated secretory pathway |
| GO:00326 | 0.00135 | 2.69 | 9.65426 | 19 | 45 | regulation of chemokine production |
| GO:00075 | 0.0014 | 1.92 | 23.1702 | 37 | 108 | female pregnancy |
| GO:00108 | 0.00143 | 1.63 | 43.5514 | 62 | 203 | lipid localization |
| GO:00508 | 0.00147 | 1.41 | 92.0372 | 118 | 429 | coagulation |
| GO:00160 | 0.00147 | 1.15 | 854.509 | 918 | 3983 | cellular component organization |
| GO:00457 | 0.00151 | 2.23 | 14.8032 | 26 | 69 | positive regulation of angiogenesis |
| GO:19016 | 0.00154 | 1.36 | 117.138 | 146 | 546 | response to nitrogen compound |
| GO:00487 | 0.00156 | 1.99 | 20.1667 | 33 | 94 | tissue remodeling |
| GO:00326 | 0.00157 | 2.84 | 8.36702 | 17 | 39 | interleukin-12 production |
| GO:00327 | 0.0016 | 4.59 | 3.8617 | 10 | 18 | positive regulation of interleukin-1 beta production |
| GO:00985 | 0.0016 | 4.59 | 3.8617 | 10 | 18 | detection of external biotic stimulus |
| GO:00224 | 0.00161 | 1.30 | 161.762 | 195 | 754 | reproductive process |
| GO:00550 | 0.00162 | 1.45 | 74.6596 | 98 | 348 | cation homeostasis |
| GO:19025 | 0.00166 | 1.35 | 117.353 | 146 | 547 | positive regulation of intracellular signal transduction |
| GO:00027 | 0.00167 | 2.94 | 7.7234 | 16 | 36 | regulation of B cell mediated immunity |
| GO:00192 | 0.00168 | 1.66 | 37.9734 | 55 | 177 | regulation of lipid metabolic process |
| GO:00713 | 0.00172 | 1.57 | 48.9149 | 68 | 228 | cellular response to lipid |
| GO:00063 | 0.00174 | 1.82 | 26.6028 | 41 | 124 | DNA packaging |
| GO:00510 | 0.00175 | 1.44 | 77.4486 | 101 | 361 | regulation of secretion |
| GO:00230 | 0.00176 | 1.37 | 104.051 | 131 | 485 | signal transduction by phosphorylation |
| GO:00345 | 0.00176 | 3.06 | 7.07979 | 15 | 33 | protein localization to chromosome |
| GO:00019 | 0.00177 | 4.04 | 4.50532 | 11 | 21 | negative regulation of cell-matrix adhesion |
| GO:00025 | 0.00177 | 4.04 | 4.50532 | 11 | 21 | production of molecular mediator involved in inflammatory response |
| GO:00095 | 0.00177 | 4.04 | 4.50532 | 11 | 21 | detection of biotic stimulus |
| GO:00516 | 0.00177 | 4.04 | 4.50532 | 11 | 21 | mitochondrion localization |
| GO:00313 | 0.00177 | 2.00 | 19.523 | 32 | 91 | negative regulation of defense response |
| GO:00075 | 0.00178 | 1.40 | 91.6082 | 117 | 427 | blood coagulation |
| GO:00323 | 0.0018 | 1.75 | 30.6791 | 46 | 143 | regulation of Rho GTPase activity |
| GO:00000 | 0.00181 | 11.01 | 1.71631 | 6 | 8 | ribosomal small subunit assembly |
| GO:00199 | 0.00181 | 11.01 | 1.71631 | 6 | 8 | cGMP-mediated signaling |
| GO:00353 | 0.00181 | 11.01 | 1.71631 | 6 | 8 | wound healing, spreading of epidermal cells |
| GO:00380 | 0.00181 | 11.01 | 1.71631 | 6 | 8 | peptidyl-tyrosine autophosphorylation |
| GO:00488 | 0.00181 | 11.01 | 1.71631 | 6 | 8 | chemical homeostasis within a tissue |
| GO:00031 | 0.00182 | 2.38 | 12.0142 | 22 | 56 | endothelium development |
| GO:00015 | 0.00183 | 3.22 | 6.43617 | 14 | 30 | retinoid metabolic process |
| GO:00461 | 0.00185 | 3.67 | 5.14894 | 12 | 24 | polyol biosynthetic process |
| GO:00550 | 0.00186 | 3.41 | 5.79255 | 13 | 27 | cardiac muscle tissue growth |
| GO:00001 | 0.00188 | 1.38 | 99.7606 | 126 | 465 | MAPK cascade |
| GO:00608 | 0.00189 | 1.84 | 25.1011 | 39 | 117 | regulation of canonical Wnt signaling pathway |
| GO:00351 | 0.00189 | 2.10 | 16.5195 | 28 | 77 | regulation of tube size |
| GO:00508 | 0.00189 | 2.10 | 16.5195 | 28 | 77 | regulation of blood vessel size |
| GO:00420 | 0.0019 | 1.78 | 28.3191 | 43 | 132 | T cell proliferation |
| GO:00075 | 0.00192 | 1.39 | 92.6809 | 118 | 432 | hemostasis |
| GO:00025 | 0.00199 | 1.44 | 74.2305 | 97 | 346 | leukocyte differentiation |
| GO:00355 | 0.00201 | 1.19 | 382.094 | 429 | 1781 | intracellular signal transduction |
| GO:00441 | 0.00201 | 5.87 | 2.78901 | 8 | 13 | regulation of growth of symbiont in host |
| GO:00441 | 0.00201 | 5.87 | 2.78901 | 8 | 13 | negative regulation of growth of symbiont in host |
| GO:00441 | 0.00201 | 5.87 | 2.78901 | 8 | 13 | modulation of growth of symbiont involved in interaction with host |
| GO:00441 | 0.00201 | 5.87 | 2.78901 | 8 | 13 | negative regulation of growth of symbiont involved in interaction with host |
| GO:00301 | 0.00206 | 1.68 | 34.9699 | 51 | 163 | regulation of Wnt signaling pathway |
| GO:00512 | 0.00208 | 1.16 | 682.449 | 740 | 3181 | establishment of localization |
| GO:00023 | 0.00211 | Inf | 0.85816 | 4 | 4 | MHC class II protein complex assembly |
| GO:00025 | 0.00211 | Inf | 0.85816 | 4 | 4 | peptide antigen assembly with MHC class II protein complex |
| GO:00605 | 0.00211 | Inf | 0.85816 | 4 | 4 | cartilage morphogenesis |
| GO:00066 | 0.00216 | 2.19 | 14.3741 | 25 | 67 | icosanoid metabolic process |
| GO:19015 | 0.00216 | 2.19 | 14.3741 | 25 | 67 | fatty acid derivative metabolic process |
| GO:00423 | 0.00219 | 2.72 | 8.58156 | 17 | 40 | vasoconstriction |
| GO:00516 | 0.00219 | 1.18 | 440.878 | 490 | 2055 | cellular localization |
| GO:00434 | 0.00221 | 1.50 | 57.9255 | 78 | 270 | positive regulation of MAPK cascade |
| GO:00431 | 0.00223 | 18.34 | 1.28723 | 5 | 6 | surfactant homeostasis |
| GO:00520 | 0.00223 | 18.34 | 1.28723 | 5 | 6 | pathogen-associated molecular pattern dependent induction by symbiont of host innate immune response |
| GO:00521 | 0.00223 | 18.34 | 1.28723 | 5 | 6 | positive regulation by symbiont of host innate immune response |
| GO:00521 | 0.00223 | 18.34 | 1.28723 | 5 | 6 | modulation by symbiont of host innate immune response |
| GO:00521 | 0.00223 | 18.34 | 1.28723 | 5 | 6 | pathogen-associated molecular pattern dependent modulation by symbiont of host innate immune response |
| GO:00522 | 0.00223 | 18.34 | 1.28723 | 5 | 6 | pathogen-associated molecular pattern dependent induction by organism of innate immune response of other organism involved in symbiotic interaction |
| GO:00523 | 0.00223 | 18.34 | 1.28723 | 5 | 6 | positive regulation by organism of innate immune response in other organism involved in symbiotic interaction |
| GO:00523 | 0.00223 | 18.34 | 1.28723 | 5 | 6 | modulation by organism of innate immune response in other organism involved in symbiotic interaction |
| GO:00523 | 0.00223 | 18.34 | 1.28723 | 5 | 6 | pathogen-associated molecular pattern dependent modulation by organism of innate immune response in other organism involved in symbiotic interaction |
| GO:00525 | 0.00223 | 18.34 | 1.28723 | 5 | 6 | positive regulation by organism of immune response of other organism involved in symbiotic interaction |
| GO:00525 | 0.00223 | 18.34 | 1.28723 | 5 | 6 | positive regulation by symbiont of host immune response |
| GO:00609 | 0.00223 | 18.34 | 1.28723 | 5 | 6 | positive regulation of macrophage cytokine production |
| GO:00610 | 0.00223 | 18.34 | 1.28723 | 5 | 6 | negative regulation of wound healing |
| GO:00703 | 0.00223 | 18.34 | 1.28723 | 5 | 6 | response to lipoteichoic acid |
| GO:00712 | 0.00223 | 18.34 | 1.28723 | 5 | 6 | cellular response to lipoteichoic acid |
| GO:20011 | 0.00223 | 18.34 | 1.28723 | 5 | 6 | regulation of interleukin-10 secretion |
| GO:00434 | 0.00225 | 1.40 | 87.7465 | 112 | 409 | regulation of MAPK cascade |
| GO:00066 | 0.00227 | 2.01 | 18.2358 | 30 | 85 | neutral lipid metabolic process |
| GO:00550 | 0.00233 | 1.46 | 67.5798 | 89 | 315 | metal ion homeostasis |
| GO:00513 | 0.00234 | 1.26 | 199.95 | 235 | 932 | regulation of hydrolase activity |
| GO:00019 | 0.00238 | 2.31 | 12.2287 | 22 | 57 | regulation of cell-matrix adhesion |
| GO:00512 | 0.00238 | 1.59 | 43.5514 | 61 | 203 | positive regulation of lymphocyte activation |
| GO:00525 | 0.00248 | 1.51 | 54.7074 | 74 | 255 | regulation of peptidase activity |
| GO:00484 | 0.00252 | 1.22 | 272.25 | 312 | 1269 | cell development |
| GO:00315 | 0.00253 | 2.90 | 7.29433 | 15 | 34 | G2 DNA damage checkpoint |
| GO:00604 | 0.00274 | 4.08 | 4.07624 | 10 | 19 | regulation of heart growth |
| GO:00512 | 0.00276 | 1.49 | 56.6383 | 76 | 264 | negative regulation of multicellular organismal process |
| GO:00457 | 0.00277 | 1.78 | 26.3883 | 40 | 123 | regulation of angiogenesis |
| GO:00075 | 0.0028 | 3.18 | 6.00709 | 13 | 28 | embryo implantation |
| GO:00517 | 0.00281 | 1.90 | 20.8103 | 33 | 97 | interaction with host |
| GO:00362 | 0.00286 | 3.67 | 4.71986 | 11 | 22 | granulocyte activation |
| GO:00000 | 0.00287 | 3.39 | 5.36348 | 12 | 25 | regulation of transcription involved in G1/S transition of mitotic cell cycle |
| GO:00310 | 0.00287 | 3.39 | 5.36348 | 12 | 25 | chromatin remodeling at centromere |
| GO:00097 | 0.00289 | 1.25 | 204.456 | 239 | 953 | response to endogenous stimulus |
| GO:00423 | 0.00289 | 1.25 | 196.089 | 230 | 914 | regulation of phosphorylation |
| GO:00442 | 0.00294 | 1.42 | 74.2305 | 96 | 346 | small molecule biosynthetic process |
| GO:00158 | 0.00301 | 6.42 | 2.35993 | 7 | 11 | norepinephrine transport |
| GO:00443 | 0.00301 | 6.42 | 2.35993 | 7 | 11 | wound healing, spreading of cells |
| GO:00466 | 0.00301 | 6.42 | 2.35993 | 7 | 11 | regulation of gamma-delta T cell activation |
| GO:00485 | 0.00301 | 6.42 | 2.35993 | 7 | 11 | post-embryonic organ development |
| GO:00905 | 0.00301 | 6.42 | 2.35993 | 7 | 11 | epiboly involved in wound healing |
| GO:00972 | 0.00301 | 6.42 | 2.35993 | 7 | 11 | cellular response to toxic substance |
| GO:00712 | 0.00302 | 1.80 | 24.8865 | 38 | 116 | cellular response to biotic stimulus |
| GO:00000 | 0.00306 | 2.25 | 12.4433 | 22 | 58 | mitotic sister chromatid segregation |
| GO:00030 | 0.00307 | 1.56 | 44.8387 | 62 | 209 | muscle system process |
| GO:00008 | 0.00308 | 2.15 | 13.945 | 24 | 65 | sister chromatid segregation |
| GO:00027 | 0.00313 | 1.84 | 22.5266 | 35 | 105 | regulation of leukocyte mediated immunity |

| | | | | | | |
|---|---|---|---|---|---|---|
| GO:00022 | 0.00316 | 1.41 | 76.1613 | 98 | 355 | activation of immune response |
| GO:00712 | 0.00319 | 1.90 | 20.1667 | 32 | 94 | cellular response to lipopolysaccharide |
| GO:00326 | 0.00328 | 2.67 | 8.15248 | 16 | 38 | regulation of interleukin-12 production |
| GO:00447 | 0.0033 | 1.29 | 147.603 | 177 | 688 | single organism reproductive process |
| GO:00424 | 0.0033 | 2.41 | 10.2979 | 19 | 48 | odontogenesis of dentin-containing tooth |
| GO:00326 | 0.00331 | 2.03 | 16.305 | 27 | 76 | interferon-gamma production |
| GO:00610 | 0.00331 | 2.03 | 16.305 | 27 | 76 | regulation of wound healing |
| GO:00435 | 0.00335 | 1.30 | 133.014 | 161 | 620 | regulation of kinase activity |
| GO:00602 | 0.00336 | 1.36 | 95.8989 | 120 | 447 | regulation of cell development |
| GO:00009 | 0.0034 | 1.32 | 121.215 | 148 | 565 | cell morphogenesis involved in differentiation |
| GO:00329 | 0.00342 | 2.09 | 14.8032 | 25 | 69 | collagen metabolic process |
| GO:00069 | 0.00344 | 2.27 | 11.7996 | 21 | 55 | smooth muscle contraction |
| GO:00525 | 0.00347 | 1.50 | 52.7766 | 71 | 246 | regulation of endopeptidase activity |
| GO:00024 | 0.00356 | 1.84 | 21.883 | 34 | 102 | production of molecular mediator of immune response |
| GO:00108 | 0.00356 | 1.84 | 21.883 | 34 | 102 | regulation of cell-substrate adhesion |
| GO:00028 | 0.00356 | 2.76 | 7.50887 | 15 | 35 | regulation of immunoglobulin mediated immune response |
| GO:00072 | 0.00356 | 2.76 | 7.50887 | 15 | 35 | glutamate receptor signaling pathway |
| GO:00610 | 0.00357 | 1.41 | 73.8014 | 95 | 344 | muscle structure development |
| GO:00066 | 0.00359 | 1.98 | 17.1631 | 28 | 80 | triglyceride metabolic process |
| GO:00507 | 0.00371 | 1.37 | 89.8918 | 113 | 419 | positive regulation of immune response |
| GO:00442 | 0.00375 | 2.04 | 15.6613 | 26 | 73 | multicellular organismal macromolecule metabolic process |
| GO:00163 | 0.00378 | 1.19 | 323.31 | 364 | 1507 | phosphorylation |
| GO:00160 | 0.00381 | 4.89 | 3.00355 | 8 | 14 | detection of bacterium |
| GO:00711 | 0.00381 | 4.89 | 3.00355 | 8 | 14 | protein localization to chromatin |
| GO:00108 | 0.00384 | 2.86 | 6.86525 | 14 | 32 | negative regulation of cell-substrate adhesion |
| GO:00022 | 0.00386 | 2.30 | 11.156 | 20 | 52 | T cell activation involved in immune response |
| GO:00421 | 0.00386 | 2.30 | 11.156 | 20 | 52 | B cell proliferation |
| GO:00066 | 0.00386 | 1.94 | 18.0213 | 29 | 84 | acylglycerol metabolic process |
| GO:00064 | 0.00388 | 1.22 | 243.073 | 279 | 1133 | protein phosphorylation |
| GO:00315 | 0.00388 | 1.54 | 44.4096 | 61 | 207 | cell-substrate adhesion |
| GO:00512 | 0.00389 | 1.54 | 45.2677 | 62 | 211 | positive regulation of cellular component movement |
| GO:00025 | 0.00391 | 1.30 | 132.585 | 160 | 618 | immune system development |
| GO:00508 | 0.00391 | 2.19 | 12.6578 | 22 | 59 | regulation of coagulation |
| GO:00458 | 0.00392 | 1.31 | 125.291 | 152 | 584 | regulation of protein kinase activity |
| GO:00485 | 0.00392 | 1.31 | 125.291 | 152 | 584 | hematopoietic or lymphoid organ development |
| GO:19013 | 0.00393 | 1.72 | 27.6755 | 41 | 129 | regulation of vasculature development |
| GO:00019 | 0.00399 | 1.27 | 163.908 | 194 | 764 | regulation of protein phosphorylation |
| GO:00486 | 0.004 | 1.28 | 148.246 | 177 | 691 | anatomical structure formation involved in morphogenesis |
| GO:00719 | 0.00405 | 1.84 | 21.2394 | 33 | 99 | negative regulation of protein serine/threonine kinase activity |
| GO:00068 | 0.00405 | 1.24 | 197.376 | 230 | 920 | ion transport |
| GO:00027 | 0.00406 | 2.50 | 9.01064 | 17 | 42 | regulation of cytokine production involved in immune response |
| GO:00193 | 0.00406 | 2.50 | 9.01064 | 17 | 42 | arachidonic acid metabolic process |
| GO:00341 | 0.00406 | 2.50 | 9.01064 | 17 | 42 | regulation of tissue remodeling |
| GO:00000 | 0.00408 | 1.99 | 16.5195 | 27 | 77 | transition metal ion transport |
| GO:00511 | 0.00411 | 1.20 | 281.046 | 319 | 1310 | regulation of cellular component organization |
| GO:00073 | 0.00418 | 1.20 | 284.908 | 323 | 1328 | nervous system development |
| GO:00072 | 0.00421 | 1.27 | 152.108 | 181 | 709 | cell-cell signaling |
| GO:00610 | 0.00425 | 4.13 | 3.64716 | 9 | 17 | myeloid leukocyte cytokine production |
| GO:00519 | 0.00429 | 1.38 | 80.4521 | 102 | 375 | regulation of nervous system development |
| GO:00096 | 0.0043 | 3.15 | 5.57801 | 12 | 26 | response to fungus |
| GO:00440 | 0.0043 | 3.15 | 5.57801 | 12 | 26 | modification by symbiont of host morphology or physiology |
| GO:00023 | 0.00432 | 2.33 | 10.5124 | 19 | 49 | cytokine production involved in immune response |
| GO:00303 | 0.00435 | 1.54 | 42.9078 | 59 | 200 | positive regulation of cell migration |
| GO:00027 | 0.00438 | 2.12 | 13.516 | 23 | 63 | regulation of production of molecular mediator of immune response |
| GO:00442 | 0.00439 | 1.94 | 17.3777 | 28 | 81 | multicellular organismal metabolic process |
| GO:00485 | 0.00441 | 1.14 | 695.106 | 748 | 3240 | positive regulation of biological process |
| GO:00097 | 0.00442 | 1.31 | 119.284 | 145 | 556 | response to hormone |
| GO:00158 | 0.00442 | 1.79 | 22.9557 | 35 | 107 | organic hydroxy compound transport |
| GO:00325 | 0.00443 | 3.37 | 4.9344 | 11 | 23 | response to progesterone |
| GO:00433 | 0.00443 | 3.37 | 4.9344 | 11 | 23 | mast cell degranulation |
| GO:00550 | 0.00443 | 3.37 | 4.9344 | 11 | 23 | regulation of cardiac muscle tissue development |
| GO:00024 | 0.00443 | 3.67 | 4.29078 | 10 | 20 | neutrophil mediated immunity |
| GO:00327 | 0.00443 | 3.67 | 4.29078 | 10 | 20 | positive regulation of interleukin-12 production |
| GO:00421 | 0.00443 | 3.67 | 4.29078 | 10 | 20 | neutrophil activation |
| GO:00464 | 0.00443 | 3.67 | 4.29078 | 10 | 20 | membrane lipid catabolic process |
| GO:00086 | 0.00445 | 7.34 | 1.93085 | 6 | 9 | pyrimidine-containing compound salvage |
| GO:00305 | 0.00445 | 7.34 | 1.93085 | 6 | 9 | negative regulation of bone mineralization |
| GO:00324 | 0.00445 | 7.34 | 1.93085 | 6 | 9 | detection of molecule of bacterial origin |
| GO:00425 | 0.00445 | 7.34 | 1.93085 | 6 | 9 | positive regulation of tumor necrosis factor biosynthetic process |
| GO:00430 | 0.00445 | 7.34 | 1.93085 | 6 | 9 | pyrimidine nucleoside salvage |
| GO:00433 | 0.00445 | 7.34 | 1.93085 | 6 | 9 | neutrophil degranulation |
| GO:00435 | 0.00445 | 7.34 | 1.93085 | 6 | 9 | positive regulation of DNA damage response, signal transduction by p53 class mediator |
| GO:00610 | 0.00445 | 7.34 | 1.93085 | 6 | 9 | positive regulation of myeloid leukocyte cytokine production involved in immune response |
| GO:00712 | 0.00445 | 7.34 | 1.93085 | 6 | 9 | cellular response to antibiotic |
| GO:20000 | 0.00445 | 7.34 | 1.93085 | 6 | 9 | regulation of Wnt signaling pathway, planar cell polarity pathway |
| GO:20007 | 0.00445 | 7.34 | 1.93085 | 6 | 9 | regulation of cardiac muscle cell differentiation |
| GO:00067 | 0.00448 | 2.56 | 8.36702 | 16 | 39 | terpenoid metabolic process |
| GO:00074 | 0.00449 | 1.44 | 62.0018 | 81 | 289 | sensory organ development |
| GO:00109 | 0.00459 | 1.22 | 230.629 | 265 | 1075 | regulation of cell death |
| GO:00197 | 0.00467 | 1.34 | 100.404 | 124 | 468 | cellular homeostasis |
| GO:00002 | 0.00467 | 1.34 | 95.8989 | 119 | 447 | nuclear division |
| GO:00083 | 0.00468 | 1.91 | 18.2358 | 29 | 85 | regulation of cell shape |
| GO:00104 | 0.00468 | 1.68 | 28.7482 | 42 | 134 | negative regulation of peptidase activity |
| GO:00070 | 0.00474 | 1.40 | 73.5869 | 94 | 343 | mitotic nuclear division |
| GO:00075 | 0.00476 | 1.42 | 65.6489 | 85 | 306 | heart development |
| GO:00455 | 0.00478 | 1.24 | 195.23 | 227 | 910 | regulation of cell differentiation |
| GO:00507 | 0.00479 | 1.40 | 72.7287 | 93 | 339 | regulation of neurogenesis |
| GO:00973 | 0.0048 | 1.57 | 38.8316 | 54 | 181 | response to alcohol |
| GO:00487 | 0.00482 | 1.81 | 21.4539 | 33 | 100 | cardiac muscle tissue development |
| GO:00706 | 0.00483 | 2.36 | 9.86879 | 18 | 46 | negative regulation of leukocyte proliferation |
| GO:00435 | 0.00487 | 1.55 | 41.406 | 57 | 193 | skin development |
| GO:00026 | 0.00489 | 1.50 | 48.2713 | 65 | 225 | regulation of immune effector process |
| GO:00072 | 0.0049 | 1.53 | 43.1223 | 59 | 201 | Ras protein signal transduction |
| GO:00158 | 0.00492 | 2.62 | 7.7234 | 15 | 36 | monoamine transport |
| GO:00327 | 0.00492 | 2.62 | 7.7234 | 15 | 36 | positive regulation of interferon-gamma production |
| GO:00456 | 0.00492 | 2.62 | 7.7234 | 15 | 36 | positive regulation of neuron differentiation |
| GO:00352 | 0.00497 | 2.23 | 11.3706 | 20 | 53 | segmentation |
| GO:00335 | 0.00498 | 1.95 | 16.734 | 27 | 78 | unsaturated fatty acid metabolic process |
| GO:00516 | 0.00509 | 1.17 | 384.025 | 426 | 1790 | establishment of localization in cell |
| GO:00343 | 0.00511 | 1.62 | 33.039 | 47 | 154 | cell junction assembly |
| GO:00022 | 0.00523 | 1.84 | 19.9521 | 31 | 93 | lymphocyte activation involved in immune response |
| GO:00109 | 0.00531 | 1.68 | 28.1046 | 41 | 131 | negative regulation of endopeptidase activity |
| GO:00430 | 0.00537 | 1.40 | 71.227 | 91 | 332 | positive regulation of programmed cell death |
| GO:00970 | 0.00537 | 2.71 | 7.07979 | 14 | 33 | regulation of plasma lipoprotein particle levels |
| GO:00329 | 0.00539 | 2.40 | 9.22518 | 17 | 43 | negative regulation of mononuclear cell proliferation |
| GO:00506 | 0.00539 | 2.40 | 9.22518 | 17 | 43 | negative regulation of lymphocyte proliferation |
| GO:00028 | 0.00548 | 1.81 | 20.8103 | 32 | 97 | regulation of adaptive immune response |
| GO:00400 | 0.00549 | 1.50 | 46.7695 | 63 | 218 | positive regulation of locomotion |
| GO:00714 | 0.00583 | 1.26 | 152.323 | 180 | 710 | cellular response to endogenous stimulus |
| GO:00430 | 0.00583 | 1.22 | 222.262 | 255 | 1036 | regulation of programmed cell death |
| GO:00329 | 0.00587 | 1.24 | 188.58 | 219 | 879 | cellular component morphogenesis |
| GO:00017 | 0.00589 | 5.14 | 2.57447 | 7 | 12 | microglial cell activation |
| GO:00027 | 0.00589 | 5.14 | 2.57447 | 7 | 12 | positive regulation of B cell mediated immunity |
| GO:00028 | 0.00589 | 5.14 | 2.57447 | 7 | 12 | positive regulation of immunoglobulin mediated immune response |
| GO:00454 | 0.00589 | 5.14 | 2.57447 | 7 | 12 | regulation of interleukin-6 biosynthetic process |
| GO:00521 | 0.00589 | 5.14 | 2.57447 | 7 | 12 | response to defenses of other organism involved in symbiotic interaction |
| GO:00522 | 0.00589 | 5.14 | 2.57447 | 7 | 12 | response to host defenses |
| GO:00751 | 0.00589 | 5.14 | 2.57447 | 7 | 12 | response to host |
| GO:00905 | 0.00589 | 5.14 | 2.57447 | 7 | 12 | epiboly |
| GO:00550 | 0.00594 | 1.76 | 22.5266 | 34 | 105 | transition metal ion homeostasis |
| GO:00217 | 0.00596 | 1.58 | 34.9699 | 49 | 163 | developmental maturation |
| GO:00421 | 0.00597 | 2.01 | 14.5887 | 24 | 68 | positive regulation of T cell proliferation |
| GO:00900 | 0.00597 | 2.01 | 14.5887 | 24 | 68 | negative regulation of canonical Wnt signaling pathway |
| GO:00301 | 0.006 | 1.47 | 51.2748 | 68 | 239 | regulation of cell adhesion |
| GO:00061 | 0.00603 | 1.30 | 114.778 | 139 | 535 | regulation of nucleotide metabolic process |
| GO:00026 | 0.00611 | 1.55 | 38.4025 | 53 | 179 | negative regulation of immune system process |
| GO:00711 | 0.00611 | 1.55 | 38.4025 | 53 | 179 | DNA conformation change |
| GO:00508 | 0.00613 | 1.49 | 46.984 | 63 | 219 | regulation of T cell activation |
| GO:00026 | 0.00623 | 1.81 | 20.1667 | 31 | 94 | negative regulation of leukocyte activation |
| GO:00149 | 0.00624 | 2.94 | 5.79255 | 12 | 27 | smooth muscle cell migration |
| GO:00435 | 0.00624 | 2.94 | 5.79255 | 12 | 27 | regulation of phosphatidylinositol 3-kinase activity |
| GO:00068 | 0.00633 | 2.16 | 11.5851 | 20 | 54 | cellular iron ion homeostasis |
| GO:00485 | 0.00642 | 1.14 | 625.381 | 674 | 2915 | positive regulation of cellular process |
| GO:00300 | 0.00643 | 1.44 | 55.7801 | 73 | 260 | myeloid cell differentiation |
| GO:00017 | 0.00643 | 9.17 | 1.50177 | 5 | 7 | natural killer cell proliferation |

| | | | | | | |
|---|---|---|---|---|---|---|
| GO:00022 | 0.00643 | 9.17 | 1.50177 | 5 | 7 | microglial cell activation involved in immune response |
| GO:00066 | 0.00643 | 9.17 | 1.50177 | 5 | 7 | creatine metabolic process |
| GO:00520 | 0.00643 | 9.17 | 1.50177 | 5 | 7 | modulation by symbiont of host defense response |
| GO:00522 | 0.00643 | 9.17 | 1.50177 | 5 | 7 | modulation by organism of defense response of other organism involved in symbiotic interaction |
| GO:00525 | 0.00643 | 9.17 | 1.50177 | 5 | 7 | positive regulation by symbiont of host defense response |
| GO:00525 | 0.00643 | 9.17 | 1.50177 | 5 | 7 | positive regulation by organism of defense response of other organism involved in symbiotic interaction |
| GO:00525 | 0.00643 | 9.17 | 1.50177 | 5 | 7 | modulation by organism of immune response of other organism involved in symbiotic interaction |
| GO:00525 | 0.00643 | 9.17 | 1.50177 | 5 | 7 | modulation by symbiont of host immune response |
| GO:00726 | 0.00643 | 9.17 | 1.50177 | 5 | 7 | interleukin-10 secretion |
| GO:00430 | 0.00656 | 1.39 | 69.9397 | 89 | 326 | positive regulation of apoptotic process |
| GO:19005 | 0.00658 | 1.30 | 114.135 | 138 | 532 | regulation of purine nucleotide metabolic process |
| GO:00022 | 0.00661 | 3.11 | 5.14894 | 11 | 24 | mast cell activation involved in immune response |
| GO:00024 | 0.00661 | 3.11 | 5.14894 | 11 | 24 | mast cell mediated immunity |
| GO:00451 | 0.00661 | 3.11 | 5.14894 | 11 | 24 | regulation of bone resorption |
| GO:00508 | 0.00661 | 3.11 | 5.14894 | 11 | 24 | negative regulation of B cell activation |
| GO:00109 | 0.00662 | 1.37 | 75.3032 | 95 | 351 | positive regulation of cell death |
| GO:00015 | 0.00664 | 4.19 | 3.21809 | 8 | 15 | response to protozoan |
| GO:00422 | 0.00664 | 4.19 | 3.21809 | 8 | 15 | ribosomal large subunit biogenesis |
| GO:00441 | 0.00664 | 4.19 | 3.21809 | 8 | 15 | growth involved in symbiotic interaction |
| GO:00441 | 0.00664 | 4.19 | 3.21809 | 8 | 15 | growth of symbiont involved in interaction with host |
| GO:00441 | 0.00664 | 4.19 | 3.21809 | 8 | 15 | growth of symbiont in host |
| GO:00327 | 0.00677 | 1.14 | 599.422 | 647 | 2794 | RNA biosynthetic process |
| GO:00712 | 0.00678 | 2.01 | 13.945 | 23 | 65 | cellular response to metal ion |
| GO:00434 | 0.0068 | 1.48 | 48.0567 | 64 | 224 | regulation of MAP kinase activity |
| GO:00457 | 0.00685 | 3.34 | 4.50532 | 10 | 21 | respiratory burst |
| GO:00105 | 0.0069 | 3.67 | 3.8617 | 9 | 18 | regulation of vascular endothelial growth factor production |
| GO:00330 | 0.0069 | 3.67 | 3.8617 | 9 | 18 | regulation of T cell differentiation in thymus |
| GO:00507 | 0.0069 | 3.67 | 3.8617 | 9 | 18 | interleukin-1 beta secretion |
| GO:19004 | 0.0069 | 3.67 | 3.8617 | 9 | 18 | regulation of glutamate receptor signaling pathway |
| GO:00308 | 0.00706 | 2.31 | 9.43972 | 17 | 44 | regulation of B cell proliferation |
| GO:00447 | 0.00707 | 1.64 | 28.5337 | 41 | 133 | multi-multicellular organism process |
| GO:00429 | 0.00716 | 1.21 | 220.332 | 252 | 1027 | regulation of apoptotic process |
| GO:00063 | 0.00731 | 1.87 | 17.1631 | 27 | 80 | nucleosome assembly |
| GO:00159 | 0.00736 | 2.57 | 7.29433 | 14 | 34 | long-chain fatty acid transport |
| GO:00161 | 0.00736 | 2.57 | 7.29433 | 14 | 34 | diterpenoid metabolic process |
| GO:00466 | 0.00736 | 2.57 | 7.29433 | 14 | 34 | response to antibiotic |
| GO:00350 | 0.00763 | 1.57 | 34.5408 | 48 | 161 | regulation of Rho protein signal transduction |
| GO:20001 | 0.00772 | 1.49 | 43.9805 | 59 | 205 | positive regulation of cell motility |
| GO:00105 | 0.00783 | 1.26 | 138.592 | 164 | 646 | positive regulation of phosphorus metabolic process |
| GO:00459 | 0.00783 | 1.26 | 138.592 | 164 | 646 | positive regulation of phosphate metabolic process |
| GO:00467 | 0.00785 | 1.19 | 276.541 | 311 | 1289 | heterocycle catabolic process |
| GO:00028 | 0.00807 | 1.81 | 18.8794 | 29 | 88 | regulation of adaptive immune response based on somatic recombination of immune receptors built from immunoglobulin superfamily domains |
| GO:00484 | 0.00808 | 2.21 | 10.2979 | 18 | 48 | oogenesis |
| GO:00230 | 0.00809 | 1.44 | 52.7766 | 69 | 246 | signal release |
| GO:00614 | 0.00813 | 1.62 | 28.7482 | 41 | 134 | connective tissue development |
| GO:00068 | 0.00821 | 1.13 | 666.144 | 714 | 3105 | transport |
| GO:00343 | 0.00833 | 1.88 | 16.5195 | 26 | 77 | adherens junction organization |
| GO:00301 | 0.00834 | 1.97 | 14.1596 | 23 | 66 | positive regulation of Wnt signaling pathway |
| GO:00330 | 0.00834 | 1.97 | 14.1596 | 23 | 66 | T cell differentiation in thymus |
| GO:00301 | 0.00841 | 1.78 | 19.7376 | 30 | 92 | negative regulation of Wnt signaling pathway |
| GO:00480 | 0.00871 | 1.53 | 37.3298 | 51 | 174 | antigen processing and presentation of peptide antigen |
| GO:00018 | 0.00876 | 14.67 | 1.0727 | 4 | 5 | complement activation, lectin pathway |
| GO:00022 | 0.00876 | 14.67 | 1.0727 | 4 | 5 | T cell activation via T cell receptor contact with antigen bound to MHC molecule on antigen presenting cell |
| GO:00023 | 0.00876 | 14.67 | 1.0727 | 4 | 5 | MHC protein complex assembly |
| GO:00024 | 0.00876 | 14.67 | 1.0727 | 4 | 5 | dendritic cell antigen processing and presentation |
| GO:00025 | 0.00876 | 14.67 | 1.0727 | 4 | 5 | peptide antigen assembly with MHC protein complex |
| GO:00026 | 0.00876 | 14.67 | 1.0727 | 4 | 5 | regulation of dendritic cell antigen processing and presentation |
| GO:00326 | 0.00876 | 14.67 | 1.0727 | 4 | 5 | interleukin-23 production |
| GO:00326 | 0.00876 | 14.67 | 1.0727 | 4 | 5 | regulation of interleukin-23 production |
| GO:00328 | 0.00876 | 14.67 | 1.0727 | 4 | 5 | positive regulation of natural killer cell proliferation |
| GO:00336 | 0.00876 | 14.67 | 1.0727 | 4 | 5 | negative regulation of cell adhesion mediated by integrin |
| GO:00608 | 0.00876 | 14.67 | 1.0727 | 4 | 5 | establishment of blood-brain barrier |
| GO:00703 | 0.00876 | 14.67 | 1.0727 | 4 | 5 | response to bacterial lipopeptide |
| GO:00712 | 0.00876 | 14.67 | 1.0727 | 4 | 5 | cellular response to bacterial lipoprotein |
| GO:00712 | 0.00876 | 14.67 | 1.0727 | 4 | 5 | cellular response to bacterial lipopeptide |
| GO:00715 | 0.00876 | 14.67 | 1.0727 | 4 | 5 | dopaminergic neuron differentiation |
| GO:00903 | 0.00876 | 14.67 | 1.0727 | 4 | 5 | phagosome acidification |
| GO:00512 | 0.00877 | 1.84 | 17.3777 | 27 | 81 | negative regulation of lymphocyte activation |
| GO:00300 | 0.00879 | 1.28 | 120.571 | 144 | 562 | hemopoiesis |
| GO:00076 | 0.0088 | 1.35 | 78.7358 | 98 | 367 | behavior |
| GO:00468 | 0.00883 | 2.75 | 6.00709 | 12 | 28 | regulation of bone remodeling |
| GO:00066 | 0.00889 | 1.22 | 193.085 | 222 | 900 | lipid metabolic process |
| GO:00098 | 0.0089 | 1.23 | 166.911 | 194 | 778 | regulation of catabolic process |
| GO:00442 | 0.00893 | 1.18 | 277.184 | 311 | 1292 | cellular nitrogen compound catabolic process |
| GO:00162 | 0.00893 | 1.92 | 15.0177 | 24 | 70 | regulation of striated muscle tissue development |
| GO:00507 | 0.00905 | 2.12 | 11.156 | 19 | 52 | positive regulation of cytokine secretion |
| GO:00000 | 0.00911 | 5.50 | 2.14539 | 6 | 10 | DNA replication checkpoint |
| GO:00062 | 0.00911 | 5.50 | 2.14539 | 6 | 10 | DNA unwinding involved in DNA replication |
| GO:00109 | 0.00911 | 5.50 | 2.14539 | 6 | 10 | regulation of macrophage cytokine production |
| GO:00311 | 0.00911 | 5.50 | 2.14539 | 6 | 10 | positive regulation of microtubule polymerization |
| GO:00329 | 0.00911 | 5.50 | 2.14539 | 6 | 10 | inositol phosphate biosynthetic process |
| GO:00345 | 0.00911 | 5.50 | 2.14539 | 6 | 10 | protein localization to kinetochore |
| GO:00357 | 0.00911 | 5.50 | 2.14539 | 6 | 10 | endothelial cell chemotaxis |
| GO:00424 | 0.00911 | 5.50 | 2.14539 | 6 | 10 | gamma-delta T cell differentiation |
| GO:00518 | 0.00911 | 5.50 | 2.14539 | 6 | 10 | negative regulation of focal adhesion assembly |
| GO:00519 | 0.00911 | 5.50 | 2.14539 | 6 | 10 | regulation of attachment of spindle microtubules to kinetochore |
| GO:00701 | 0.00911 | 5.50 | 2.14539 | 6 | 10 | negative regulation of biomineral tissue development |
| GO:00970 | 0.00911 | 5.50 | 2.14539 | 6 | 10 | cellular response to thyroid hormone stimulus |
| GO:19018 | 0.00911 | 5.50 | 2.14539 | 6 | 10 | negative regulation of cell junction assembly |
| GO:19027 | 0.00911 | 5.50 | 2.14539 | 6 | 10 | negative regulation of cell cycle G2/M phase transition |
| GO:19033 | 0.00911 | 5.50 | 2.14539 | 6 | 10 | negative regulation of adherens junction organization |
| GO:00485 | 0.00915 | 1.44 | 50.4167 | 66 | 235 | response to steroid hormone |
| GO:00313 | 0.00927 | 1.24 | 151.25 | 177 | 705 | regulation of cellular catabolic process |
| GO:00328 | 0.00951 | 1.69 | 23.1702 | 34 | 108 | regulation of microtubule-based process |
| GO:00062 | 0.00955 | 2.88 | 5.36348 | 11 | 25 | pyrimidine nucleobase metabolic process |
| GO:00519 | 0.00955 | 2.88 | 5.36348 | 11 | 25 | catecholamine transport |
| GO:00447 | 0.00962 | 1.28 | 116.28 | 139 | 542 | multi-organism reproductive process |
| GO:00508 | 0.00972 | 1.55 | 34.1117 | 47 | 159 | positive regulation of T cell activation |
| GO:00650 | 0.00973 | 1.67 | 24.0284 | 35 | 112 | protein-DNA complex assembly |
| GO:00068 | 0.00978 | 1.51 | 38.4025 | 52 | 179 | lipid transport |
| GO:19013 | 0.00983 | 1.18 | 281.475 | 315 | 1312 | organic cyclic compound catabolic process |
| GO:00017 | 0.00987 | Inf | 0.64362 | 3 | 3 | type IIa hypersensitivity |
| GO:00017 | 0.00987 | Inf | 0.64362 | 3 | 3 | regulation of type IIa hypersensitivity |
| GO:00017 | 0.00987 | Inf | 0.64362 | 3 | 3 | positive regulation of type IIa hypersensitivity |
| GO:00022 | 0.00987 | Inf | 0.64362 | 3 | 3 | innate immune response activating cell surface receptor signaling pathway |
| GO:00022 | 0.00987 | Inf | 0.64362 | 3 | 3 | response to molecule of fungal origin |
| GO:00023 | 0.00987 | Inf | 0.64362 | 3 | 3 | serotonin production involved in inflammatory response |
| GO:00024 | 0.00987 | Inf | 0.64362 | 3 | 3 | serotonin secretion involved in inflammatory response |
| GO:00024 | 0.00987 | Inf | 0.64362 | 3 | 3 | type II hypersensitivity |
| GO:00025 | 0.00987 | Inf | 0.64362 | 3 | 3 | serotonin secretion by platelet |
| GO:00027 | 0.00987 | Inf | 0.64362 | 3 | 3 | cell surface pattern recognition receptor signaling pathway |
| GO:00027 | 0.00987 | Inf | 0.64362 | 3 | 3 | Fc receptor mediated inhibitory signaling pathway |
| GO:00028 | 0.00987 | Inf | 0.64362 | 3 | 3 | regulation of type II hypersensitivity |
| GO:00028 | 0.00987 | Inf | 0.64362 | 3 | 3 | positive regulation of type II hypersensitivity |
| GO:00030 | 0.00987 | Inf | 0.64362 | 3 | 3 | involuntary skeletal muscle contraction |
| GO:00066 | 0.00987 | Inf | 0.64362 | 3 | 3 | platelet activating factor biosynthetic process |
| GO:00069 | 0.00987 | Inf | 0.64362 | 3 | 3 | response to lipid hydroperoxide |
| GO:00109 | 0.00987 | Inf | 0.64362 | 3 | 3 | regulation of isomerase activity |
| GO:00109 | 0.00987 | Inf | 0.64362 | 3 | 3 | positive regulation of isomerase activity |
| GO:00215 | 0.00987 | Inf | 0.64362 | 3 | 3 | rhombomere 5 development |
| GO:00305 | 0.00987 | Inf | 0.64362 | 3 | 3 | embryonic genitalia morphogenesis |
| GO:00311 | 0.00987 | Inf | 0.64362 | 3 | 3 | rRNA 3'-end processing |
| GO:00329 | 0.00987 | Inf | 0.64362 | 3 | 3 | regulation of muscle filament sliding |
| GO:00341 | 0.00987 | Inf | 0.64362 | 3 | 3 | toll-like receptor 7 signaling pathway |
| GO:00482 | 0.00987 | Inf | 0.64362 | 3 | 3 | negative regulation of isotype switching to IgE isotypes |
| GO:00486 | 0.00987 | Inf | 0.64362 | 3 | 3 | embryonic hindgut morphogenesis |
| GO:00514 | 0.00987 | Inf | 0.64362 | 3 | 3 | response to cortisol |
| GO:00604 | 0.00987 | Inf | 0.64362 | 3 | 3 | positive regulation of penile erection |
| GO:00604 | 0.00987 | Inf | 0.64362 | 3 | 3 | positive regulation of heart growth |
| GO:00605 | 0.00987 | Inf | 0.64362 | 3 | 3 | intestinal epithelial cell maturation |
| GO:00703 | 0.00987 | Inf | 0.64362 | 3 | 3 | detection of bacterial lipopeptide |
| GO:19022 | 0.00987 | Inf | 0.64362 | 3 | 3 | regulation of delayed rectifier potassium channel activity |
| GO:20006 | 0.00987 | Inf | 0.64362 | 3 | 3 | regulation of histone H3-K9 acetylation |
| GO:20011 | 0.00987 | Inf | 0.64362 | 3 | 3 | regulation of T cell activation via T cell receptor contact with antigen bound to MHC molecule on antigen presenting cell |
| GO:00345 | 0.00988 | 2.45 | 7.50887 | 14 | 35 | centromere complex assembly |

| | | | | | | |
|---|---|---|---|---|---|---|
| GO:00508 | 0.00988 | 2.45 | 7.50887 | 14 | 35 | neuromuscular process controlling balance |
| GO:00509 | 0.00996 | 2.04 | 12.0142 | 20 | 56 | neuromuscular process |
| GO:00071 | 0.00997 | 1.23 | 161.762 | 188 | 754 | enzyme linked receptor protein signaling pathway |

| GOCCID | Pvalue | OddsRatio | ExpCount | Count | Size | Term |
|---|---|---|---|---|---|---|
| GO:00226 | 1.19E-25 | 9.988475 | 19.36542 | 66 | 91 | cytosolic ribosome |
| GO:00226 | 1.49E-17 | 13.29512 | 10.64034 | 39 | 50 | cytosolic large ribosomal subunit |
| GO:00443 | 2.12E-15 | 4.199675 | 27.87769 | 69 | 131 | ribosomal subunit |
| GO:00719 | 3.68E-14 | 1.4517 | 632.2489 | 781 | 2971 | cell periphery |
| GO:00058 | 7.96E-14 | 1.448673 | 616.5012 | 762 | 2897 | plasma membrane |
| GO:00159 | 2.16E-11 | 5.118873 | 15.10928 | 41 | 71 | large ribosomal subunit |
| GO:00444 | 2.77E-11 | 2.874321 | 37.87961 | 77 | 178 | cytosolic part |
| GO:00226 | 6.38E-11 | 9.162394 | 8.086658 | 27 | 38 | cytosolic small ribosomal subunit |
| GO:00306 | 4.40E-10 | 3.605769 | 21.70629 | 50 | 102 | endocytic vesicle membrane |
| GO:00444 | 7.68E-10 | 1.478122 | 313.0388 | 405 | 1471 | plasma membrane part |
| GO:00056 | 1.48E-08 | 1.653025 | 142.1549 | 202 | 668 | extracellular space |
| GO:00055 | 3.95E-08 | 1.313193 | 614.586 | 720 | 2888 | extracellular region |
| GO:00058 | 4.44E-08 | 1.568465 | 169.607 | 232 | 797 | integral component of plasma membrane |
| GO:00058 | 4.56E-08 | 2.310972 | 41.92294 | 75 | 197 | ribosome |
| GO:00312 | 4.73E-08 | 1.553449 | 177.4809 | 241 | 834 | intrinsic component of plasma membrane |
| GO:00059 | 1.25E-07 | 1.887211 | 72.1415 | 113 | 339 | focal adhesion |
| GO:00306 | 1.49E-07 | 7.858158 | 5.95859 | 19 | 28 | clathrin-coated endocytic vesicle membrane |
| GO:00059 | 2.10E-07 | 1.861907 | 72.77992 | 113 | 342 | cell-substrate adherens junction |
| GO:00057 | 2.83E-07 | 1.729454 | 94.2734 | 139 | 443 | vacuole |
| GO:00300 | 3.47E-07 | 1.837257 | 73.41834 | 113 | 345 | cell-substrate junction |
| GO:00059 | 4.33E-07 | 1.767843 | 83.42026 | 125 | 392 | adherens junction |
| GO:00426 | 4.98E-07 | 22.29057 | 2.979295 | 12 | 14 | MHC class II protein complex |
| GO:00444 | 5.75E-07 | 1.293714 | 545.4238 | 637 | 2563 | extracellular region part |
| GO:00701 | 6.15E-07 | 1.743796 | 85.54833 | 127 | 402 | anchoring junction |
| GO:00003 | 1.86E-06 | 1.706977 | 84.6971 | 124 | 398 | lytic vacuole |
| GO:00057 | 1.86E-06 | 1.706977 | 84.6971 | 124 | 398 | lysosome |
| GO:00453 | 2.07E-06 | 5.31686 | 7.23543 | 20 | 34 | clathrin-coated endocytic vesicle |
| GO:00319 | 2.24E-06 | 1.273413 | 556.0641 | 643 | 2613 | membrane-bounded vesicle |
| GO:00319 | 2.73E-06 | 1.268102 | 570.1094 | 657 | 2679 | vesicle |
| GO:00301 | 3.04E-06 | 2.118096 | 38.30522 | 65 | 180 | endocytic vesicle |
| GO:00310 | 3.66E-06 | 1.981399 | 46.17907 | 75 | 217 | extracellular matrix |
| GO:00300 | 5.69E-06 | 1.450213 | 171.7351 | 223 | 807 | cell junction |
| GO:00444 | 6.56E-06 | 1.78509 | 62.778 | 95 | 295 | vacuolar part |
| GO:00312 | 9.87E-06 | 1.805427 | 57.67064 | 88 | 271 | cell leading edge |
| GO:00159 | 1.48E-05 | 3.161363 | 12.98121 | 28 | 61 | small ribosomal subunit |
| GO:00432 | 2.35E-05 | 3.324598 | 11.27876 | 25 | 53 | lysosomal lumen |
| GO:00099 | 2.92E-05 | 1.55565 | 96.82708 | 133 | 455 | cell surface |
| GO:00325 | 3.14E-05 | 4.156818 | 7.661044 | 19 | 36 | trans-Golgi network membrane |
| GO:00057 | 3.39E-05 | 3.123207 | 12.12999 | 26 | 57 | vacuolar lumen |
| GO:00055 | 4.62E-05 | 1.971603 | 35.75154 | 58 | 168 | proteinaceous extracellular matrix |
| GO:00453 | 0.000145 | 2.571257 | 15.10928 | 29 | 71 | phagocytic vesicle |
| GO:00312 | 0.000175 | 2.224547 | 21.06787 | 37 | 99 | leading edge membrane |
| GO:00429 | 0.000225 | 1.299172 | 236.0027 | 283 | 1109 | cell projection |
| GO:00312 | 0.000348 | 1.178732 | 737.5883 | 808 | 3466 | intrinsic component of membrane |
| GO:00444 | 0.000376 | 2.42369 | 15.10928 | 28 | 71 | extracellular matrix part |
| GO:00007 | 0.000529 | 22.24247 | 1.489647 | 6 | 7 | condensed nuclear chromosome kinetochore |
| GO:00007 | 0.000618 | 6.677573 | 2.979295 | 9 | 14 | condensed nuclear chromosome, centromeric region |
| GO:00058 | 0.000622 | 2.942316 | 9.150692 | 19 | 43 | kinesin complex |
| GO:00160 | 0.000642 | 1.170144 | 723.543 | 790 | 3400 | integral component of membrane |
| GO:00432 | 0.000757 | 1.200218 | 448.5967 | 504 | 2108 | extracellular organelle |
| GO:00650 | 0.000757 | 1.200218 | 448.5967 | 504 | 2108 | extracellular membrane-bounded organelle |
| GO:00700 | 0.000757 | 1.200218 | 448.5967 | 504 | 2108 | extracellular vesicular exosome |
| GO:00055 | 0.000943 | 2.907847 | 8.725078 | 18 | 41 | collagen trimer |
| GO:00160 | 0.000963 | 1.315193 | 162.7972 | 198 | 765 | cytoplasmic membrane-bounded vesicle |
| GO:00426 | 0.001063 | 4.049125 | 4.894556 | 12 | 23 | MHC protein complex |
| GO:00300 | 0.001306 | 1.84106 | 27.02646 | 42 | 127 | lamellipodium |
| GO:00444 | 0.00143 | 1.147793 | 893.7885 | 959 | 4200 | membrane part |
| GO:00306 | 0.001497 | 1.489113 | 65.75729 | 88 | 309 | cytoplasmic vesicle membrane |
| GO:00314 | 0.001607 | 1.285558 | 177.268 | 212 | 833 | cytoplasmic vesicle |
| GO:00017 | 0.001611 | 1.80013 | 28.09049 | 43 | 132 | ruffle |
| GO:00125 | 0.001624 | 1.473404 | 68.52378 | 91 | 322 | vesicle membrane |
| GO:00427 | 0.001684 | 3.248537 | 6.384203 | 14 | 30 | presynaptic membrane |
| GO:00058 | 0.001693 | 1.839327 | 25.74962 | 40 | 121 | trans-Golgi network |
| GO:00715 | 0.001717 | 3.711321 | 5.107363 | 12 | 24 | integral component of lumenal side of endoplasmic reticulum membrane |
| GO:00985 | 0.001717 | 3.711321 | 5.107363 | 12 | 24 | lumenal side of endoplasmic reticulum membrane |
| GO:00985 | 0.001717 | 3.711321 | 5.107363 | 12 | 24 | lumenal side of membrane |
| GO:00331 | 0.001734 | 11.12011 | 1.702454 | 6 | 8 | proton-transporting V-type ATPase, V0 domain |

| | | | | | | |
|---|---|---|---|---|---|---|
| GO:00057 | 0.001782 | 1.545413 | 52.35047 | 72 | 246 | vacuolar membrane |
| GO:00974 | 0.001946 | 1.321304 | 134.9195 | 165 | 634 | neuron part |
| GO:00452 | 0.002213 | 1.42993 | 76.18483 | 99 | 358 | synapse |
| GO:00306 | 0.002253 | 2.520958 | 10.00192 | 19 | 47 | phagocytic vesicle membrane |
| GO:00430 | 0.002337 | 1.346721 | 111.9364 | 139 | 526 | neuron projection |
| GO:00057 | 0.00267 | 3.710298 | 4.681749 | 11 | 22 | multivesicular body |
| GO:00444 | 0.002689 | 1.40306 | 81.93061 | 105 | 385 | cytoplasmic vesicle part |
| GO:00056 | 0.002759 | 2.271065 | 12.34279 | 22 | 58 | basement membrane |
| GO:00353 | 0.003609 | 4.943984 | 2.979295 | 8 | 14 | microtubule plus-end |
| GO:00163 | 0.003852 | 1.731503 | 26.81365 | 40 | 126 | basolateral plasma membrane |
| GO:00325 | 0.004144 | 3.400761 | 4.894556 | 11 | 23 | neuron projection membrane |
| GO:00444 | 0.00436 | 1.480044 | 53.20169 | 71 | 250 | synapse part |
| GO:00007 | 0.004393 | 1.890938 | 18.9398 | 30 | 89 | condensed chromosome, centromeric region |
| GO:00306 | 0.004473 | 2.151096 | 12.76841 | 22 | 60 | clathrin-coated vesicle membrane |
| GO:00057 | 0.004541 | 1.510646 | 47.2431 | 64 | 222 | lysosomal membrane |
| GO:00009 | 0.005613 | 5.189755 | 2.553681 | 7 | 12 | condensed chromosome outer kinetochore |
| GO:00432 | 0.005852 | 1.547658 | 39.15645 | 54 | 184 | receptor complex |
| GO:00007 | 0.006883 | 1.85858 | 17.87577 | 28 | 84 | condensed chromosome kinetochore |
| GO:00432 | 0.007486 | 1.79889 | 19.57822 | 30 | 92 | apical junction complex |
| GO:00332 | 0.007868 | 1.858254 | 17.23735 | 27 | 81 | axon part |
| GO:00306 | 0.008745 | 5.558923 | 2.128068 | 6 | 10 | axolemma |
| GO:00301 | 0.009365 | 1.72423 | 21.49348 | 32 | 101 | clathrin-coated vesicle |

| GOMFID | Pvalue | OddsRatio | ExpCount | Count | Size | Term |
|---|---|---|---|---|---|---|
| GO:00048 | 1.37E-12 | 1.826313 | 155.1989 | 234 | 728 | receptor activity |
| GO:00037 | 3.57E-11 | 3.181061 | 30.69868 | 66 | 144 | structural constituent of ribosome |
| GO:00380 | 8.96E-09 | 1.73506 | 119.8101 | 176 | 562 | signaling receptor activity |
| GO:00048 | 1.21E-08 | 1.78915 | 103.8212 | 156 | 487 | transmembrane signaling receptor activity |
| GO:00048 | 5.96E-07 | 1.50656 | 172.2537 | 229 | 808 | signal transducer activity |
| GO:00600 | 5.96E-07 | 1.50656 | 172.2537 | 229 | 808 | molecular transducer activity |
| GO:00051 | 6.16E-06 | 1.643748 | 89.11146 | 127 | 418 | structural molecule activity |
| GO:00037 | 2.87E-05 | 1.419607 | 164.1527 | 210 | 770 | sequence-specific DNA binding transcription factor activity |
| GO:00010 | 3.14E-05 | 1.416912 | 164.3659 | 210 | 771 | nucleic acid binding transcription factor activity |
| GO:00508 | 0.000108 | 2.09596 | 26.64816 | 45 | 125 | cell adhesion molecule binding |
| GO:00422 | 0.000188 | 2.048541 | 26.43498 | 44 | 124 | peptide binding |
| GO:00508 | 0.0002 | 3.70935 | 7.2483 | 17 | 34 | extracellular matrix binding |
| GO:00332 | 0.000249 | 1.995271 | 27.5009 | 45 | 129 | amide binding |
| GO:00038 | 0.000342 | 2.754594 | 11.51201 | 23 | 54 | antigen binding |
| GO:00432 | 0.000361 | 4.940095 | 4.476891 | 12 | 21 | laminin binding |
| GO:00301 | 0.000373 | 9.870663 | 2.345038 | 8 | 11 | low-density lipoprotein particle binding |
| GO:00302 | 0.000373 | 9.870663 | 2.345038 | 8 | 11 | lipoprotein particle receptor activity |
| GO:00718 | 0.000425 | 6.172139 | 3.410965 | 10 | 16 | lipoprotein particle binding |
| GO:00718 | 0.000425 | 6.172139 | 3.410965 | 10 | 16 | protein-lipid complex binding |
| GO:00323 | 0.000474 | 12.95142 | 1.918668 | 7 | 9 | MHC class II receptor activity |
| GO:00050 | 0.000535 | 22.19587 | 1.492297 | 6 | 7 | low-density lipoprotein receptor activity |
| GO:00083 | 0.000626 | 6.663938 | 2.984594 | 9 | 14 | signaling pattern recognition receptor activity |
| GO:00381 | 0.000626 | 6.663938 | 2.984594 | 9 | 14 | pattern recognition receptor activity |
| GO:00080 | 0.000875 | 1.805058 | 31.33824 | 48 | 147 | microtubule binding |
| GO:00341 | 0.001289 | 8.633333 | 2.131853 | 7 | 10 | apolipoprotein binding |
| GO:00151 | 0.001353 | 2.781009 | 8.953783 | 18 | 42 | amino acid transmembrane transporter activity |
| GO:00051 | 0.001363 | 2.166122 | 16.20208 | 28 | 76 | integrin binding |
| GO:00192 | 0.001863 | 2.434217 | 11.29882 | 21 | 53 | kinase inhibitor activity |
| GO:00170 | 0.002062 | Inf | 0.852741 | 4 | 4 | ceramidase activity |
| GO:00433 | 0.002355 | 4.758914 | 3.410965 | 9 | 16 | proteoglycan binding |
| GO:00050 | 0.002463 | 1.857845 | 23.02401 | 36 | 108 | Ras guanyl-nucleotide exchange factor activity |
| GO:00302 | 0.002709 | 3.418568 | 5.329632 | 12 | 25 | protein serine/threonine kinase inhibitor activity |
| GO:00037 | 0.002737 | 2.392375 | 10.87245 | 20 | 51 | RNA polymerase II distal enhancer sequence-specific DNA binding transcription factor activity |
| GO:00048 | 0.002737 | 2.392375 | 10.87245 | 20 | 51 | protein kinase inhibitor activity |
| GO:00055 | 0.0028 | 2.62544 | 8.740597 | 17 | 41 | collagen binding |
| GO:00050 | 0.002818 | 2.171796 | 13.85704 | 24 | 65 | Rho guanyl-nucleotide exchange factor activity |
| GO:00198 | 0.003503 | 1.957242 | 17.90757 | 29 | 84 | growth factor binding |
| GO:00055 | 0.004064 | 4.163594 | 3.62415 | 9 | 17 | 1-phosphatidylinositol binding |
| GO:00306 | 0.004123 | 1.782941 | 23.66357 | 36 | 111 | protein binding, bridging |
| GO:00426 | 0.004202 | 3.393823 | 4.903262 | 11 | 23 | peptide antigen binding |
| GO:00165 | 0.004222 | 3.701662 | 4.263706 | 10 | 20 | cyclin-dependent protein serine/threonine kinase regulator activity |
| GO:00507 | 0.004303 | 7.397005 | 1.918668 | 6 | 9 | RAGE receptor binding |
| GO:00428 | 0.004471 | 1.25147 | 178.2229 | 209 | 836 | identical protein binding |
| GO:00051 | 0.004529 | 1.803684 | 22.17127 | 34 | 104 | cytokine activity |
| GO:00380 | 0.005035 | 2.422952 | 9.166968 | 17 | 43 | cargo receptor activity |
| GO:00015 | 0.005918 | 2.96211 | 5.756003 | 12 | 27 | beta-amyloid binding |
| GO:00600 | 0.00598 | 1.70049 | 25.79542 | 38 | 121 | binding, bridging |
| GO:00048 | 0.006246 | 9.243528 | 1.492297 | 5 | 7 | triglyceride lipase activity |
| GO:00351 | 0.006371 | 4.228433 | 3.197779 | 8 | 15 | histone kinase activity |
| GO:00037 | 0.007184 | 1.749918 | 21.95809 | 33 | 103 | motor activity |
| GO:00055 | 0.008191 | 1.746469 | 21.31853 | 32 | 100 | glycosaminoglycan binding |
| GO:00053 | 0.008243 | 1.934667 | 14.92297 | 24 | 70 | organic acid transmembrane transporter activity |
| GO:00151 | 0.008379 | 2.776673 | 5.969188 | 12 | 28 | L-amino acid transmembrane transporter activity |
| GO:00019 | 0.008441 | 2.133676 | 11.08564 | 19 | 52 | glycoprotein binding |
| GO:00018 | 0.008552 | 14.78528 | 1.065926 | 4 | 5 | lipopolysaccharide receptor activity |
| GO:00197 | 0.008552 | 14.78528 | 1.065926 | 4 | 5 | immunoglobulin receptor activity |
| GO:00336 | 0.008552 | 14.78528 | 1.065926 | 4 | 5 | receptor serine/threonine kinase binding |
| GO:00431 | 0.008552 | 14.78528 | 1.065926 | 4 | 5 | single-stranded DNA-dependent ATPase activity |
| GO:00706 | 0.008552 | 14.78528 | 1.065926 | 4 | 5 | transmembrane receptor protein serine/threonine kinase binding |
| GO:00051 | 0.009335 | 2.468805 | 7.461485 | 14 | 35 | Rho GTPase activator activity |
| GO:00049 | 0.00968 | Inf | 0.639556 | 3 | 3 | anaphylatoxin receptor activity |
| GO:00049 | 0.00968 | Inf | 0.639556 | 3 | 3 | N-formyl peptide receptor activity |
| GO:00052 | 0.00968 | Inf | 0.639556 | 3 | 3 | amine transmembrane transporter activity |
| GO:00199 | 0.00968 | Inf | 0.639556 | 3 | 3 | interleukin-1 binding |
| GO:00302 | 0.00968 | Inf | 0.639556 | 3 | 3 | very-low-density lipoprotein particle receptor activity |
| GO:00310 | 0.00968 | Inf | 0.639556 | 3 | 3 | troponin T binding |
| GO:00320 | 0.00968 | Inf | 0.639556 | 3 | 3 | clathrin light chain binding |
| GO:00356 | 0.00968 | Inf | 0.639556 | 3 | 3 | AP-3 adaptor complex binding |
